# A mathematical synthesis of genetics, development, and evolution

**DOI:** 10.64898/2026.02.25.707927

**Authors:** Mauricio González-Forero

## Abstract

Mathematically integrating genetics, development, and evolution is a longstanding challenge. A prevalent approach is quantitative genetics underpinned by the infinitesimal model, which considers a simplification of development, where phenotypes are described as the sum of many independent gene contributions. While this approach enjoys predictive success under controlled conditions, it is of interest to relax the model to consider more realistic development. Here I develop general mathematical theory that integrates sexual, discrete, multilocus genetics, developmental dynamics, and evolution. This yields an exact method to describe the evolutionary dynamics of allele frequencies and linkage disequilibria in multilocus systems and the associated evolutionary dynamics of mean phenotypes constructed via arbitrarily complex developmental dynamics. The theory shows that development affects evolution under realistic genetics by shaping the fitness landscape of allele frequencies and linkage disequilibria and by constraining adaptation to an admissible evolutionary manifold where mean phenotypes and phenotype (co-)variances can be developed. I derive a first-order approximation of this exact method, which yields equations in gradient form describing change in allele frequency, linkage disequilibria, and mean phenotypes as constrained, sometimes-adaptive topographies. Both the exact and approximated equations describe long-term phenotypic and genetic evolution. I provide worked examples to illustrate the methods. The theory obtained can be interpreted as an extension of population genetics, with some similarities with quantitative genetics under the infinitesimal model but with fundamental differences. The theory may improve prediction accuracy and shed light on empirical observations that have been paradoxical under previous theory, but appear less paradoxical in this theory.

## Introduction

Lewontin (1970) identified three necessary conditions for adaptive evolution, namely, variation, selection, and inheritance. Variation depends on factors such as mutation, migration, and development. The dependence of variation on development arises since development, defined as the process that constructs the phenotype over life, constructs the phenotypic variation upon which selection acts. Mathematical theory integrating aspects of variation such as mutation and migration with selection and inheritance, particularly Mendelian genetics, is extensively developed. However, mathematical theory integrating development, selection, and inheritance is much less developed. For instance, population genetics theory often integrates genetics and selection to describe genetic evolution with a limited consideration of development, without accounting for complex developmental dynamics where multiple genes interact non-linearly to affect the construction process of multiple phenotypes. On the other hand, quantitative genetics integrates genetics and selection to describe phenotypic evolution, but assuming a simplified developmental process, where phenotypes are the sum of scaled and independent contributions of many loci plus normally distributed residuals, which entails that the (multivariate) phenotype tends to be (multivariate) normally distributed as the number of loci increases (Fisher 1918; Barton *et al*. 2017). This assumption is known as Fisher’s infinitesimal model and it underpins much of the theory of the response to selection in quantitative genetics (Walsh and Lynch 2018) (Ch. 24).

In contrast to the infinitesimal model, developmental biologists have long found that phenotypes are the product of complex interactions between genes, environment, developmental history, and developmental stochasticity (Barresi and Gilbert 2020). Despite this discrepancy, the infinitesimal model is commonly justified by its ability to yield satisfactory predictions for short term evolution, particularly under controlled conditions possible in animal and plant breeding (Walsh and Lynch 2018). However, the infinitesimal model also makes predictions both in short and long term evolution that are inconsistent with data, including regarding the maintenance of genetic variation (Bürger 2000; Walsh and Lynch 2018; Charlesworth 2026), the evolutionary stasis of wild populations despite persistent selection and genetic variation (Kingsolver and Diamond 2011; Kirkpatrick 2009; Merilä *et al*. 2001), the rarity of stabilising selection (Kingsolver *et al*. 2001; Kingsolver and Diamond 2011), and the long term predictive ability of genetic variation (Tsuboi *et al*. 2024). Thus, both the infinitesimal model’s simplified description of development and its incorrect predictions in various situations indicate that it may be worthwhile to relax the infinitesimal model to understand phenotypic evolution (Nelson *et al*. 2013).

The simplification of development used in quantitative genetics with the infinitesimal model has provided a basis to analyse how development and selection interact. Specifically, the infinitesimal model and another assumption regarding the ability of breeding value to predict offspring phenotype imply the Lande equation, which partitions the response to selection into selection and heritable covariation, respectively described by the selection gradient and the **G** matrix of additive genetic covariation (Lande 1979; Lande and Arnold 1983). As development affects the generation of phenotypic variation that selection acts on, the Lande equation entails that development affects adaptive phenotypic evolution by affecting additive genetic covariation (Cheverud 1984), for instance, biasing evolution to proceed along “genetic lines of least resistance”, which are directions of highest additive genetic variation (Schluter 1996). Under the common assumptions that there is a single mean fitness peak and that the **G** matrix is invertible, the latter of which entails that there is genetic variation in all directions of phenotype space, this biasing of variation by development has a transient effect as in the long run evolutionary trajectories converge to the local mean fitness peak regardless of how biased the trajectory is and regardless of ancestral conditions as illustrated below. Empirical work has sought to determine whether genetic variation is absent in some directions of phenotype space, but this requires establishing that **G** has eigenvalues that are exactly zero, which is statistically infeasible (Kirkpatrick 2009).

Long standing efforts have sought to enhance mathematical evolutionary theory to further integrate aspects of development and selection. These include Alberch *et al*.’s (1979) description of development and evolution as the coupling of developmental dynamics of phenotypes with evolution as change in parameters modulating these developmental dynamics, Wagner’s (1984) analysis of the biased distribution of additive genetic covariation due to pleiotropy, Slatkin’s (1987) model integrating developmental dynamics of a phenotype with evolution of additive genetic parameters modulating such developmental dynamics, Rice’s (2002) tensor decomposition of a phenotype that is an arbitrarily complex function of underlying factors such as genes or environment, whose evolution leads to phenotypic evolution, and Parvinen *et al*.’s (2013) formulation of a phenotype’s developmental dynamics modulated by a genetically controlled, age-dependent trait under simplified genetics. A variety of other approaches have been formulated including those of Atchley and Hall (1991), Hansen and Wagner (2001), and Morrissey (2015). However, a common theme is a lack of mathematical theory simultaneously describing arbitrarily complex developmental dynamics, allele frequency change in multiple loci, and their evolution due to selection. That is, there is a lack of mathematical theory synthesising genetics, development, and evolution by selection.

Accordingly, there is a persistent disconnect in the mathematical tools to describe phenotypic and genetic evolution. On the one hand, equations describing phenotypic evolution, typically under the infinitesimal model, depend on additive genetic variances and covariances, which depend on allele frequencies whose change is not simultaneously considered (e.g., Service and Rose 1985; Barton and Turelli 1987; Turelli 1988). Rather than describing allele frequency changes jointly with phenotypic evolution, various approaches are commonly taken to avoid the need of describing allele frequency change, often motivated by the difficulty of knowing the genetic basis of complex phenotypes. These include following the evolution of additive genetic covariances with equations that do not describe changes in allele frequencies or linkage disequilibrium (e.g., Lande and Arnold 1983; Barton and Turelli 1987; Phillips and Arnold 1989; Arnold 1992; Gavrilets and Hastings 1994; Carter *et al*. 2005; Débarre *et al*. 2014; Mullon and Lehmann 2019) or assuming that additive genetic covariances change slowly relative to changes in mean phenotypes (Barton *et al*. 2017; Hill 2017; Walsh and Lynch 2018). Overall, phenotypic evolution is typically studied without considering genetic evolution and is thus confined to short-term evolution where genetic evolution remains small (Walsh and Lynch 2018).

On the other hand, mathematical descriptions of genetic evolution often consider allele frequency change for either a few loci with possibly non-additive effects on the phenotype (Nagylaki 1992; Ewens 2004), with many loci with additive effects on the phenotype (Barton and Turelli 1987), or with many loci but with a limited attempt to describe the evolution of the mean phenotypes (Barton and Turelli 1991; Felsenstein 2019). Consequently, mathematical theory describing genetic evolution does not typically consider the complexities of the developmental dynamics where many genes interact non-linearly to affect the construction process of multiple phenotypes.

This disconnect, where either phenotypic or genetic evolution is mathematically described but not both, may limit evolutionary understanding. For instance, the field of evolutionary developmental biology, or evo-devo, seeks to understand the interplay of development and evolution, focusing on two fundamental questions: how development evolves and how development affects evolution (Nuño de la Rosa and Müller 2021; Pavličev 2021). While the former question has seen much progress, the latter has been more challenging, arguably in part due to the mathematical disconnect between phenotypic and genetic evolution. Indeed, development gives rise to the genotype-phenotype map, the mathematical relationship between genotype and phenotype, but such disconnect entails that mathematical treatments of phenotypic and genetic evolution have focused on the evolution of separate ends of the genotype-phenotype map, potentially limiting their ability to see the evolutionary effects of the map itself.

Recent work formulated a mathematical synthesis of development, evolution, and simplified genetics (González-Forero 2024b). Specifically, that work formulated mathematical theory that simultaneously describes genetic and phenotypic evolution, for arbitrarily many phenotypes constructed via arbitrarily complex developmental dynamics influenced by arbitrarily many loci, but assuming simplified genetics, specifically, clonal reproduction and a continuum of alleles under rare and weak mutation. This theory found that development has two fundamental evolutionary effects: it co-shapes with selection the fitness landscape of the genes and it imposes necessary constraints on adaptation as evolution can only take place on a region (manifold) where development can produce mean phenotypes (González-Forero 2023). These two effects mean that development co-defines evolutionary outcomes with selection, rather than only influencing the evolutionary trajectories via genetic correlations as suggested by quantitative genetics under the infinitesimal model, according to which phenotypic variation can be produced anywhere on the phenotypic space. These conclusions were arrived at by deriving an equation that describes simplified genetic evolution of arbitrarily many loci by means of a discrete-time function-valued canonical equation of adaptive dynamics (Dieckmann and Law 1996) and by subjecting such an equation to a constraint in the form of a general dynamic equation that describes the arbitrarily complex development of a multivariate phenotype. However, there remains a lack of general mathematical theory that synthesises genetics, development, and evolution, specifically by integrating genetic and phenotypic evolution under sexual, discrete, and multilocus genetics and arbitrarily complex developmental dynamics. Such a theory would enable assessing whether the conclusions of how development affects evolution arrived at with simplified genetics hold with more realistic genetics. It would also provide a theory of phenotypic evolution that moves beyond the infinitesimal model.

Here I formulate general mathematical theory that integrates genetic and phenotypic evolution under arbitrary, realistic genetics and arbitrarily complex developmental dynamics. By “general”, I mean that the theory makes relatively few assumptions and so it applies to a broad class of data and models. Specifically, I allow for sexual reproduction, arbitrary numbers of loci and of discrete alleles, and arbitrary, evolving linkage disequilibria, where genes affect the arbitrarily complex, evolving developmental dynamics of arbitrarily many phenotypes, allowing for complexities such as recombination, epistasis, and pleiotropy in arbitrarily many loci. Despite allowing for such complexities, the resulting mathematical theory is highly tractable, both analytically and numerically. I follow an analogous strategy to that outlined above, where I derive general equations describing realistic genetic evolution for arbitrarily many loci and subject such equations to a constraint describing general developmental dynamics.

Results recover the two fundamental evolutionary effects of development as previously found with simplified genetics (González-Forero 2023). First, development determines the admissible region (manifold) on the fitness landscape where evolution can take place, that is, where mean phenotypes can be developed. Second, development co-shapes with selection the fitness landscape from the perspective of the genes. These two effects are of minor importance under the infinitesimal model where phenotypes are the sum of effects of many genes as in that case all phenotypes can be developed so the manifold covers the entire phenotype space and the co-shaping of the genetic fitness landscape is immaterial. However, the two effects are substantially more important when phenotypes are more complex. Results also provide mathematical theory describing both phenotypic and genetic evolution under complex development, without assuming the infinitesimal model. Thus, results provide methods to analyse data regarding phenotypic and genetic evolution under milder assumptions than those made by standard population and quantitative genetics.

The paper has the following structure, moving from more general sections involving fewer assumptions to more particular sections involving more assumptions. First, I derive general equations, termed generator equations, that describe the change in a multivariate mean trait in terms of what is commonly called the selection gradient. These generator equations are as general as the Price (1970) equation in multivariate form and generalise the Lande equation (Lande 1979; Lande and Arnold 1983) to allow for arbitrary trait distributions and any form of inheritance, including non-additive genetic inheritance, transmission bias (e.g., due to recombination and linkage disequilibrium), and non-genetic inheritance. These generator equations have wide biological and non-biological applicability and yield various insights, including regarding how arbitrary forms of inheritance may impact evolution, such as by possibly inducing maladaptation by selection. I call these equations “generator” because they enable one to generate, or derive, dynamically sufficient equations in gradient form that describe long-term multivariate evolution as an adaptive topography, where mean fitness increases over generations.

Second, I apply the generator equations to describe genetic evolution and subject these equations to a constraint describing development, thus describing the genetic evolutionary and phenotypic developmental (evo-devo) dynamics. This yields a general and exact description of the evo-devo dynamics. This description presents a principle of constrained evolution by selection, where evolution is constrained to occur within an admissible manifold determined by development. However, this general and exact description is dynamically insufficient in that it depends on variables whose evolution are not described, a feature shared with the Price and Lande equations. Yet, the general and exact description yields a method to derive exact and dynamically sufficient equations for the evo-devo dynamics under realistic genetics once additional assumptions are made, as illustrated in the examples.

Third, to describe phenotypic evolution as the climbing of a fitness landscape, I formulate a first-order approximation of the evo-devo dynamics assuming small allelic variation and weak selection. This obtains general, approximated equations in gradient form that describe genetic and phenotypic evolution as a modified, constrained, sometimes-adaptive topography, where development necessarily imposes hard evolutionary constraints and where mean fitness may increase but also decrease due to complexities in parent-offspring covariation disallowed by the infinitesimal model. Similarly to the Lande equation, these approximated equations describe selection response as being partitioned into selection and covariation where phenotypes are built under arbitrarily complex developmental dynamics. In doing so, these approximations yield notions resembling those of quantitative genetics, such as “mechanistic” additive genetic (MAG) covariances, but with important differences including that they are not within-generation variances, but parent-offspring covariances and so could in principle be negative or asymmetric. I illustrate that although the approximations give roughly correct results in some cases, the approximations can give substantially incorrect results in other cases.

Fourth, I provide worked examples that illustrate the methods, specifically, by obtaining dynamically sufficient equations for the evo-devo dynamics by making additional assumptions. These examples include the derivation of an exact equation in gradient form describing changes in allele frequency and linkage disequilibrium for two bi-allelic loci, which seems to have been previously unavailable.

Fifth, I end by discussing this work including its relationship to previous theory, limitations, how it could be applied to data, and possible future applications.

## Materials and methods

### Generator equations

In this section, I use a multivariate version of the Price (1970) equation to derive two general equations that I refer to as the primary and secondary generator equations, which describe the change of a mean vector between sets (e.g., parent and offspring generations) in terms of what is typically called the selection gradient. The derivation of each of these generator equations applies up to three linear regressions to the multivariate Price equation without loss of generality and so the equations are as general as the multivariate Price equation. The key part of the primary generator equation was essentially obtained by Rice (2004b) (his Eq. 7.13) who assumed normal distributions and so the infinitesimal model, which we do not. This key part is a generalisation of a previous univariate version (Frank 1997) and of the Lande equation (Lande 1979; Lande and Arnold 1983).

We begin by setting up definitions and assumptions of the multivariate Price equation, briefly derived in the Supplementary Information (SI) section S1 following Frank (1997). Consider two sets, referred to as parent and offspring sets. Each member of the parent set is described by an array of values listed in a column vector 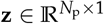, where *N*_p_ is the number of properties considered (e.g., the *i*-th entry of **z** may be each individual’s body size at a certain age, or it may be the discrete-valued gene content at a given locus). Thus, **z** may describe the phenotype or genotype, or anything else; in particular, its entries need not be quantitative traits, and may be meristic traits for instance. Similarly, each member of the offspring set is described by an array of values listed in a vector 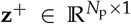, for the same properties considered. The vectors **z** and **z**^+^ are thus multivariate random variables that may be combinations of discrete or continuous values but in the same probability space (I use the common abuse of notation of denoting a random variable and its realisation with the same symbol). We assume, as the Price equation does, that the first and second moments of the random variables we consider exist (e.g., their distribution is not Cauchy, which is sometimes used in evolutionary modelling; Gupte *et al*. 2023). We do not make any other assumption regarding the genetic basis of these variables or their distribution, or whether they pertain to a biological population. Now, let each offspring be somehow linked to at least one parent (e.g., via biological parent-offspring relationships). Let *p*(**z, z**^+^) be the joint probability distribution of **z** and **z**^+^. The marginal distribution of **z** is *p*_**z**_ (**z**) ≡ ∫ *p*(**z, z**^+^)d**z**^+^ (this multiple integral can be replaced by sums or combinations of integrals and sums depending on whether each dimension of the probability space is discrete or continuous). The marginal distribution of 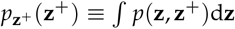. Let **z**^*′*^ be the conditional expectation of the offspring values **z**^+^ given that at least one of their parents has values 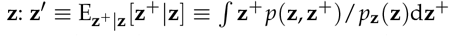. By the definition of conditional expectation, the expected offspring value **z**^*′*^ is a random variable that is a function of the parent value **z**, so we sometimes denote this by **z**^*′*^ (**z**). Let *p*_**z**_*′* (**z**) be the distribution of **z**^*′*^. Change in the phenotype due to transmission is Δ**z** = **z**^*′*^ − **z**. Define relative fitness as 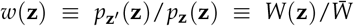, which implicitly defines absolute fitness *W*(**z**), and where mean absolute fitness is 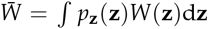. This fitness notion of the Price equation is general without regard to what causes differences in fitness, so selection is here a generalised notion that includes both standard selection and drift (Rice 2004b Ch. 6).

The change in mean vector from the parental set to the offspring set is then given by the multivariate Price equation in its standard form (SI section S1):

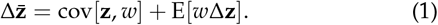

Throughout, E and cov without subscripts denote expectation and covariance over the parent distribution *p*_**z**_ (**z**), so for instance E[**z**^*′*^] = ∫ *p*_**z**_ (**z**)**z**^*′*^ (**z**)d**z** is not the average vector in the offspring population (Frank 1997), which is instead 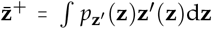. The symbols cov[**x, y**] denote a matrix, called the cross-covariance matrix of the column vectors **x** and **y** of possibly different dimension, whose *ij*-th entry is the covariance between the *i*-th entry of **x** and the *j*-th entry of **y**, namely, cov[*x*_*i*_, *y*_*j*_] (hence, cov[**x, y**] is not symmetric in general and its transpose is cov[**x, y**]^⊺^ = cov[**y, x**]); in particular, the matrix cov[**x, x**] is called the covariance (or variance-covariance) matrix of **x**, whose diagonal lists the variances of the entries of **x**.

We now rewrite the multivariate Price equation in a useful form, also following Frank (1997). The two terms in the Price equation above are traditionally interpreted as describing selection and transmission bias respectively. The first term is the selection differential:

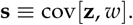

However, the second term depends on fitness so it incorporates aspects of selection. To isolate further selection from transmission bias, the Price equation can be rearranged to take the form (SI section S1)

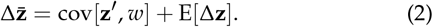

This form more neatly separates change in mean vector into the total response to selection in the first term and pure transmission bias in the second term, as the latter does not depend on fitness. The usefulness of the univariate version of (2) has been noted by various authors (e.g., Frank 1997; Rice 2004b; Heywood 2005).

We now partition the total selection response in the Price equation (2) into selection and transmission. Following Frank (1997) and without loss of generality, consider the linear regression of expected offspring values on parent values:

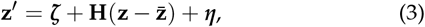

where 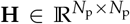 is the matrix of regression coefficients of 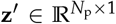 on 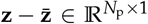, ***ζ*** is a vector of intercepts, and 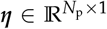 is the associated vector of residuals, with ***ζ*** and **H** estimated via least squares. The intercepts are to be simultaneously estimated with the slopes via least squares so that the expected value of residuals is zero. The predictor variables are 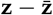 rather than **z** so that the expected value of Δ**z** does not depend on **H**. Hence, E[**z**^*′*^] = ***ζ***, using E[***η***] = **0**, which is guaranteed because the intercepts ***ζ*** are estimated via least squares. The regression (3) is without loss of generality because of least squares; specifically, the regression does not assume linearity, as non-linear effects are in the residuals and we do not require that the linear regression is a good fit. Since **z**^*′*^ is the conditional expectation of offspring values **z**^+^ given parent values **z**, then perfect linear fit (***η*** = **0**) would be guaranteed by least squares if parent values **z** and offspring values **z**^+^ were jointly multivariate normal (i.e., if *p*(**z, z**^+^) were multivariate normal), as in the infinitesimal model, but we do not assume this. However, as illustrated in examples below, perfect fit emerges when **z** is gene content due to the linear nature of Mendelian inheritance.

I refer to **H** as the heredity matrix. As shown in the next section after introducing the infinitesimal model, this matrix is a generalisation of heritability to any form of inheritance (genetic or otherwise). The *ij*-th entry of this matrix, *H*_*ij*_, gives the partial regression coefficient of the *i*-th expected offspring value 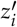 on the *j*-th parent value deviation 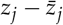. The diagonal entries *H*_*ii*_ give the “within-trait heredity” of trait *i*, whereas off-diagonal entries *H*_*ij*_ for *i* ≠ *j* give the “between-trait heredity” of trait *i* from trait *j*. Whereas univariate heritability, denoted by *h*^2^, is non-negative, both within- and between-trait heredities could be negative, as entries of **H** may be negative, since the corresponding parent-offspring regressions could in principle have negative slopes. For instance, in burying beetles, parents providing poor parental care have offspring providing better parental care (Potticary *et al*. 2024), indicating negative within-trait heredity; also in burying beetles, parents’ remaining near larvae without feeding them is negatively associated with offspring performance (Lock *et al*. 2004), indicating negative between-trait heredity. Moreover, between-trait heredity is not necessarily symmetric (*H*_*ij*_ ≠ *H*_*ji*_ for *i* ≠ *j* in general), as the matrix **H** may be asymmetric, since the regression coefficient of the *i*-th offspring trait on the *j*-th parent trait (e.g., offspring brain size on parent body size) may be different from that of the *j*-th offspring trait on the *i*-th parent trait (e.g., offspring body size on parent brain size). See also Rice 2004b (pp. 197-205) who already identified these features.

We can now write the total selection response as a multivariate breeder’s equation generalised to any form of inheritance. Using (3), the total selection response is

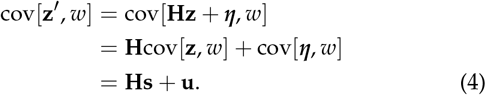

where **Hs** is the selection response from linear inheritance and **u** is the selection response from non-linear inheritance, given by

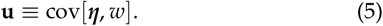

The selection response from linear inheritance **Hs** in (4) is a generalisation of the multivariate breeder’s equation (whereby selection response equals **GP**^−1^**s**; first derived in text after Eq. 7a of Lande 1979) to any form of inheritance described by **H**. Eq. (5) is also a multivariate generalisation of a general, univariate form of the breeder’s equation obtained by Heywood (2005). Interestingly, heredity **H** emerges in (4) from the transmission bias term that is often assumed to be zero (e.g., in standard equations for allele frequency change in one locus, as shown below). Indeed, in SI section S2 we show that the transmission bias in the standard form of the multivariate Price equation (1) is

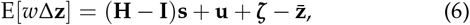

which contains a term (**H** − **I**)cov[**z**, *w*] that cancels the selection term in (1), leaving a selection response **H**cov[**z**, *w*]. Using (3), we also have that pure transmission bias is then

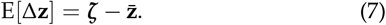

Although the multivariate breeder’s equation makes many assumptions (Lande and Arnold 1983 and Walsh and Lynch 2018), Eq. (4) does not make any assumption other than those involved in the multivariate Price equation, particularly as the regression (3) is without loss of generality.

Next, we write the total selection response under any form of inheritance in terms of what is traditionally called the selection gradient. To do this and without loss of generality, consider the linear regression of relative fitness on the vector **z**:

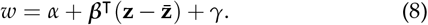

The vector 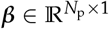 is the vector of partial regression coefficients of fitness on 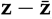, as defined by Lande and Arnold (1983). Although ***β*** is traditionally called selection gradient, I will refer to ***β*** as the selection regression coefficient and will reserve the term selection gradient for a gradient of mean fitness, which points in the direction of steepest mean fitness increase. Under some assumptions ***β*** is indeed a selection gradient, but in many cases ***β*** is not a selection gradient and so does not point in the direction of steepest fitness increase (e.g., with “frequency dependent” fitness; Walsh and Lynch 2018, p. 1313). Using (8) and the fact that residuals are uncorrelated with predictors because of least-squares, the selection differential becomes

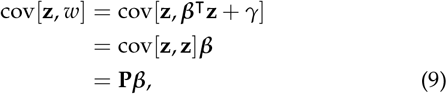

where **P** ≡ cov[**z, z**] is the covariance matrix of the vector **z**. Thus, the selection regression coefficient is ***β*** = **P**^−1^cov[**z**, *w*] if **P** is invertible (Lande and Arnold 1983). The total selection response (4) then becomes

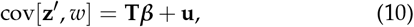

where we define the transmission matrix as

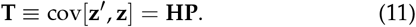

Then, the heredity matrix is **H** = **TP**^−1^ if **P** is invertible. This expression for heredity is analogous to that for multivariate heritability **GP**^−1^ (Lande 1979), with **T** in place of **G**.

As shown in the next section after introducing the infinitesimal model, the transmission matrix **T** is a generalisation of the additive genetic covariance matrix **G** to any form of inheritance. The matrix **T** is the cross-covariance matrix of expected offspring values with parent values, including not only additive genetic covariation but also non-additive genetic covariation and non-genetic covariation (e.g., via epigenetic, cytoplasmic, cultural, or ecological inheritance; Jablonka and Lamb 2014; Lala *et al*. 2024). The transmission matrix **T** was already found by Rice (2004b) (his *C* in his Eq. 7.13).

To account for non-linear inheritance, consider now the regression of parent fitness on expected offspring phenotype

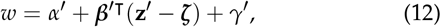

with a different regression coefficient ***β***^*′*^ of offspring traits that I will refer to as the selection pointer, and with intercept *α*^*′*^ and residual *γ*^*′*^. Using equation (12) and (3) in (5) and the fact that residuals are uncorrelated with predictors because of least-squares, the selection response from non-linear inheritance is

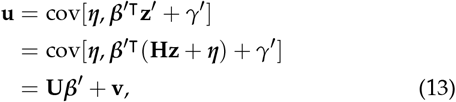

where we define the covariance matrix of parent-offspring residuals as

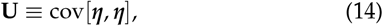

and the selection response from non-linear selection (described by *γ*^*′*^) under non-linear inheritance (described by ***η***) as

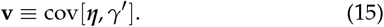

Note that analogously, from (5) and (8), the selection response from non-linear inheritance is

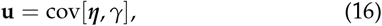

because predictors and residuals are uncorrelated by least squares. We can more compactly account for non-linear inheritance as follows. Using equation (12) and the fact that residuals are uncorrelated with predictors, the total selection response becomes

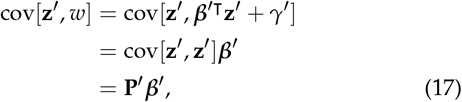

where **P**^*′*^ cov[**z**^*′*^, **z**^*′*^] is the covariance matrix of the expected offspring phenotype. Thus, ***β***^*′*^ = **P**^*′*−1^cov[**z**^*′*^, *w*] if **P**^*′*^ is invertible. Additionally, from (3), it follows that the offspring phenotypic covariance matrix is

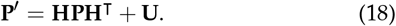

Therefore, using (2), (10), and (17), we obtain two general *generator equations* describing the change in the mean vector between the parent and offspring sets:

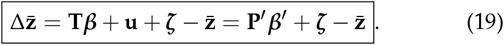

The primary (left-most) generator equation partitions total selection response into selection response from linear inheritance (**T*β***) and from non-linear inheritance (**u**). The first term in the primary generator equation describing selection response from linear inheritance was already written without complete derivation by Rice (2004b) (his Eq. 7.13) assuming normal distributions and so the infinitesimal model (whereby **u** = **0**). The primary generator equation in turn partitions selection response from linear inheritance into selection (***β***), variation (**P**), and linear inheritance (**H**) via (11), matching Lewontin’s (1970) three necessary conditions for evolution by selection, generalising previous results for univariate phenotypes under the infinitesimal model (Eq. 10 of Frank 1997) (see also Okasha 2006). The secondary (right-most) generator equation (19) describes total selection response compactly as the product between offspring variation (**P**^*′*^) and offspring selection (***β***^*′*^). This product accounts for both linear and non-linear inheritance and linear and non-linear selection. The generator equations do not make any assumptions that are not already included in the multivariate Price equation, and so are as general as that equation, applying broadly across biological and non-biological systems.

### Lande equation as a particular case

The general form of the Lande equation (i.e., 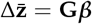; Lande and Arnold 1983) follows from the primary generator equation by making two assumptions. To see this, let us first consider the infinitesimal model. Let **z** be a phenotype and consider a vector of genotype content 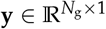, where *N*_g_ is the number of loci or genetic predictors (e.g., the *i*-th entry of **y**, *y*_*i*_, is 0, 1, or 2 depending on whether the individual’s genotype is aa, Aa, or AA in the *i*-th locus). Without loss of generality, consider the linear regression of phenotype **z** on genotype content **y**:

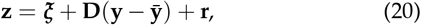

where 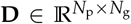 is the matrix of regression coefficients of 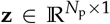 on 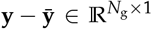, ***ξ*** is a vector of intercepts, and 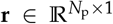 is the associated vector of residuals, with ***ξ*** and estimated simultaneously via least squares. Taking the expectation yields 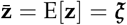. The breeding value of the phenotype **z** is defined as

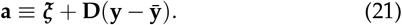

The *ij*-th entry of the matrix 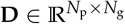 is the partial regression coefficient of the *i*-th phenotype on the *j*-th genetic predictor. Eq. (20) is the starting point of Fisher’s (1918) infinitesimal model, where the coefficients in **D** are called the additive effects of allelic substitution (Lynch and Walsh 1998, p. 72), typically denoted as *α* but we use **D** to keep a homogeneous notation where capital bold letters refer to matrices throughout and to highlight that this matrix describes development in the sense of describing the effects of genotype on phenotype. Wagner (1984) called **D** the developmental matrix (and denoted it by **B**). Thus, the additive genetic covariance matrix of the phenotype **z** is

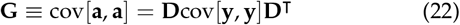

(e.g., eq. + of Wagner 1984 or eqs. 3.5 of Barton and Turelli 1987). From (22), it follows that the **G** matrix is singular if there are fewer loci than phenotypes (i.e., *N*_g_ < *N*_p_) (see also Altenberg 1995). This is because any matrix with fewer rows than columns is singular (Horn and Johnson 2013, section 0.5 second line) so if *N*_g_ < *N*_p_, then **D**^⊺^ is singular; in such a case, since for any singular matrix **B**, any well-defined product **AB** is a singular matrix (indeed, if **Bv** = **0** for **v** = **0**, then **ABv** ≠ **0** but **v** ≠ **0**), then **G** is singular.

The regression (20) can always be made, so it is done without loss of generality. However, the infinitesimal model involves additional assumptions including that the *y*_*i*_ are independent (so no linkage disequilibrium), that the effects of the genotype on the phenotype *D*_*ij*_ are of order 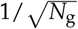, that *N*_g_ approaches infinity, and that residuals **r** are multivariate normal. By the central limit theorem, this entails that **z** and **a** are jointly and marginally multivariate normal.

We now obtain the Lande equation in general form from the primary generator equation by making two assumptions made by Lande and Arnold (1983): (I) Assume that the expected offspring vector is the expected breeding value given the parent vector: **z**^*′*^ = E[**a z**] (see text after their Eq. 6a). Then, the transmission matrix equals the **G** matrix: **T** ≡ cov[**z**^*′*^, **z**] = cov[E[**a z**], **z**] = cov[**a, a** + **r**] = cov[**a, a**] = **G**, where we use the fact that cov[**a, r**] = **0** because predictors and residuals are uncorrelated by least squares. Assumption (I) also entails that there is no pure transmission bias, 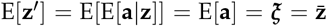. (II) Assume that the parent vector **z** and its breeding value **a** are jointly multivariate normal, for instance, by assuming the infinitesimal model. By least squares and assumption (I), assumption (II) entails that the parent-offspring regression (3) is exactly linear: ***η*** = **0**. Consequently, there is no selection response from non-linear inheritance, **u** = cov[***η***, *w*] = **0**. Hence, from these two assumptions, the primary generator equation becomes the Lande equation in general form: 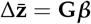.

The gradient form of the Lande (1979) equation (i.e., 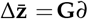 ln 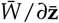) follows by making further assumptions and so is less general (SI section S4; see also Walsh and Lynch 2018 p. 1313). Specifically, if **z** is multivariate normal, then the selection regression coefficient equals the average fitness gradient, ***β*** = E[*∂w*/*∂***z**] (Eq. 9 and previous unnumbered equation of Lande and Arnold 1983). The assumption that **z** is multivariate normal would be automatically met if one assumes the infinitesimal model, but not necessarily if one only assumes that **z** and **a** are jointly multivariate normal. If, additionally, absolute fitness is independent of the mean phenotype (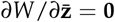 ; often described as fitness being frequency independent, Walsh and Lynch 2018 p. 1313), then the selection regression coefficient equals the selection gradient, 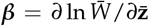. Note it is more general to use the average fitness gradient than the selection gradient, and more general still to use the selection regression coefficient. The series of assumptions made by the general and gradient forms of the Lande equation are not made by the generator equations. In particular, the assumptions (I-II) of expected offspring phenotype equalling breeding value and normal distributions are not typically met in our genetic treatment below.

Thus, the transmission matrix **T** is a generalisation of the additive genetic covariance matrix **G**: the two are equal under assumption (I), that is, if the expected offspring phenotype given parent phenotype is the expected breeding value given parent phenotype. However, assumption (I) does not always hold, as illustrated below in a simple situation of a single phenotype controlled by a single biallelic locus under random mating, where 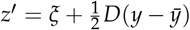 so *z*^*′*^ ≠ E[*a z*] and *T* = *G*/2 (see also Heywood 2005).

Consequently, heredity **H** is a generalisation of multivariate heritability **GP**^−1^. We saw that **H** = **TP**^−1^ if **P** is invertible, so heredity equals multivariate heritability if assumption (I) holds. Yet, for the example just mentioned where this assumption does not hold and *T* = *G*/2, we have that heredity is half heritability: *H* = *h*^2^/2. This is consistent with classic results where the regression of expected offspring phenotype on parent phenotype (our heredity) is half the regression of offspring phenotype on mid-parent phenotype, which is heritability under these assumptions (Fisher 1918, Eq. 10.134 of Nagylaki 1992, Eq. IX-56 of Felsenstein 2019). Hence, **T** and **H** are respective generalisations of **G** and **GP**^−1^ to any form of inheritance.

The generator equations allow for a wider variety of evolutionary dynamics than those described by the Lande equation. Lande (1979) proved that under his gradient equation there is a basic principle of multivariate phenotypic evolution, whereby evolution by selection proceeds as the climbing of an adaptive topography such that mean fitness cannot decrease with weak selection and invertible **G** (his Eq. 8b). Fitness cannot decrease in the Lande equation because **G** is a covariance matrix, so it is symmetric and its eigenvalues are always real and positive (under the infinitesimal model, otherwise the normality of **a** does not hold), which entails that mean (Gaussian) fitness peaks are locally stable equilibria. In contrast, **T** is not a covariance matrix, but a cross-covariance matrix as **z**^*′*^ and **z** are different in general, so it may be asymmetric (e.g., as illustrated by Rice 2004a for a plastic trait where parents and offspring have different environments; third equality in his Eq. 26). Then, some of the eigenvalues of **T** could be zero, negative, or complex, even if all the entries of **T** are non-negative. If **T** has at least one negative or complex eigenvalue, then the selection response **T*β*** alone could cause: mean fitness to decrease, mean fitness peaks to be saddle points or unstable equilibria, limit or dampened evolutionary cycles, or evolutionary chaos. Although these features are not possible in the gradient Lande equation, they are known to happen in various evolutionary models (Feldman and Cavalli-Sforza 1976; Parvinen 2005; Doebeli and Ispolatov 2014) and are allowed by the primary generator equation given the broad structure of **T**.

### Adaptability and respondability

We now define two quantities whose sign indicates whether selection responses are locally adaptive or not, in the sense of increasing mean fitness. We thus define two quantities, namely adaptability, which measures the ability of the system to adapt, and respondability, which measures the ability of the system to respond to selection. Since the selection response cannot be maladaptive under the gradient Lande equation, there is no distinction between adaptability and respondability under that equation. But since **T** might yield maladaptation, such distinction occurs here.

The response to selection from linear inheritance locally yields maladaptation if the angle between **T***∂* ln 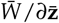 and *∂* ln is greater than 90^°^. To assess the angle between **T***∂* ln 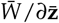 and *∂* ln 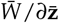, note that the space of **z** may be non-Euclidean because the entries of **z** may have different units (e.g., some in mass units and others in length units). Consider then the non-dimensionalising metric **Ĩ**, which is a diagonal matrix whose *i*-th diagonal entry is 1 divided by the squared units of the *i*-th entry of **z**. Let us then define adaptability as the (non-Euclidean) dot product between the selection response from linear inheritance under the selection gradient and the selection gradient:

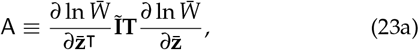

which is a non-dimensional real number. This A is a modified, non-normalised generalisation of evolvability sensu Hansen and Houle (2008) (their Eq. 1). The angle between the selection response from linear inheritance and the selection gradient is less than 90^°^ if adaptability is positive, equal to 90^°^ if adaptability is zero, or greater than 90^°^ if adaptability is negative. There is thus local maladaptation by linear selection from linear inheritance if adaptability is negative. Maladaptation by selection is not possible with the Lande equation because **G** is symmetric and its eigenvalues are all non-negative since it is a covariance matrix. Because **T** may be asymmetric but is square, the sign of adaptability is determined by whether the symmetric part of **T**, defined as 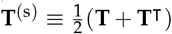, is respectively positive definite (all its eigenvalues are positive), positive semi-definite (all eigen-values are positive or zero), or indefinite (all eigenvalues are positive, zero, or negative), respectively. Thus, adaptability is necessarily positive if all the eigenvalues of **T**^(s)^ are positive, but adaptability can be negative if at least one of these eigenvalues is negative. The definition (23a) resembles some definitions of energy in physics, which measures the ability of a system to do work. Analogously, Eq. (23a) measures the ability of the system to adapt under linear inheritance.

Given that the total response to selection is given by (17), we similarly define respondability as

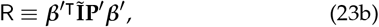

which is non-negative because **P**^*′*^ is a covariance matrix and so is positive semi-definite. Hence, the total selection response (**P**^*′*^ ***β***^*′*^) points in the direction (≤ 90^°^) of the selection pointer ***β***^*′*^, which is the reason for giving it that label. Yet, ***β***^*′*^ may not point in the direction of the selection gradient, and so it may not point in the direction of steepest increase in mean fitness. Thus, respondability does not necessarily measure the ability of the system to adapt, but the ability of the system to respond to selection, possibly maladaptively.

### How development comes in

The generator equations alone are useful to understand shortterm phenotypic evolution but they are insufficient to understand the role of development in evolution and to describe longterm phenotypic evolution. Indeed, if **z** in the generator equations is a phenotypic vector, the equations could be used to understand and predict short term evolutionary change in the mean phenotypes if its components are empirically estimated (particularly, **z, z**^+^, *p*(**z, z**^+^), and *p*_**z**_*′* (**z**), from which all other components follow, including fitness). However, the generator equations applied to phenotypes assume that the phenotypes have already been constructed, which limits the possibility of understanding the role of development on evolution. Moreover, the generator equations applied to phenotypes are generally dynamically insufficient: as examples below show, the generator equations applied to phenotypes depend on allele frequency but do not describe allele frequency change. Hence, the generator equations alone applied to phenotypes do not describe long-term phenotypic evolution as they cannot be iterated over many generations, over which allele frequencies change. This is a common feature with the Lande equation (Barton and Turelli 1987; Turelli 1988), whose iteration over generations implicitly assumes negligible allele frequency change in general, which restricts it to short-term evolution (Walsh and Lynch 2018, pp. 504, 879).

A traditional approach to address this dynamic insufficiency is not to follow allele frequency change but to treat the phenotypic or additive genetic covariance matrix as a dynamic variable itself. For comparison with this traditional approach, we derive an equation describing the unconstrained change in the covariance matrix of vector **z** allowing for any form of inheritance and transmission bias and so generalising the corresponding equation of Lande and Arnold (1983) (following the approach of Lande and Arnold (1983) and using ideas from the Price equation, but not the Price equation itself; SI section S3). We obtain that the unconstrained change from parent to offspring sets in the covariance matrix of vector **z** is

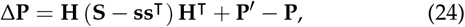

where **S** ≡ cov[**Z**, *w*] is the selection differential of the squared deviation and 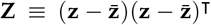 is the squared deviation of the parent vector **z** from the mean. Equation (24) assumes that there is no selection response from non-linear inheritance cov[***η***, *w*] = **0** and similarly that cov[***η*, z***w*] = **0** (e.g., if **z**^*′*^ and **z** are jointly multivariate normal, so ***η*** = **0**). If **z** is multivariate normal, then the selection differential is given in terms of the average fitness gradient, **s** = **P**E[*∂w*/*∂***z**] (SI section S4), and the selection differential of the squared deviation is given in terms of the average fitness Hessian matrix, **S** = **P**E[*∂*^2^*w*/*∂***z***∂***z**^⊺^]**P** (SI section S5) (Eq. 14b of Lande and Arnold 1983). If, additionally, absolute fitness is independent of the mean vector 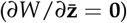 and the phenotype covariance matrix is independent of the mean vector 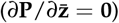, then the average fitness Hessian equals the selection Hessian, 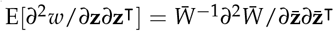 (SI section S5). Then, equation (24) recovers Lande and Arnold’s (1983) equation describing the change in phenotype covariance (their eq. 13a; see also their eq. 12) under the assumptions in this paragraph and if there is perfect heredity and no pure transmission bias (i.e., if **H** = **I** and **z**^*′*^ = **z**, so **P**^*′*^ = **P**). Our **P**^*′*^ − **P** describes the changes in phenotypic covariation resulting from reproduction and so it would include a matrix **M** used by Phillips and Arnold (1989) to describe phenotypic covariation generated by mutation, but we do not explore this here.

In this paper, to integrate genetic and phenotypic evolution, we apply the generator equations to haplotypes subject to a constraint describing development. This allows one to study the role and evolution of development, to solve the problem of dynamic insufficiency of the generator equations, yielding exact equations describing both genetic and phenotypic evolutionary change, and to describe the evolution of mean phenotypes and covariance matrices as emergent properties modulated by genetic evolution.

### Evo-devo dynamics under diploid genetics

We now outline a strategy to obtain dynamically sufficient equations describing both genetic and phenotypic evolutionary change. The key of the strategy is to apply one of the generator equations to haplotypes rather than phenotypes and to subject such haplotype generator equation to a constraint describing phenotype construction. This strategy can be implemented in two equivalent ways. On the one hand, one can use a generator equation to follow changes in allele frequencies and linkage dis-equilibria. On the other hand, one can instead and equivalently follow changes in haplotype (or gamete) frequencies as is traditional in multi locus approaches (Kimura 1956; Lewontin and Kojima 1960; Bodmer and Parsons 1962; Barton and Turelli 1991). These two implementations are mathematically equivalent, and each one has pros and cons in practice. The former takes some extra effort in setting it up, but yields overall simpler and biologically insightful expressions. We thus present the allele frequency approach in the main text and outline the haplotype frequency implementation in SI section S9.3.

We now build a vector **x** of haplotype content to study the evolution of its mean. The specific form of the vector may depend on the model, but for specificity consider the following scheme. Consider a haplotype **k** = (*k*_1_, *k*_2_, …, *k*_*n*_) ℍ, whose *i*-th entry *k*_*i*_ is the allele (i.e., some letter) present at locus *i* and ℍ is a set of possible haplotypes. Origin or loss of loci could be described by using a symbol (e.g., ∅) in the relevant entries of **k** that denotes locus absence in a particular haplotype (say of parents or sex chromosomes). Let *p*_**k**_ be the frequency of the haplotype **k** in the population. Let 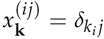 be the gene con-tent in haplotype **k** at locus *i* for allele *j*, where *δ*_*ij*_ is Kronecker’s delta (i.e., 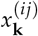 is 0 or 1 if allele *j* is absent or present at locus *i* in haplotype **k**). Thus, the frequency of allele *j* in locus *i* is the mean gene content: 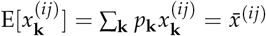(Price 1970). Moreover, the frequency of the haplotype **k** is the mean product of gene content across loci: 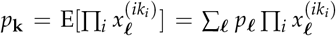 (since the product is always zero except when ***ℓ*** = **k**, in which case the product is one). Writing this expectation of a product in terms of the product of expectations yields

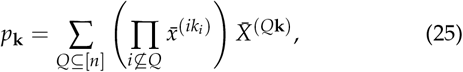

where [*n*] = {1, …, *n*} is the set of all loci in haplotype **k**, *Q* is any subset of [*n*] including the empty set and [*n*], 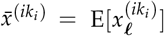 is the frequency of the allele *k*_*i*_ in locus *i*, 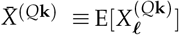 is the coefficient of linkage disequilibrium of the subset of loci *Q* in haplotype **k**, and 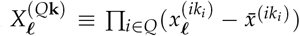 is the haplotype content deviation of haplotype ***ℓ*** relative to the subset of loci *Q* in haplotype **k** (Eq. (25) follows from the distributive property of multiplication and addition, as shown with assistance from Gemini in SI section S7). So, to describe allele frequency change, we must follow the evolution of both allele frequencies and linkage disequilibria (Kimura 1956; Lewontin and Kojima 1960; Bodmer and Parsons 1962; Barton and Turelli 1991). Then, for haplotype **k**, we build a vector of haplotype content **x**_**k**_ whose first entries are the gene content 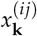 across loci (*i*) and alleles (*j*), and subsequent entries are the haplotype content deviation 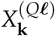 over all subsets *Q* of two or more loci (if *Q* contains no or one locus, 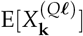 is 1 or 0, respectively, so we do not need to track these cases) and over all haplotypes ***ℓ*** that yield an undetermined 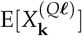 (as illustrated in example 4, for two loci, there is only one such undetermined 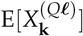 over ***ℓ***). Therefore, the mean haplotype content 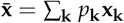 lists all allele frequencies and linkage disequilibrium coefficients across loci. Let *N*_g_ be the number of entries of 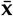 (i.e., number of alleles times number of loci times number of linkage disequilibrium coefficients). As following allele frequencies and linkage disequilibria is equivalent to following haplotype frequencies, the number of variables would be the same in the latter case.

Applying the primary generator equation (19) to haplotype content **x**_**k**_, the change in allele frequencies and linkage disequilibria is given by

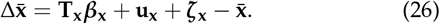

The subscript **x** is used to denote that the corresponding quantity is for haplotype content. The haplotype generator equation (26) is a first ingredient to derive dynamically sufficient equations for phenotypic evolution as they describe the underlying evolutionary dynamics of allele frequencies and linkage disequilibria. For the examples below, the residual of regressing offspring haplotypes on parent haplotypes is zero (***η***_**x**_ = **0**) due to the linearity of Mendelian inheritance. This entails that expected offspring haplotypes are exactly a linear transformation of parent haplotypes, and that for haplotypes the selection response from non-linear inheritance is zero (**u**_**x**_ = **0**).

Because ***β***_**x**_ is now the regression coefficient of fitness on haplotype content, this selection coefficient measures total genetic selection, that is, the total effects of haplotype content on fitness. Importantly, total genetic selection depends on development separately from selection. To see this, we now calculate this regression coefficient. From (8), the regression coefficient ***β***_**x**_ is defined by the linear regression of the relative fitness of haplotype **k** on haplotype content:

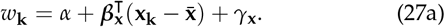

For diploids, the relative fitness of genotype **kl** is *w*_**kl**_. Without loss of generality, let the first subscript in *w*_**kl**_ refer to the maternally inherited haplotype. In general, fitness is asymmetric (*w*_**kl**_ ≠ *w*_**lk**_, e.g., due to sex chromosomes, where if a haplotype **l** in mammals includes the Y chromosome and so is not maternally inherited, *w*_**kl**_ may be positive but *w*_**lk**_ may be zero or undefined). Let *q*_**k**_ be the probability that haplotype **k** is the maternal one in a diploid genotype. Then, the relative fitness of haplotype **k** is

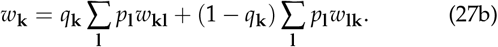

Let us now consider the linear regression of *w*_**kl**_ on phenotype and haplotype content, given by

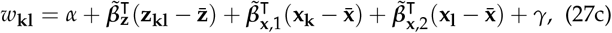

where 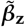 and 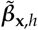 are respectively the partial regression coefficients of fitness on phenotype and haplotype content in the *h*-th haplotype of the individual. These regression coefficients are often interpreted as measuring direct selection on phenotype and haplotype, respectively (although as shown in SI section S9.2, this interpretation is problematic). In turn, the linear regression of phenotype **z**_**kl**_ on haplotype content is

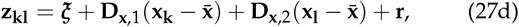

where 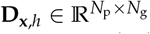 is the matrix of regression coefficients of the phenotype on the haplotype content in the *h*-th haplotype of the individual. Substituting (27d) into (27c), this into (27b), simplifying, and setting the result equal to (27a) yields the selection regression coefficient of the haplotype

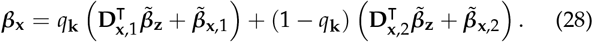

Thus, total genetic selection ***β***_**x**_ depends on development separately from selection, since **D**_**x**,*h*_ measures the effects of alleles on phenotype without measuring selection.

To describe phenotypic evolution, we subject the haplotype generator equation (26) to a constraint describing phenotype construction, that is, development. I will consider development both implicitly in terms of the genotype-phenotype map and explicitly via developmental dynamics.

Let us first consider development implicitly. That is, we consider development by subjecting genetic evolution (26) to the static developmental constraint

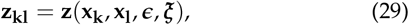

which gives the phenotype of a diploid individual with genotype **kl**. The function **z**(*·*) is the genotype-phenotype map, which depends on the genotype content **x**_**k**_ and **x**_**l**_, on the environment ***ϵ*** experienced by the individual, and on developmental stochasticity ***ξ***, where ***ϵ*** and ***ξ*** are random vectors with the former referring to variables external to the individual (e.g., temperature) and the latter to internal variables that are possibly unknown or unmeasurable. The function **z**(*·*) can be arbitrarily complex, non-linear, and discontinuous. Let *p*_**kl*ϵξ***_ be the joint probability distribution of genotype **kl**, environment ***ϵ***, and developmental stochasticity ***ξ***. Thus, the phenotype of genotype **kl** is a random variable, so not all individuals with that genotype necessarily have the same phenotype. The mean phenotype is

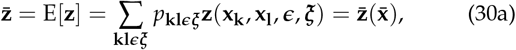

and the phenotype covariance matrix is

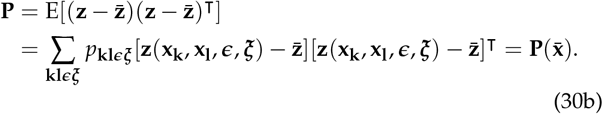

As examples below illustrate, the mean phenotype and phenotype covariance matrix are generally functions 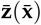 and 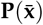 of allele frequencies because the distribution *p*_**kl*ϵξ***_ is a function of allele frequencies. I call 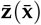 the mean genotype-phenotype map and 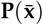 the mean genotype-covariance map. These mean maps can evolve (e.g., 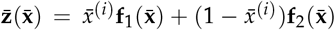 may evolve from function **f**_1_ to function **f**_2_ as the allele frequency 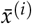 at locus *i* evolves from 1 to 0).

That these mean maps are functions of allele frequencies is a key point that contrasts with common practice under the infinitesimal model, where the mean phenotype and phenotype covariance matrix are not taken as functions of allele frequencies but as parameters that selection can modify freely. Instead, here, the mean phenotype and phenotype covariance matrix evolve as the mean haplotype content 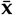 evolves. This entails that there is a hard constraint over evolution given by development: as allele frequencies change, the mean genotype-phenotype and mean genotype-covariance maps must be satisfied. This hard constraint persists as the mean maps evolve, because the newly evolved maps still need to be satisfied. If the mean genotypephenotype map evolves so that it intersects with mean fitness peaks, the constraint becomes inactive, but the hard constraint persists as the map must still be satisfied.

Alternatively, let us consider development explicitly. That is, we now consider development by subjecting genetic evolution (26) to the dynamic developmental constraint

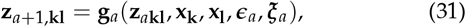

with initial condition **z**_1**kl**_ = **g**_0_(**x**_**k**_, **x**_**l**_, ***ϵ***_0_, ***ξ***_0_). This dynamic constraint gives the phenotype **z**_*a*+1,**kl**_ at age *a* + ∈ {1 2, …, *N*_a_} for genotype **kl** via a recurrence equation. The symbols ***ϵ***_*a*_ and ***ξ***_*a*_ are random vectors respectively describing the environment and developmental stochasticity experienced at age *a* ∈ {0, …, *N*_a_ − 1}. The function **g**_*a*_ for *a* ∈ { 0, …, *N*_a_} is the developmental map, which can be arbitrarily complex, non-linear, and discontinuous. The developmental dynamics (31) occur within each evolutionary step of (26). The lifetime phenotype of genotype **kl** is the concatenation of the phenotypes at each age: 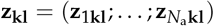, where the semicolon denotes a line break or concatenation. Thus, denoting by *N*_p_ the number of phenotypes at each age (i.e., **z**_*a***kl**_ has that many entries), the lifetime phenotype **z**_**kl**_ has *N*_p_ *N*_a_ entries. Analytically or numerically solving the recurrence (31) yields the genotype-phenotype map (29), which can be substituted into (30) to obtain the mean genotype-phenotype map and mean genotype-covariance map.

Thus, constraining the genetic evolutionary dynamics (26) with the phenotypic developmental dynamics (31) to compute phenotype mean and covariance (30) describes the evo-devo dynamics, enabling a dynamically sufficient description of the change in the phenotype mean and covariance. In the examples below, the constraining of evolution by development affects the selection regression coefficients on the haplotype ***β***_**x**_ and 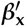. The equations describing phenotype construction, either implicitly (29) or explicitly (31), can be understood as specifying the developmental constraints, that is, the constraints that the phenotype must meet as a result of how it is constructed.

Using the developmental map has advantages over the genotype-phenotype map. Indeed, the genotype-phenotype map is the solution of the developmental map, so non-linearities in the developmental map easily make it infeasible to write the genotype-phenotype map **z**(*·*) analytically. So for realistic developmental maps, it is more tractable to operate with **g**_*a*_ than **z**(*·*). Moreover, developmental feedbacks, where phenotypes affect themselves at later ages, are not made apparent by the genotype-phenotype map but can be readily studied with the developmental map. As system of recurrence equations can describe ordinary and partial differential equations once time and space is suitably discretised, the developmental dynamics (31) can describe a variety of complex models of development, including gene regulatory networks (Alon 2020) and spatially explicit models of morphological development (Deutsch and Dormann 2017), possibly such as existing models of tooth cusps (Salazar-Ciudad and Jernvall 2010), digit number (Sheth *et al*. 2012), and leaf shapes (Runions *et al*. 2017), once such models have been arranged into the form of this equation.

### First-order approximation in gradient form

The above scheme provides an exact method to describe the evo-devo dynamics. However, it does not describe phenotypic evolution as an adaptive topography. To describe phenotypic evolution under arbitrarily complex development and realistic genetics as an adaptive topography and to gain additional analytical insight, we now obtain an approximation of the exact evo-devo dynamics by assuming small haplotype variation and weak selection. This approximation yields equations describing genetic and phenotypic evolution in a constrained gradient form and so implying the climbing of a fitness landscape under some situations, with some similarity to the gradient Lande equation, but with important differences as the infinitesimal model is not assumed. Differences include that the constraining matrix is always singular in long term phenotypic evolution so mean fitness peaks may not be reached, and that the constraining matrix may be asymmetric so selection could reduce mean fitness and there may be evolutionary oscillations or chaos, which are forbidden under the Lande equation.

Obtaining accurate predictions under the assumption of small haplotype variation is made difficult, but not impossible, by the discrete nature of haplotype content. Taking a first-order approximation of a function of discrete variables is appropriate, as what is required is that the approximated function is differentiable, regardless of whether its variables are continuous (e.g., first-order approximations are commonly exploited to approximate the means of discrete probability distributions). However, difficulties arise for two reasons. First, the approximation cannot be made arbitrarily accurate by taking infinitesimally small deviations due to the discrete variables. Second, the approximated function is not uniquely defined: infinitely many differentiable functions can be used to describe a function of discrete variables, so there is a choice to be made for which differentiable function to use, and the choice influences how well the approximation works. A rule of thumb may be to use a differentiable fitness function and genotype-phenotype map whose expectations yield the exact mean fitness and mean genotype-phenotype map as functions of allele frequency. As illustrated in examples below, the approximation can give broadly correct results in some circumstances, in which case its analytical insight can be exploited. However, the approximation’s results can be substantially incorrect in other circumstances and so should be compared with the exact results before concluding based on the approximation. As discussed below, it is at present not straightforward to characterise under what conditions the approximation works.

For these first order approximations, I will assume that there is no developmental stochasticity, environmental stochasticity, or endogenous and exogenous environmental change, which are strong assumptions entailing genetic determinism, but I will do so to simplify the presentation. So I will assume the genotypephenotype map to be of the form

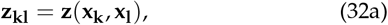

and the developmental map to be of the form

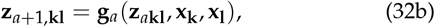

with initial condition **z**_1**kl**_ = **g**_0_(**x**_**k**_, **x**_**l**_). The genotype-phenotype and developmental maps here can also depend on constant parameters, which are not noted in the notation for simplicity.

#### Approximated evo-devo dynamics, including gradient equation for genetic evolution

We now derive a gradient equation that describes the approximated change in allele frequencies and linkage disequilibria as an adaptive topography for arbitrarily many alleles and loci, assuming that fitness, the genotype-phenotype map, and the developmental map are continuously differentiable functions, and that there is no meiotic drive or that the phenotype is symmetric with respect to maternal and paternal haplotypes.

We will use the chain rule in matrix calculus notation (Caswell 2019), which for a composition of multivariate functions **f**(**g**(**x**)) is (González-Forero 2024b)

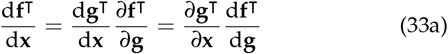

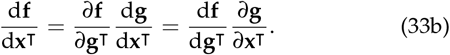

The second line is the transpose of the first line.

From the Price equation (2), the change in the mean haplotype content is

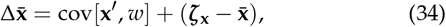

where 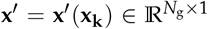 is the expected haplotype content among offspring conditional on a parent having haplotype content **x**_**k**_ and ***ζ***_**x**_ = E[**x**^*′*^]. The covariance term in this equation is cov[**x**^*′*^, *w*] = ∑_**k**_ *p*_**k**_**x**^*′*^ (**x**_**k**_)(*w*_**k**_ − 1), where *w*_**k**_ is the relative fitness of haplotype **k**.

We assume that the relative fitness of genotype **kl**, *w*_**kl**_ = *w*(**z**_**kl**_), is a continuously differentiable function of the phenotype **z**_**kl**_. We also assume that the genotype-phenotype map **z**(**x**_**k**_, **x**_**l**_) and the developmental map **g**_*a*_ (**z**_*a***kl**_, **x**_**k**_, **x**_**l**_) are continuously differentiable with respect to their arguments. As discussed above, this involves a choice of functions among infinitely many continuously differentiable functions that describe the exact, discrete-valued maps. We take the first-order approximation of relative fitness *w*_**kl**_ = *w*(**z**(**x**_**k**_, **x**_**l**_)) with respect to haplotype content around mean haplotype content:

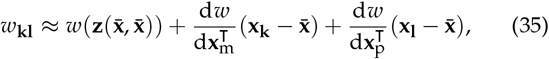

where we define the total selection gradients of maternal and paternal haplotypes as

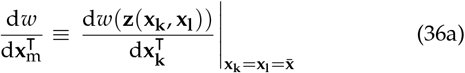

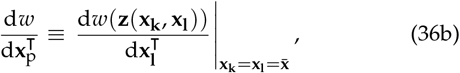

and similarly for the total effects of maternal and paternal haplotypes on the phenotype (i.e., 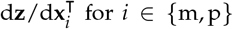). The approximation (35) improves if the haplotype contents **x**_**k**_ and **x**_**l**_ are near the mean haplotype content. From the chain rule in matrix calculus notation (33a), we have that the total selection gradient of the haplotype inherited from parent *i* ∈ {m, p} is

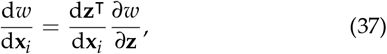

where we denote

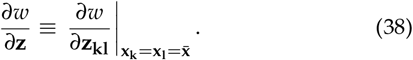

Using (35) in (27b) applying (37), the relative fitness of haplotype **k** is to first order

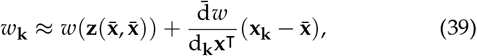

where we define the sex-averaged total derivative operator as

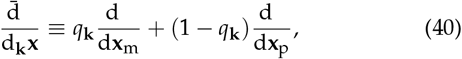

so the sex-averaged total selection gradient of the haplotype **k** is

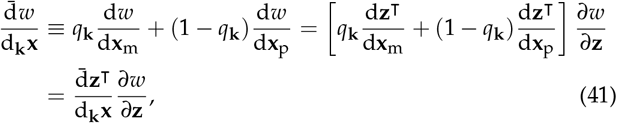

and where we used the facts that 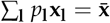 and that the total selection gradients (36) are independent of the haplotypes **k** and **l** because they are evaluated at 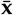.

Using (39), the total selection response is to first order

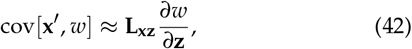

where the covariance between expected offspring haplotype and the expected phenotypic effect of parental haplotype is

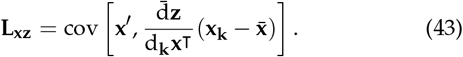

Substituting (42) into (34) yields

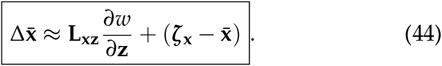

Eq. (44) will be useful to obtain an approximated gradient equation for phenotypic evolution, but is not a gradient equation for genetic evolution (notice the gradient on its right-hand side is not with respect to **x** but **z**).

Instead, we can obtain a less general approximated gradient equation for change in allele frequency and linkage disequilibrium in multiple loci by further assuming either no meiotic drive or that the phenotype is symmetric with respect to maternal or paternal haplotypes. Indeed, if *q*_**k**_ = *q* is constant over haplotypes **k** (i.e., there is no meiotic drive), then the sex-averaged total effect of the haplotype on the phenotype is independent of **k** and can be pulled out of the covariance on the right-hand size of (43), which using (37) yields

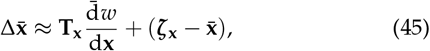

where the transmission matrix of haplotype content is

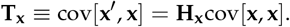

and where the sex-averaged total selection gradient of the haplotype is

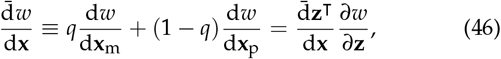

the latter equality following from (41). Eq. (45) also results if the phenotype is haplotype-symmetric (**z**_**kl**_ = **z**_**lk**_), where the sex-averaged total selection gradient of the haplotype equals the total selection gradient of the maternal or paternal haplotype. Note that selection response from non-linear inheritance **u**_**x**_ does not appear in (45) as a result of the first-order approximation.

Eq. (45) is an approximated gradient equation for change in allele frequencies and linkage disequilibria in arbitrarily many loci. In principle, the haplotype transmission matrix **T**_**x**_ may be asymmetric and so may have negative or complex eigenvalues, entailing that mean fitness could decrease in multilocus systems as a result of selection even with haplotype-symmetric fitness. However, we will see that this is not possible with two biallelic loci under random mating, so to first order the Moran (1964) phenomenon of mean fitness decrease in two-locus systems (Felsenstein 2019, p. 376) is not caused by this property of the transmission matrix but by pure transmission bias (example 4).

As allele frequencies and linkage disequilibria evolve, the developmental dynamics of the mean phenotype are to first-order given by

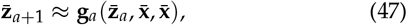

with initial condition 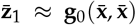. Equations (44) and (47) jointly describe the evo-devo dynamics to first order of approximation. The developmental constraint (32) affects the genetic evolutionary dynamics (44) via the total selection gradient of the haplotype (37).

#### Approximated gradient equation for short-term phenotypic evolution

We now derive a gradient equation for approximated phenotypic evolution as an adaptive topography allowing for any arbitrary genotype-phenotype map or developmental map, assuming these maps are continuously differentiable. There may be meiotic drive and the phenotype may be asymmetric with respect to maternal and paternal haplotypes provided that fitness does not directly depend on haplotype content. These gradient equations involving the phenotype have a similar structure to the gradient Lande equation, but with important differences.

Taking its Taylor expansion with respect to haplotype around the mean haplotype, the phenotype of an individual with genotype **kl** is to first order

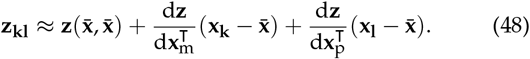

Also, to first-order, the mean phenotype is

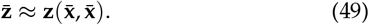

Applying the chain rule (33b) to equation (49), the change in the mean phenotype is to first order:

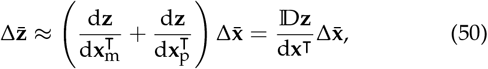

where we denote the total effect of the genotype on the phenotype as

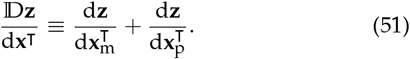

Equation (50) partitions the approximated change in the mean phenotype into that due to change in the mean haplotype content and the associated response of the mean phenotype.

Substituting (34) into (50) using (42) yields

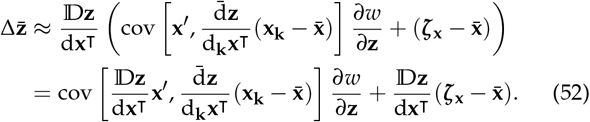

This is a gradient equation for approximated phenotypic evolution under any continuous and genetically deterministic genotype-phenotype map in sexual diploids.

We now interpret the covariance matrix premultiplying the selection gradient in Equation (52). First, analogously to (27b), we define the phenotype of haplotype **k** as **z**_**k**_ ≡ *q*_**k**_ ∑_**l**_ *p*_**l**_**z**_**kl**_ + (1 − *q*_**k**_) ∑_**l**_ *p*_**l**_**z**_**lk**_. Using here the first-order approximation of the phenotype (48) yields

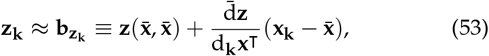

where we used the facts that 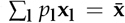 and the derivatives in (48) are independent of **l** because they are evaluated at 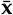. We call 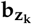 the mechanistic breeding value of the phenotype of haplotype **k**. Next, we consider the expected offspring haplotype from parent haplotype **k**:

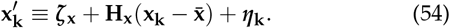

Then, we define the predicted expected offspring phenotype as that developed under the expected offspring haplotypes, 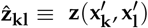, and take its first order approximation around the mean haplotype:

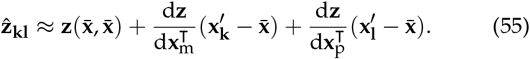

Now, we define the total expected offspring phenotype predicted from parental haplotype **k** as 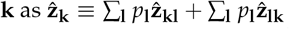 (note *q*_**k**_ is not used here in order to recover the matrix premultiplying the selection gradient in (52); thus, 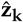 is a “total” rather than an average). Substituting here the first order approximation of the predicted expected offspring phenotype 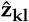 (55) yields

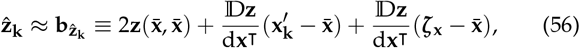

where in the last term we used (54). We call 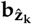 the mechanistic breeding value of the phenotype among offspring of parents with haplotype **k**. Finally, we define the mechanistic additive genetic (MAG) cross-covariance matrix as the cross-covariance matrix of offspring-parent mechanistic breeding values (L for legacy):

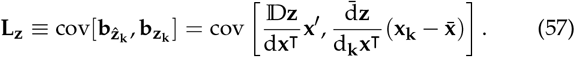

Note that this matrix is not a covariance matrix but a cross-covariance matrix (i.e., the vectors 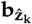 and 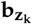 are different in general). Thus, the main diagonal entries of **L**_**z**_ are not variances but covariances and so can be negative. This means that, even for a single phenotype, this does not define a mechanistic additive genetic *variance* but a covariance between parent and offspring mechanistic breeding values. The MAG cross-covariance matrix involves haplotype variation (**x**_**k**_), haplotype heredity (**x**^*′*^), construction of parental phenotype subject to selection 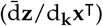, and offspring phenotype reconstruction (D**z**/d**x**^⊺^).

Hence, using (57) in (52), we obtain a gradient equation for approximated phenotypic evolution under any continuous and genetically deterministic genotype-phenotype map in sexual diploids:

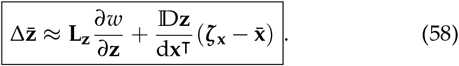

The MAG cross-covariance matrix **L**_**z**_ is analogous to Lande’s **G** matrix, but it has different properties. In particular, it is not defined in terms of regression coefficients so it does not generally satisfy the standard partitioning of phenotypic variation (**P** = **G** + **E**, but 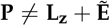, where 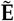 is the cross-covariance matrix of the errors of the first-order approximations of predicted-offspring and parent phenotypes of a haplotype). Moreover, in contrast to **G, L**_**z**_ may be asymmetric and so may have negative or complex eigenvalues, because **L**_**z**_ is a cross-covariance matrix rather than a covariance matrix. This entails that selection may reduce mean fitness, fitness peaks may be saddle points, and there may be oscillations and chaos. This gradient equation uses a selection gradient as in Iwasa and Pomiankowski (1991), defined as a derivative with respect to individual trait values evaluated at mean trait values rather than with respect to mean trait values.

Now, the approximated gradient phenotypic Equation (58) is dynamically insufficient because it depends on allele frequency and linkage disequilibria but does not describe the change in allele frequency nor linkage disequilibria. Thus, the equation only describes short-term evolution. That is, the equation predicts the change in mean phenotype from the current time to the next, given that allele frequencies and linkage disequilibria are known at the current time, so it does not predict changes in the mean phenotype at further times. To describe long-term phenotypic evolution as an adaptive topography, we need a dynamically sufficient gradient equation describing change in the mean phenotype, to which we now turn.

#### Approximated gradient equation for long-term phenotypic evolution

We now obtain a gradient equation for approximated long-term phenotypic evolution under any continuous and genetically deterministic genotype-phenotype map in sexual diploids.

As stated, the approximated gradient phenotypic equation is dynamically insufficient because it does not describe genetic evolution. We solve this problem by coupling it with the approximated gradient equation for genetic evolution (44) so we have the coupled system:

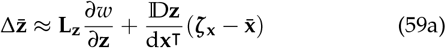

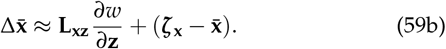

This system is overdetermined if the mean phenotype is substituted for the mean genotype-phenotype map: there are more equations than unknowns in the sense that mean phenotypes are fully determined functions of the mean haplotype, so the only unknowns are the mean haplotype. This overdeterminacy means that (59a) is unnecessary if the mean genotype-phenotype map is applied. However, the coupled system (59) is not overdetermined if the mean phenotypes are treated as unknowns by not substituting for the mean genotype-phenotype map. We now consider this latter approach as this allows us to obtain a description of long-term evolution of the mean phenotype as an adaptive topography, long term as the mean haplotype evolves.

To obtain this long-term phenotypic adaptive topography, we can write system (59) in a more compact way. To do this, we define mechanistic breeding values of the haplotypes. Let us define the mechanistic breeding value of haplotype **k** as

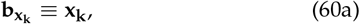

and the mechanistic breeding value of the haplotype among offspring of parents with haplotype **k** as

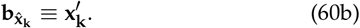

These renaming of variables allows us to form the parent and offspring mechanistic breeding values of phenotype and haplotype as

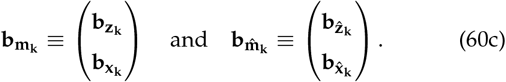

Hence, the MAG cross-covariance matrix of the phenotype and haplotype is

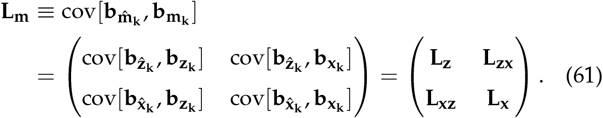

Writing **m**_**kl**_ = (**z**_**kl**_, **x**_**k**_)^⊺^, we then obtain that the system can be written as a gradient equation for the approximated change in the mean phenotype, allele frequencies, and linkage disequilibria:

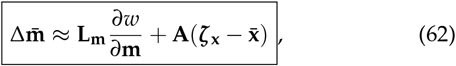

where the selection gradient of the phenotype and haplotype is

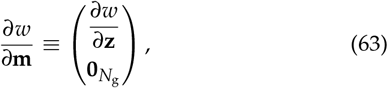

the total effects of the genotype on offspring phenotype and haplotype are

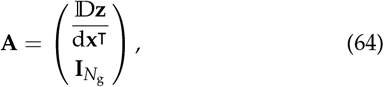

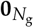 is the zero vector of dimension *N*_g_ because we assumed that relative fitness is a direct function of the phenotype but not of the haplotypes, and 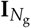 is the *N*_g_ *× N*_g_ identity matrix. Eq. (62) is an approximated equation for long-term phenotypic and genetic evolution in gradient form. It is long-term because this gradient system involving the mean phenotype is now dynamically sufficient, as it describes the evolution of all dynamic variables involved.

We can make further relatively general statements if we assume that either there is no meiotic drive (*q*_**k**_ = *q* is constant with respect to haplotypes **k**) or that the phenotype is haplotype-symmetric (**z**_**kl**_ = **z**_**lk**_). In such a case, we can write **L**_**m**_ as

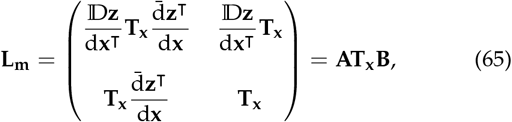

where the sex-averaged total effects of the haplotype on parent phenotype and haplotype are

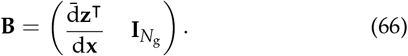

Hence, without meiotic drive or with haplotype-symmetric phenotypes, the long-term evolution Equation (62) entails that there are necessarily absolute constraints on long-term adaptation. This follows because from Eq. (66), the matrix 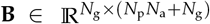 has fewer rows than columns, so it is singular (Horn and Johnson 2013, p. 14) (i.e., **Bv** = **0** does not imply **v** = **0** for any vector **v** of suitable dimension). Consequently, **L**_**m**_ is singular and its rank is at most *N*_g_, while its dimension is (*N*_p_ *N*_a_ + *N*_g_) *×* (*N*_p_ *N*_a_ + *N*_g_), so it has *N*_p_ *N*_a_ eigenvalues that are exactly zero. This singularity means that there are directions in the evolutionary space (mean phenotype and haplotype, 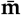) in which there is no MAG covariation and so evolutionary change in those directions is not possible. Intuitively, this is because not any mean phenotype can be produced by the developmental system, regardless of the developmental system, as long as there is some genotype-phenotype map.

Still assuming no meiotic drive or haplotype-symmetric phenotype, we can rearrange the approximated long-term evolution Equation (62) as

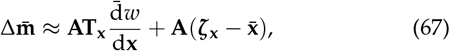

where the sex-averaged total selection gradient of the haplotype is

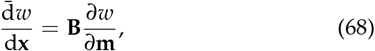

because of (66), (63), and (46), recovering the forms of the extended selection gradient of Morrissey (2014) and the total selection gradient of the genotype of González-Forero (2024b).

The alternative long-term evolution Equation (67) entails that selection response vanishes when total genetic selection vanishes 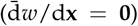 if there are no absolute haplotypic constraints (i.e., if **T**_**x**_ is invertible). This is because the matrix **A** is always nonsingular, as the product **Av** = **0** for any vector **v** of suitable dimension entails that **v** = **0**. Indeed, writing **A** = (**A**_1_; **A**_2_), we have that **Av** = **0** entails that **A**_1_**v** = **0** and **A**_2_**v** = **0**, but **A**_2_ = **I**, so **v** = **0**. Hence, while the direct selection gradient *∂w*/*∂***m** is insufficient to identify when selection response vanishes, total genetic selection 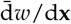 is sufficient if there are no absolute haplotypic constraints.

There are two important biological implications of absolute constraints on long-term adaptation. First, as constraints are absolute, evolutionary equilibria do not only depend on selection but also on development. Under the gradient Lande equation, a typical assumption is that there are no absolute constraints (i.e., **G** is invertible), which means that there is additive genetic variation in all directions of the evolutionary space. Thus, under this equation and without absolute constraints, evolutionary equilibria depend only on selection but not on additive genetic covariation (i.e., **G***∂* ln 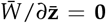 implies that *∂* ln 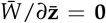). In contrast, under (62) and (65), absolute constraints are unavoidable (**L**_**m**_ is always singular), so even without transmission bias, evolutionary equilibria depend both on selection and on MAG covariation and so on development: the selection gradient *∂w*/*∂***m** is not generally zero when the selection response in (62) vanishes. That evolutionary equilibria depend on development can also be seen with the alternative arrangement (67), according to which evolutionary equilibria occur when total genetic selection vanishes if there are no absolute haplotypic constraints and no pure transmission bias. From Eq. (37), total genetic selection depends on both direct selection and development via the total effects of the haplotype on the phenotype. Thus, development and selection co-define evolutionary equilibria.

The importance that evolutionary equilibria are partly co-defined by development is illustrated by a model of brain size evolution where an equation such as (62) without transmission bias applies (González-Forero 2024a). In that model, the brain and body sizes of seven different hominins, from small-brained Australopiths to large-brained Neanderthal and *Homo sapiens*, evolve by differences in development (i.e., via a version of **A** and **B** with simplified genetics) but not in direct selection (i.e., *∂w*/*∂***z**) despite monotonic fitness (González-Forero 2024a). This is possible because of the singularity of **L**_**m**_, since if the constraining matrix were invertible, differences in development that do not affect the selection gradient cannot change the evolutionary outcomes with a single-peaked fitness.

Second, as constraints are absolute under under (62) and (65), evolutionary history matters, or more specifically, the evolutionary outcomes depend on the ancestral conditions. Under the gradient Lande equation with a single fitness peak and without absolute constraints, the evolutionary outcome is the mean fitness peak regardless of the ancestral condition. In contrast, with absolute constraints, the evolutionary outcomes depend on the ancestral conditions, even with a single fitness peak, so evolutionary history matters for the outcome. This is because with a singular constraining matrix, there are infinite evolutionary equilibria if there is any, and the one that is attained depends on the initial conditions. Indeed, evolutionary equilibria are given by setting total genetic selection to zero, but this provides only *N*_g_ equations while there are *N*_g_ + *N*_p_ unknowns. The particular equilibrium attained depends on the mean genotype-phenotype map, which provides the remaining equations, so that if the mean genotype-phenotype map at the initial condition changes (e.g., due to developmental stochasticity), so does the ensuing evolutionary trajectory and the equilibrium attained (González-Forero 2023). That absolute constraints entail that outcomes depend on the initial conditions was shown by Kirkpatrick and Lofsvold (1992) for a function-valued trait under stabilising selection and a constant and singular constraining function; in SI section S6, I show this for vector-valued traits under stabilising selection and a constant, singular, positive semi-definite, and diagonalisable constraining matrix, with a single zero eigenvalue. That absolute constraints entail that history matters has also been illustrated in the previously mentioned model of brain size evolution, where the evolved brain and body sizes over ontogeny strongly depend on the ancestral (i.e., evolutionarily initial) conditions of genotypic traits because of the singularity of the constraining matrix, despite fitness being monotonic and the selection gradient never vanishing in this model (González-Forero 2024a). This dependence of outcomes on history could be exploited to infer ancestral genotypes and phenotypes over a single lineage.

We can use the approximated long-term evolution Equation (62) to approximate adaptability to assess whether the selection response is locally adaptive to first order. Thus, adaptability is to first order given by

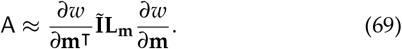

Since we are assuming that there is no direct selection for haplotypes (63), this approximation reduces to

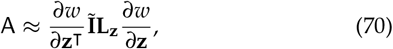

which is the approximated adaptability for the short-term evolution Equation (58). This approximated adaptability could still be negative, so selection may yield maladaptation, because **L**_**z**_ is a cross-covariance matrix. The approximated long-term evolution Equation (62) also entails that pure transmission bias 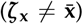 could decrease or increase mean fitness, depending on the total effects of the haplotype on the phenotype (via **A**), which depends on development rather than selection (for an example where plasticity increases mean fitness by acting through an analogous term, see González-Forero 2023).

Overall, Equation (62) yields a description of the approximated long-term evolutionary dynamics as a constrained sometimes-adaptive topography.

#### Sensitivity of the phenotype to genetic change

The above approximated equations depend on the sensitivity of the phenotype to change in the maternal or paternal haplotype, namely, d**z**^⊺^/d**x**_*i*_ (for *i* ∈ {m, p}). If a genotype-phenotype map of the form (32a) is available, deriving such derivatives is straightforward. Alternatively, if a developmental map of the form (32b) is available, the following formulas (derived in González-Forero 2024b) enable derivation of such sensitivity and provide analytical insight. Sensitivities can also be computed numerically with automatic differentiation or backpropagation.

From the chain rule (33a), the sensitivity of the phenotype **z** to change in the maternal or paternal haplotype **x**_*i*_ (for *i* ∈ {m, p}) is

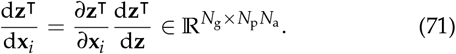

The matrix of direct effects of the haplotype on the phenotype is

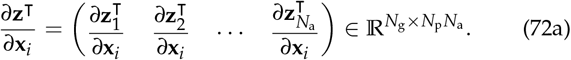

The *a*-th block entry of this matrix is

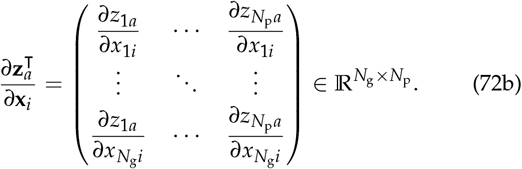

As in (36), the *jk*-th entry of this matrix for the maternal and paternal haplotype (*i* ∈ {m, p}) is

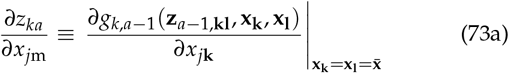

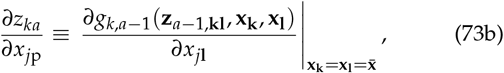

for *a* ∈ {2, …, *N*_a_}, obtained by differentiating Eq. (32b), where *x*_*j***k**_ is the *j*-th entry of **x**_**k**_. For *a* = 1, the analogous matrix is obtained by differentiating the initial condition **z**_1**kl**_.

In turn, the matrix of developmental feedback is the sensitivity of the phenotype to phenotypic change:

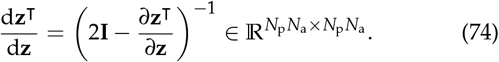

This matrix describes how phenotypes affect each other throughout development. This matrix is always invertible and has the form of a classic formula of total effects of variables on themselves (Greene 1977), but here it has underlying structure due to the recurrence equations defined by the developmental map. The matrix of direct effects of the phenotype on the phenotype at all ages is

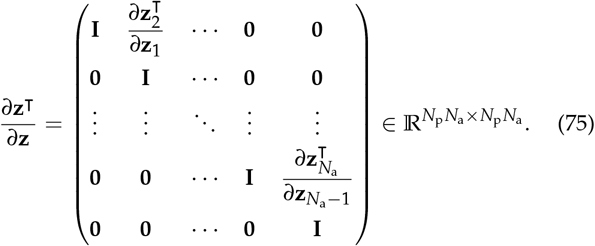

The (*a, a* + 1)-st block entry of this matrix is the matrix of direct effects of the phenotype at age *a* on the phenotype at age *a* + 1

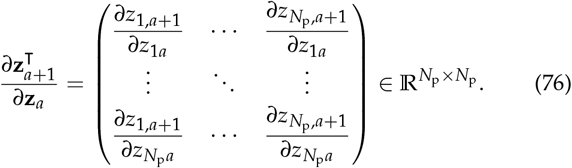

As in (73), the *a*

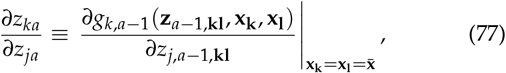

for *a* ∈ {2, …, *N*_a_}, obtained by differentiating Eq. (32b), whereas for *a* = 1 the analogous matrix is obtained by differentiating the initial condition **z**_1**kl**_.

Partitioning the matrix (71) into *N*_a_ blocks of size *N*_g_ *× N*_p_, the *a*-th block entry gives the total effects of perturbing the haplotype on the phenotype at age *a* (that is, 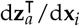, whose *jk*- th entry gives the total effect of the perturbing the *j*-th haplotypic trait on the *k*-th phenotype at age *a*). Similarly, partitioning the matrix (74) into 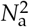 blocks of size *N*_p_ *× N*_p_, the *aj*-th block entry gives the total effects of perturbing the phenotype at age *a* on the phenotype at age *j* (that is, 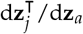, whose *ik*-th entry gives the total effect of perturbing the *i*-th phenotype at age *a* on the *k*-th phenotype at age *j*, d*z*_*kj*_ /d*z*_*ia*_).

## Results

### Examples

We now provide examples illustrating the methods above. To keep this paper of reasonable length, these examples do not aim to illustrate the full range of possibilities afforded by the methods. Instead, the examples are designed to be as simple as possible to illustrate the basic workings of the theory and relate it with standard results, recovering some modifications of classic results such as an adjustment to Wright’s equation for allele frequency change in a biallelic locus and obtaining some new results such an exact gradient equation for change in allele frequency and linkage disequilibrium in two biallelic loci. Thus, the examples in the main text consider implicit development, with additional examples in the SI, some of which consider explicit development (SI section S9). A Mathematica notebook with code to generate all figures and do routine derivations for these examples is available as Supplementary Material.

### Example 1: joint evolution of mean phenotype and phenotypic variance under quantitative genetics: evolution of ubiquitous perfection

A standard approach to study the evolution of phenotypes and additive genetic variances and covariances is to treat them as unconstrained dynamic variables by coupling dynamic equations describing change in mean phenotype, variances and covariances (Lande and Arnold 1983). In this example, we illustrate the results of taking this classic approach, to compare it with our approach. Specifically, we use equations under quantitative genetics assumptions (specifically, the infinitesimal model) coupling phenotype and variance evolution to study the evolution of mean phenotype and phenotype variance. We recover the standard result that stabilising selection depletes phenotypic variation under the infinitesimal model (Robertson 1956, Barton1986 1986 and Walsh and Lynch 2018 Example 5.6 and p. 1016). This contrasts with population genetic models where phenotypic variation can be maintained with stabilising selection (Bürger 2000). In subsequent examples, we will recover the population genetics results with our approach, and conclude that the discrepancies arise because the infinitesimal model allows organisms to develop any phenotype, effectively removing any constraints.

Consider a single phenotype *z* that is normally distributed (e.g., under the infinitesimal model) with mean 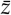 and variance *P* and assume perfect heredity and no pure transmission bias (*H* = 1 and *z′* = *z*). Consider a Gaussian absolute fitness function *W* = exp[−(*z* − *θ*)^2^/*σ*^2^]; hence, mean fitness is 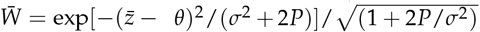, with optimum mean phenotype at *θ*. Hence, from (19) and (24), we have the coupled dynamical system

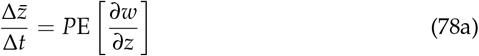

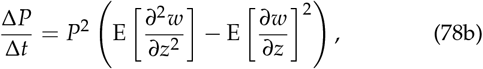

where the average fitness gradient is

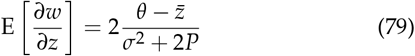

and the average fitness Hessian is

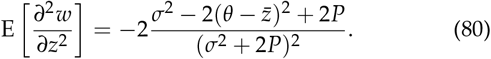

Hence, system (78) is

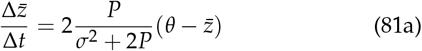

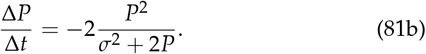

Rescaling time to *τ* = 2*Pt*/(*σ*^2^ + 2*P*), the dynamical system reduces to

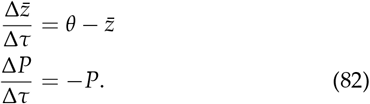

Therefore, the population always attains the optimum phenotype and phenotypic variance vanishes given enough time if there is initially some phenotypic variation (otherwise the normality assumption does not hold; Fig. 1). To put it simply, everyone is perfect in the long run. A common extension of this approach is to include mutation (an **M** added to the equation of Δ**P**, analogous to our **P***′* − **P**; Phillips and Arnold 1989), which yields a mutation-selection balance with small phenotypic variance at equilibrium.

**Figure 1.**
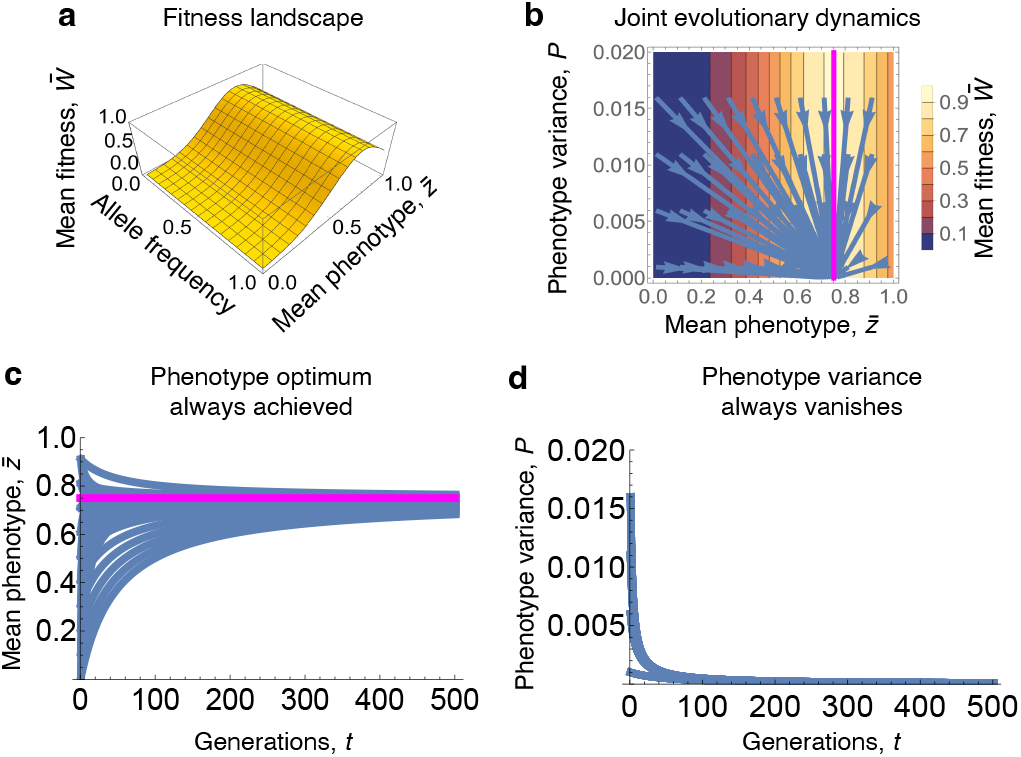
Example 1: Evolution of one phenotype under quantitative genetics. **a** Fitness landscape. **b** Evolutionary dynamics of the mean phenotype and phenotype variance under system (81), which respectively converge to the optimum phenotype and no phenotypic variance. **c,d** The dynamics over time. Parameter values: *θ* = 0.75 and *σ*^2^ = 0.1. Mean fitness is plotted at *P* = 0.01.

### Example 2: generator equation coefficients under one diallelic locus: heredity differs from heritability

In this example, we illustrate how to calculate the coefficients of the generator equations applied to one phenotype influenced by one diallelic, autosomal locus in a large diploid population under random mating and non-overlapping generations. We also compare these coefficients to standard coefficients, specifically, additive genetic variance and heritability.

Let 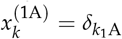 be the gene content for allele A in haplotype *k* ∈ ℍ = {a, A}, where 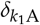 is the Kronecker delta. That is, the gene content for allele A in haplotype a is 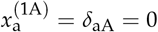 and in haplotype A is 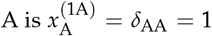. Let *p*_*k*_ be the frequency of haplotype *k* ∈ H in the population. Thus, the frequency of allele A is the mean gene content 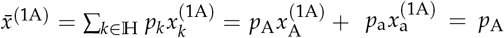 (Price 1970). For readability, we will drop the superscripts of *x* in this example from now on since there is only one locus and two alleles, so 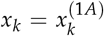, and 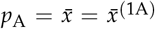. The variance in gene content is then

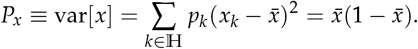

This is half the variance in genotype content (Eq. IX-27 of Felsenstein (2019)) because *x* is here haplotype content that goes up to one rather than genotype content that goes up to two.

Let *z*_*kl*_ be the phenotype of an individual with genotype *kl* ∈ G = {aa, aA, Aa, AA} and, following Lynch and Walsh (1998) (their Fig. 4.6), let it be given by the genotype-phenotype map

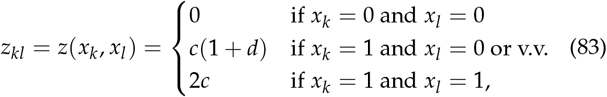

where *c* ∈ ℝ scales the phenotypic value and *d* ∈ ℝ is the coefficient of dominance, so the phenotype is haplotype-symmetric (Fig. 2a-g). In the approximated method, we will consider the following differentiable genotype-phenotype map that meets this definition, but note it is not the only one that does:

**Figure 2.**
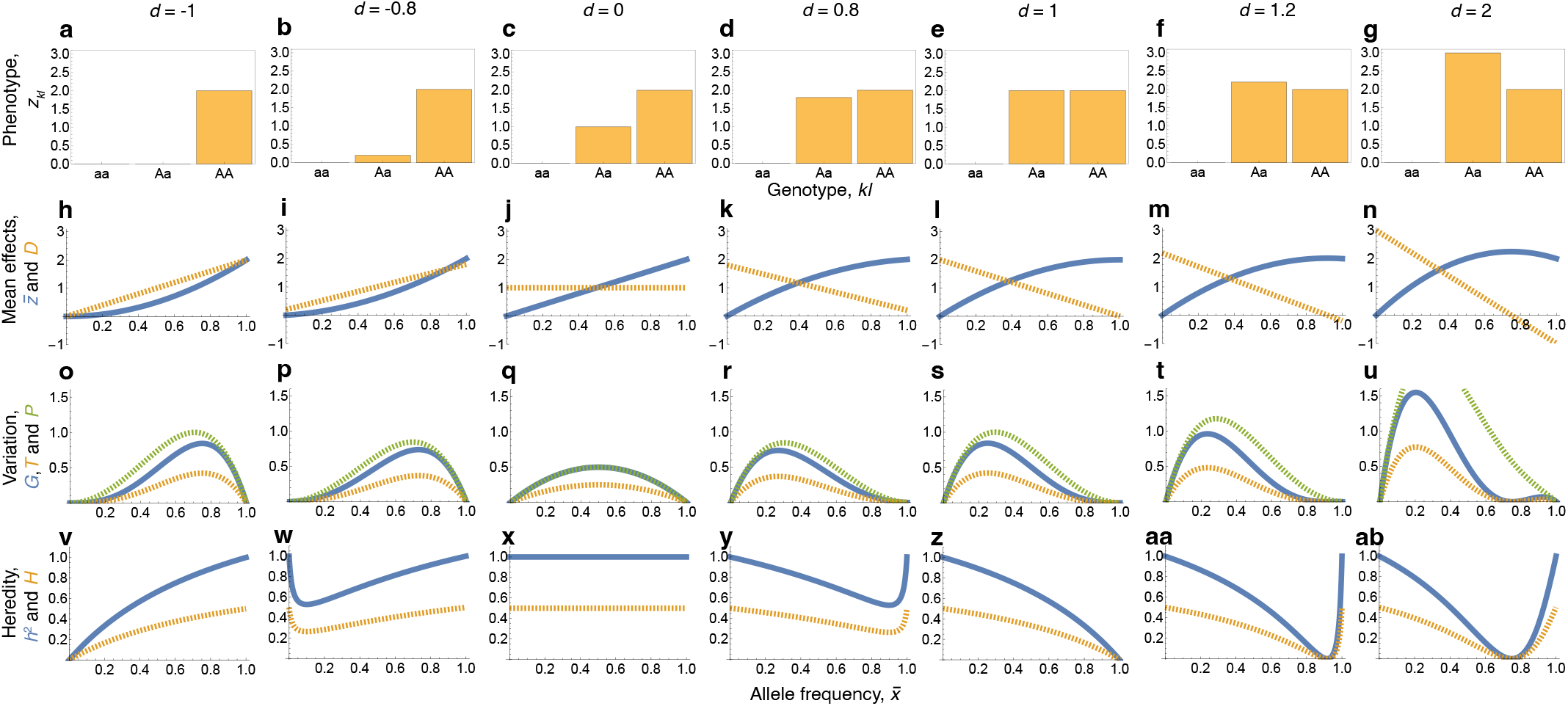
Example 2: Genotype-phenotype map and evolutionary coefficients for one phenotype influenced by one biallelic locus in a randomly mating diploid population. In all panels, *c* = 1.

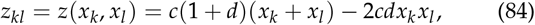

so when *d* = 0 the allele A has additive effects on the phenotype (*z*_*kl*_ = *c*(*x*_*k*_ + *x*_*l*_)) and when *d* = −1 the allele A is recessive and has multiplicative effects (*z*_*kl*_ = 2*cx*_*k*_*x*_*l*_). Let *p*_*kl*_ be the frequency of genotype *kl*. Given random mating, genotypic frequencies are in Hardy-Weinberg equilibrium, whereby 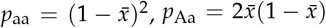, and 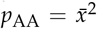. Following Lynch and Walsh (1998) (their *N* in p. 65), we also define genotype content *y*_*kl*_ as being 0 if *kl* = aa, 1 if *kl* = aA or Aa, or 2 if *kl* = AA.

Thus, we have the following distributions for haplotype, genotype, and phenotype. The haplotype content *x*_*k*_ is a Bernoulli distributed random variable that takes the value 1 with probability 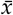. Genotype content *y*_*kl*_ is a binomially distributed random variable with two Bernoulli trials (one for each haplotype in ℍ). The phenotype value *z*_*kl*_ is a random variable whose distribution is binomial if *d* = 0 and *c* = 2, but is not binomial otherwise.

#### Phenotype mean, variance and development coefficients

The mean phenotype is

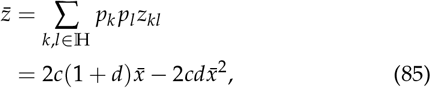

which reduces to 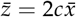 if *d* = 0. Thus, 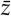 is a function of allele frequency 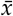, so as allele frequency evolves, the mean phenotype can only take values allowed by this equation.

To compare our coefficients with additive genetic variance and heritability, we compute the coefficients involved in such quantities. Such coefficients are those arising from the linear regression of phenotype *z*_*kl*_ on genotype content *y*_*kl*_, with slope *D* and intercept *ξ* (20). From least squares, they are

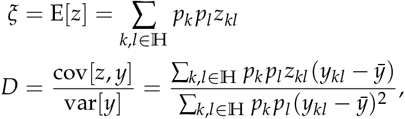

where 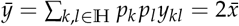. This yields

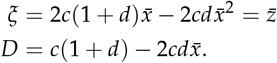

This *D* is Fisher’s average effect of allelic substitution (Eq. 4.10b of Lynch and Walsh 1998 after substituting their *a* for our *c*, their *k* for our *d*, their *p*_1_ for 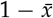, and their *p*_2_ for 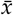). The coefficients *ξ* and *D* also depend on allele frequency 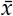. Note that 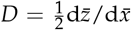. If *d* = 0, they reduce to 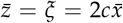 and *D* = *c* (Fig. 2h-n).

In accordance with Lynch and Walsh (1998) (their eqs. 4.11 and 4.12), the variance of the phenotype is

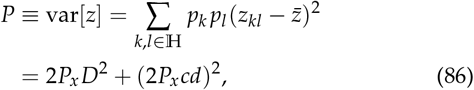

where the first term is the additive genetic variance and the second term is the dominance genetic variance. With additive effects (*d* = 0), the phenotype variance reduces to the additive genetic variance 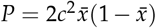. Thus, the phenotypic variance *P* is a function of allele frequency 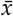, rather than an unconstrained variable that can take any positive value.

#### Transmission coefficients

We now compute the conditional expectations of offspring phenotype given parent phenotype: *z′* (*z*) = E_*z*_+ |_*z*_ [*z*^+ |^*z*]. With random mating, the expected phenotype of offspring given that a parent has phenotype *z*_*aa*_ is

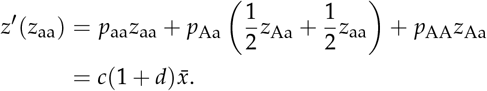

This is because a parent aa mates with aa with probability *p*_aa_, in which case all their offspring have phenotype *z*_aa_; it mates with Aa with probability *p*_Aa_, in which case half their offspring have phenotype *z*_Aa_ and half have phenotype *z*_aa_; and it mates with AA with probability *p*_AA_, in which case all their offspring have phenotype *z*_Aa_; see also Frank (1997) (his Appendix A). Similarly, the expected phenotype of offspring from a parent with phenotype *z*_Aa_ is

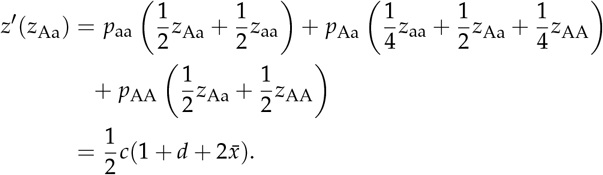

Finally, the expected phenotype of offspring from a parent with phenotype *z*_AA_ is

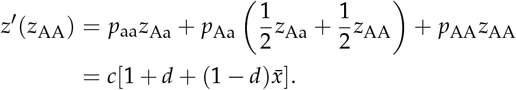

Thus, the covariance between expected offspring phenotype and parent phenotype is

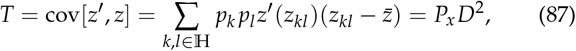

which reduces to 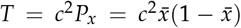 if *d* = 0, and also depends on allele frequency. Here *T* is non-negative (Fig. 2o-u). Yet, transmission *T* can be slightly negative in this same scenario but under selfing rather than random mating (SI section S9.1).

Regarding the regression of expected offspring phenotype on parent phenotype (with intercept *ζ* and slope *H*), from least squares, it follows that

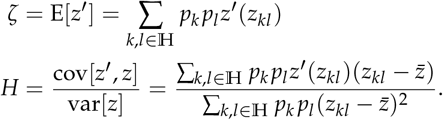

Plugging the values specified, this yields

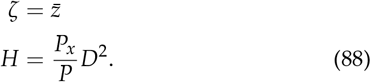

Thus, there is no pure transmission bias in the phenotype 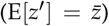 and heredity *H* depends on allele frequency 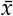. If the allelic effect is additive, that is *d* = 0, *H* reduces to *H* = 1/2, but if *d* ≠ 0, then generally *H* ≠ 1/2 (Fig. 2v-ab). Although heredity *H* is here non-negative and smaller than one, heredity can be negative or greater than one under selfing, in contrast to heritability (SI section S9.1).

The variance in expected offspring phenotype is

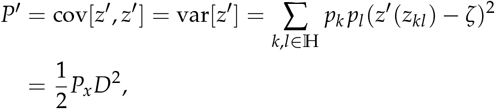

which reduces to 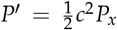 if *d* = 0. Note that *P′* = *H*^2^ *P* for *d* = 0 but *P′* ≠ *H*^2^ *P* for *d* ≠ 0. Thus, from (18), if *d* = 0, then var[*η*] = 0, so the residuals of the regression of expected offspring phenotype on parent phenotype do not vary and the regression exactly predicts expected offspring phenotype. Instead, if *d* ≠ 0, these residuals vary (var[*η*] ≠ 0) and the regression does not exactly predict expected offspring phenotype.

Using (5), the lack of variance in residuals *η* means that there cannot be selection response from non-linear inheritance for the phenotype (*u* = 0). Although this only holds under additive allelic effects of gene content on the phenotype (*d* = 0), in examples below this holds in general for haplotype content, which is trivially additive, so there cannot be selection response from non-linear inheritance for haplotype content.

#### Heritability

The primary generator equation depends on transmission **T** and heredity **H** = **TP**^−1^, whereas the Lande equation depends on the additive genetic covariance matrix **G** and multivariate heritability **GP**^−1^. We now compare heredity *H* and heritability in this example.

Additive genetic variance is by definition

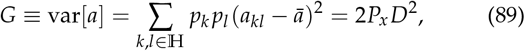

where breeding value is the best linear prediction of the phenotype from genotype content, 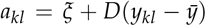, and 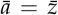. Eq. (89) recovers classic expressions for the additive genetic variance for a phenotype controlled by a single diallelic locus under random mating (Eq. 4.12a of Lynch and Walsh 1998 after substituting their *α* for our *D*). From Eqs. (87) and (89), we have that in this example 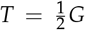 (Fig. 2o-u), which recovers the standard result (Eq. IX-53 of Felsenstein 2019). Also, *G* reduces to 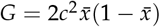 if *d* = 0 and so depends on allele frequency. It can be checked that here 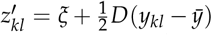, so assumption (I) of the Lande equation (i.e., *z′* = E[*a*|*z*]) does not hold here.

Narrow sense heritability is

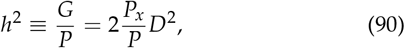

which reduces to *h*^2^ = 1 if *d* = 0. Both heritability and heredity *H* may be zero under particular combinations of dominance and allele frequencies (Fig. 2v,z,aa,ab). From (88) and (90), 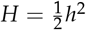 in this example, recovering the classic result that the parent-offspring regression is half heritability (Fisher 1918, Eq. 10.134 of Nagylaki 1992, and Eq. IX-56 of Felsenstein 2019). The midparent offspring regression does equal heritability in this situation (Eq. IX-61 of Felsenstein 2019). This halving of heritability into the heredity that results in the exact total selection response had previously been identified (Heywood 2005).

### Example 3: evo-devo dynamics of one phenotype influenced by one biallelic locus under implicit development: evolution of common imperfection

In this example, we now use our approach to study the evolution of one phenotype influenced by one diallelic locus. To do this, consider again the scenario of the previous example, with one haplotype-symmetric phenotype influenced by a single biallelic locus in a large diploid randomly mating population having non-overlapping generations. We first use the exact method and then use the approximated method.

#### Exact method: constraints affect outcomes

We have the mean genotype-phenotype map (85) and the mean genotype-variance map (86), so to model here the evolution of the mean phenotype and phenotype variance, it is enough to model the evolution of allele frequency and compute the resulting mean phenotype and phenotype variance as emergent properties. Thus, if we know how allele frequency 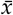 changes, we know how the mean phenotype and phenotype variance change for a given *d*. There is no need for an additional equation describing the change in mean phenotype or phenotype variance, except to understand the different factors that affect their evolution.

From the primary generator equation applied to haplotypes (26), allele frequency change for a single diallelic locus is in general given by

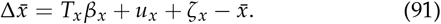

We now proceed to compute the elements of this equation.

The covariance between expected offspring and parent haplotypes is

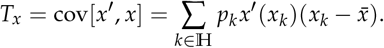

As *x*_*k*_ is the gene content of haplotype *k*, then *x′* (*x*_*k*_) is the expected gene content among the offspring haplotypes of parent haplotype with gene content *x*_*k*_. With random mating, the probability that haplotype a pairs with haplotype a is *p*_a_, in which case all their offspring haplotypes are a with gene content *x*_a_. Using this reasoning, we have that offspring haplotypes are

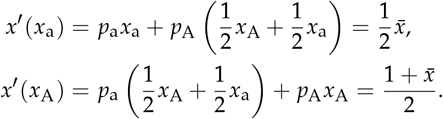

Hence, the covariance between expected offspring and parent haplotypes is

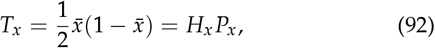

where the rightmost equality follows from (11), so the heredity of the haplotype is *H*_*x*_ = 1/2.

The expected haplotype content in offspring is

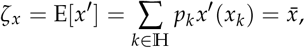

so there is no change in allele frequency due to pure transmission bias in the haplotype.

The variance in expected offspring haplotypes is

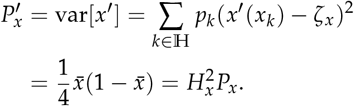

Hence, from (18), it follows that there is no variance in the residuals of the regression of expected offspring on parent haplotypes (var[*η*_*x*_] = 0), so such regression predicts expected offspring haplotype exactly. Thus, using (5), there cannot be selection response from non-linear inheritance for haplotypes (*u*_*x*_ = 0).

In turn, from least squares, the selection regression coefficient of the haplotype is *β*_*x*_ = cov[*x, w*]/var[*x*]. The relative fitness of genotype *kl* is 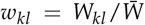, and mean relative fitness is 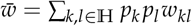, so 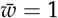. Assuming haplotype-symmetric fitness, *w*_*kl*_ = *w*_*lk*_, the haplotype selection differential is

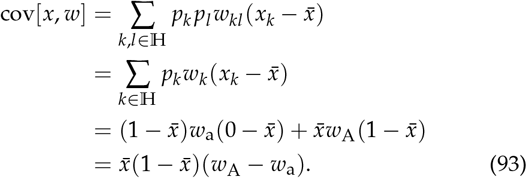

This is the standard expression for change in allele frequency for a biallelic locus in diploids (Eq. II34 of Felsenstein 2019), but is not exactly the expression for change in allele frequency we recover as shown as follows. The selection regression coefficient of the haplotype is then

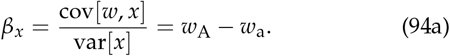

Similarly, the selection pointer of the haplotype is

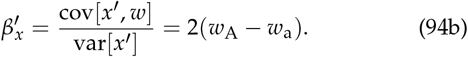

From Eqs. (91) and (92), allele frequency change is here half the haplotype selection differential

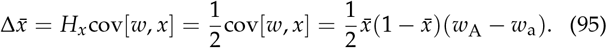

Hence, change in allele frequency is here half Wright’s classic result, with a 1/2 dilution due to imperfect heredity of haplotype content. This contrasting result emerges from considering elements of selection that are part of transmission bias in the Price equation (Eq. (6)), which leads to an adjusted measurement of inheritance using heredity rather than heritability (Heywood 2005). Noting that mean absolute fitness is 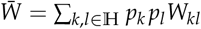 and assuming that absolute fitness *W*_*kl*_ is haplotype-symmetric (*W*_*kl*_ = *W*_*lk*_) and independent of allele frequency 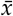, it can be checked that the selection gradient of allele frequency is

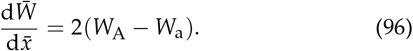

Hence, using (94a), the selection regression coefficient of the haplotype is one-half of the selection gradient of allele frequency:

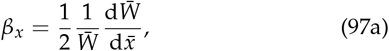

and the selection pointer of the haplotype equals the selection gradient of allele frequency:

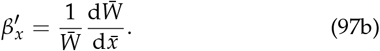

Also, the adaptability of the haplotype is

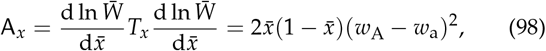

which is non-negative. Thus, selection response of the haplotype cannot locally decrease mean fitness here.

Hence, since here 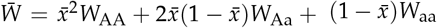, allele frequency change can be written as the gradient equation

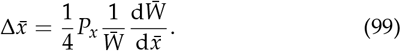

As previously noted, the right-hand side of this equation equals one half of the classic Wright (1937) equation for allele frequency change (Eq. 5.5b of Walsh and Lynch 2018; Eq. II-115 of Felsenstein 2019), because of the correction by haplotype heredity *H*_*x*_ (Heywood 2005).

Specifying fitness makes Eq. (95) dynamically sufficient to model the evolution of allele frequency, mean phenotype, and phenotypic variance, as the only dynamic variable that affects the latter two is allele frequency. Let the absolute fitness of genotype *kl* be

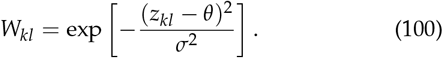

Thus, mean absolute fitness is 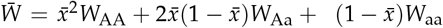, where each *W*_*kl*_ is a non-linear function of *z*_*kl*_. Note that mean absolute fitness is computed over a binomial distribution of genotypes with two Bernoulli trials (one for each haplotype), rather than over a normal distribution. Consequently, mean absolute fitness is here a function of the allele frequency and the phenotypes of all individuals but not of mean fitness, because of the non-linearity of this fitness function and the binomial, rather than normal, distribution of genotypes. To note this, let us write 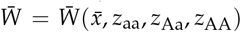. Hence, the relative fitnesses of alleles are functions of the allele frequency and of the phenotypes of all individuals, but not of mean fitness, so let us write 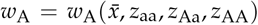 and 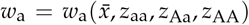. Therefore, the dynamics of allele frequency are given by

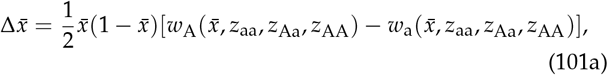

subject to the static developmental constraint

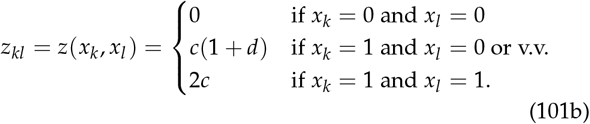

Equations (101) describe the constrained evolutionary dynamics of allele frequency, and the evolutionary dynamics of the mean phenotype emerge by using the mean genotype-phenotype map (85). Numerical solutions under some parameter values are in Fig. 3a-o.

**Figure 3.**
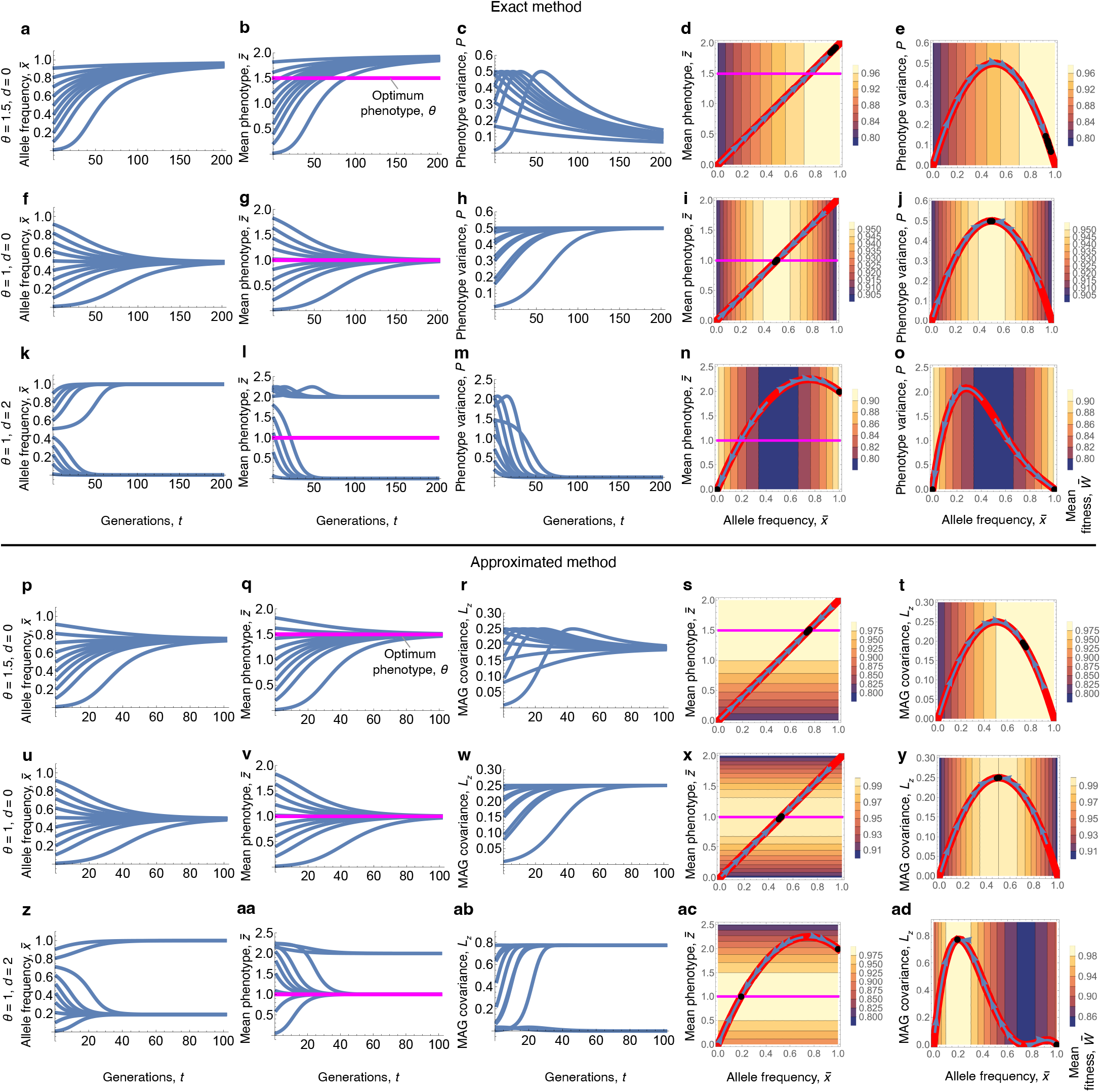
Example 3: Evolution of one phenotype influenced by one biallelic locus with implicit development. **a-o**, Exact method, solving system (101). In red is the developmental constraint (admissible evolutionary manifold). Evolutionary trajectories (blue lines) only occur along this path. Parameter values: *c* = 1 and *σ*^2^ = 10 in all panels (weak selection is used to compare with the weak-selection approximation below); **a-e**, *θ* = 1.5 and *d* = 0, **f-j**, *θ* = 1 and *d* = 0, and **k-o**, *θ* = 1 and *d* = 2. In **a-e**, the phenotype optimum is not achieved. In **f-j**, the phenotype variance increases. In **k-o**, different mean phenotypes evolve depending on the initial condition. None of these results are possible under the Lande equations. In **d,i,n**, the red lines are the mean phenotype 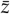 as a function of allele frequency 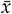. In **e,j,o**, the red lines are the phenotypic variance *P* as a function of allele frequency 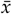. **p-ad**, Approximated method, assuming small haplotype variation and weak selection. Plotted are the same scenarios as in **a-o**, solving system (110). In **p-t**, the phenotype optimum is achieved, in contrast to **a-e.** In **u-y**, the MAG covariance increases; this case is a close approximation of **f-j**. In **z-ad**, different mean phenotypes evolve depending on the initial condition, as in **z-ad**, but the phenotype optimum is achieved for some trajectories, in contrast to **z-ad**. The approximations are somewhat poor, except in the cases where the optimum is achieved in the exact method. In **s,x,ac**, the red lines are 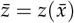. In **t,y,ad**, the red lines are *L*_*z*_ as a function of 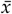. MAG: mechanistic additive genetic.

We highlight a series of contrasts with quantitative genetics under the infinitesimal model as illustrated in our example 1. First, the population evolves either away from the optimum or to the optimum but preserving phenotypic variation; simply put, some or all individuals are imperfect in the end. Internal equilibria for allele frequency (i.e., for 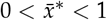) are found by solving for 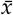 in *W*_A_ = *W*_a_, or in 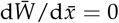, which yields

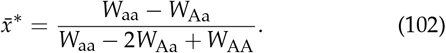

Substituting (100) and (101b) into (102), there is a single internal allele frequency equilibrium, given by

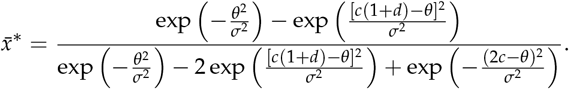

Evaluating the mean phenotype 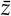 (85) at this 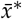 does not generally yield the optimum phenotype *θ*, not even with *d* = 0 (Fig. 3b,l). The reason is that the internal equilibria of (99) occur when the selection gradient of *allele frequency* vanishes, not necessarily when the selection gradient of the *mean phenotype* vanishes. Indeed, Equation (99) does not describe the evolution of the mean phenotype as the climbing of a mean fitness landscape in phenotype space. This is not possible in this model because mean absolute fitness cannot be written as a function of the mean phenotype in this model. Thus, (99) does not describe an adaptive topography for the mean phenotype. In contrast, if the phenotype is multivariate normal, mean fitness can be written as a function of the mean phenotype. Yet, as shown below, the approximated method for this example does describe an adaptive topography for the mean phenotype.

Second, the evolutionary trajectories of the mean phenotype and phenotypic variance are not scattered over the space with arbitrary initial conditions (as in Fig. 1b), but follow the trajectory specified by the developmental constraint (Fig. 3d,e,i,j,n,o). Third, phenotype variance can increase transiently (Fig. 3c) or permanently (Fig. 3h), which can be understood as a result of the phenotype variance being constrained to evolve along a path where the phenotypic variance must increase for mean fitness to increase (Fig. 3e,j). Fourth, populations can evolve different mean phenotypes and allele frequencies depending on initial conditions, because of disruptive total genetic selection created by the genotype-phenotype map (Fig. 3k,l). Indeed, even though there is only stabilising direct selection for the phenotype, the non-linearity of the genotype-phenotype map reshapes the fitness landscape experienced by genes to make total genetic selection disruptive (blue mean fitness valley in Fig. 3n).

To analyse why the phenotypic covariance can increase in these results, let us derive an equation describing the constrained change in the phenotypic variance. Taking the derivative of Eq. (86) with respect to time, we obtain after simplification

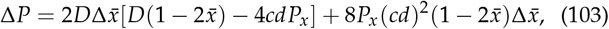

where the first term is the change in the additive genetic variance and the second one is the change in the dominance genetic variance. With additive allelic effects (*d* = 0), this reduces to 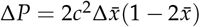, where phenotypic variance increases if allele frequency increases but is less than 1/2 or if allele frequency decreases but is greater than 1/2. This differs substantially from the equation describing the unconstrained change in phenotypic covariance in general (24) and in particular for example 1 (81b). Specifically, according to (103), the phenotypic variance can increase under stabilising direct selection on the phenotype, which cannot happen under the unconstrained equation.

This series of contrasts agree with the previously known contrasts between quantitative genetics models underpinned by many loci with additive effects and population genetics models involving a few loci with potentially non-additive effects (Bürger 2000). As illustrated in example 1, under the gradient Lande equation with a single mean fitness peak and invertible **G**, stabilising selection leads to the phenotypic optimum and depletes phenotypic variation regardless of the ancestral conditions. Yet, as illustrated in the current example, in population genetics models with a single mean fitness peak, stabilising selection may not lead to the optimum nor deplete phenotypic variation and outcomes depend on ancestral conditions.

A fundamental reason for this series of contrasts is that quantitative genetics under the infinitesimal model imposes no active constraints on the mean phenotype and phenotype variance, so selection is free to shape them. In contrast, in population genetics models, where allele frequency change is followed, the mean phenotype and phenotype variance are specific functions of allele frequency that must be satisfied. In effect, the normality assumption in quantitative genetics implicitly allows for the mean phenotype and phenotype variance to take any value, without constraints. Intuitively, the reason is that a finite number of phenotypes is specified by the additive contribution of an infinite number of independent loci, so any phenotype combination can be produced by development by tuning the weight of the contribution of each locus.

We have illustrated here how to obtain an exact, dynamically sufficient description of both genetic and phenotypic evolution for one phenotype influenced by a single diallelic locus by applying a generator equation to allele frequency change coupled with a developmental constraint. Alternatively, it is also possible to obtain an exact, dynamically sufficient description of both genetic and phenotypic evolution for one phenotype influenced by a single diallelic locus by applying a generator equation to both allele frequency and mean phenotype. However, doing so does not yield a gradient system in this example, so such an approach is not here conducive to understanding phenotypic evolution as an adaptive topography (SI section S9.2).

#### Approximated method: constraints as singular matrix

We now analyse the same scenario using the approximated method, which can describe phenotypic evolution as an adaptive topography assuming small haplotype variation around the mean haplotype and weak selection.

Since in this example we are assuming that the phenotype is haplotype-symmetric, then Eq. (45) applies. Thus, the change in allele frequency for a single biallelic locus is to first order

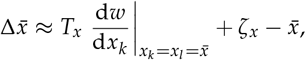

where the total selection gradient for the haplotype is

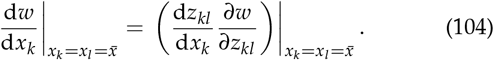

We have seen that 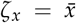, so there is no pure transmission bias in haplotype content in this example. We also have that 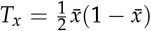 and following (84) we choose the continuously differentiable genotype-phenotype map

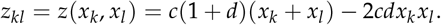

Thus, to first order, the mean phenotype is

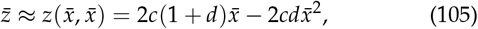

which equals the exact mean phenotype (85), so our choice of the continuous genotype-phenotype map meets our rule of thumb. Hence, the total effect of the haplotype on the phenotype is

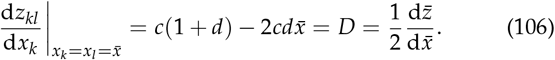

Allele frequency change then reduces to

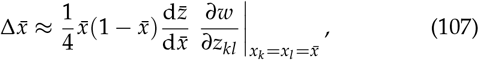

in close analogy with (99) but now this equation depends on a fitness gradient with respect to *phenotype*, so it describes changes in allele frequency as the climbing of a fitness landscape in phenotype space. Thus, there are three types of allele frequency equilibria to first order: first, if there is no haplotype variation (*P*_*x*_ = 0); second, if there is no effect of allele frequency change on mean phenotype 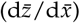; and third, if there is no direct phenotypic selection 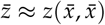.

Hence, if there is haplotype variation, evolutionary outcomes are not only given by setting the selection gradient of the phenotype to zero, but by setting the total selection gradient of the haplotype to zero given the developmental constraint:

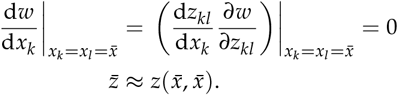

The evolutionary outcomes thus depend on the genotype-phenotype map, both through d*z*_*kl*_ /d*x*_*k*_ and through the constraint 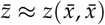.

As before, let absolute fitness be *W*(*z*_*kl*_) = exp[− (*z*_*kl*_ −*θ*) /*σ*]. From our assumption of small haplotype variation, mean fitness is 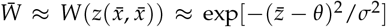 to first order. Then, the selection gradient of the phenotype is

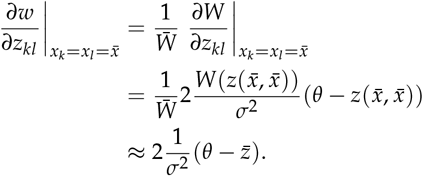

Thus, the three types of allele frequency equilibria occur as follows. First, when haplotype variance vanishes, which is at 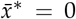 or 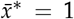. Second, when there is no effect of allele frequency change on mean phenotype (*D* = 0), which is at 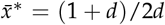 if *d* ≠ 0 and requires |*d*| > 1 for this 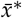 to be between zero and one. Substituting this allele frequency into 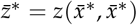 yields 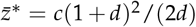, which is independent of the optimum *θ*. Third, when the mean phenotype is optimal, at 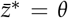. Substituting the latter into 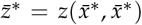, the optimum phenotype equilibrium 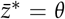 occurs to first order with the allele frequency

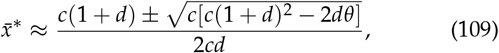

which by L’Hôpital’s rule reduces to 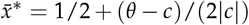 if *d* tends to zero and *c* ≠ 0. If the quantity (109) is greater than one or smaller than zero, the optimum phenotype is not attainable, again to first order.

Allele frequency change then reduces to

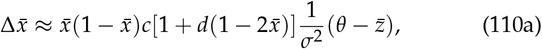

subject to the static developmental constraint

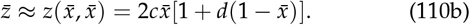

Numerical solutions for this example (110) under the approximated method are in Fig. 3p-ad. In these solutions, the approximated method tends to overestimate the ability of the system to converge to the optimum phenotype, sometimes giving qualitatively highly incorrect results (e.g., with the optimum phenotype reached and MAG covariance maintained with overdominance under some initial conditions, whereas in the exact method the optimum phenotype is not achieved and phenotypic variance vanishes; Fig. 3l,m,aa,ab). In the case where the approximated method qualitatively recovers the exact results of evolution toward the optimum phenotype, the approximated method also recovers an increase in MAG covariance under stabilising total genetic selection (Fig. 3w), the analogous of which was not possible with quantitative genetics (example 1).

To analyse this model as an adaptive topography for the phenotype, we now consider the approximated gradient Equation (58) for short-term phenotypic evolution. For a single, haplotype-symmetric phenotype influenced by a single biallelic locus, we have that this equation is

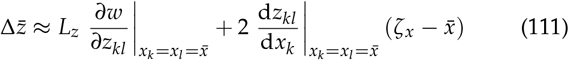

where, from (57), the MAG covariance of the phenotype is

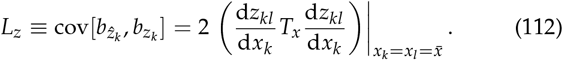

Hence, using the previously calculated quantities, we obtain that *L*_*z*_ = *T* and

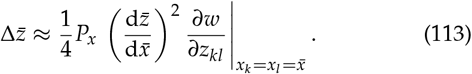

This expression has a Lande form with the selection gradient sensu Iwasa and Pomiankowski (1991), additive genetic variance sensu Barton and Turelli (1987), and a 1/4 correction as found above for the gradient equation of allele frequency change (99). As noted before, the haplotype variance (*P*_*x*_) and the total effect of allele frequency on the mean phenotype 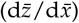 depend on allele frequency. Indeed, Eq. (113) is

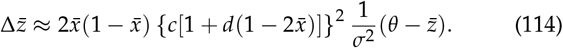

This dynamical system is underdetermined: it has more unknowns (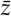 and 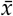) than equations, so it does not specify a unique evolutionary trajectory. Thus, the system is dynamically insufficient. To have a dynamically sufficient system, we also must describe allele frequency change.

If we track both phenotypic and genetic evolution, then from (62) the evolutionary change is given by

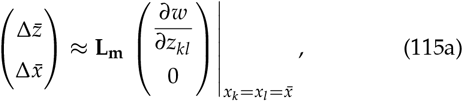

where from (65) the MAG covariance matrix of phenotype and haplotype is

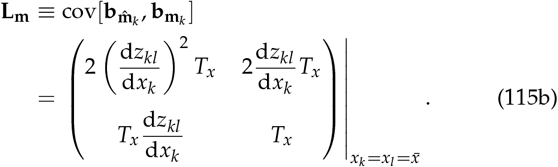

To check whether **L**_**m**_ is singular, take its determinant:

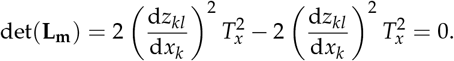

Therefore, **L**_**m**_ is always singular, regardless of the genotype-phenotype map and how it evolves. The matrix **L**_**m**_ is asymmetric and it can be checked that one of the eigenvalues of the symmetric part of **L**_**m**_ can be negative, so adaptability could be negative here to first order; that is, selection could yield mal-adaptation here. Indeed, adaptability is here, to first order,

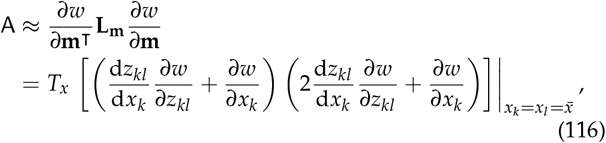

where we have allowed for non-zero direct selection for haplotype content (we had assumed this was zero to simplify our derivations of the approximate gradient equations, but allowing for this to be non-zero yields the same Equation (62) if the phenotype is haplotype-symmetric). This adaptability could be negative if direct selection for haplotype content is non-zero and direct selection for the phenotype and haplotype have opposite signs. However, under our assumption that direct selection for haplotype content is zero, adaptability reduces to

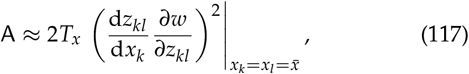

which is non-negative and so maladaptation by selection is not possible, to first order. Thus, the approximated selection response **L**_**m**_ ∂*w*/∂**m** points at ≤ 90*◦* from the selection gradient ∂*w*/∂**m**, regardless of the genotype-phenotype map and how it evolves in this system involving a single phenotype under selection whose development is influenced by a single biallelic locus under random mating.

Since **L**_**m**_ is always singular, the approximated long-term phenotypic evolution does not generally reach a local fitness peak. In this simple example, this is only possible under the trivial situation where the MAG covariance of the phenotype is zero. Yet, phenotypic evolution may stop under broader conditions with persistent selection and persistent MAG covariation in more complex examples (e.g., involving more traits or ages). Phenotypic evolution stopping with persistent selection is not possible in the Lande equation under the common assumption that its **G** matrix is non-singular, which means that evolution can only stop when selection vanishes.

We then have the system

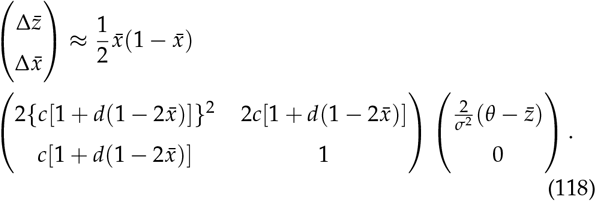

If we do not substitute for the mean genotype-phenotype map, we now have a system that is determined as it has two unknowns and two equations and that gives a description of long-term phenotypic evolution as an adaptive topography. It can thus be solved to obtain a unique evolutionary trajectory. The system is coupled: the dynamics of allele frequency depend on the mean phenotype and the dynamics of the mean phenotype depend on allele frequency. So both equations must be solved simultaneously. Since **L**_**m**_ is singular, the evolutionary trajectory and outcomes depend on the initial conditions, so evolutionary history matters. Solving the system with the initial condition for the mean phenotype given by the static developmental constraint, so 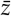 at the initial evolutionary time is given by (105), yields the evolutionary trajectories and outcomes found by solving the evodevo dynamics (110), shown in Fig. 3p-ad. Yet, adding noise to the initial condition of the phenotype yields different trajectories and outcomes due to the singularity of **L**_**m**_ (Fig. 4; see also phase portraits in González-Forero 2023 Fig. 1j,k,l).

### Example 4: evo-devo dynamics of two phenotypes influenced two biallelic loci under implicit development: evolution of ubiquitous imperfection

We now consider a scenario with two haplotype-symmetric phenotypes influenced by two biallelic loci under implicit development, for a large randomly mating sexual diploid population with non-overlapping generations. As before, we first analyse it with the exact method and then with the approximated one.

#### Exact method: constraints in higher dimensions

We begin by specifying our notation to the case of two loci. Let locus 1 have alleles a and A, and locus 2 have alleles b and B. Let 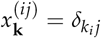 be the gene content in haplotype **k** = (*k*_1_, *k*_2_) ∈ ℍ = {ab, aB, Ab, AB} for allele *j* in locus *i* ∈ {1, 2}. That is, the gene content in haplotype **k** for allele A in locus 1 is 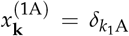 and the gene content in haplotype **k** for allele B in locus 2 is 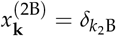, which yields the values in Table 1.

**Table 1.**
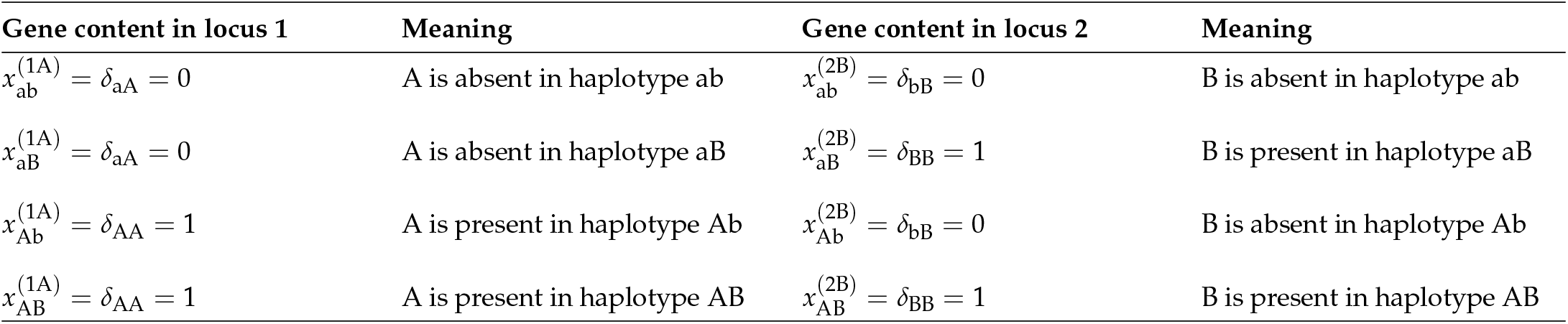
Gene content in two locus example.

Let *p*_**k**_ be the frequency of haplotype **k**. Then, the frequency of allele *j* in locus *i* is

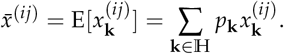

Specifically, the frequency of allele A is

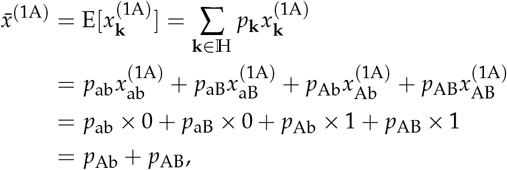

and the frequency of allele B is 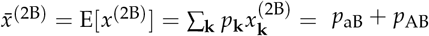.

Moreover, the frequency of haplotype **k** is given by

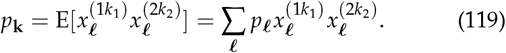

For instance, the frequency of haplotype ab is

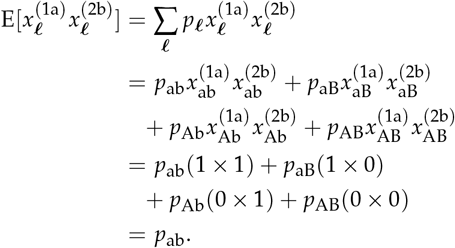

Hence, since E[*XY*] = E[*X*]E[*Y*] + cov[*X, Y*], the frequency of haplotype **k** is

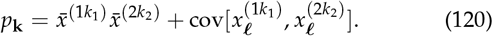

In SI section S8, we recover that the latter covariances involve only one degree of freedom called the coefficient of linkage disequilibrium:

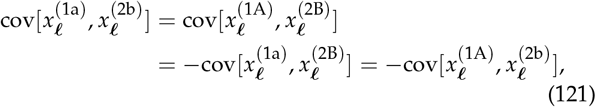

which is a previously known relationship (Nagylaki 1992, Eq. 8.69). The haplotype content deviation of haplotype **k** among the subset of loci {1, 2} of haplotype AB is

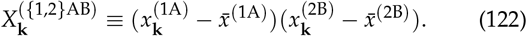

As we will be concerned with only one of these, for readability, we will drop the superscript and write *X*_**k**_ for 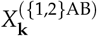. Hence, the expected value of this quantity is 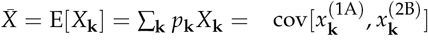, which is the coefficient of linkage disequilibrium. Then, we obtain the standard expressions for haplotype frequencies: 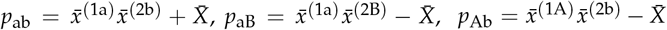, and 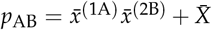 (Lewontin and Kojima 1960). It can be checked that 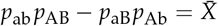.

As allele frequencies and linkage disequilibrium coevolve in multilocus systems, to obtain a dynamically sufficient system for allele frequency change, we build the vector **x**_**k**_ of haplotype content for haplotype **k** formed by the haplotype’s gene content at both loci and by the haplotype content deviation, namely, 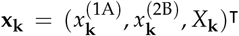. Hence, the mean haplotype content lists the allele frequencies at both loci and the linkage disequilibrium between loci: 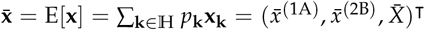. Applying the generator equations (19), the change in allele frequencies and linkage disequilibrium is given by

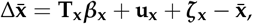

whose components we derive now.

For simplicity, we denote the frequency of allele A in locus 1 as 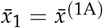 and of allele B in locus 2 as 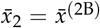. Then, the covariance matrix of haplotype content is

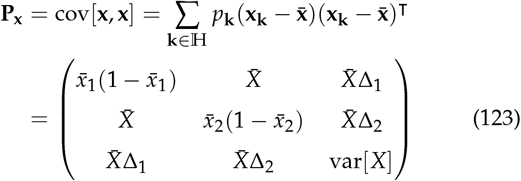

where the difference of allele frequencies in locus *i* is 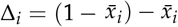 and the variance of haplotype content deviation is

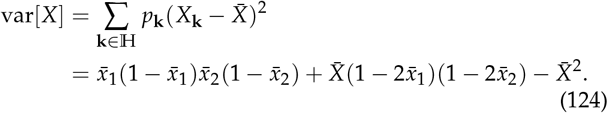

Note that if only allele frequency change were followed, the off-diagonal entries of such **P**_**x**_ would contain the linkage disequilibrium coefficient, so the system would be dynamically insufficient; consequently, we must follow the dynamics of linkage disequilibrium (Kimura 1956; Lewontin and Kojima 1960; Bodmer and Parsons 1962).

Next, we compute the conditional expectation of offspring haplotype content **x***′* (**x**_**k**_) given parental haplotype content **x**_**k**_.

The expected gene content in locus *i* for allele *j* among the offspring haplotypes given parental haplotype content **x**_**k**_ is

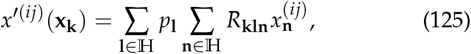

where *R*_**kln**_ is the probability that genotype **kl** produces gametes with haplotype **n**. This probability is given by:

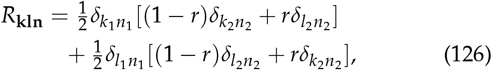

where *r* ∈ [0, 1/2] is the recombination frequency between the two loci and *δ*_*kl*_ is the Kronecker delta (eq. 8.9 of Nagylaki 1992). Similarly, the expected haplotype content deviation among the offspring haplotypes given parental haplotype content **x**_**k**_ is

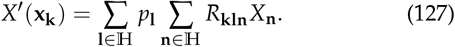

We now form the vector of expected offspring haplotype content conditional on parental haplotype content: **x***′* (**x**_**k**_) = (*x*^*′*(1A)^, *x*^*′*(2B)^, *X*^*′*^)^⊺^. Doing the indicated calculations, this vector takes the values

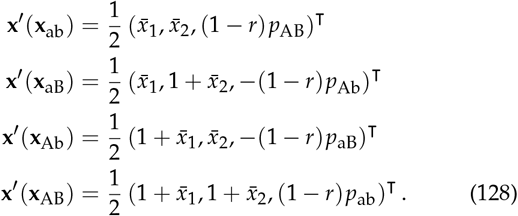

Hence, the mean offspring haplotype content is

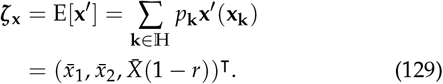

Pure transmission bias in haplotype content is then

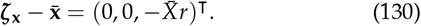

Thus, there is pure transmission bias in haplotype content if the two loci recombine (i.e., if *r* ≠ 0) and there is linkage disequilibrium (i.e., 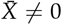).

Then, the transmission matrix of haplotype content is

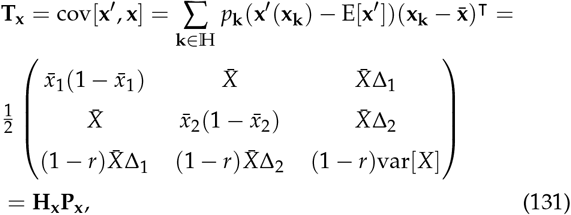

where the heredity matrix is

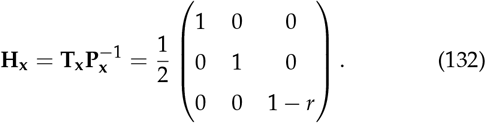

The transmission matrix **T**_**x**_ is asymmetric but it can be checked that the eigenvalues of its symmetric part are non-negative, given the constraint that 0 ≤ *p*_**k**_ ≤ 1 for all **k** ∈ ℍ. Hence, the selection response from linear inheritance points in the direction (≤ 90*◦*) of the selection regression coefficient of the haplotype, ***β***_**x**_. Moreover, the determinant of 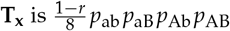, so **T**_**x**_ is invertible if all haplotype frequencies are non-zero, in which case the selection response from linear inheritance vanishes if and only if ***β***_**x**_ = **0**.

We next find that there cannot be selection response of haplotype content from non-linear inheritance, because Mendelian genetics is linear. From (3), we have that the residual of regressing expected offspring haplotype on parent haplotype is 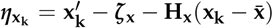. Doing this calculation yields that 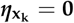 for all **k** ∈ ℍ, so the expected offspring haplotype is exactly given by its linear regression on parent haplotype content. Hence, from (5), there cannot be selection response of haplotype content from non-linear inheritance, that is, **u**_**x**_ = **0**. Also, since 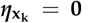, the covariance of parent-offspring residuals of haplotype content is zero, **U**_**x**_ = cov[***η***_**x**_, ***η***_**x**_] = **0**, which using (18) entails that the covariance matrix of expected offspring haplotype is 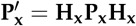. We can also obtain the covariance matrix of expected offspring haplotype from its definition:

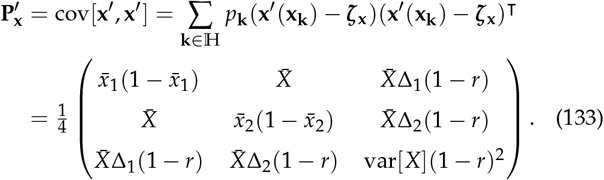

The selection regression coefficient of haplotype content is

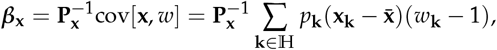

where *w*_**k**_ = ∑_**l** ∈ℍ_ *p*_**l**_ *w*_**kl**_ is the fitness of haplotype **k** and *w*_**kl**_ is the fitness of genotype **kl**, for **k** ∈ ℍ and **l** ∈ ℍ. Doing this calculation yields

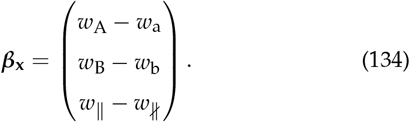

where the allele-specific fitnesses are

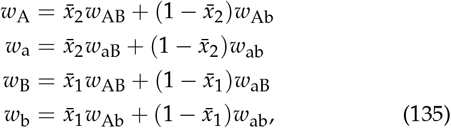

the fitness of the coupling haplotypes is *w*_∥_ = *w*_ab_ + *w*_AB_ and the fitness of the repulsion haplotypes is *w*_∦_ = *w*_aB_ + *w*_Ab_. I am unaware of an expression for the regression of fitness on haplotype content for two biallelic loci under linkage disequilibrium, as in (134). This expression suggests that a generalisation to more loci might be tractable (recall that for one biallelic locus we have the classic *β*_*x*_ = *w*_A_ − *w*_a_, Eq. (94a)).

Mean absolute fitness is

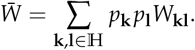

Then, assuming that absolute fitness *W*_**kl**_ is independent of allele frequency 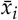 and of linkage disequilibrium 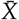 and that fitness is haplotype-symmetric *W*_**kl**_ = *W*_**lk**_, it can be checked that the selection regression coefficient of haplotype content is

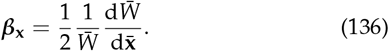

The adaptability of haplotype content is then

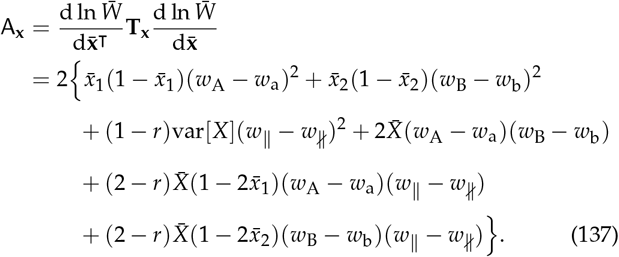

Since 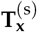 is positive semi-definite, this adaptability is non-negative and selection cannot decrease fitness here.

The change in allele frequencies and linkage disequilibrium then becomes

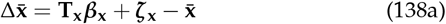

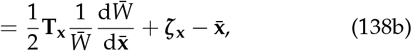

where, from (130), pure transmission bias is 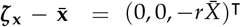. Eq. (138b) is a gradient equation describing the exact dynamics of allele frequency and linkage disequilibrium in two loci, which to my knowledge remained unavailable (Walsh and Lynch 2018, p. 127-128). This equation simplifies substantially if linkage disequilibrium is absent, with **T**_**x**_ becoming diagonal and pure transmission bias becoming zero, which suggests that a generalisation of this equation to arbitrarily many biallelic loci under linkage equilibrium might have a simple structure. Eqs. (138) are rather simple, enabling analytical insight given (130), (131), and (134). In contrast, the analogous equations under the haplotype frequency formulation are rather complex, limiting analytical insight (SI section S9.3). Moreover, Eqs. (138) are not particular cases of the Lande equation, including because **T**_**x**_ is asymmetric and there is pure transmission bias.

Some implications follow. Since 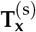 is positive semi-definite, the selection response of mean haplotype content cannot decrease mean fitness. Thus, the Moran (1964) phenomenon of mean fitness decrease with two loci must be due to pure transmission bias with both recombination and linkage disequilibrium. Note also that because of pure transmission bias (with 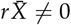), at evolutionary equilibrium we have 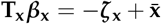. So, at equilibrium and if all haplotype frequencies are non-zero, we have 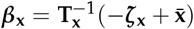, which may be different from zero. If at least one haplotype frequency is zero, then **T**_**x**_ is singular, in which case if there is any equilibrium, there is an infinite number of them, only one of which involves a zero selection regression coefficient. Then, in general a local mean fitness peak with respect to mean haplotype content may not be reached. For specific situations where linkage disequilibrium occurs at stable equilibria in this scenario, see Nagylaki (1992), p. 188.

So far in this example we have only considered genetic evolution. To analyse phenotypic evolution, suppose each individual has two phenotypes influenced by the two loci with analogous genotype-phenotype maps as in examples 2 and 3, but with contributions from both loci. Then, let *z*_**kl***m*_ be the *m*-th phenotype of an individual with genotype **kl**, having the genotype-phenotype map:

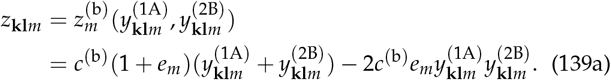

Here, 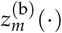 is a function that maps contributions between loci to the *m*-th phenotype, 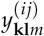 is the contribution of locus *i* to the *m*th phenotype, *e*_*m*_ is the epistasis coefficient between the loci for that phenotype, and *c*^(b)^ is a scaling factor. Let the contribution of locus *i* be defined analogously as

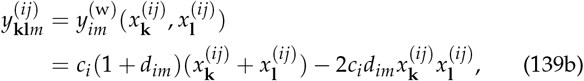

where 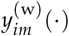 is a function that maps the contribution of locus *i* to phenotype *m, d*_*im*_ is the dominance-pleiotropy coefficient for locus *i* and phenotype *m* (pleiotropy is thus generally present in this model), and *c*_*i*_ is a scaling factor for the contribution of locus *i*. The phenotypic vector for an individual of genotype **kl** is **z**_**kl**_ = (*z*_**kl**1_, *z*_**kl**2_)^⊺^ (so, the genotype-phenotype map is **z**(·) = **z**^(b)^ *◦* **y**^(w)^(·), or for the *m*-th phenotype, it is 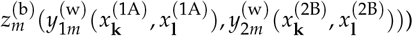. As before, the mean phenotype is

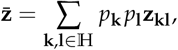

and the phenotypic covariance matrix is

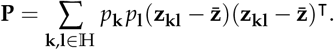

Consider the case of additive allelic effects between loci on phenotype 1 and multiplicative allelic effects between loci on phenotype 2 with additive allelic effects within loci (*e*_1_ = 0, *e*_2_ = −1, and *d*_11_ = *d*_12_ = *d*_21_ = *d*_22_ = 0, with *c*^(b)^ = *c*_1_ = *c*_2_ = 1). Then, the static developmental constraint for phenotype 1 reduces to

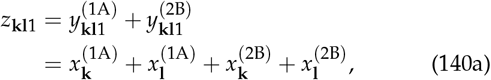

so phenotype 1 is fully additive. In turn, the static developmental constraint for phenotype 2 reduces to

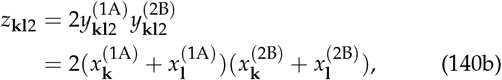

so phenotype 1 is additive within loci and multiplicative between loci. Consequently, the mean phenotype is

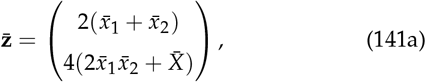

and the phenotypic covariance matrix is

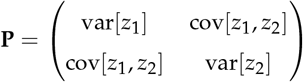

with

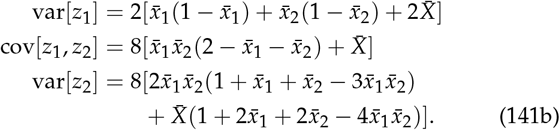

The genotype-phenotype map (140) produces the individual phenotypes, the distribution of phenotypes, and the admissible manifold for mean phenotypes plotted in Fig. 5. Thus, only 6 phenotypic values can be developed and the mean phenotype can only take values in a restricted region that changes as linkage disequilibrium evolves.

**Figure 4.**
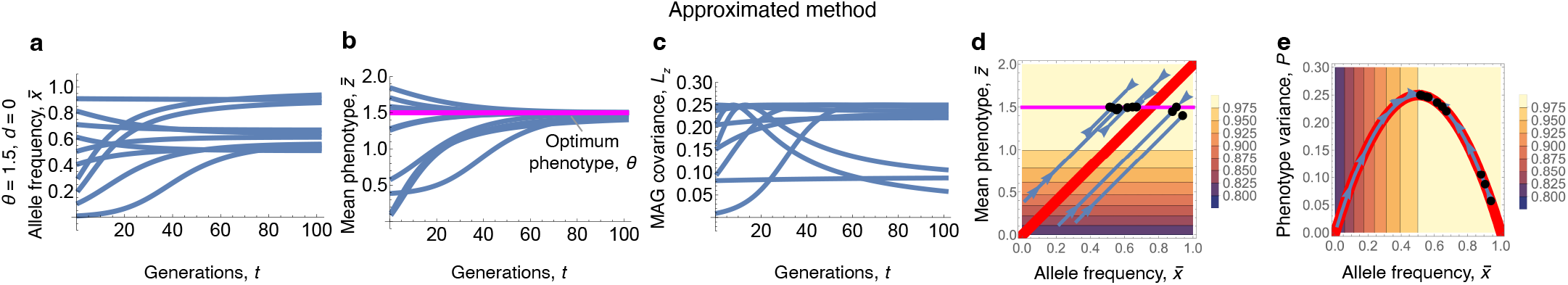
Absolute constraints on adaptation entail that evolutionary outcomes depend on ancestral conditions. Here we add developmental stochasticity to the ancestral conditions of Fig. 3p-t. Plots are the solution of Eq. (118), setting the ancestral condition of the phenotype to 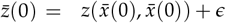, where *ϵ* is a uniformly distributed random real number between −0.5 and 0.5. Parameter values are as in Fig. 3p-t. **a-c**, Evolutionary trajectories and outcomes of allele frequency and phenotype variance differ from Fig. 3p-r. **d**, In red is the developmental constraint (admissible evolutionary manifold) under the initial condition 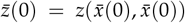. In **d**, the evolutionary trajectories are parallel to such developmental constraint (red) due to the singularity of **L**_**m**_. There is an infinite number of equilibria along the phenotype optimum (magenta). The equilibrium attained is the intersection of this line of equilibria with the developmental constraint, which has been perturbed by the ancestral condition. The evolutionary outcome of the mean phenotype does not depend on the ancestral condition in this example, but this is only due to the simplicity of the example. For further illustration, see phase portraits in González-Forero (2023) Fig. 1j,k,l and different mean phenotype outcomes in González-Forero (2024a).

**Figure 5.**
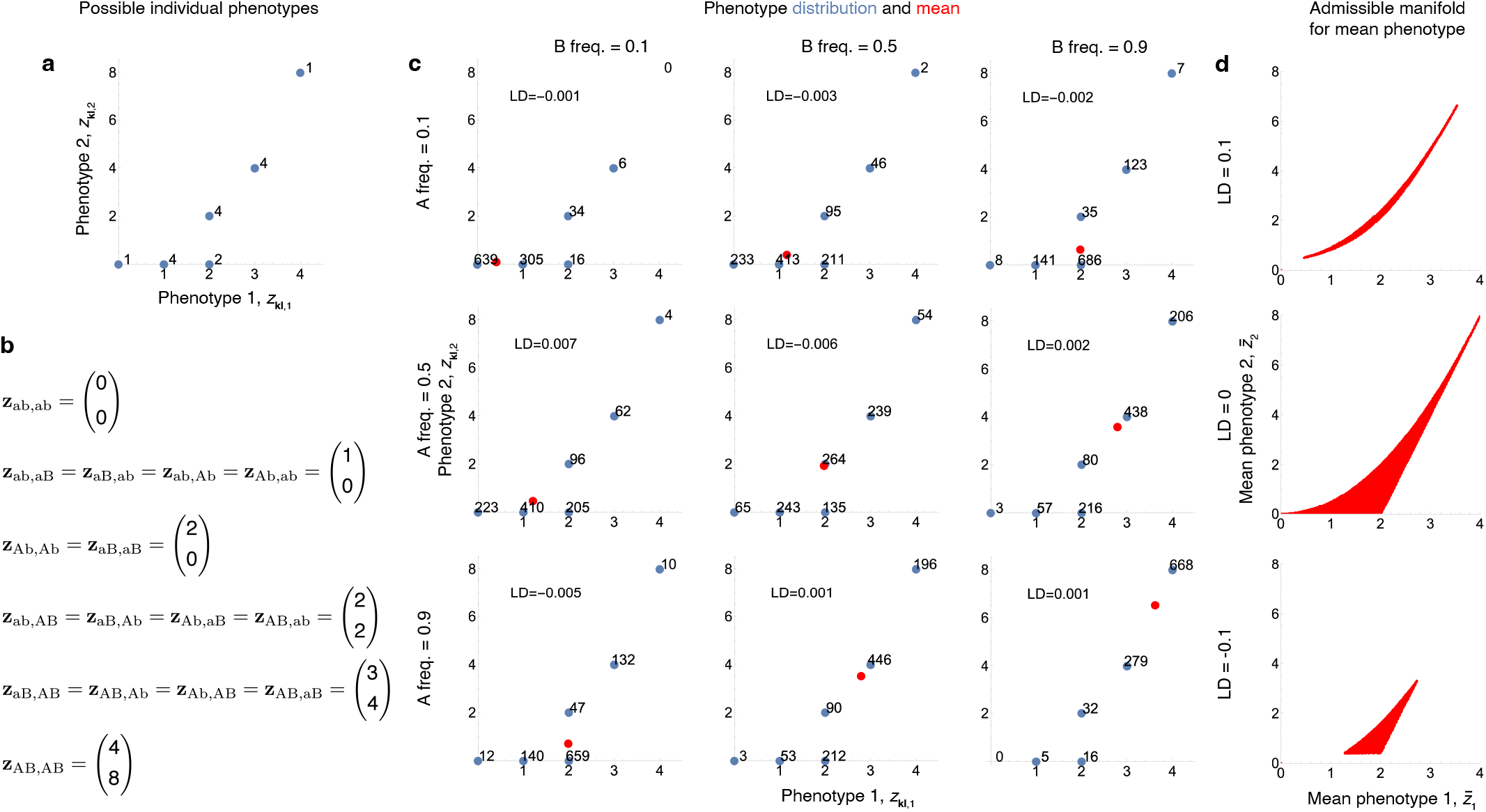
Genotype-phenotype map for example 4 and resulting admissible manifold for mean phenotype. **a**, Possible phenotypes, with numbers indicating the number of genotypes that produce the corresponding phenotype. **b**, Phenotypes produced by the different genotypes. **c**, Distribution of phenotypes that result when sampling 1000 genotypes, each allele sampled from a Bernoulli distribution with parameter given by the allele frequency indicated in the rows and columns of the panels. Blue dots give the resulting phenotypes with numbers denoting the number of genotypes with the corresponding phenotype. Red dots give the average phenotype. The inset text gives the resulting linkage disequilibrium (LD, or 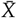) in the corresponding panel. **d**, Admissible manifold for mean phenotype. The red region gives all possible points where the mean phenotype can occur for any combination of allele frequencies and for the indicated linkage disequilibrium, under the constraint that haplotype frequencies are between zero and one. Thus, linkage disequilibrium changes the size, shape, and position of the admissible manifold.

Let the absolute fitness of genotype **kl** be Gaussian: 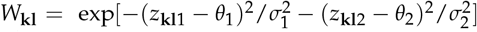. Numerical results of the equation (138a) describing change in allele frequency subject to the static developmental constraints (140) are in Fig. 6a-h. For the parameter values used, allele frequencies converge to the mean fitness peak with respect to mean haplotype content (Fig. 6a). Mean phenotypes are constrained to evolve within the developmental constraint, which is the admissible manifold (red region) that is given by the genotype-phenotype map under any combination of allele frequencies and linkage disequilibrium provided that haplotype frequencies are between zero and one (Fig. 6b). Whereas the admissible manifold was one dimensional path in the one locus examples above, the admissible manifold is here three dimensional because **x** is three dimensional: two loci and one linkage disequilibrium; for visualisation, we plot this manifold in two dimensions by evaluating linkage disequilibrium at initial and final times. The outcome of phenotypic evolution is a point on the admissible manifold that is closest to the optimum phenotype (Fig. 6b). Yet, as linkage disequilibrium evolves, the admissible manifold moves away from rather than towards the optimum phenotype. Phenotypic variation does not disappear despite stabilising selection (Fig. 6h), in contrast to quantitative genetics (Barton and Turelli 1987) (and our example 1) but consistently with previous models of two loci with additive effects (Gale and Kearsey 1968; Bürger 2000). Simply put, individuals vary, but nobody is ever perfect. A substantially higher phenotypic variance is maintained for the non-additive phenotype, even after standardizing the phenotypic variances by squared phenotype means. Despite the active developmental constraint, phenotypic covariances are strongly positive, emphasizing that constraints or trade-offs are not revealed by negative phenotypic or genetic correlations (Roff and Fairbairn 2007; Houle 1991).

#### Approximated method: simplified constraints

The above exact method describes genetic evolution as an adaptive topography, but does not describe phenotypic evolution as an adaptive topography. To examine whether *phenotypic* evolution can here be understood as an adaptive topography and gain further analytical insight, consider now the approximated method applied to example 4, where two phenotypes are influenced by two loci.

Assuming small haplotype variation and weak selection and since the phenotype is here haplotype-symmetric, from (45), the change in mean haplotype content is to first order

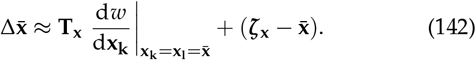

We have already derived **T**_**x**_ and ***ζ***_**x**_, so let us derive the total selection gradient of haplotype content. It is given by

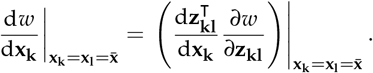

As before, letting alleles within loci have additive effects (i.e., *d*_*im*_ = 0 for the two loci *i* and two phenotypes *m*), the total effects of the haplotype on the phenotype are

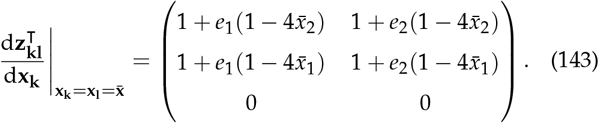

The row of zeros arises because the genotype-phenotype map does not depend on the haplotype content deviation, *X*. It can be checked that postmultiplying the matrix (143) by a column vector (*a, b*)^⊺^, setting the result to zero, and solving for *a* and *b* does not imply that *a* = *b* = 0 if 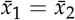 or if *e*_1_ = *e*_2_, in which case the matrix (143) is non-singular. Hence, the total selection gradient of the haplotype may vanish with non-zero phenotypic selection if allele frequencies are the same at both loci 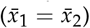 or if epistasis is the same for both phenotypes (*e*_1_ = *e*_2_). Letting there be no epistasis for the first trait and multiplicative epistasis for the second trait (i.e., *e*_1_ = 0 and *e*_2_ = −1), the total effects further reduce to

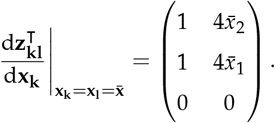

Letting fitness be 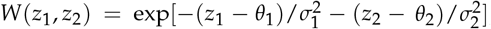, mean fitness is 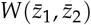 to first order, where the mean phenotypes are to first order

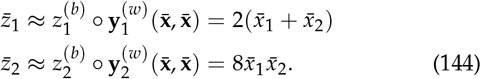

Thus, to first order, the mean phenotype only depends on the allele frequencies at the two loci but does not depend on linkage disequilibrium, so the admissible manifold is to first order two dimensional rather than three dimensional. To first order, the selection gradient of the phenotype is

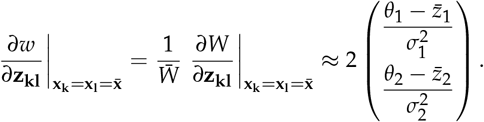

Hence, total genetic selection reduces to

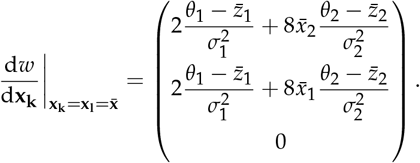

Setting this total selection gradient to zero yields two internal equilibria. First, one equilibrium at the optimum phenotype. Second, another equilibrium when allele frequencies are the same at both loci 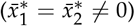 and the mean phenotype 2 is

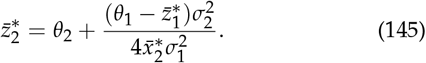

This equilibrium is thus a line of equilibria in allele frequency space (where 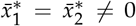) and another line of equilibria in mean phenotype space, where (145) holds.

This approximation is qualitatively accurate relative to the exact method (Fig. 6i-r). As in the exact method, this system never reaches the optimum phenotype, even though mechanistic additive genetic covariation is maintained for both phenotypes, particularly for the non-additive phenotype (Fig. 6j,p). Mechanistic additive genetic covariation in the direction of selection vanishes, because fitter mean phenotypes cannot be produced by the developmental process: mechanistic additive genetic covariation becomes orthogonal to the direction of selection (at 90 degrees) (Fig. 6q,r). The outcome is at a mean fitness peak within the admissible manifold, that is, a locally highest point in the fitness landscape within the admissible region (Fig. 6j). As in the exact method, individuals vary and nobody is perfect in the end.

To analyse phenotypic evolution as the climbing of a fitness landscape, from (58) we have the approximated gradient equation for the phenotype:

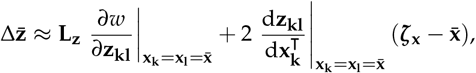

where 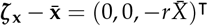 and

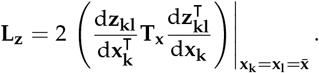

For the scaling and dominance coefficients used, we have that det 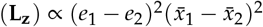, so **L**_**z**_ is singular if allele frequencies are the same in both loci 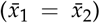 or if epistasis is the same for both phenotypes (*e*_1_ = *e*_2_). Also, this **L**_**z**_ is symmetric and does not depend on the recombination frequency *r*, despite **T**_**x**_ being asymmetric and depending on *r*. The reason is that the row of zeros in (143) makes **L**_**z**_ independent from the third column and third row of **T**_**x**_; recall this row of zeros arises because the phenotype does not depend on the haplotype content deviation. For *e*_1_ = 0 and *e*_2_ = −1, the matrix is

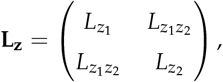

where

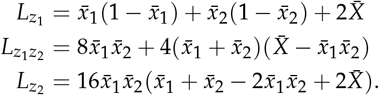

Since this matrix is symmetric, the dynamics are characterised by whether it is positive semi-definite, which it is, given the constraint that haplotype frequencies are between zero and one. Thus, this recovers the exact result that selection response cannot decrease mean fitness in this example. Yet, as before, this equation for phenotypic evolution as an adaptive topography is dynamically insufficient because it depends on allele frequency and linkage disequilibrium but does not describe their change.

If we track both phenotypic and genetic evolution, then from (62) the evolutionary change is to first order given by

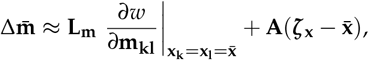

where

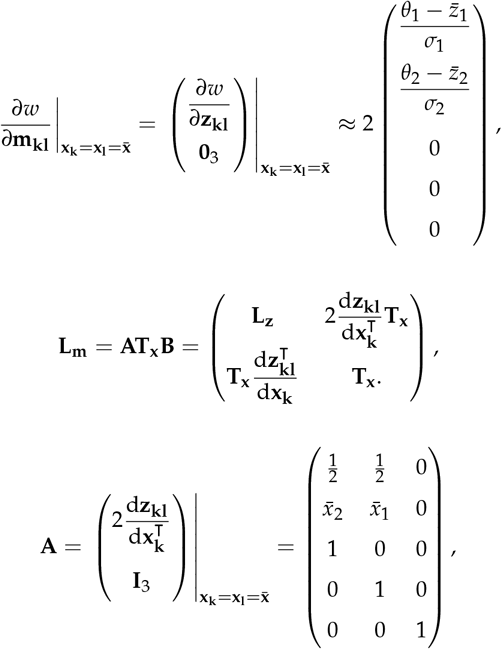

and

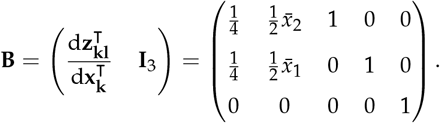

This system is now dynamically sufficient as it contains as many equations as unknowns, so it describes long-term phenotypic and genetic evolution to first order of approximation. The matrix **L**_**m**_ is singular and so direct selection can persist despite vanishing pure transmission bias, as observed numerically (Fig. 6j). Indeed, selection eventually aligns with the direction of no MAG covariation and becomes orthogonal to the direction of most MAG covariation (Fig. 6q). Since there is no direct selection on haplotype content, adaptability is determined by **L**_**z**_ rather than **L**_**m**_. The matrix **L**_**z**_ is symmetric and positive semi-definite because the haplotype content deviation does not affect the phenotype, so adaptability is non-negative. So we recover the above exact result that selection cannot decrease mean fitness in this approximation. Hence, if a Moran phenomenon is obtained over long-term evolution with this system, it must also be due to pure transmission bias.

## Discussion

### Overview

We derived general mathematical theory that integrates genetics, development, and evolution, specifically where the infinitesimal model is relaxed to describe arbitrarily complex development and is coupled with multilocus genetic inheritance, selection, and evolution. The theory thus presents a generalised understanding of phenotypic variation, inheritance, and evolution. This theory yields a principle of developmentally constrained phenotypic adaptation that contrasts with the notion of developmentally unconstrained phenotypic adaptation that is provided by quantitative genetics theory under the infinitesimal model (Fig. 7; see also González-Forero 2023). We showed that this constrained phenotypic adaptation occurs in the simplest population genetics models and carries over to arbitrary complexity, where the constraints do not disappear under the infinitesimal model but become inactive as the admissible manifold covers all the phenotypic space. Overall, the theory finds that development affects evolution by shaping the total (i.e., genetic) fitness landscape and by constraining evolution to an admissible manifold where mean phenotypes and higher moments can be developed. This entails that development plays a major evolutionary role, including by co-defining with selection the evolutionary outcomes. These basic evolutionary effects of development had been identified before using simplified genetics (González-Forero 2023, 2024b) and are recovered here with realistic genetics.

**Figure 6.**
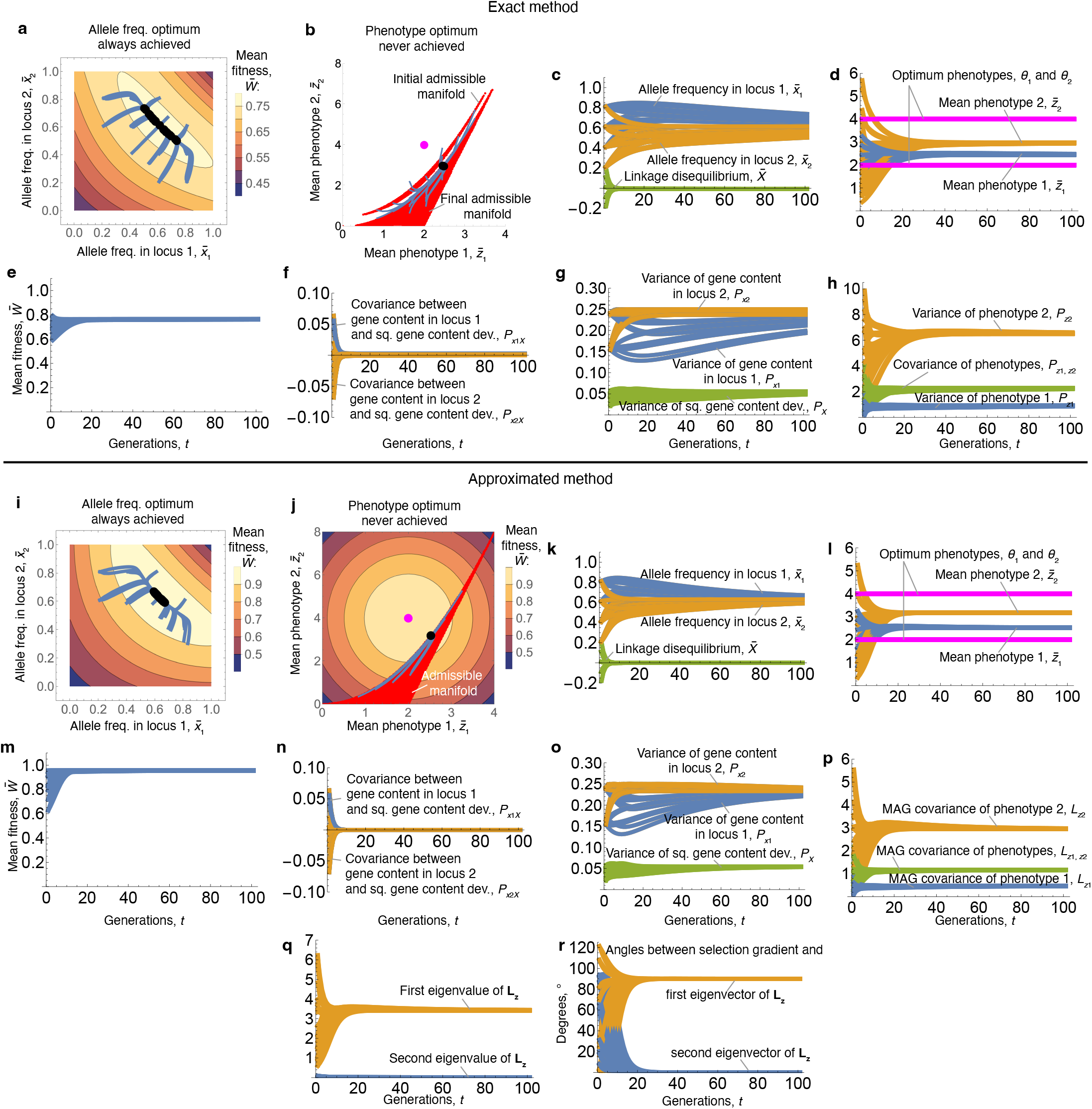
Example 4: Evolution of two phenotypes influenced by two biallelic loci under implicit development. **a-h**, Exact method. **a**, Allele frequency change over the contour of the (total) fitness landscape vs allele frequency. The values at final evolutionary times are black dots. **b**, Mean phenotype change over the admissible evolutionary manifold (red), which is all the mean phenotypes that would result with any allele frequencies and linkage disequilibrium. The manifold is three dimensional but it is two-dimensional in the plot because there are two loci, and one linkage disequilibrium coefficient evaluated at the initial and final times. The initial and final manifolds are for initial conditions 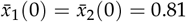 and 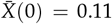 (chosen for illustration; other initial conditions yield slightly different initial and final admissible developmental manifolds, which can be seen by some trajectories falling outside the initial manifold plotted). The optimum phenotype is in magenta. Parameter values: *r* = 0.5, *e*_1_ = 0, *e*_2_ = −1, *d*_11_ = *d*_12_ = *d*_21_ = *d*_22_ = 0, *c*^(b)^ = *c*_1_ = *c*_2_ = 1, *θ*_1_ = 2, *θ*_2_ = 4, 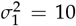, and 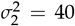. Initial conditions are all combinations of 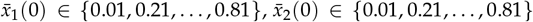, and 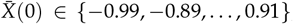, if the initial frequencies of all haplotypes **k** satisfy 0 ≤ *p*_**k**_(0) ≤ 1. In **a**, mean fitness is evaluated at zero linkage disequilibrium 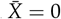. **i-r**, Approximated method. Panel descriptions are as in **a-h**, except that the approximations 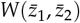 for mean fitness and 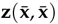 for mean phenotype are used, as well as the following differences: **j**, The contour is the approximated mean fitness 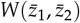, which is now a function of the mean phenotype. The admissible manifold (red) is the approximated mean phenotypes 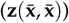 under all possible allele frequencies given the constraint that haplotype frequencies are between zero and one. **p**, Mechanistic additive genetic (MAG) covariances **L**_**z**_. **q**, Eigenvalues of **L**_**z**_, giving how much MAG covariation changes selection in the direction of the corresponding eigenvectors of **L**_**z**_; selection in the direction of the second eigenvector is largely nullified. **r**, Angles in degrees between the selection gradient 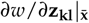 and the eigenvectors of **L**_**z**_ associated to the eigenvalues in **q**; selection aligns with the direction of no MAG covariation and becomes orthogonal to the direction of most MAG covariation.

**Figure 7.**
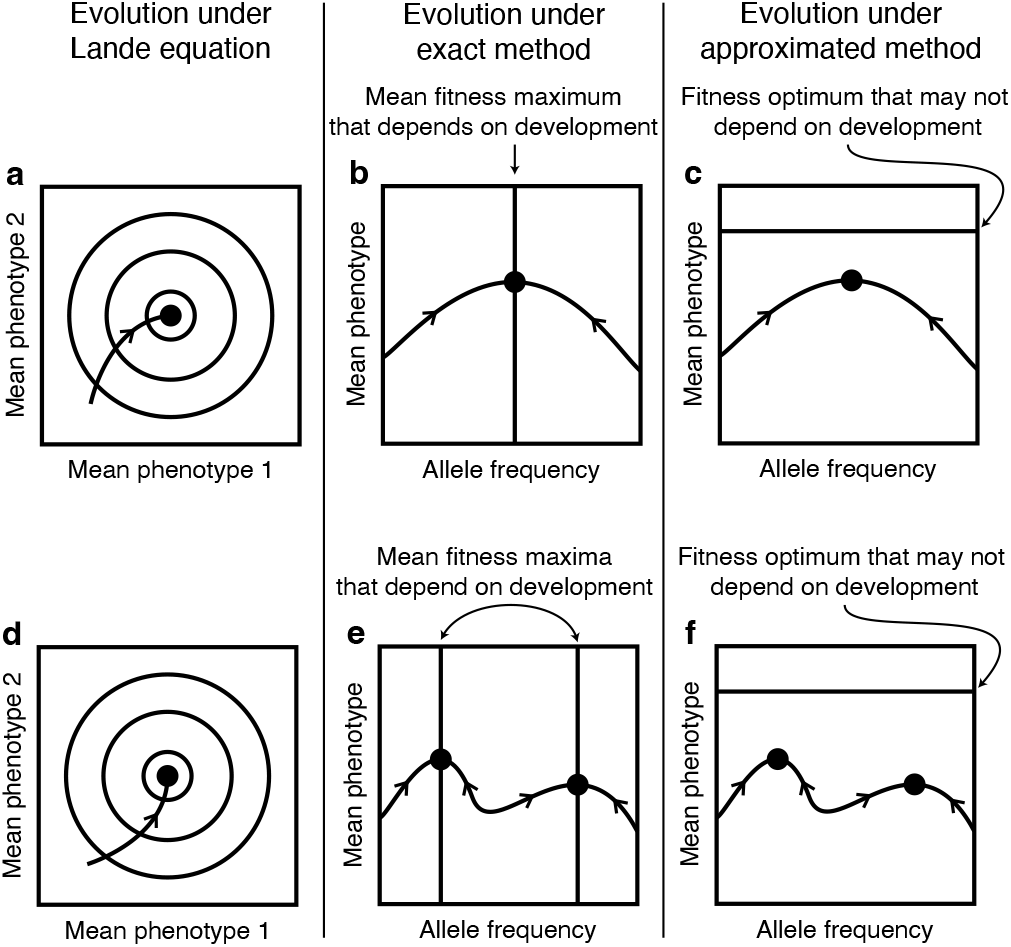
Evo-devo adaptation. **a**, Schematic description of adaptive phenotypic evolution under the Lande equation in gradient form, where the evolutionary space is the mean phenotype and the local optimum of the mean (multivariate) phenotype is reached (using the typical assumption that the **G** matrix is non-singular). Circles are mean fitness contours, with mean fitness increasing toward the center. **b**, Schematic description of adaptive phenotypic evolution under our exact method describing evolutionary and developmental dynamics. The curve is the mean genotype-phenotype map, which relates allele frequency to mean phenotype. This curve gives the developmental constraint and evolution is constrained to occur within that path (admissible evolutionary manifold). In the exact method, mean fitness is in general a function of allele frequency rather than of mean phenotype; I refer to this as the total fitness landscape of the mean haplotype. Mean fitness has a local peak at the straight vertical line and decreases away from it. If 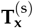 is positive definite and there is no pure transmission bias for the haplotype, the evolution of mean phenotypes increases mean fitness, so evolution converges to the path peak (black dot). **c**, Schematic description of adaptive phenotypic evolution under the approximated evolutionary and developmental dynamics. To first-order of approximation, mean fitness is now a function of the mean phenotype; I refer to this as the direct fitness landscape of the mean phenotype. Thus, mean fitness has a local peak at the straight horizontal line and decreases away from it. If 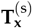 is positive definite and there is no pure transmission bias for the haplotype, evolution converges to the path peak (black dot). These panels illustrate why development is of minor evolutionary importance under the Lande equation, but is of major importance under our evo-devo dynamics approach. Specifically, in **d**, a change in development relative to **a** affects genetic correlations and so the transient trajectory, but not the outcome (black dot), which is independent of the initial conditions and of development. In contrast, in **e,f**, a change in development relative to **b,c** affects the developmental constraint (curve) and the total fitness landscape (so vertical lines in **e**), thereby affecting the outcomes (black dots), which depend on the initial conditions and on development.

To integrate realistic genetics, we first obtained general generator equations that describe change in multivariate traits, generalising the general form of the Lande equation (Lande and Arnold 1983) including with a transmission matrix **T** identified by Rice (2004b) (his Eq. 7.13) that generalises the matrix of additive genetic covariation, **G**, to any form of inheritance. In particular, **T** allows for non-normal distributions of the covariates involved, non-additive genetic inheritance, non-genetic inheritance, and organismal maladaptation. The latter is because, in contrast to **G, T** is not a covariance matrix but a cross-covariance matrix so it is not necessarily symmetric. Consequently, **T*β*** can point away (i.e., at an angle > 90*◦*) from ***β***, whereas **G*β*** cannot.

The generality of the generator equations allowed them to be applied to describe change in allele frequencies and linkage disequilibrium, and so to describe long-term phenotypic evolution as a response to genetic evolution. This application provides analytical expressions of multilocus evolution as a modified adaptive topography, which remained unavailable to my knowledge. The example with two loci afforded novel analysis of the Moran (1964) phenomenon of mean fitness decrease in two loci, a phenomenon that is not possible with the Lande equation.

We derived approximated gradient equations describing long-term genetic and phenotypic evolution as modified, constrained adaptive topographies, to first order of approximation. The approximated gradient equations have similar forms to the gradient form of the Lande (1979) equation, but with important differences including that the associated mechanistic additive genetic cross-covariance matrix may be asymmetric and that the matrix is always singular under haplotype-symmetric phenotypes in the long-term evolutionary system describing both genetic and phenotypic evolution. That the matrix is asymmetric entails that selection may yield transient or permanent organismal maladaptation, with potentially complex dynamics that are forbidden by the Lande equation in gradient form, such as saddle points at fitness peaks, evolutionary cycles, and chaos.

That the matrix is singular entails that evolutionary outcomes depend on ancestral conditions even with a single fitness peak and that development plays a major evolutionary role by co-defining with selection the evolutionary outcomes, neither of which is the case with the Lande equation in gradient form, particularly under the typical assumption of non-singular **G**. The evolutionary relevance of development arises because it determines which phenotypes can be developed, so it influences both the admissible evolutionary manifold and the total selection gradient of the haplotype, which determines when the selection response vanishes. These qualitative features hold both in the exact and approximated methods, but the approximated gradient equations facilitate analyses.

Thus, this work yields a theory of constrained evolutionary dynamics, where development imposes necessarily absolute constraints on adaptation. In this theory, evolution proceeds within the admissible evolutionary manifold determined by development. This manifold may have determinant evolutionary effects, for instance, by forcing phenotypic variances to increase while mean fitness increases (example 3).

This constrained adaptation contrasts with quantitative genetics under the infinitesimal model, where there are effectively no active constraints and so stabilising selection leads to the local optimum mean phenotype and depletes variation. The key reason for the discrepancy is that by assuming normal distributions, or an infinite number of loci contributing additively to the phenotype, quantitative genetics under the infinitesimal model entails that development has no active constraints. Indeed, the infinitesimal model entails that any phenotype and any phenotype combination can be developed, from minus infinity to plus infinity. Accordingly, previous results have found that increasing the number of loci that additively affect a single trait under stabilising selection rapidly decreases the variation that is maintained (Bürger and Gimelfarb 1999). As the number of loci increases, the phenotype space covered by the admissible evolutionary manifold expands without bounds given additivity and that only one trait is considered. This yields no restrictions for selection, producing monomorphic, perfect phenotypes under stabilising selection. The infinitesimal model is relaxed in the present work, where phenotypes can be of any form including through the construction of arbitrarily complex developmental dynamics, yielding contrasting results so the admissible evolutionary manifold does not necessarily cover all the evolutionary space. This is consistent with previous results where increasing genetic variation is maintained as the number of loci increases under epistasis (Gimelfarb 1989) and substantial genetic variation is maintained for arbitrarily many additively acting loci under pleiotropy (Gimelfarb 1996). Our results thus help unify these previously contrasting results suggesting that the developmental or genotype-phenotype map can constrain stabilising selection so that it leads to variable, possibly imperfect phenotypes.

One might expect that as many more loci are considered, normal distributions are approached and developmental constraints become inactive. Illustration that this is not necessarily the case is provided by a model applying the version of this theory under simplified genetics, where over 1400 loci influence the non-linear developmental dynamics of four phenotypes and the developmental constraints are still determinant (González-Forero 2024a). In that model, changing parameters that affect the developmental dynamics but not selection leads to the evolution of the brain and body sizes of seven different hominins, from small-brained Australopiths to large-brained Neanderthal and *Homo sapiens*. These seven different evolutionary outcomes occur in that model by changing developmental constraints alone, without changing direct selection, so the constraints are not only determinant there but of key explanatory role, despite phenotypes being influenced by over 1400 loci.

### Relationship to previous theory

The present theory departs from previous theory in various ways. On the one hand, the theory offers an extension of population genetics to describe both genetic and phenotypic evolution considering arbitrarily complex development. On the other hand, the theory relaxes the assumption of the infinitesimal model, which has been at the core of quantitative genetics. Under small haplotype variation and weak selection, the present theory recovers objects that resemble those of quantitative genetics and are labelled here with “mechanistic” qualifiers (specifically, mechanistic breeding values and mechanistic additive genetic cross-covariance matrices). However, by relaxing the infinitesimal model to describe developmental dynamics and by describing genetic evolution, the present theory departs from quantitative genetics in important ways. For instance, the theory allows for not only quantitative but also discrete or meristic phenotypes, for example, if the developmental dynamics display bifurcations, leading to distinct developmental outcomes (e.g., digit numbers) with slight changes in gene content or in parameters affecting the developmental dynamics. Moreover, whereas quantitative genetics seeks to describe phenotypic evolution without describing genetic evolution, the theory here explicitly considers genetic evolution to describe long term evolution. Furthermore, the objects recovered here that resemble those of quantitative genetics have importantly different properties from those of quantitative genetics and so should not be confused. So, to avoid confusion, I have referred to the present theory as evo-devo dynamics, given its features distinguishing it from standard population and quantitative genetics. As outlined in the introduction, this theory has long roots that can be traced back at least to Alberch *et al*. (1979), Wagner (1984), Slatkin (1987), Rice (2002), Parvinen *et al*. (2013), and Morrissey (2015). A mathematical integration of arbitrarily complex developmental dynamics and evolution under simplified genetics was previously available (González-Forero 2024b), and the present theory offers an additional integration to realistic genetics.

### Limitations

Some limitations are the following. Although the generator equations apply to any form of inheritance, the exact method considered only genetic inheritance. The generator equations can be used to extend the method to non-genetic inheritance. Moreover, although the generator equations and the exact method apply under stochastic development and endogenous or exogenous environmental change, the approximated method is so far genetically deterministic, as it considered deterministic genotype-phenotype and developmental maps without endogenous or exogenous environmental change. Consequently, in the approximated method, knowing an individual’s genes and genotype-phenotype or developmental map implies one knows the individual’s phenotype. This is a strong simplification that should be relaxed in the near future. Some relaxation of these simplifications is available in work formulating evo-devo dynamics under simplified genetics but allowing for endogenous and exogenous environmental change (González-Forero 2024b).

The approximated part of the theory involves a choice of continuously differentiable maps among infinitely many possible ones, and this choice affects how well the approximation works. Due to this and the difficulty of meeting the assumption of small haplotype variation with discrete-valued haplotypes, the approximate method has modest success replicating the exact dynamics. At present, it is not straightforward to anticipate when the approximated method is reliable. For instance, correct results can arise with both linear (example 3, Fig. 3f-j,u-y) and non-linear development (example 4, Fig. 6) and incorrect results may arise with both linear (example 3, Fig. 3a-e,p-t) and nonlinear development (example 3, Fig. 3k-o,z-ad), despite fitness being non-linear in all cases. Ideally, the approximations should for now only be relied upon when it has been confirmed that they do recover the exact dynamics for the particular situation examined, at least qualitatively.

The evo-devo dynamics theory previously formulated under simplified genetics involves analogous approximations to the approximated method derived here (González-Forero 2024b). Hence, it remains to be checked whether the application of such evo-devo dynamics to specific models (e.g., González-Forero 2024a) yields good approximations of what would result using the exact theory introduced here.

More broadly, our results suggest caution regarding modelling phenotypic evolution without describing genetic evolution. Such approaches include quantitative genetics, adaptive dynamics, and behavioural ecology as used in evolutionary game theory, social evolution, and inclusive fitness theory, often under the label of phenotypic gambit. Our observation that predictions can be substantially affected by considering genetic evolution and its mapping to phenotypic evolution suggests caution regarding reliance on purely phenotypic approaches.

### Applying the theory to data

The generator equations can be readily applied to empirical data to understand short-term evolution. The basic elements to be empirically estimated are parent vectors **z**, offspring vectors **z**^+^, the joint probability distribution of parent and offspring vectors *p*(**z, z**^+^), and the probability distribution of the conditional expectation of offspring vectors *p*_**z**_*′* (**z**). From these basic elements, all the components of the generator equations follow, including fitness. In particular, the transmission matrix **T** may be easier to estimate empirically than **G** as it does not require the isolation of additive genetic effects. Moreover, the empirical application of the generator equations may help improve predictions of shortterm change made with the breeder or Lande equations, which make more assumptions.

The application of evo-devo dynamics, both in its exact and approximate forms, to data to understand long-term evolution may benefit from taking a simulation-based inference approach (Cranmer *et al*. 2020), which has been characteristic of population genetics but less so of phenotypic evolution. This application requires that development is modelled; that is, that expressions for the genotype-phenotype or developmental maps are used, either from first principles or learned from data. This stands to benefit from rapidly advancing methods to infer genotype-phenotype and developmental maps (Martí-Gómez *et al*. 2026), including those that learn dynamic equations from data (Brunton *et al*. 2016; Course and Nair 2023; Prokop and Gelens 2026).

### Possible future applications

A number of empirical observations have been paradoxical in the light of previous theory such as the maintenance of genetic variation (Bürger 2000; Walsh and Lynch 2018; Charlesworth 2026), the paradox of stasis (Kingsolver and Diamond 2011; Kirkpatrick 2009; Merilä *et al*. 2001), the rarity of stabilising selection (Kingsolver *et al*. 2001; Kingsolver and Diamond 2011), and the paradox of predictability (Tsuboi *et al*. 2024). Reasons include that quantitative genetics under the infinitesimal model predicts that stabilising selection necessarily depletes variation, that evolution by selection converges to fitness peaks where directional selection vanishes and becomes stabilising, and that the effects of genetic variation on evolution are transient.

These predictions are not made by the theory presented. Specifically, by relaxing the infinitesimal model so phenotypes are possibly non-linear functions of gene content, we find that development imposes necessarily absolute constraints that may be active, so not every phenotype can be developed. Consequently, the current theory does not predict that stabilising selection necessarily depletes variation, nor that evolution converges to fitness peaks where directional selection vanishes and becomes stabilising, nor that the effects of genetic variation are transient. An important avenue for future work is to examine to what extent the present theory can account for the empirical observations that have been paradoxical with previous theory.

## Supporting information

Computer code

## Data availability

I affirm that all data necessary for confirming the conclusions of the article are present within the article, figures, and supplementary material.

## Acknowledgments

I thank Reinhard Bürger, Michael Lynch, Valeria Montano, Mihaela Pavličev, and Günter Wagner for discussion that helped improve the paper. I also thank Steven Frank, Sam Yeaman, and two anonymous reviewers for detailed comments and criticism of the manuscript that were very helpful to improve it greatly. I thank Kevin Lala (formerly Laland) for access to literature and to Mathematica. I thank the Konrad Lorenz Institute for Evolution and Cognition Research and its community for financial and scholarly support that made this paper possible.

## Literature cited

Alberch P, Gould SJ, Oster GF, Wake DB. 1979. Size and shape in ontogeny and phylogeny. Paleobiology. 5:296–317.

Alon U. 2020. An Introduction to Systems Biology. Taylor and Francis. Boca Raton, FL, USA. second edition.

Altenberg L. 1995. Genome growth and the evolution of the genotype-phenotype map, In: Banzhaf W, Eeckman FH, editors, Evolution and biocomputation, Springer-Verlag. volume 899 of Lecture Notes in Computer Science. pp. 205–259.

Arnold SJ. 1992. Constraints on phenotypic evolution. Am. Nat. 140:S85–S107.

Atchley WR, Hall BK. 1991. A model for development and evolution of complex morphological structures. Biol. Rev 66:101–157.

Barresi MJF, Gilbert SF. 2020. Developmental Biology. Oxford Univ. Press. Oxford, UK. 12th edition.

Barton NH, Etheridge AM, Véber A. 2017. The infinitesimal model: definition, derivation, and implications. Theor. Popul. Biol. 118:50–73.

Barton NH, Turelli M. 1987. Adaptive landscapes, genetic distance and the evolution of quantitative characters. Genet. Res. 49:157–173.

Barton NH, Turelli M. 1991. Natural and sexual selection on many loci. Genetics. 127:229–255.

Barton1986. 1986. The maintenance of polygenic variability by a balance between mutation and stabilising selection. Genet. Res. 47.

Bodmer WF, Parsons PA. 1962. Linkage and recombination in evolution. Advan. Genet. 11:1–100.

Brunton SL, Proctor JL, Kutz JN. 2016. Discovering governing equations from data by sparse identification of nonlinear dynamical systems. Proc. Nat. Acad. Sci. USA. 113:3932–3937.

ürger R. 2000. The Mathematical Theory of Selection, Recombination, and Mutation. Wiley. Chichester, UK.

ürger R, Gimelfarb A. 1999. Genetic variation maintained in multilocus models of additive quantitative traits under stabilizing selection. Genetics. 152:807–820.

Carter AJR, Hermisson J, Hansen TF. 2005. The role of epistatic interactions in the response to selection and the evolution of evolvability. Theor. Popul. Biol. 68:179–196.

Caswell H. 2019. Sensitivity Analysis: Matrix Methods in Demography and Ecology. Springer Open. Cham, Switzerland.

Charlesworth B. 2026. Is the fundamental theorem of natural selection of any use? Evolution.

Cheverud JM. 1984. Quantitative genetics and developmental constraints on evolution by selection. J. Theor. Biol. 110:155–171.

Course K, Nair PB. 2023. State estimation of a physical system with unknown governing equations. Nature. 622:261–267.

Cranmer K, Brehmer J, Louppe G. 2020. The frontier of simulation-based inference. Proc. Natl. Acad. Sci. USA. 117:30055–30062.

Débarre F, Nuismer SL, Doebeli M. 2014. Multidimensional (co)evolutionary stability. Am. Nat. 184:158–171.

Deutsch A, Dormann S. 2017. Cellular Automaton Modeling of Biological Pattern Formation. Birkhäuser. Boston, MA, USA. second edition.

Dieckmann U, Law R. 1996. The dynamical theory of coevolution: a derivation from stochastic ecological processes. J. Math. Biol. 34:579–612.

Doebeli M, Ispolatov I. 2014. Chaos and unpredictability in evolution. Evolution. 68:1365–1373.

Ewens WJ. 2004. Mathematical Population Genetics: I. Theoretical Introduction. Springer. New York. second edition.

Feldman MW, Cavalli-Sforza LL. 1976. Cultural and biological evolutionary processes, selection for a trait under complex transmission. Theor. Popul. Biol. 9:238–259.

elsenstein J. 2019. Theoretical Evolutionary Genetics. University of Washington. Seattle, WA.

Fisher RA. 1918. XV.—The correlation between relatives on the supposition of Mendelian inheritance. Trans. Roy. Soc. Edinb. 52:399–433.

Frank SA. 1997. The Price equation, Fisher’s fundamental theorem, kin selection, and causal analysis. Evolution. 51:1712–1729.

Gale JS, Kearsey MJ. 1968. Stable equilibria under stabilising selection in the absence of dominance. Heredity. 23:553–561.

Gavrilets S, Hastings A. 1994. A quantitative-genetic model for selection on developmental noise. Evolution. 48:1478–1486.

Gimelfarb A. 1989. Genotypic variation for a quantitative character maintained under stabilizing selection without mutations: epistasis. Genetics. 123:217–227.

Gimelfarb A. 1996. Pleiotropy as a factor maintaining genetic variation in quantitative characters under stabilizing selection. Genet. Res. 68:65–73.

González-Forero M. 2023. How development affects evolution. Evolution. 77:562–579.

González-Forero M. 2024a. Evolutionary-developmental (evodevo) dynamics of hominin brain size. Nat. Hum. Behav. 8:1321–1333.

González-Forero M. 2024b. A mathematical framework for evodevo dynamics. Theor. Popul. Biol. 155:24–50.

Greene VL. 1977. An algorithm for total and indirect causal effects. Political Methodology. 4:369–381.

Gupte PR, Netz C, Weissing FJ. 2023. The joint evolution of animal movement and competition strategies. Am. Nat. 202:E65–E82.

Hansen TF, Houle D. 2008. Measuring and comparing evolvability and constraint in multivariate characters. J. Evol. Biol. 21:1201–1219.

Hansen TF, Wagner GP. 2001. Modeling genetic architecture: a multilinear theory of gene interaction. Theor. Popul. Biol. 59:61–86.

Heywood JS. 2005. An exact form of the breeder’s equation for the evolution of a quantitative trait under natural selection. Evolution. 59:2287–2298.

Hill WG. 2017. “Conversion” of epistatic into additive genetic variance in finite populations and possible impact on longterm selection response. J. Anim. Breed. Genet. 134:196–201.

Horn RA, Johnson CR. 2013. Matrix Analysis. Cambridge Univ. Press. New York, NY, USA. second edition.

Houle D. 1991. Genetic covariance of fitness correlates: what genetic correlations are made of and why it matters. Evolution. 45:630–648.

Iwasa Y, Pomiankowski A. 1991. The evolution of costly mate preferences II. The “handicap” principle. Evolution. 45:1431–1442.

Jablonka E, Lamb MJ. 2014. Evolution in Four Dimensions. The MIT Press. London, England. revised edition.

Kimura M. 1956. A model of a genetic system which leads to closer linkage under natural selection. Evolution. 10:278–287.

Kingsolver JG, Diamond SE. 2011. Phenotypic selection in natural populations: What limits directional selection? Am. Nat. 177:346–357.

Kingsolver JG, Hoekstra HE, Hoekstra JM, Berrigan D, Vignieri SN et al. 2001. The strength of phenotypic selection in natural populations. Am. Nat. 157:245–261.

Kirkpatrick M. 2009. Patterns of quantitative genetic variation in multiple dimensions. Genetica. 136:271–284.

Kirkpatrick M, Lofsvold D. 1992. Measuring selection and constraint in the evolution of growth. Evolution. 46:954–971.

Lala KN, Uller T, Feiner N, Feldman MW, Gilbert SF. 2024. Evolution Evolving. Princeton Univ. Press.

Lande R. 1979. Quantitative genetic analysis of multivariate evolution applied to brain: body size allometry. Evolution. 34:402–416.

Lande R, Arnold SJ. 1983. The measurement of selection on correlated characters. Evolution. 37:1210–1226.

Lewontin RC. 1970. The units of selection. Annu. Rev. Ecol. Syst. 1:1–18.

Lewontin RC, Kojima K. 1960. The evolutionary dynamics of complex polymorphisms. Evolution. 14:458–472.

Lock JE, Smiseth PT, Moore AJ. 2004. Selection, inheritance, and the evolution of parent-offspring interactions. Am. Nat. 164:13–24.

Lynch M, Walsh B. 1998. Genetics and Analysis of Quantitative Traits. Sinauer. Sunderland, MA, USA.

Martí-Gómez C, Zhou J, Chen WC, Stoltzfus A, Kinney JB et al. 2026. Inference and visualization of complex genotype–phenotype maps. Mol. Biol. Evol. 43:msag023.

Merilä J, Sheldon B, Kruuk L. 2001. Explaining stasis: microevolutionary studies in natural populations. Genetica. 112:199–222.

Moran PAP. 1964. On the nonexistence of adaptive topographies. Ann. Hum. Genet. Lond. 27:383–393.

Morrissey MB. 2014. Selection and evolution of causally covarying traits. Evolution. 68:1748–1761.

Morrissey MB. 2015. Evolutionary quantitative genetics of nonlinear developmental systems. Evolution. 69:2050–2066.

Mullon C, Lehmann L. 2019. An evolutionary quantitative genetics model for phenotypic (co)variances under limited dispersal, with an application to socially synergistic traits. Evolution. 73:1695–1728.

Nagylaki T. 1992. Theoretical Population Genetics. Springer-Verlag. Berlin, Germany.

Nelson RM, Pettersson ME, Carlborg Ö. 2013. A century after Fisher: time for a new paradigm in quantitative genetics. Trends Genet. 29:669–676.

Nuño de la Rosa L, Müller GB. 2021. A reference guide to evodevo, In: Nuño de la Rosa L, Müller GB, editors, Evolutionary Developmental Biology: A Reference Guide, Springer. Cham, Switzerland. pp. 3–12.

Okasha S. 2006. Evolution and the levels of selection. Clarendon Press. Oxford, UK.

Parvinen K. 2005. Evolutionary suicide. Acta Biotheor. 53:241–264.

Parvinen K, Heino M, Dieckmann U. 2013. Function-valued adaptive dynamics and optimal control theory. J. Math. Biol. 67:509–533.

Pavličev M. 2021. Developmental evolutionary biology (devoevo), In: Nuño de la Rosa L, Müller GB, editors, Evolutionary Developmental Biology: A Reference Guide, Springer. Cham, Switzerland. pp. 1033–1046.

Phillips PC, Arnold SJ. 1989. Visualizing multivariate selection. Evolution. 43:1209–1222.

Potticary AL, Cunningham CB, Moore AJ. 2024. Offspring overcome poor parenting by being better parents. J. Evol. Biol. 37.

Price GR. 1970. Selection and covariance. Nature. 227:520–521.

Prokop B, Gelens L. 2026. Data-driven discovery of dynamical models in biology. Nat. Rev. Phys.

Rice SH. 2002. A general population genetic theory for the evolution of developmental interactions. Proc. Natl. Acad. Sci. USA. 99:15518–15523.

Rice SH. 2004a. Developmental associations between traits: covariance and beyond. Genetics. 166:513–526.

Rice SH. 2004b. Evolutionary Theory. Sinauer. Sunderland, MA, USA.

Robertson A. 1956. The effect of selection against extreme deviants based on deviation or homozygosis. J. Genet. 54.

Roff DA, Fairbairn DJ. 2007. The evolution of trade-offs: where are we? J. Evol. Biol. 20:433–447.

Runions A, Tsiantis M, Prusinkiewicz P. 2017. A common developmental program can produce diverse leaf shapes. New Phytol. 216:401–418.

Salazar-Ciudad I, Jernvall J. 2010. A computational model of teeth and the developmental origins of morphological variation. Nature. 464:583–586.

Schluter D. 1996. Adaptive radiation along genetic lines of least resistance. Evolution. 50:1766–1774.

Service PM, Rose MR. 1985. Genetic covariation among life-history components: the effect of novel environments. Evolution. 39:943–945.

Sheth R et al. 2012. Hox genes regulate digit patterning by controlling the wavelength of a Turing-type mechanism. Science. 338:1476–1480.

Slatkin M. 1987. Quantitative genetics of heterochrony. Evolution. 41:799–811.

Tsuboi M et al. 2024. The paradox of predictability provides a bridge between micro- and macroevolution. J. Evol. Biol. 37:1413–1432.

Turelli M. 1988. Phenotypic evolution, constant covariances, and the maintenance of additive variance. Evolution. 42:1342–1347.

Wagner GP. 1984. On the eigenvalue distribution of genetic and phenotypic dispersion matrices: Evidence for a nonrandom organization of quantitative character variation. J. Math. Biol. 21:77–95.

Walsh B, Lynch M. 2018. Evolution and Selection of Quantitative Traits. Oxford Univ. Press. Oxford, UK.

Wright S. 1937. The distribution of gene frequencies in populations. Proc. Natl. Acad. Sci. USA. 23:307–320.

