## Supplementary Information for "A mathematical synthesis of genetics, development, and evolution"

#### Contents

**S1 Derivation of the multivariate Price equation in two forms** **S2**

**S2 Proof that transmission bias contains total selection response** **S2**

**S3 Derivation of equation for unconstrained change in P** **S3**

**S4 Selection regression coefficient under quantitative genetics assumptions** **S4**

**S5 Derivation of selection Hessian under quantitative genetics assumptions** **S6**

**S6 Proof for a special case that a singular constraining matrix entails that evolutionary outcomes depend on initial conditions** **S7**

**S7 Derivation of haplotype frequency in terms of allele frequency and linkage disequilibria** **S8**

**S8 Derivation that there is only one linkage disequilibrium coefficient with two biallelic loci** **S10**

**S9 Additional examples** **S13**

S9.1 Example 2 continued: under selfing, transmission matrix can be slightly negative . . . . . . . . . . S13

S9.2 Example 3 continued: exact method by coupling genetic and phenotypic evolution in generator equations . . . . . . . . S13

S9.3 Example 4 continued: following haplotype frequencies rather than allele frequencies and linkage disequilibrium . . . . . . . . S16

S9.4 Example 5: evo-devo dynamics of one phenotype influenced by one biallelic locus under explicit development, without weakening of selection with age . . . . . . . . S18

S9.5 Example 6: evo-devo dynamics of one phenotype influenced by one biallelic locus under explicit development, with weakening of selection with age . . . . . . . . S22

**References** **S25**

### S1 Derivation of the multivariate Price equation in two forms

This appendix follows the univariate approach of Frank (1997) to obtain a multivariate Price equation. The derivation is essentially the same, but the multivariate treatment has additional implications. The vector $\mathbf{z}$ may have a continuous or discrete probability distribution, or a combination of continuous and discrete. In this appendix, we use integrals for the former case and these can be replaced by sums or combinations of integrals and sums for the latter cases. Since $\mathbf{z}'$ is the expectation of offspring values $\mathbf{z}^+$ conditional on parent values $\mathbf{z}$, then $\mathbf{z}'$ is a function of $\mathbf{z}$, so we sometimes write $\mathbf{z}'(\mathbf{z})$. By the law of total expectation, according to which the expectation of a conditional expectation equals the mean of the conditioned variable, we have that the mean of $\mathbf{z}^+$ in the offspring set is $\bar{\mathbf{z}}^+ = \int \mathbf{z}'(\mathbf{z})p_{\mathbf{z}'}(\mathbf{z})\mathrm{d}\mathbf{z}$, where the overbar denotes average. Hence, the change in mean values from the parent set to the offspring set is

$$\Delta\bar{\mathbf{z}} = \int p_{\mathbf{z}'}(\mathbf{z})\mathbf{z}'(\mathbf{z})\mathrm{d}\mathbf{z} - \int p_{\mathbf{z}}(\mathbf{z})\mathbf{z}\mathrm{d}\mathbf{z}, . \tag{S1}$$

Relative fitness is $w(\mathbf{z}) \equiv p_{\mathbf{z}'}(\mathbf{z})/p_{\mathbf{z}}(\mathbf{z}) \equiv W(\mathbf{z})/\bar{W}$, which implicitly defines absolute fitness $W(\mathbf{z})$. Then, the change in mean phenotype is

$$\begin{aligned}\Delta\bar{\mathbf{z}} &= \int w(\mathbf{z})p_{\mathbf{z}}(\mathbf{z})(\Delta\mathbf{z}+\mathbf{z})\mathrm{d}\mathbf{z} - \int p_{\mathbf{z}}(\mathbf{z})\mathbf{z}\mathrm{d}\mathbf{z}\\ &= \int p_{\mathbf{z}}(\mathbf{z})\mathbf{z}(w(\mathbf{z})-1)\mathrm{d}\mathbf{z} + \int w(\mathbf{z})p_{\mathbf{z}}(\mathbf{z})\Delta\mathbf{z}\mathrm{d}\mathbf{z}\\ &= \mathrm{cov}[\mathbf{z}, w] + \mathrm{E}[w\Delta\mathbf{z}].\end{aligned} \tag{S2}$$

In this derivation, we used the standard Price equation convention that E and cov refer to expectation and covariance over the parent distribution $p_{\mathbf{z}}(\mathbf{z})$. We also used two facts. First, the average relative fitness over the parent distribution is one: $\bar{w} = \mathrm{E}[w] = \int p_{\mathbf{z}}(\mathbf{z})p'_{\mathbf{z}}(\mathbf{z})/p_{\mathbf{z}}(\mathbf{z})\mathrm{d}\mathbf{z} = 1$. Second, for any two random vectors $\mathbf{x}$ and $\mathbf{y}$ with joint probability distribution $p(\mathbf{x},\mathbf{y})$, we have

$$\mathrm{cov}[\mathbf{x},\mathbf{y}] \equiv \int p(\mathbf{x},\mathbf{y})(\mathbf{x}-\bar{\mathbf{x}})(\mathbf{y}-\bar{\mathbf{y}})^\intercal = \int p(\mathbf{x},\mathbf{y})\mathbf{x}(\mathbf{y}-\bar{\mathbf{y}})^\intercal - \bar{\mathbf{x}}\int p(\mathbf{x},\mathbf{y})(\mathbf{y}-\bar{\mathbf{y}})^\intercal = \int p(\mathbf{x},\mathbf{y})\mathbf{x}(\mathbf{y}-\bar{\mathbf{y}})^\intercal, \tag{S3}$$

which gives a shorthand definition of covariance that we use repeatedly throughout without reference to this equation. Eq. (S2) is the multivariate Price equation in standard form describing evolutionary change as the sum of the effects of selection and transmission.

Next, we rearrange the Price equation to separate more neatly the effects of selection and transmission bias. To do this, consider the transmission bias term in the Price equation (S2). Completing terms to obtain covariances yields

$$\begin{aligned}\mathrm{E}[w\Delta\mathbf{z}] &= \mathrm{E}[(w-1)\Delta\mathbf{z}] + \mathrm{E}[\Delta\mathbf{z}]\\ &= \mathrm{cov}[\Delta\mathbf{z}, w] + \mathrm{E}[\Delta\mathbf{z}].\end{aligned}$$

Hence, the Price equation becomes

$$\begin{aligned}\Delta\bar{\mathbf{z}} &= \mathrm{cov}[\mathbf{z}, w] + \mathrm{cov}[\Delta\mathbf{z}, w] + \mathrm{E}[\Delta\mathbf{z}]\\ &= \mathrm{cov}[\mathbf{z}+\Delta\mathbf{z}, w] + \mathrm{E}[\Delta\mathbf{z}]\\ &= \mathrm{cov}[\mathbf{z}', w] + \mathrm{E}[\Delta\mathbf{z}].\end{aligned} \tag{S4}$$

This form neatly separates selection from transmission bias as the second term does not depend on fitness.

### S2 Proof that transmission bias contains total selection response

Transmission bias in the multivariate Price equation in standard form is

$$\mathrm{E}[w\Delta\mathbf{z}] = \mathrm{E}[w\{\boldsymbol{\zeta} + \mathbf{H}(\mathbf{z}-\bar{\mathbf{z}}) + \boldsymbol{\eta} - \mathbf{z}\}]. \tag{S5}$$

Adding and subtracting $\bar{\mathbf{z}}$ yields

$$\begin{aligned}\mathrm{E}[w\Delta\mathbf{z}] &= \mathrm{E}[w\{\boldsymbol{\zeta} + \mathbf{H}(\mathbf{z}-\bar{\mathbf{z}}) + \boldsymbol{\eta} - \mathbf{z} + \bar{\mathbf{z}} - \bar{\mathbf{z}}\}]\\ &= \mathrm{E}[w\{\mathbf{H}(\mathbf{z}-\bar{\mathbf{z}}) + \boldsymbol{\eta} - \mathbf{z} + \bar{\mathbf{z}}\}] + \boldsymbol{\zeta} - \bar{\mathbf{z}}\\ &= \mathrm{E}[w\{(\mathbf{H}-\mathbf{I})(\mathbf{z}-\bar{\mathbf{z}}) + \boldsymbol{\eta}\}] + \mathrm{E}[\Delta\mathbf{z}].\end{aligned} \tag{S6}$$

Recalling that $\mathrm{E}[\boldsymbol{\eta}] = \mathbf{0}$ by least squares, this is

$$\begin{aligned}\mathrm{E}[w\Delta\mathbf{z}] &= (\mathbf{H}-\mathbf{I})\mathrm{cov}[\mathbf{z}, w] + \mathrm{cov}[\boldsymbol{\eta}, w] + \mathrm{E}[\Delta\mathbf{z}] \\ &= (\mathbf{H}-\mathbf{I})\mathbf{s} + \mathbf{u} + \boldsymbol{\zeta} - \bar{\mathbf{z}}.\end{aligned} \tag{S7}$$

Hence, the selection term in the Price equation in its standard form is exactly cancelled by a term in the transmission bias term:

$$\begin{aligned}\Delta\bar{\mathbf{z}} &= \mathrm{cov}[\mathbf{z}, w] + \mathrm{E}[w\Delta\mathbf{z}] \\ &= \mathrm{cov}[\mathbf{z}, w] + (\mathbf{H}-\mathbf{I})\mathrm{cov}[\mathbf{z}, w] + \mathrm{cov}[\boldsymbol{\eta}, w] + +\mathrm{E}[\Delta\mathbf{z}] \\ &= \mathbf{H}\mathrm{cov}[\mathbf{z}, w] + \mathrm{cov}[\boldsymbol{\eta}, w] + \mathrm{E}[\Delta\mathbf{z}].\end{aligned} \tag{S8}$$

### S3 Derivation of equation for unconstrained change in P

We now derive an equation for the unconstrained change in the phenotypic covariance matrix **P** allowing for transmission bias. To do this, we follow Lande and Arnold (1983) with ideas from the Price equation (Frank, 1997) although without using the Price equation itself. Lande and Arnold (1983) assume no transmission bias in the phenotype (i.e., the equation before their equation 13a should have $z'$ instead of $z$ to consider transmission bias), but we allow for transmission bias (i.e., for $\Delta\mathbf{z} \neq \mathbf{0}$).

In our derivation of the Price equation, we had that from the law of total expectation, the expected vector of traits in the offspring set is $\bar{\mathbf{z}}^{+} = \int p_{\mathbf{z}'}(\mathbf{z})\mathbf{z}'(\mathbf{z})\mathrm{d}\mathbf{z}$ (section S1). The covariance matrix of $\mathbf{z}$ in the offspring set is thus

$$\mathbf{P}^{+} = \int p_{\mathbf{z}'}(\mathbf{z})[\mathbf{z}'(\mathbf{z}) - \bar{\mathbf{z}}^{+}][\mathbf{z}'(\mathbf{z}) - \bar{\mathbf{z}}^{+}]^{\top}\mathrm{d}\mathbf{z}. \tag{S9}$$

Expanding the product, this becomes

$$\mathbf{P}^{+} = \int p_{\mathbf{z}'}(\mathbf{z})[\mathbf{z}'\mathbf{z}'^{\top} - \mathbf{z}'\bar{\mathbf{z}}^{+\top} - \bar{\mathbf{z}}^{+}\mathbf{z}'^{\top} + \bar{\mathbf{z}}^{+}\bar{\mathbf{z}}^{+\top}]\mathrm{d}\mathbf{z}.$$

Distributing the integral and simplifying yields

$$\begin{aligned}\mathbf{P}^{+} &= \int p_{\mathbf{z}'}(\mathbf{z})\mathbf{z}'\mathbf{z}'^{\top}\mathrm{d}\mathbf{z} - \int p_{\mathbf{z}'}(\mathbf{z})\mathbf{z}'\bar{\mathbf{z}}^{+\top}\mathrm{d}\mathbf{z} - \int p_{\mathbf{z}'}(\mathbf{z})\bar{\mathbf{z}}^{+}\mathbf{z}'^{\top}\mathrm{d}\mathbf{z} + \int p_{\mathbf{z}'}\bar{\mathbf{z}}^{+}\bar{\mathbf{z}}^{+\top}\mathrm{d}\mathbf{z} \\ &= \int p_{\mathbf{z}'}(\mathbf{z})\mathbf{z}'\mathbf{z}'^{\top}\mathrm{d}\mathbf{z} - \bar{\mathbf{z}}^{+}\bar{\mathbf{z}}^{+\top} - \bar{\mathbf{z}}^{+}\bar{\mathbf{z}}^{+\top} + \bar{\mathbf{z}}^{+}\bar{\mathbf{z}}^{+\top} \\ &= \int p_{\mathbf{z}'}(\mathbf{z})\mathbf{z}'\mathbf{z}'^{\top}\mathrm{d}\mathbf{z} - \bar{\mathbf{z}}^{+}\bar{\mathbf{z}}^{+\top}.\end{aligned} \tag{S10}$$

Now using Eq. 3, we have that

$$\begin{aligned}\mathbf{z}'\mathbf{z}'^{\top} &= [\boldsymbol{\zeta} + \mathbf{H}(\mathbf{z}-\bar{\mathbf{z}}) + \boldsymbol{\eta}][\boldsymbol{\zeta} + \mathbf{H}(\mathbf{z}-\bar{\mathbf{z}}) + \boldsymbol{\eta}]^{\top} \\ &= \boldsymbol{\zeta}\boldsymbol{\zeta}^{\top} + \boldsymbol{\zeta}(\mathbf{z}-\bar{\mathbf{z}})^{\top}\mathbf{H}^{\top} + \boldsymbol{\zeta}\boldsymbol{\eta}^{\top} \\ &\quad + \mathbf{H}(\mathbf{z}-\bar{\mathbf{z}})\boldsymbol{\zeta}^{\top} + \mathbf{H}(\mathbf{z}-\bar{\mathbf{z}})(\mathbf{z}-\bar{\mathbf{z}})^{\top}\mathbf{H}^{\top} \\ &\quad + \mathbf{H}(\mathbf{z}-\bar{\mathbf{z}})\boldsymbol{\eta}^{\top} \\ &\quad + \boldsymbol{\eta}\boldsymbol{\zeta}^{\top} + \boldsymbol{\eta}(\mathbf{z}-\bar{\mathbf{z}})^{\top}\mathbf{H}^{\top} + \boldsymbol{\eta}\boldsymbol{\eta}^{\top}.\end{aligned}$$

Hence, since $p_{\mathbf{z}'}(\mathbf{z}) = p_{\mathbf{z}}(\mathbf{z})w(\mathbf{z})$, we have

$$\begin{aligned}\int p_{\mathbf{z}'}(\mathbf{z})\mathbf{z}'\mathbf{z}'^{\top}\mathrm{d}\mathbf{z} &= \int p_{\mathbf{z}}(\mathbf{z})w(\mathbf{z})\mathbf{z}'\mathbf{z}'^{\top}\mathrm{d}\mathbf{z} \\ &= \boldsymbol{\zeta}\boldsymbol{\zeta}^{\top} + \boldsymbol{\zeta}\mathrm{cov}[\mathbf{z}^{\top}, w]\mathbf{H}^{\top} + \boldsymbol{\zeta}\mathrm{cov}[\boldsymbol{\eta}^{\top}, w] \\ &\quad + \mathbf{H}\mathrm{cov}[\mathbf{z}, w]\boldsymbol{\zeta}^{\top} + \mathbf{H}(\mathrm{cov}[\mathbf{Z}, w] + \mathbf{P})\mathbf{H}^{\top} \\ &\quad + \mathbf{H}\mathrm{cov}[w(\mathbf{z})(\mathbf{z}-\bar{\mathbf{z}}), \boldsymbol{\eta}] \\ &\quad + \mathrm{cov}[\boldsymbol{\eta}, w]\boldsymbol{\zeta}^{\top} + \mathrm{cov}[\boldsymbol{\eta}, w(\mathbf{z})(\mathbf{z}-\bar{\mathbf{z}})]\mathbf{H}^{\top} \\ &\quad + \mathrm{cov}[\boldsymbol{\eta}, \boldsymbol{\eta}].\end{aligned}$$

From Eq. 18, we have that $\mathbf{P}' = \mathbf{HPH}^\intercal + \mathrm{cov}[\boldsymbol{\eta}, \boldsymbol{\eta}]$. Then, using $\mathbf{s} = \mathrm{cov}[\mathbf{z}, w]$, $\mathbf{S} = \mathrm{cov}[\mathbf{Z}, w]$, and $\mathbf{u} = \mathrm{cov}[\boldsymbol{\eta}, w]$, we have

$$\begin{aligned}
&\int p_{\mathbf{z}}(\mathbf{z})\, w(\mathbf{z}) \mathbf{z}' \mathbf{z}'^\intercal \mathrm{d}\mathbf{z} \\
&= \boldsymbol{\zeta}(\boldsymbol{\zeta}^\intercal + \mathbf{s}^\intercal \mathbf{H}^\intercal + \mathbf{u}^\intercal) \\
&\quad + \mathbf{Hs}\boldsymbol{\zeta}^\intercal + \mathbf{HSH}^\intercal \\
&\quad + \mathbf{H}\mathrm{cov}[w(\mathbf{z})(\mathbf{z} - \bar{\mathbf{z}}), \boldsymbol{\eta}] \\
&\quad + \mathbf{u}\boldsymbol{\zeta}^\intercal + \mathrm{cov}[\boldsymbol{\eta}, w(\mathbf{z})(\mathbf{z} - \bar{\mathbf{z}})]\mathbf{H}^\intercal \\
&\quad + \mathbf{P}'.
\end{aligned}$$

From Eqs. 2, 4 and 7, note that $\boldsymbol{\zeta} + \mathbf{Hs} + \mathbf{u} = \Delta\bar{\mathbf{z}} + \bar{\mathbf{z}}$. Then,

$$\begin{aligned}
&\int p_{\mathbf{z}}(\mathbf{z})\, w(\mathbf{z}) \mathbf{z}' \mathbf{z}'^\intercal \mathrm{d}\mathbf{z} \\
&= \boldsymbol{\zeta}(\Delta\bar{\mathbf{z}} + \bar{\mathbf{z}})^\intercal \\
&\quad + (\Delta\bar{\mathbf{z}} + \bar{\mathbf{z}} - \boldsymbol{\zeta})\boldsymbol{\zeta}^\intercal + \mathbf{HSH}^\intercal \\
&\quad + \mathbf{H}\mathrm{cov}[w(\mathbf{z})(\mathbf{z} - \bar{\mathbf{z}}), \boldsymbol{\eta}] \\
&\quad + \mathrm{cov}[\boldsymbol{\eta}, w(\mathbf{z})(\mathbf{z} - \bar{\mathbf{z}})]\mathbf{H}^\intercal \\
&\quad + \mathbf{P}'.
\end{aligned} \tag{S11}$$

Substituting (S11) into (S10) noting that $\bar{\mathbf{z}}^+ = \Delta\bar{\mathbf{z}} + \bar{\mathbf{z}}$ yields

$$\begin{aligned}
\mathbf{P}^+ &= \boldsymbol{\zeta}(\Delta\bar{\mathbf{z}} + \bar{\mathbf{z}})^\intercal \\
&\quad + (\Delta\bar{\mathbf{z}} + \bar{\mathbf{z}} - \boldsymbol{\zeta})\boldsymbol{\zeta}^\intercal + \mathbf{HSH}^\intercal \\
&\quad + \mathbf{H}\mathrm{cov}[w(\mathbf{z})(\mathbf{z} - \bar{\mathbf{z}}), \boldsymbol{\eta}] \\
&\quad + \mathrm{cov}[\boldsymbol{\eta}, w(\mathbf{z})(\mathbf{z} - \bar{\mathbf{z}})]\mathbf{H}^\intercal \\
&\quad + \mathbf{P}' \\
&\quad - (\Delta\bar{\mathbf{z}} + \bar{\mathbf{z}})(\Delta\bar{\mathbf{z}} + \bar{\mathbf{z}})^\intercal.
\end{aligned}$$

Now note that the 1st, 2nd, and 7th factors can be factored as

$$\begin{aligned}
&\boldsymbol{\zeta}(\Delta\bar{\mathbf{z}} + \bar{\mathbf{z}})^\intercal + (\Delta\bar{\mathbf{z}} + \bar{\mathbf{z}} - \boldsymbol{\zeta})\boldsymbol{\zeta}^\intercal - (\Delta\bar{\mathbf{z}} + \bar{\mathbf{z}})(\Delta\bar{\mathbf{z}} + \bar{\mathbf{z}})^\intercal \\
&= -(\Delta\bar{\mathbf{z}} + \bar{\mathbf{z}} - \boldsymbol{\zeta})(\Delta\bar{\mathbf{z}} + \bar{\mathbf{z}})^\intercal + (\Delta\bar{\mathbf{z}} + \bar{\mathbf{z}} - \boldsymbol{\zeta})\boldsymbol{\zeta}^\intercal \\
&= -(\Delta\bar{\mathbf{z}} + \bar{\mathbf{z}} - \boldsymbol{\zeta})(\Delta\bar{\mathbf{z}} + \bar{\mathbf{z}} - \boldsymbol{\zeta})^\intercal \\
&= -(\mathbf{Hs} + \mathbf{u})(\mathbf{Hs} + \mathbf{u})^\intercal.
\end{aligned}$$

Substituting back yields

$$\begin{aligned}
\mathbf{P}^+ &= \mathbf{HSH}^\intercal - (\mathbf{Hs} + \mathbf{u})(\mathbf{Hs} + \mathbf{u})^\intercal \\
&\quad + \mathbf{H}\mathrm{cov}[w(\mathbf{z})(\mathbf{z} - \bar{\mathbf{z}}), \boldsymbol{\eta}] \\
&\quad + \mathrm{cov}[\boldsymbol{\eta}, w(\mathbf{z})(\mathbf{z} - \bar{\mathbf{z}})]\mathbf{H}^\intercal \\
&\quad + \mathbf{P}'.
\end{aligned}$$

Assuming that there is no selection response from non-linear inheritance $\mathbf{u} = \mathbf{0}$ and similarly that $\mathrm{cov}[\boldsymbol{\eta}, w(\mathbf{z})(\mathbf{z} - \bar{\mathbf{z}})] = \mathbf{0}$ (e.g., if $\mathbf{z}'$ and $\mathbf{z}$ are jointly multivariate normal, so $\boldsymbol{\eta} = \mathbf{0}$), this reduces to

$$\mathbf{P}^+ = \mathbf{HSH}^\intercal - \mathbf{Hss}^\intercal \mathbf{H}^\intercal + \mathbf{P}'.$$

Factoring and noting that $\Delta\mathbf{P} = \mathbf{P}^+ - \mathbf{P}$, we finally obtain

$$\Delta\mathbf{P} = \mathbf{H}\left(\mathbf{S} - \mathbf{ss}^\intercal\right)\mathbf{H}^\intercal + \mathbf{P}' - \mathbf{P}.$$

### S4 Selection regression coefficient under quantitative genetics assumptions

We now show how the selection regression coefficient $\boldsymbol{\beta}$ relates to the selection gradient following Lande and Arnold (1983). To do this, we assume that the phenotype $\mathbf{z}$ has a multivariate normal distribution $p_{\mathbf{z}}(\mathbf{z})$, and we seek to calculate the average fitness gradient

$$\begin{aligned}\mathrm{E}\left[\frac{\partial w}{\partial \mathbf{z}}\right] &= \int_{-\infty}^{\infty} p_{\mathbf{z}}(\mathbf{z})\frac{\partial w}{\partial \mathbf{z}}\mathrm{d}\mathbf{z} \\ &= -\int_{-\infty}^{\infty} \frac{\partial p_{\mathbf{z}}}{\partial \mathbf{z}} w(\mathbf{z})\mathrm{d}\mathbf{z},\end{aligned}$$

where the last equality follows from integration by parts and the fact that the normal distribution vanishes at $\pm\infty$. Now, because $p_{\mathbf{z}}$ is the multivariate normal distribution, we have

$$\begin{aligned}\frac{\partial p_{\mathbf{z}}}{\partial \mathbf{z}} &= p_{\mathbf{z}}\left[-\frac{1}{2}\left(\frac{\partial(\mathbf{z}-\bar{\mathbf{z}})^{\intercal}}{\partial \mathbf{z}}\mathbf{P}^{-1}(\mathbf{z}-\bar{\mathbf{z}}) + (\mathbf{z}-\bar{\mathbf{z}})^{\intercal}\mathbf{P}^{-1}\frac{\partial(\mathbf{z}-\bar{\mathbf{z}})}{\partial \mathbf{z}}\right)\right] \\ &= p_{\mathbf{z}}\left[-\frac{1}{2}\left(\mathbf{P}^{-1}(\mathbf{z}-\bar{\mathbf{z}}) + (\mathbf{z}-\bar{\mathbf{z}})^{\intercal}\mathbf{P}^{-1}\right)\right] \\ &= -p_{\mathbf{z}}\mathbf{P}^{-1}(\mathbf{z}-\bar{\mathbf{z}}).\end{aligned}$$

Hence, the average fitness gradient is

$$\begin{aligned}\mathrm{E}\left[\frac{\partial w}{\partial \mathbf{z}}\right] &= \mathbf{P}^{-1}\int_{-\infty}^{\infty} p_{\mathbf{z}}(\mathbf{z})(\mathbf{z}-\bar{\mathbf{z}})w(\mathbf{z})\mathrm{d}\mathbf{z} \\ &= \mathbf{P}^{-1}\mathrm{cov}[\mathbf{z}, w]. \qquad \text{(S12)}\end{aligned}$$

Then, the selection differential is

$$\mathrm{cov}[\mathbf{z}, w] = \mathbf{P}\mathrm{E}\left[\frac{\partial w}{\partial \mathbf{z}}\right]. \qquad \text{(S13)}$$

As $\mathbf{P}$ is invertible by the normality assumption, using equation (9) yields

$$\boldsymbol{\beta} = \mathrm{E}\left[\frac{\partial w}{\partial \mathbf{z}}\right]. \qquad \text{(S14)}$$

Now we relate the average fitness gradient to the gradient of mean fitness. Since $\bar{W} = \int W(\mathbf{z})p_{\mathbf{z}}(\mathbf{z})\mathrm{d}\mathbf{z}$, we have that

$$\frac{\partial \bar{W}}{\partial \bar{\mathbf{z}}} = \int\left(\frac{\partial W(\mathbf{z})}{\partial \bar{\mathbf{z}}}p_{\mathbf{z}}(\mathbf{z}) + W(\mathbf{z})\frac{\partial p_{\mathbf{z}}(\mathbf{z})}{\partial \bar{\mathbf{z}}}\right)\mathrm{d}\mathbf{z}.$$

Assuming that $\mathbf{z}$ is multivariate normal, then

$$\begin{aligned}\frac{\partial p_{\mathbf{z}}}{\partial \bar{\mathbf{z}}} &= p_{\mathbf{z}}\left[-\frac{1}{2}\left(-\mathbf{P}^{-1}(\mathbf{z}-\bar{\mathbf{z}}) - (\mathbf{z}-\bar{\mathbf{z}})^{\intercal}\mathbf{P}^{-1}\right)\right] \\ &= p_{\mathbf{z}}\mathbf{P}^{-1}(\mathbf{z}-\bar{\mathbf{z}}). \qquad \text{(S15)}\end{aligned}$$

Hence, assuming that $\partial W(\mathbf{z})/\partial\bar{\mathbf{z}} = \mathbf{0}$, we have

$$\begin{aligned}\frac{\partial \bar{W}}{\partial \bar{\mathbf{z}}} &= \int_{-\infty}^{\infty} W(\mathbf{z})p_{\mathbf{z}}(\mathbf{z})\mathbf{P}^{-1}(\mathbf{z}-\bar{\mathbf{z}})\mathrm{d}\mathbf{z} \\ &= \mathbf{P}^{-1}\int_{-\infty}^{\infty} W(\mathbf{z})p_{\mathbf{z}}(\mathbf{z})(\mathbf{z}-\bar{\mathbf{z}})\mathrm{d}\mathbf{z} \qquad \text{(S16a)} \\ &= \mathbf{P}^{-1}\mathrm{cov}[\mathbf{z}, W] \qquad \text{(S16b)} \\ &= \bar{W}\mathrm{E}\left[\frac{\partial w}{\partial \mathbf{z}}\right]. \qquad \text{(S16c)}\end{aligned}$$

where the last equation follows from (S12).

Then, we obtain the form of the selection gradient of Lande (1979) so the average fitness gradient is the proportional gradient of mean absolute fitness:

$$\mathrm{E}\left[\frac{\partial w}{\partial \mathbf{z}}\right] = \frac{1}{\bar{W}}\frac{\partial \bar{W}}{\partial \bar{\mathbf{z}}} = \frac{\partial \ln \bar{W}}{\partial \bar{\mathbf{z}}}. \qquad \text{(S17)}$$

#### S5 Derivation of selection Hessian under quantitative genetics assumptions

We now show how the selection differential for $\mathbf{Z}$, $\mathbf{S} = \mathrm{cov}[\mathbf{Z}, w]$, relates to the average fitness Hessian following Lande and Arnold (1983). To do this, we assume that the phenotype $\mathbf{z}$ has the multivariate normal distribution $p_{\mathbf{z}}(\mathbf{z})$, and we seek to calculate the average fitness Hessian

$$\begin{aligned}
\mathrm{E}\left[\frac{\partial^2 w}{\partial \mathbf{z} \partial \mathbf{z}^\intercal}\right] &= \int_{-\infty}^{\infty} p_{\mathbf{z}}(\mathbf{z}) \frac{\partial^2 w}{\partial \mathbf{z} \partial \mathbf{z}^\intercal} \mathrm{d}\mathbf{z} \\
&= \int_{-\infty}^{\infty} p_{\mathbf{z}}(\mathbf{z}) \frac{\partial}{\partial \mathbf{z}} \frac{\partial w}{\partial \mathbf{z}^\intercal} \mathrm{d}\mathbf{z} \\
&= -\int_{-\infty}^{\infty} \frac{\partial p_{\mathbf{z}}(\mathbf{z})}{\partial \mathbf{z}} \frac{\partial w}{\partial \mathbf{z}^\intercal} \mathrm{d}\mathbf{z} \\
&= \int_{-\infty}^{\infty} \frac{\partial^2 p_{\mathbf{z}}}{\partial \mathbf{z} \partial \mathbf{z}^\intercal} w(\mathbf{z}) \mathrm{d}\mathbf{z},
\end{aligned}$$

where the third and fourth equalities follow from doing integration by parts twice and the fact that the normal distribution vanishes at $\pm\infty$. Now, since $p_{\mathbf{z}}$ is the multivariate normal distribution, we have

$$\begin{aligned}
\frac{\partial}{\partial \mathbf{z}} \frac{\partial p_{\mathbf{z}}}{\partial \mathbf{z}^\intercal} &= \frac{\partial}{\partial \mathbf{z}} \left(-p_{\mathbf{z}}(\mathbf{z})(\mathbf{z}-\bar{\mathbf{z}})^\intercal \mathbf{P}^{-1}\right) \\
&= -\left(\frac{\partial p_{\mathbf{z}}(\mathbf{z})}{\partial \mathbf{z}} (\mathbf{z}-\bar{\mathbf{z}})^\intercal + p_{\mathbf{z}}(\mathbf{z}) \frac{\partial (\mathbf{z}-\bar{\mathbf{z}})^\intercal}{\partial \mathbf{z}}\right) \mathbf{P}^{-1} \\
&= -\left(-p_{\mathbf{z}}(\mathbf{z}) \mathbf{P}^{-1} (\mathbf{z}-\bar{\mathbf{z}})(\mathbf{z}-\bar{\mathbf{z}})^\intercal + p_{\mathbf{z}}(\mathbf{z}) \mathbf{I}\right) \mathbf{P}^{-1} \\
&= p_{\mathbf{z}}(\mathbf{z}) \left(\mathbf{P}^{-1} (\mathbf{z}-\bar{\mathbf{z}})(\mathbf{z}-\bar{\mathbf{z}})^\intercal - \mathbf{I}\right) \mathbf{P}^{-1}
\end{aligned}$$

Hence, the average fitness Hessian is

$$\begin{aligned}
\mathrm{E}\left[\frac{\partial^2 w}{\partial \mathbf{z} \partial \mathbf{z}^\intercal}\right] &= \int_{-\infty}^{\infty} p_{\mathbf{z}}(\mathbf{z}) \left(\mathbf{P}^{-1} (\mathbf{z}-\bar{\mathbf{z}})(\mathbf{z}-\bar{\mathbf{z}})^\intercal - \mathbf{I}\right) w(\mathbf{z}) \mathrm{d}\mathbf{z} \mathbf{P}^{-1} \\
&= \mathbf{P}^{-1} \int_{-\infty}^{\infty} p_{\mathbf{z}}(\mathbf{z}) (\mathbf{z}-\bar{\mathbf{z}})(\mathbf{z}-\bar{\mathbf{z}})^\intercal w(\mathbf{z}) \mathrm{d}\mathbf{z} \mathbf{P}^{-1} \\
&\quad - \int_{-\infty}^{\infty} p_{\mathbf{z}}(\mathbf{z}) w(\mathbf{z}) \mathrm{d}\mathbf{z} \mathbf{P}^{-1} \\
&= \mathbf{P}^{-1} \left[\mathrm{cov}[(\mathbf{z}-\bar{\mathbf{z}})(\mathbf{z}-\bar{\mathbf{z}})^\intercal, w] + \mathbf{P}\right] \mathbf{P}^{-1} \\
&\quad - \mathbf{P}^{-1} \\
&= \mathbf{P}^{-1} \mathrm{cov}[\mathbf{Z}, w] \mathbf{P}^{-1}.
\end{aligned} \tag{S18}$$

Therefore, if the phenotype $\mathbf{z}$ is multivariate normal, the selection differential for $\mathbf{Z}$ is

$$\mathbf{S} \equiv \mathrm{cov}[\mathbf{Z}, w] = \mathbf{P}\mathrm{E}\left[\frac{\partial^2 w}{\partial \mathbf{z} \partial \mathbf{z}^\intercal}\right] \mathbf{P}, \tag{S19}$$

as shown by Lande and Arnold (1983).

Hence, equation (21) for the unconstrained change in the covariance matrix of $\mathbf{z}$ becomes

$$\begin{aligned}
\Delta \mathbf{P} &= \mathbf{H}\left(\mathbf{P}\mathrm{E}\left[\frac{\partial^2 w}{\partial \mathbf{z} \partial \mathbf{z}^\intercal}\right] \mathbf{P} - \mathbf{P}\mathrm{E}\left[\frac{\partial w}{\partial \mathbf{z}}\right] \mathrm{E}\left[\frac{\partial w}{\partial \mathbf{z}^\intercal}\right] \mathbf{P}\right) \mathbf{H}^\intercal \\
&\quad + \mathbf{P}' - \mathbf{P} \\
&= \mathbf{H}\mathbf{P}\left(\mathrm{E}\left[\frac{\partial^2 w}{\partial \mathbf{z} \partial \mathbf{z}^\intercal}\right] - \mathrm{E}\left[\frac{\partial w}{\partial \mathbf{z}}\right] \mathrm{E}\left[\frac{\partial w}{\partial \mathbf{z}^\intercal}\right]\right) \mathbf{P}\mathbf{H}^\intercal \\
&\quad + \mathbf{P}' - \mathbf{P}.
\end{aligned}$$

Recalling that $\mathbf{T} = \mathbf{H}\mathbf{P}$ (Eq. 11) and that $\mathbf{P}$ is symmetric, we obtain

$$\begin{aligned}
\Delta \mathbf{P} &= \mathbf{T}\left(\mathrm{E}\left[\frac{\partial^2 w}{\partial \mathbf{z} \partial \mathbf{z}^\intercal}\right] - \mathrm{E}\left[\frac{\partial w}{\partial \mathbf{z}}\right] \mathrm{E}\left[\frac{\partial w}{\partial \mathbf{z}^\intercal}\right]\right) \mathbf{T}^\intercal \\
&\quad + \mathbf{P}' - \mathbf{P}.
\end{aligned} \tag{S20}$$

Now we relate the average fitness Hessian to the Hessian of mean fitness. Assuming that $\mathbf{z}$ is multivariate normal and that $\partial W(\mathbf{z})/\partial\bar{\mathbf{z}} = \mathbf{0}$, using (S16b) we have that

$$\frac{\partial^2 \bar{W}}{\partial\bar{\mathbf{z}}\partial\bar{\mathbf{z}}^\intercal} = \frac{\partial}{\partial\bar{\mathbf{z}}}\left(\frac{\partial\bar{W}}{\partial\bar{\mathbf{z}}^\intercal}\right) = \frac{\partial}{\partial\bar{\mathbf{z}}}\left(\mathrm{cov}[\mathbf{z}^\intercal, W]\mathbf{P}^{-1}\right).$$

Assuming that $\mathbf{P}$ is independent of the mean phenotype ($\partial\mathbf{P}/\partial\bar{\mathbf{z}} = \mathbf{0}$), then

$$\begin{aligned}\frac{\partial^2 \bar{W}}{\partial\bar{\mathbf{z}}\partial\bar{\mathbf{z}}^\intercal} &= \frac{\partial}{\partial\bar{\mathbf{z}}}\left(\int_{-\infty}^{\infty} W(\mathbf{z})p_{\mathbf{z}}(\mathbf{z})(\mathbf{z}-\bar{\mathbf{z}})^\intercal \mathrm{d}\mathbf{z}\right)\mathbf{P}^{-1} \\ &= \int_{-\infty}^{\infty} W(\mathbf{z})\left(-p_{\mathbf{z}}(\mathbf{z}) + \frac{\partial p}{\partial\bar{\mathbf{z}}}(\mathbf{z}-\bar{\mathbf{z}})^\intercal\right)\mathrm{d}\mathbf{z}\mathbf{P}^{-1}.\end{aligned}$$

Using (S15), this becomes

$$\begin{aligned}\frac{\partial^2 \bar{W}}{\partial\bar{\mathbf{z}}\partial\bar{\mathbf{z}}^\intercal} &= -\bar{W}\mathbf{P}^{-1} + \int_{-\infty}^{\infty} W(\mathbf{z})p_{\mathbf{z}}(\mathbf{z})\mathbf{P}^{-1}(\mathbf{z}-\bar{\mathbf{z}})(\mathbf{z}-\bar{\mathbf{z}})^\intercal \mathrm{d}\mathbf{z}\mathbf{P}^{-1} \\ &= -\bar{W}\mathbf{P}^{-1} \\ &\quad + \mathbf{P}^{-1}\int_{-\infty}^{\infty}(W(\mathbf{z}) - \bar{W})p_{\mathbf{z}}(\mathbf{z})(\mathbf{z}-\bar{\mathbf{z}})(\mathbf{z}-\bar{\mathbf{z}})^\intercal \mathrm{d}\mathbf{z}\mathbf{P}^{-1} \\ &\quad + \mathbf{P}^{-1}\int_{-\infty}^{\infty}\bar{W}p_{\mathbf{z}}(\mathbf{z})(\mathbf{z}-\bar{\mathbf{z}})(\mathbf{z}-\bar{\mathbf{z}})^\intercal \mathrm{d}\mathbf{z}\mathbf{P}^{-1} \\ &= -\bar{W}\mathbf{P}^{-1} \\ &\quad + \mathbf{P}^{-1}\mathrm{cov}[\mathbf{Z}, W]\mathbf{P}^{-1} \\ &\quad + \bar{W}\mathbf{P}^{-1}\mathbf{P}\mathbf{P}^{-1} \\ &= \mathbf{P}^{-1}\mathrm{cov}[\mathbf{Z}, W]\mathbf{P}^{-1}.\end{aligned}$$

Using (S18) here yields

$$\mathrm{E}\left[\frac{\partial^2 w}{\partial\mathbf{z}\partial\mathbf{z}^\intercal}\right] = \frac{1}{\bar{W}}\frac{\partial^2 \bar{W}}{\partial\bar{\mathbf{z}}\partial\bar{\mathbf{z}}^\intercal}. \tag{S21}$$

Substituting (S17) and (S21) into (S20), the unconstrained change in the covariance matrix of $\mathbf{z}$ becomes

$$\begin{aligned}\Delta\mathbf{P} = \mathbf{T}&\left(\frac{1}{\bar{W}}\frac{\partial^2 \bar{W}}{\partial\bar{\mathbf{z}}\partial\bar{\mathbf{z}}^\intercal} - \frac{\partial\ln\bar{W}}{\partial\bar{\mathbf{z}}}\frac{\partial\ln\bar{W}}{\partial\bar{\mathbf{z}}^\intercal}\right)\mathbf{T}^\intercal \\ &+ \mathbf{P}' - \mathbf{P}.\end{aligned} \tag{S22}$$

By further assuming perfect heredity $\mathbf{H} = \mathbf{I}$ and no pure transmission bias $\mathbf{z}' = \mathbf{z}$ so $\mathbf{P}' = \mathbf{P}$, we recover the classic equation describing the change in $\mathbf{P}$ due to selection (eq. 15a of Lande and Arnold 1983).

### S6 Proof for a special case that a singular constraining matrix entails that evolutionary outcomes depend on initial conditions

We prove this assuming a constant constraining matrix. Consider the matrix differential equation

$$\frac{\mathrm{d}\mathbf{z}}{\mathrm{d}t} = \mathbf{C}(\boldsymbol{\theta} - \mathbf{z}), \tag{S23}$$

where $\mathbf{z} \in \mathbb{R}^{(n+1)\times 1}$ is an evolving vector, $\mathbf{C} \in \mathbb{R}^{(n+1)\times(n+1)}$ is a constant, positive semi-definite, singular, and diagonalisable matrix with $n$ distinct eigenvalues that are positive and 1 eigenvalue that is exactly zero, and $\boldsymbol{\theta}$ is a constant vector (the optimum). We will solve this differential equation and prove that as time tends to infinity, the vector $\mathbf{z}$ tends to a value that depends on the initial conditions because $\mathbf{C}$ is singular. We consider only one distinct eigenvalues because we will prove this using diagonalisation, and strict diagonalisation is only possible with non-repeated eigenvalues. A more general proof for repeated eigenvalues would be more involved, presumably requiring Jordan normal forms.

We use the standard strategy of solving first the diagonalised system, which is easier. So, we first diagonalise $\mathbf{C}$ as

$$\mathbf{V}^{-1}\mathbf{C}\mathbf{V} = \mathbf{U}, \tag{S24}$$

where $\mathbf{U}$ is a diagonal matrix listing in its main diagonal the eigenvalues of $\mathbf{C}$ and $\mathbf{V}$ is an invertible matrix whose $i$-th column gives the eigenvector $\mathbf{v}_i$ associated to the $i$-th eigenvalue listed in $\mathbf{U}$. For simplicity, let $\mathbf{U}$ list the positive eigenvalues first and then the zero eigenvalue. Thus,

$$\mathbf{C} = \mathbf{V}\mathbf{U}\mathbf{V}^{-1}. \tag{S25}$$

Now consider the change of variables given by

$$\mathbf{z} = \mathbf{V}\mathbf{y} \tag{S26a}$$
$$\boldsymbol{\theta} = \mathbf{V}\boldsymbol{\theta}_{\mathbf{y}} \tag{S26b}$$

for some vector $\mathbf{y} = \mathbf{V}^{-1}\mathbf{z}$. Taking the time derivative of $\mathbf{y}$, we obtain

$$\begin{aligned}\frac{\mathrm{d}\mathbf{y}}{\mathrm{d}t} &= \mathbf{V}^{-1}\frac{\mathrm{d}\mathbf{z}}{\mathrm{d}t}\\ &= \mathbf{V}^{-1}\mathbf{C}(\boldsymbol{\theta} - \mathbf{z})\\ &= \mathbf{V}^{-1}\mathbf{V}\mathbf{U}\mathbf{V}^{-1}\mathbf{V}(\boldsymbol{\theta}_{\mathbf{y}} - \mathbf{y})\\ &= \mathbf{U}(\boldsymbol{\theta}_{\mathbf{y}} - \mathbf{y}).\end{aligned} \tag{S27}$$

Since $\mathbf{U}$ is diagonal, this is a system of decoupled differential equations. Hence, the solution of this system is

$$\mathbf{y}(t) = \boldsymbol{\theta}_{\mathbf{y}} - \exp(-\mathbf{U}t)(\boldsymbol{\theta}_{\mathbf{y}} - \mathbf{y}(0)), \tag{S28}$$

where $\mathbf{y}(0)$ is $\mathbf{y}$ at time $t = 0$ and $\exp(-\mathbf{U}t)$ is a diagonal matrix whose $i$-th diagonal entry is $\exp(-\lambda_i t)$, where $\lambda_i$ is the $i$-th eigenvalue of $\mathbf{C}$ as listed in $\mathbf{U}$. Since the last $\lambda_i$ is zero, the last entry of the diagonal of $\exp(-\mathbf{U}t)$ is one. Hence, taking the product yields

$$y_i(t) = \begin{cases}\theta_{y_i} - \exp(-\lambda_i t)(\theta_{y_i} - y_i(0)) & \text{if } i \le n\\ y_i(0) & \text{if } i = n+1.\end{cases} \tag{S29}$$

Thus, the $y_i$ corresponding to the zero eigenvalue does not evolve and remains at its initial value. In the limit as $t \to \infty$, we obtain

$$y_i(\infty) = \begin{cases}\theta_{y_i} & \text{if } i \le n\\ y_i(0) & \text{if } i = n+1.\end{cases} \tag{S30}$$

Applying (S30) to (S26a) yields that the $i$-th variable converges to the following value as time advances

$$z_i(\infty) = \sum_{j=1}^{n} V_{ij}\theta_{y_j} + V_{i,n+1}y_{n+1}(0). \tag{S31}$$

Since from (S26b), the optimum is $\theta_i = \sum_{j=1}^{n+1} V_{ij}\theta_{y_j} = \sum_{j=1}^{n} V_{ij}\theta_{y_j} + V_{i,n+1}\theta_{y_{n+1}}$, we obtain

$$z_i(\infty) = \theta_i + V_{i,n+1}[y_{n+1}(0) - \theta_{y_{n+1}}]. \tag{S32}$$

Therefore, as time tends to infinity, $\mathbf{z}$ converges to a value that: depends on the initial conditions and on the constraining matrix $\mathbf{C}$ (because the value depends on $V_{i,n+1}$) and that it is not generally the optimum.

### S7 Derivation of haplotype frequency in terms of allele frequency and linkage disequilibria

We have that haplotype frequency $p_{\mathbf{k}}$ is the expected value of the product of random variables. Here we seek to write such haplotype frequency in terms of allele frequencies and linkage disequilibrium coefficients. We do so by moving the expected value inside the product of random values as follows.

The derivation from here to equation (S33e) was produced by Gemini and is written here in my own words. The prompt used was:

```
Let's say I want to compute the expectation of a product of random variables E[\prod_{i=1}^n
X_i]. I want to write this expectation in terms of \prod_{i=1}^n E[X_i] plus covariance
terms. What would that look like?
```

Let $Y_i \in \mathbb{R}$ for $i \in \{1, \ldots, n\}$ be $n$ random variables. We seek to write the expected value of the product of these random variables

$$\mathrm{E}\left[\prod_{i=1}^{n} Y_i\right] \tag{S33a}$$

by moving the expected value inside the product. Let us write each random variable's deviation from the mean as $\tilde{Y}_i = Y_i - \mu_i$, where $\mu_i = \mathrm{E}[Y_i]$ is the mean of each random variable, so the expectation becomes

$$\mathrm{E}\left[\prod_{i=1}^{n} Y_i\right] = \mathrm{E}\left[\prod_{i=1}^{n} (\mu_i + \tilde{Y}_i)\right]. \tag{S33b}$$

Note that the distributive property of multiplication and addition is

$$\prod_{i=1}^{n} (a_i + b_i) = \sum_{Q \subseteq [n]} \left(\prod_{j \notin Q} a_j\right)\left(\prod_{i \in Q} b_i\right), \tag{S33c}$$

where $[n] = \{1, \ldots, n\}$ is the set listing all the random variables and $Q$ is any subset of $[n]$ including the empty set and $[n]$ itself. So, applying the distributive property, the expectation we seek becomes

$$\mathrm{E}\left[\prod_{i=1}^{n} (\mu_i + \tilde{Y}_i)\right] = \mathrm{E}\left[\sum_{Q \subseteq [n]} \left(\prod_{j \notin Q} \mu_j\right)\left(\prod_{i \in Q} \tilde{Y}_i\right)\right]. \tag{S33d}$$

Moving the expectation inside, noting that the $\mu$'s are not random variables, yields

$$\mathrm{E}\left[\prod_{i=1}^{n} Y_i\right] = \sum_{Q \subseteq [n]} \left(\prod_{j \notin Q} \mu_j\right) \mathrm{E}\left[\prod_{i \in Q} \tilde{Y}_i\right]. \tag{S33e}$$

The proof generated by Gemini ends here. Gemini stated that this formula is textbook knowledge and provided various references, but the references provided do not seem to have this exact formula.

Now, substituting $Y_i = x_{\boldsymbol{\ell}}^{(ik_i)}$ where each vector $\boldsymbol{\ell}$ denotes a different realisation of the random variable (i.e., a different haplotype), so $\mu_i = \bar{x}^{(ik_i)}$, Eq. (S33e) becomes

$$p_{\mathbf{k}} = \mathrm{E}\left[\prod_{i=1}^{n} x_{\boldsymbol{\ell}}^{(ik_i)}\right] = \sum_{Q \subseteq [n]} \left(\prod_{j \notin Q} \bar{x}^{(jk_j)}\right) \mathrm{E}\left[\prod_{i \in Q} (x_{\boldsymbol{\ell}}^{(ik_i)} - \bar{x}^{(ik_i)})\right]. \tag{S34a}$$

Denoting the haplotype content deviation as

$$X_{\boldsymbol{\ell}}^{(Q\mathbf{k})} \equiv \prod_{i \in Q} (x_{\boldsymbol{\ell}}^{(ik_i)} - \bar{x}^{(ik_i)}), \tag{S34b}$$

we obtain

$$p_{\mathbf{k}} = \sum_{Q \subseteq [n]} \left(\prod_{j \notin Q} \bar{x}^{(jk_j)}\right) \bar{X}^{(Q\mathbf{k})}, \tag{S34c}$$

where the expectation of the haplotype content deviation is

$$\bar{X}^{(Q\mathbf{k})} = \mathrm{E}[X_{\boldsymbol{\ell}}^{(Q\mathbf{k})}] = \mathrm{E}\left[\prod_{i \in Q} (x_{\boldsymbol{\ell}}^{(ik_i)} - \bar{x}^{(ik_i)})\right], \tag{S34d}$$

which is the coefficient of linkage disequilibrium of the subset of loci $Q$ in haplotype $\mathbf{k}$. The formula (S34c) is Eq. (22) in the main text.

For instance, for two loci ($n = 2$), Eq. (S34c) becomes

$$\begin{aligned}
p_{\mathbf{k}} = \mathrm{E}\left[\prod_{i=1}^{2} x_{\boldsymbol{\ell}}^{(ik_i)}\right] &= \sum_{Q \subseteq [2]} \left(\prod_{j \notin Q} \bar{x}^{(jk_j)}\right) \bar{X}^{(Q\mathbf{k})} \\
&= \left(\prod_{j \notin \emptyset} \bar{x}^{(jk_j)}\right) \bar{X}^{(\emptyset\mathbf{k})} + \left(\prod_{j \notin \{1\}} \bar{x}^{(jk_j)}\right) \bar{X}^{(\{1\}\mathbf{k})} + \left(\prod_{j \notin \{2\}} \bar{x}^{(jk_j)}\right) \bar{X}^{(\{2\}\mathbf{k})} + \left(\prod_{j \notin \{1,2\}} \bar{x}^{(jk_j)}\right) \bar{X}^{(\{1,2\}\mathbf{k})} \\
&= \prod_{j \notin \emptyset} \bar{x}^{(jk_j)} + \bar{X}^{(\{1,2\}\mathbf{k})} \\
&= \bar{x}^{(1k_1)} \bar{x}^{(2k_2)} + \bar{X}^{(\{1,2\}\mathbf{k})},
\end{aligned} \tag{S35}$$

where the third line follows by the convention that the empty product is one ($\prod_{i\in\varnothing} Y_i = 1$) and because the linkage disequilibrium coefficient is zero for single loci: $\bar{X}^{(\{i\}\mathbf{k})} = \mathrm{E}\left[(x_\ell^{(ik_i)} - \bar{x}^{(ik_i)})\right] = 0$. The last line in (S35) follows by expanding the product. Thus, Eq. (S35) writes the frequency of haplotype $\mathbf{k}$ in terms of the product of allele frequencies $\bar{x}^{(1k_1)}\bar{x}^{(2k_2)}$ and the linkage disequilibrium between the two loci $\bar{X}^{(\{1,2\}\mathbf{k})}$, which recovers the classic expression (Lewontin and Kojima, 1960).

For three loci ($n = 3$), the haplotype frequency is

$$\begin{aligned}
p_{\mathbf{k}} &= \mathrm{E}\left[\prod_{i=1}^{3} x_\ell^{(ik_i)}\right] = \sum_{Q\subseteq[3]}\left(\prod_{j\notin Q}\bar{x}^{(jk_j)}\right)\bar{X}^{(Q\mathbf{k})} \\
&= \left(\prod_{j\notin\varnothing}\bar{x}^{(jk_j)}\right)\bar{X}^{(\varnothing\mathbf{k})} + \left(\prod_{j\notin\{1\}}\bar{x}^{(jk_j)}\right)\bar{X}^{(\{1\}\mathbf{k})} + \left(\prod_{j\notin\{2\}}\bar{x}^{(jk_j)}\right)\bar{X}^{(\{2\}\mathbf{k})} + \left(\prod_{j\notin\{3\}}\bar{x}^{(jk_j)}\right)\bar{X}^{(\{3\}\mathbf{k})} \\
&\quad + \left(\prod_{j\notin\{1,2\}}\bar{x}^{(jk_j)}\right)\bar{X}^{(\{1,2\}\mathbf{k})} + \left(\prod_{j\notin\{1,3\}}\bar{x}^{(jk_j)}\right)\bar{X}^{(\{1,3\}\mathbf{k})} + \left(\prod_{j\notin\{2,3\}}\bar{x}^{(jk_j)}\right)\bar{X}^{(\{2,3\}\mathbf{k})} + \left(\prod_{j\notin\{1,2,3\}}\bar{x}^{(jk_j)}\right)\bar{X}^{(\{1,2,3\}\mathbf{k})} \\
&= \prod_{j\notin\varnothing}\bar{x}^{(jk_j)} \\
&\quad + \left(\prod_{j\notin\{1,2\}}\bar{x}^{(jk_j)}\right)\bar{X}^{(\{1,2\}\mathbf{k})} + \left(\prod_{j\notin\{1,3\}}\bar{x}^{(jk_j)}\right)\bar{X}^{(\{1,3\}\mathbf{k})} + \left(\prod_{j\notin\{2,3\}}\bar{x}^{(jk_j)}\right)\bar{X}^{(\{2,3\}\mathbf{k})} + \bar{X}^{(\{1,2,3\}\mathbf{k})} \\
&= \bar{x}^{(1k_1)}\bar{x}^{(2k_2)}\bar{x}^{(3k_3)} \\
&\quad + \bar{x}^{(3k_3)}\bar{X}^{(\{1,2\}\mathbf{k})} + \bar{x}^{(2k_2)}\bar{X}^{(\{1,3\}\mathbf{k})} + \bar{x}^{(1k_1)}\bar{X}^{(\{2,3\}\mathbf{k})} + \bar{X}^{(\{1,2,3\}\mathbf{k})}
\end{aligned} \tag{S36}$$

This writes the frequency of haplotype $\mathbf{k}$ in terms of the allele frequencies $\bar{x}^{(1k_1)}$, $\bar{x}^{(2k_2)}$, and $\bar{x}^{(3k_3)}$ at the three loci, the pairwise linkage disequilibrium coefficients between pairs of loci $\bar{X}^{(\{1,2\}\mathbf{k})}$, $\bar{X}^{(\{1,3\}\mathbf{k})}$, $\bar{X}^{(\{2,3\}\mathbf{k})}$, and the three-way linkage disequilibrium coefficient among the three loci $\bar{X}^{(\{1,2,3\}\mathbf{k})}$.

To see that following haplotype frequencies would involve the same number of variables as following allele frequencies and linkage disequilibrium, consider the following. For two biallelic loci ($n = 2$), there are 4 haplotype frequencies

$$p_{\mathrm{ab}}, p_{\mathrm{aB}}, p_{\mathrm{Ab}}, p_{\mathrm{AB}}, \tag{S37}$$

but since they add up to 1, there are 3 variables to follow. This is the same number as following allele frequencies and linkage disequilibrium (S35).

For three biallelic loci ($n = 3$), there are 8 haplotype frequencies

$$p_{\mathrm{abc}}, p_{\mathrm{abC}}, p_{\mathrm{aBc}}, p_{\mathrm{Abc}}, p_{\mathrm{aBC}}, p_{\mathrm{ABc}}, p_{\mathrm{AbC}}, p_{\mathrm{ABC}}, \tag{S38}$$

but since they add up to 1, there are 7 variables to follow. This is the same number as following allele frequencies and linkage disequilibrium (S36).

### S8 Derivation that there is only one linkage disequilibrium coefficient with two biallelic loci

To show this, we show that the covariances in gene content between loci in for two biallelic loci are equal to or the negative of the others, as previously known (Nagylaki, 1992 p. 8.69). We begin by noting that the linkage disequilibrium coefficients for two loci are covariances of gene content between the loci:

$$\bar{X}^{(\{1,2\}\mathbf{k})} = \mathrm{E}\left[\prod_{i\in\{1,2\}}(x_\ell^{(ik_i)} - \bar{x}^{(ik_i)})\right] = \mathrm{E}\left[(x_\ell^{(1k_1)} - \bar{x}^{(1k_1)})(x_\ell^{(2k_2)} - \bar{x}^{(2k_2)})\right] = \mathrm{cov}[x_\ell^{(1k_1)}, x_\ell^{(2k_2)}]. \tag{S39}$$

Consider first the linkage disequilibrium coefficient between allele a and allele b in the two loci:

$$\begin{aligned}
\bar{X}^{(\{1,2\}\text{ab})} = \operatorname{cov}[x_{\boldsymbol{\ell}}^{(1\text{a})}, x_{\boldsymbol{\ell}}^{(2\text{b})}] &= \sum_{\ell_1\in\{\text{a,A}\}}\sum_{\ell_2\in\{\text{b,B}\}} p_{\ell_1\ell_2}(x_{\ell_1\ell_2}^{(1\text{a})} - \bar{x}^{(1\text{a})})(x_{\ell_1\ell_2}^{(2\text{b})} - \bar{x}^{(2\text{b})}) \\
&= p_{\text{ab}}(x_{\text{ab}}^{(1\text{a})} - \bar{x}^{(1\text{a})})(x_{\text{ab}}^{(2\text{b})} - \bar{x}^{(2\text{b})}) \\
&\quad + p_{\text{aB}}(x_{\text{aB}}^{(1\text{a})} - \bar{x}^{(1\text{a})})(x_{\text{aB}}^{(2\text{b})} - \bar{x}^{(2\text{b})}) \\
&\quad + p_{\text{Ab}}(x_{\text{Ab}}^{(1\text{a})} - \bar{x}^{(1\text{a})})(x_{\text{Ab}}^{(2\text{b})} - \bar{x}^{(2\text{b})}) \\
&\quad + p_{\text{AB}}(x_{\text{AB}}^{(1\text{a})} - \bar{x}^{(1\text{a})})(x_{\text{AB}}^{(2\text{b})} - \bar{x}^{(2\text{b})}) \\
&= p_{\text{ab}}(1 - \bar{x}^{(1\text{a})})(1 - \bar{x}^{(2\text{b})}) \\
&\quad + p_{\text{aB}}(1 - \bar{x}^{(1\text{a})})(0 - \bar{x}^{(2\text{b})}) \\
&\quad + p_{\text{Ab}}(0 - \bar{x}^{(1\text{a})})(1 - \bar{x}^{(2\text{b})}) \\
&\quad + p_{\text{AB}}(0 - \bar{x}^{(1\text{a})})(0 - \bar{x}^{(2\text{b})}) \\
&= p_{\text{ab}}(1 - \bar{x}^{(1\text{a})})(1 - \bar{x}^{(2\text{b})}) \\
&\quad - p_{\text{aB}}(1 - \bar{x}^{(1\text{a})})\bar{x}^{(2\text{b})} \\
&\quad - p_{\text{Ab}}\bar{x}^{(1\text{a})}(1 - \bar{x}^{(2\text{b})}) \\
&\quad + p_{\text{AB}}\bar{x}^{(1\text{a})}\bar{x}^{(2\text{b})}.
\end{aligned}$$

Consider next the linkage disequilibrium coefficient between the allele A and allele B in the two loci:

$$\begin{aligned}
\bar{X}^{(\{1,2\}\text{AB})} = \operatorname{cov}[x_{\boldsymbol{\ell}}^{(1\text{A})}, x_{\boldsymbol{\ell}}^{(2\text{B})}] &= \sum_{\ell_1\in\{\text{a,A}\}}\sum_{\ell_2\in\{\text{b,B}\}} p_{\ell_1\ell_2}(x_{\ell_1\ell_2}^{(1\text{A})} - \bar{x}^{(1\text{A})})(x_{\ell_1\ell_2}^{(2\text{B})} - \bar{x}^{(2\text{B})}) \\
&= p_{\text{ab}}(x_{\text{ab}}^{(1\text{A})} - \bar{x}^{(1\text{A})})(x_{\text{ab}}^{(2\text{B})} - \bar{x}^{(2\text{B})}) \\
&\quad + p_{\text{aB}}(x_{\text{aB}}^{(1\text{A})} - \bar{x}^{(1\text{A})})(x_{\text{aB}}^{(2\text{B})} - \bar{x}^{(2\text{B})}) \\
&\quad + p_{\text{Ab}}(x_{\text{Ab}}^{(1\text{A})} - \bar{x}^{(1\text{A})})(x_{\text{Ab}}^{(2\text{B})} - \bar{x}^{(2\text{B})}) \\
&\quad + p_{\text{AB}}(x_{\text{AB}}^{(1\text{A})} - \bar{x}^{(1\text{A})})(x_{\text{AB}}^{(2\text{B})} - \bar{x}^{(2\text{B})}) \\
&= p_{\text{ab}}(0 - \bar{x}^{(1\text{A})})(0 - \bar{x}^{(2\text{B})}) \\
&\quad + p_{\text{aB}}(0 - \bar{x}^{(1\text{A})})(1 - \bar{x}^{(2\text{B})}) \\
&\quad + p_{\text{Ab}}(1 - \bar{x}^{(1\text{A})})(0 - \bar{x}^{(2\text{B})}) \\
&\quad + p_{\text{AB}}(1 - \bar{x}^{(1\text{A})})(1 - \bar{x}^{(2\text{B})}) \\
&= p_{\text{ab}}\bar{x}^{(1\text{A})}\bar{x}^{(2\text{B})} \\
&\quad - p_{\text{aB}}\bar{x}^{(1\text{A})}(1 - \bar{x}^{(2\text{B})}) \\
&\quad - p_{\text{Ab}}(1 - \bar{x}^{1\text{A}})\bar{x}^{(2\text{B})} \\
&\quad + p_{\text{AB}}(1 - \bar{x}^{(1\text{A})})(1 - \bar{x}^{(2\text{B})}) \\
&= p_{\text{ab}}(1 - \bar{x}^{(1\text{a})})(1 - \bar{x}^{(2\text{b})}) \\
&\quad - p_{\text{aB}}(1 - \bar{x}^{(1\text{a})})\bar{x}^{(2\text{b})} \\
&\quad - p_{\text{Ab}}\bar{x}^{(1\text{a})}(1 - \bar{x}^{(2\text{b})}) \\
&\quad + p_{\text{AB}}\bar{x}^{(1\text{a})}\bar{x}^{(2\text{b})} \\
&= \bar{X}^{(\{1,2\}\text{ab})}.
\end{aligned}$$

So the linkage disequilibrium coefficients $\bar{X}^{(\{1,2\}\text{ab})}$ and $\bar{X}^{(\{1,2\}\text{AB})}$ are the same.

Consider now the linkage disequilibrium coefficient between alleles a and B in the two loci:

$$
\begin{aligned}
\bar{X}^{(\{1,2\}\mathrm{aB})} &= \mathrm{cov}[x_{\ell}^{(1\mathrm{a})}, x_{\ell}^{(2\mathrm{B})}] = \sum_{\ell_1 \in \{\mathrm{a},\mathrm{A}\}} \sum_{\ell_2 \in \{\mathrm{b},\mathrm{B}\}} p_{\ell_1 \ell_2} (x_{\ell_1 \ell_2}^{(1\mathrm{a})} - \bar{x}^{(1\mathrm{a})})(x_{\ell_1 \ell_2}^{(2\mathrm{B})} - \bar{x}^{(2\mathrm{B})}) \\
&= p_{\mathrm{ab}} (x_{\mathrm{ab}}^{(1\mathrm{a})} - \bar{x}^{(1\mathrm{a})})(x_{\mathrm{ab}}^{(2\mathrm{B})} - \bar{x}^{(2\mathrm{B})}) \\
&\quad + p_{\mathrm{aB}} (x_{\mathrm{aB}}^{(1\mathrm{a})} - \bar{x}^{(1\mathrm{a})})(x_{\mathrm{aB}}^{(2\mathrm{B})} - \bar{x}^{(2\mathrm{B})}) \\
&\quad + p_{\mathrm{Ab}} (x_{\mathrm{Ab}}^{(1\mathrm{a})} - \bar{x}^{(1\mathrm{a})})(x_{\mathrm{Ab}}^{(2\mathrm{B})} - \bar{x}^{(2\mathrm{B})}) \\
&\quad + p_{\mathrm{AB}} (x_{\mathrm{AB}}^{(1\mathrm{a})} - \bar{x}^{(1\mathrm{a})})(x_{\mathrm{AB}}^{(2\mathrm{B})} - \bar{x}^{(2\mathrm{B})}) \\
&= p_{\mathrm{ab}} (1 - \bar{x}^{(1\mathrm{a})})(0 - \bar{x}^{(2\mathrm{B})}) \\
&\quad + p_{\mathrm{aB}} (1 - \bar{x}^{(1\mathrm{a})})(1 - \bar{x}^{(2\mathrm{B})}) \\
&\quad + p_{\mathrm{Ab}} (0 - \bar{x}^{(1\mathrm{a})})(0 - \bar{x}^{(2\mathrm{B})}) \\
&\quad + p_{\mathrm{AB}} (0 - \bar{x}^{(1\mathrm{a})})(1 - \bar{x}^{(2\mathrm{B})}) \\
&= -p_{\mathrm{ab}} (1 - \bar{x}^{(1\mathrm{a})}) \bar{x}^{(2\mathrm{B})} \\
&\quad + p_{\mathrm{aB}} (1 - \bar{x}^{(1\mathrm{a})})(1 - \bar{x}^{(2\mathrm{B})}) \\
&\quad + p_{\mathrm{Ab}} \bar{x}^{(1\mathrm{a})} \bar{x}^{(2\mathrm{B})} \\
&\quad - p_{\mathrm{AB}} \bar{x}^{(1\mathrm{a})} (1 - \bar{x}^{(2\mathrm{B})}) \\
&= -p_{\mathrm{ab}} (1 - \bar{x}^{(1\mathrm{a})})(1 - \bar{x}^{(2\mathrm{b})}) \\
&\quad + p_{\mathrm{aB}} (1 - \bar{x}^{(1\mathrm{a})}) \bar{x}^{(2\mathrm{b})} \\
&\quad + p_{\mathrm{Ab}} \bar{x}^{(1\mathrm{a})} (1 - \bar{x}^{(2\mathrm{b})}) \\
&\quad - p_{\mathrm{AB}} \bar{x}^{(1\mathrm{a})} \bar{x}^{(2\mathrm{b})} \\
&= -\bar{X}^{(\{1,2\}\mathrm{ab})}.
\end{aligned}
$$

So the linkage disequilibrium coefficient $\bar{X}^{(\{1,2\}\mathrm{aB})}$ is the negative of the previous two.

Consider finally the linkage disequilibrium coefficient between alleles A and b in the two loci:

$$
\begin{aligned}
\bar{X}^{(\{1,2\}\mathrm{Ab})} &= \mathrm{cov}[x_{\ell}^{(1\mathrm{A})}, x_{\ell}^{(2\mathrm{b})}] = \sum_{\ell_1 \in \{\mathrm{a},\mathrm{A}\}} \sum_{\ell_2 \in \{\mathrm{b},\mathrm{B}\}} p_{\ell_1 \ell_2} (x_{\ell_1 \ell_2}^{(1\mathrm{A})} - \bar{x}^{(1\mathrm{A})})(x_{\ell_1 \ell_2}^{(2\mathrm{b})} - \bar{x}^{(2\mathrm{b})}) \\
&= p_{\mathrm{ab}} (x_{\mathrm{ab}}^{(1\mathrm{A})} - \bar{x}^{(1\mathrm{A})})(x_{\mathrm{ab}}^{(2\mathrm{b})} - \bar{x}^{(2\mathrm{b})}) \\
&\quad + p_{\mathrm{aB}} (x_{\mathrm{aB}}^{(1\mathrm{A})} - \bar{x}^{(1\mathrm{A})})(x_{\mathrm{aB}}^{(2\mathrm{b})} - \bar{x}^{(2\mathrm{b})}) \\
&\quad + p_{\mathrm{Ab}} (x_{\mathrm{Ab}}^{(1\mathrm{A})} - \bar{x}^{(1\mathrm{A})})(x_{\mathrm{Ab}}^{(2\mathrm{b})} - \bar{x}^{(2\mathrm{b})}) \\
&\quad + p_{\mathrm{AB}} (x_{\mathrm{AB}}^{(1\mathrm{A})} - \bar{x}^{(1\mathrm{A})})(x_{\mathrm{AB}}^{(2\mathrm{b})} - \bar{x}^{(2\mathrm{b})}) \\
&= p_{\mathrm{ab}} (0 - \bar{x}^{(1\mathrm{A})})(1 - \bar{x}^{(2\mathrm{b})}) \\
&\quad + p_{\mathrm{aB}} (0 - \bar{x}^{(1\mathrm{A})})(0 - \bar{x}^{(2\mathrm{b})}) \\
&\quad + p_{\mathrm{Ab}} (1 - \bar{x}^{(1\mathrm{A})})(1 - \bar{x}^{(2\mathrm{b})}) \\
&\quad + p_{\mathrm{AB}} (1 - \bar{x}^{(1\mathrm{A})})(0 - \bar{x}^{(2\mathrm{b})}) \\
&= -p_{\mathrm{ab}} \bar{x}^{(1\mathrm{A})} (1 - \bar{x}^{(2\mathrm{b})}) \\
&\quad + p_{\mathrm{aB}} \bar{x}^{(1\mathrm{A})} \bar{x}^{(2\mathrm{b})} \\
&\quad + p_{\mathrm{Ab}} (1 - \bar{x}^{(1\mathrm{A})})(1 - \bar{x}^{(2\mathrm{b})}) \\
&\quad - p_{\mathrm{AB}} (1 - \bar{x}^{(1\mathrm{A})}) \bar{x}^{(2\mathrm{b})} \\
&= -p_{\mathrm{ab}} (1 - \bar{x}^{(1\mathrm{a})})(1 - \bar{x}^{(2\mathrm{b})}) \\
&\quad + p_{\mathrm{aB}} (1 - \bar{x}^{(1\mathrm{a})}) \bar{x}^{(2\mathrm{b})} \\
&\quad + p_{\mathrm{Ab}} \bar{x}^{(1\mathrm{a})} (1 - \bar{x}^{(2\mathrm{b})}) \\
&\quad - p_{\mathrm{AB}} \bar{x}^{(1\mathrm{a})} \bar{x}^{(2\mathrm{b})} \\
&= -\bar{X}^{(\{1,2\}\mathrm{ab})}.
\end{aligned}
$$

So the linkage disequilibrium coefficient $\bar{X}^{(\{1,2\}\mathrm{Ab})}$ is also the negative of the first two. Consequently, there is only one independent linkage disequilibrium coefficient $\bar{X} \equiv \bar{X}^{(\{1,2\}\mathrm{ab})} = \bar{X}^{(\{1,2\}\mathrm{AB})} = -\bar{X}^{(\{1,2\}\mathrm{aB})} = -\bar{X}^{(\{1,2\}\mathrm{Ab})}$.

### S9 Additional examples

#### S9.1 Example 2 continued: under selfing, transmission matrix can be slightly negative

We now illustrate the possibility of negative transmission covariance in the same situation as in example 2 but with selfing rather than random mating. So we first compute the conditional expectations of offspring phenotype given parent phenotype: $z'(z) = \mathrm{E}_{z^+|z}[z^+|z]$. With selfing, the expected phenotype of offspring given that a parent has phenotype $z_{aa}$ is

$$\begin{aligned} z'(z_{\mathrm{aa}}) &= z_{\mathrm{aa}} \\ &= 0. \end{aligned}$$

Similarly, the expected phenotype of offspring from a parent with phenotype $z_{\mathrm{Aa}}$ is

$$\begin{aligned} z'(z_{\mathrm{Aa}}) &= \frac{1}{4} z_{\mathrm{aa}} + \frac{1}{2} z_{\mathrm{Aa}} + \frac{1}{4} z_{\mathrm{AA}} \\ &= \frac{1}{2} c(2+d). \end{aligned}$$

Finally, the expected phenotype of offspring from a parent with phenotype $z_{\mathrm{AA}}$ is

$$\begin{aligned} z'(z_{\mathrm{AA}}) &= z_{\mathrm{AA}} \\ &= 2c. \end{aligned}$$

Given random meiosis, the genotype frequencies are in Hardy-Weinberg equilibrium. Thus, the covariance between expected offspring phenotype and parent phenotype is

$$T = \mathrm{cov}[z', z] = \sum_{k,l \in \mathbb{H}} p_k p_l z'(z_{kl})(z_{kl} - \bar{z}) = c^2 P_x [2 + d(3 + d - 6\bar{x} - 2dP_x)], \tag{S40}$$

which reduces to $T = 2c^2 P_x = 2c^2 \bar{x}(1-\bar{x})$ if $d = 0$. Note that here $T \neq P_x D^2$ as in the random mating case even though the same $D$ applies here because $D$ does not depend on reproduction, which is all we have changed. Here $T$ can be slightly negative for a very narrow set of allele frequencies and dominance coefficients (Fig. S1t, but it is difficult to visualise there given its small negative magnitude). For instance, $T = -0.00115347c^2$ for $\bar{x} = 0.99$ and $d = 1.2$.

Regarding the regression of expected offspring phenotype on parent phenotype (with intercept $\zeta$ and slope $H$), from least squares, it follows that

$$\begin{aligned} \zeta &= \mathrm{E}[z'] = \sum_{k,l \in \mathbb{H}} p_k p_l z'(z_{kl}) \\ H &= \frac{\mathrm{cov}[z', z]}{\mathrm{var}[z]} = \frac{\sum_{k,l \in \mathbb{H}} p_k p_l z'(z_{kl})(z_{kl} - \bar{z})}{\sum_{k,l \in \mathbb{H}} p_k p_l (z_{kl} - \bar{z})^2}. \end{aligned}$$

Plugging the values specified, this yields

$$\begin{aligned} \zeta &= c(2+d)\bar{x} - cd\bar{x}^2 \neq \bar{z} \\ H &= \frac{T}{P}. \end{aligned} \tag{S41}$$

Thus, in contrast to the random mating case, here there is pure transmission bias in the phenotype ($\mathrm{E}[z'] - \bar{z} = -cd\bar{x}(1-\bar{x})$) if there is dominance, which vanishes with additive allelic effects $d = 0$ (see also Frank 1997, his Appendix A). If the allelic effect is additive, that is $d = 0$, $H$ reduces to $H = 1$, but if $d \neq 0$, then generally $H \neq 1$ (Fig. S1v-ab). In contrast to heritability, heredity can be greater than 1 (Fig. S1w) or smaller than zero (Fig. S1aa), a point already noted by Rice (2004) (p. 205). Even though transmission $T$ can only be here slightly negative, heredity $H$ can be strongly negative due to division by small phenotypic variance $P$ (Fig. S1aa). For instance, $H = -0.97929$ for $\bar{x} = 0.99$ and $d = 1.2$.

#### S9.2 Example 3 continued: exact method by coupling genetic and phenotypic evolution in generator equations

Here we continue our example 3 in the main text using the exact method. The aim here is to determine whether a dynamically sufficient adaptive topography for the mean phenotype could be obtained by applying the generator equations, not just to the haplotype as in the main text, but to both the haplotype and phenotype. Doing

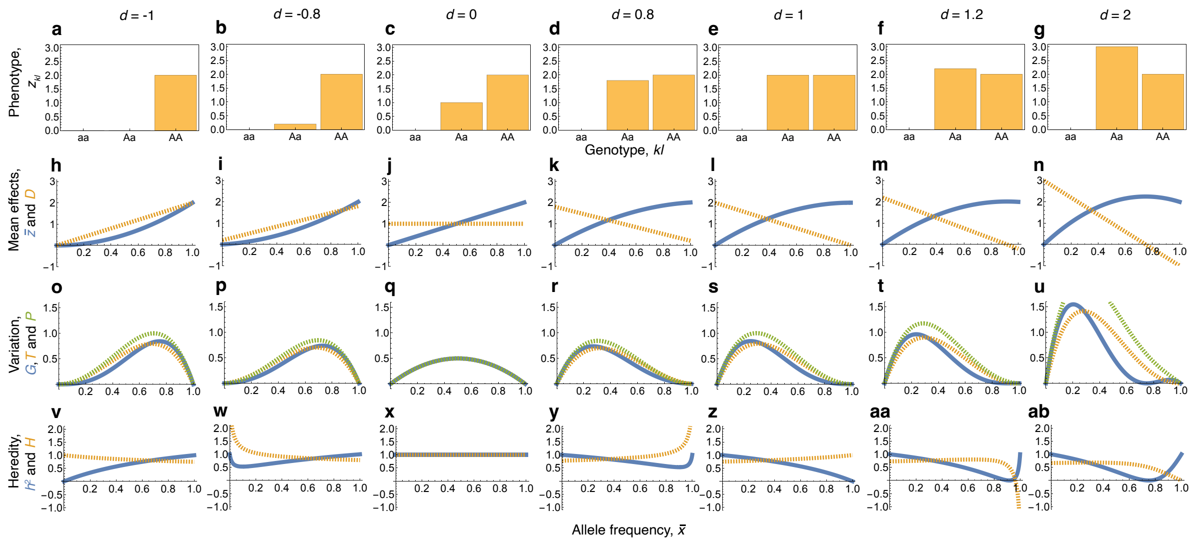


Figure S1: Example 2 continued: evolutionary coefficients under selfing. Compare with Fig. 2. The only differences with that figure are for $T$ and $H$, in panels **a-ab**. Note the differences in slope for $H$ relative to Fig. 2: in instances where heredity decreases with allele frequency here, it may increase in Fig. 2, and viceversa. Note also the negative heredity in **aa**. In all panels, $c = 1$.

so does generate a dynamically sufficient system of equations, by describing the change of both mean haplotype and mean phenotype, while being in terms of the selection regression coefficient of both. However, as we will see, the resulting system does not describe an adaptive topography for the mean phenotype.

To do this, consider the vector of phenotype and haplotype content $\mathbf{m}_{kl} = (z_{kl}, x_k)$ for genotype $kl$. Similarly, consider the expected offspring vector $\mathbf{m}'_{kl} = (z'_{kl}, x'_k)^\intercal$ and the mean vector $\bar{\mathbf{m}} = (\bar{z}, \bar{x})^\intercal$. Applying the generator equations to this vector, the evolution of the mean phenotype and allele frequency is here given by

$$\Delta\bar{\mathbf{m}} = \mathbf{T_m}\boldsymbol{\beta}_\mathbf{m} + \mathbf{u_m} = \mathbf{P}'_\mathbf{m}\boldsymbol{\beta}'_\mathbf{m},$$

since there is no pure transmission bias in the phenotype and haplotype ($\boldsymbol{\zeta}_\mathbf{m} = (\zeta, \zeta_x)^\intercal = (\bar{z}, \bar{x})^\intercal$) as described in the main text.

We first derive the components of the primary generator equation. The transmission matrix of phenotype and haplotype is

$$\mathbf{T_m} = \mathrm{cov}[\mathbf{m}', \mathbf{m}] = \sum_{k,l \in \mathbb{H}} p_k p_l \mathbf{m}'_{kl}(\mathbf{m}_{kl} - \bar{\mathbf{m}})^\intercal,$$

This yields

$$\mathbf{T_m} = P_x \begin{pmatrix} D^2 & H_x D \\ H_x D & H_x \end{pmatrix} = \begin{pmatrix} T & T_x D \\ T_x D & T_x \end{pmatrix}. \tag{S42}$$

where as before $D = c[1 + d(1 - 2\bar{x})]$, $P_x = \bar{x}(1 - \bar{x})$, and $H_x = 1/2$. Thus, $\det(\mathbf{T_m}) = P_x^2 D^2/4$, so $\mathbf{T_m}$ is invertible if $D \neq 0$ and $P_x \neq 0$. Also, $\mathbf{T_m}$ is symmetric and its eigenvalues are non-negative. Thus, the linear selection response has a direction that is less than or at 90° from the selection regression coefficient $\boldsymbol{\beta}_\mathbf{m}$. To compute the selection regression coefficient $\boldsymbol{\beta}_\mathbf{m} = \mathbf{P}_\mathbf{m}^{-1}\mathrm{cov}[\mathbf{m}, w]$, note that $\mathbf{P_m} = \sum_{k,l \in \mathbb{H}} p_k p_l (\mathbf{m}_{kl} - \bar{\mathbf{m}})(\mathbf{m}_{kl} - \bar{\mathbf{m}})^\intercal$ which yields

$$\mathbf{P_m} = \begin{pmatrix} P & P_x D \\ P_x D & P_x \end{pmatrix},$$

where $P$ is as found in Example 2. Therefore, assuming that fitnesses are symmetric (i.e., $w_{\mathrm{Aa}} = w_{\mathrm{aA}}$), we obtain

$$\boldsymbol{\beta}_\mathbf{m} = \begin{pmatrix} \dfrac{w_{\mathrm{AA}}\bar{x}(1-d) + w_{\mathrm{Aa}}(1 + d - 2\bar{x}) - w_{\mathrm{aa}}(1+d)(1-\bar{x})}{c[(1+d)^2 - 4d\bar{x}]} \\ \dfrac{2d\bar{x}(1-\bar{x})[(1+d)w_{\mathrm{AA}} - 2w_{\mathrm{Aa}} + (1-d)w_{\mathrm{aa}}]}{(1+d)^2 - 4d\bar{x}} \end{pmatrix},$$

Recall that $\beta_x = w_{\mathrm{A}} - w_{\mathrm{a}}$ in this example. It can be checked that

$$\beta_x = \tilde{\beta}_z D + \tilde{\beta}_x, \tag{S43}$$

where $\tilde{\beta}_z$ and $\tilde{\beta}_x$ are respectively the first and second entries of $\boldsymbol{\beta}_{\mathbf{m}}$ (Eq. 28 of main text). Note that we have not specified how the form of the fitness functions $w_{kl}$: they could be direct functions of the phenotype $z_{kl}$ but also of the haplotype content $x_k$ or $x_l$. Regardless, $\boldsymbol{\beta}_{\mathbf{m}}$ with $d = 0$ reduces to

$$\boldsymbol{\beta}_{\mathbf{m}} = \begin{pmatrix} \frac{1}{c}\left[w_{\mathrm{AA}}\bar{x} + w_{\mathrm{Aa}}(1-2\bar{x}) - w_{\mathrm{aa}}(1-\bar{x})\right] \\ 0 \end{pmatrix}.$$

So, even if the fitness functions are not direct functions of the phenotype $z_{kl}$ but only of haplotype content $x_k$ or $x_l$, and the phenotype is additive $d = 0$, $\tilde{\beta}_x = 0$ which would suggest that there is no direct selection for the haplotype, but there is. This indicates that the interpretation of the direct regression coefficients $\tilde{\beta}$ as estimates of direct selection is problematic.

Thus, while $\beta_x$ can be written as a selection gradient of allele frequency, it is thus not immediately clear that $\boldsymbol{\beta}_{\mathbf{m}}$ can be written in terms of a selection gradient either of allele frequency or of mean phenotype and allele frequency $\bar{\mathbf{m}} = (\bar{z}, \bar{x})^{\intercal}$.

Now, to calculate the selection response from non-linear inheritance $\mathbf{u}_{\mathbf{m}}$, from (4) in the main text, note that $\mathbf{u}_{\mathbf{m}} = \mathrm{cov}[\mathbf{m}', w] - \mathbf{H}_{\mathbf{m}}\mathrm{cov}[\mathbf{m}, w]$. The heredity matrix of the phenotype and haplotype is $\mathbf{H}_{\mathbf{m}} = \mathbf{T}_{\mathbf{m}}\mathbf{P}_{\mathbf{m}}^{-1}$, which yields

$$\mathbf{H}_{\mathbf{m}} = \begin{pmatrix} \dfrac{D^2}{2c^2[(1+d)^2 - 4d\bar{x}]} & \dfrac{2d^2 P_x D}{(1+d)^2 - 4d\bar{x}} \\ 0 & H_x \end{pmatrix},$$

which reduces to $\mathbf{H}_{\mathbf{m}} = \frac{1}{2}\mathbf{I}$ if $d = 0$. The top left entry is not the $H$ we had found and can now be negative for $d < -1$ or $d > 1$. We also obtain that the total selection response is

$$\mathrm{cov}[\mathbf{m}', w] = P_x \begin{pmatrix} D \\ H_x \end{pmatrix} \beta_x.$$

Hence, we obtain

$$\mathbf{u}_{\mathbf{m}} = \begin{pmatrix} -\dfrac{dP_x^2 D^2[(1+d)w_{\mathrm{AA}} - 2w_{\mathrm{Aa}} + (1-d)w_{\mathrm{aa}}]}{(1+d)^2 - 4d\bar{x}]} \\ 0 \end{pmatrix},$$

which is zero if $d = 0$ but not otherwise. We thus obtain that the change in the mean phenotype and allele frequency can be written as

$$\Delta\bar{\mathbf{m}} = \mathbf{T}_{\mathbf{m}}\boldsymbol{\beta}_{\mathbf{m}} + \mathbf{u}_{\mathbf{m}} = P_x \begin{pmatrix} D \\ H_x \end{pmatrix} \beta_x = \frac{1}{2} P_x \begin{pmatrix} D \\ H_x \end{pmatrix} \frac{1}{\bar{W}} \frac{\mathrm{d}\bar{W}}{\mathrm{d}\bar{x}}. \tag{S44}$$

However, this is not an adaptive topography for the mean phenotype. Specifically, evolution thus proceeds in the direction $(D, 1/2)^{\intercal}$ (not a selection gradient) and its magnitude is scaled by the selection gradient of allele frequency. In particular, the mean phenotype evolves in the opposite direction of allele frequency change when $D$ is negative, for instance, for overdominance ($d > 1$) and sufficiently high allele frequency (Fig. 2m,n). Yet, this equation does not describe the evolution of the mean phenotype as the climbing of a fitness landscape in phenotype space. This is not possible in this model because mean fitness cannot be written as a function of the mean phenotype, at least with the fitness function and genotype-phenotype map used.

To further analyse the direction of evolution, let us consider the secondary generator equation, which depends on the selection pointer that better approaches such direction. We have that the covariance matrix of offspring phenotype and haplotype is

$$\mathbf{P}'_{\mathbf{m}} = \mathrm{cov}[\mathbf{m}', \mathbf{m}'] = \sum_{k,l \in \mathbb{H}} p_k p_l \mathbf{m}'_{kl}(\mathbf{m}'_{kl} - \boldsymbol{\zeta}_{\mathbf{m}})^{\intercal}.$$

This yields

$$\mathbf{P}'_{\mathbf{m}} = \frac{1}{2} P_x \begin{pmatrix} D^2 & H_x D \\ H_x D & H_x \end{pmatrix},$$

so $\mathbf{P}'_{\mathbf{m}}$ is invertible if $D \neq 0$ and $P_x \neq 0$. In turn, the selection pointer is $\beta'_{\mathbf{m}} = \mathbf{P}'^{-1}_{\mathbf{m}}\mathrm{cov}[\mathbf{m}', w]$, which yields

$$\beta'_{\mathbf{m}} = \begin{pmatrix} 2\dfrac{w_{\mathrm{A}} - w_{\mathrm{a}}}{D} \\ 0 \end{pmatrix} = \begin{pmatrix} 1 \\ 0 \end{pmatrix} \frac{1}{D} \frac{1}{\bar{W}} \frac{\mathrm{d}\bar{W}}{\mathrm{d}\bar{x}}.$$

That the partial selection pointer of haplotype content is zero is surprising because we have not specified whether or not fitness depends directly on gene content. With this, the change in mean phenotype and allele frequency is again

$$\Delta\bar{\mathbf{m}} = \mathbf{P}'_{\mathbf{m}}\beta'_{\mathbf{m}} = \frac{1}{2}P_x \begin{pmatrix} D \\ H_x \end{pmatrix} \frac{1}{\bar{W}} \frac{\mathrm{d}\bar{W}}{\mathrm{d}\bar{x}}.$$

Since $\mathbf{P}'_{\mathbf{m}}$ is a covariance matrix, this direction of evolution is at less than $< 90°$ of the selection pointer $\beta'_{\mathbf{m}}$, whose direction is that of $(1,0)^\intercal$ (again, not a selection gradient) and its magnitude is scaled by the selection gradient of allele frequency. It can be checked that the angle between the selection pointer $\beta'_{\mathbf{m}}$ and the selection regression coefficient $\beta_{\mathbf{m}}$ can be greater than 90° because their dot product can be negative. Yet, the selection pointer $\beta'_{\mathbf{m}}$ more closely approaches the direction of evolution, particularly with non-additive allelic effects ($d \neq 0$).

#### S9.3 Example 4 continued: following haplotype frequencies rather than allele frequencies and linkage disequilibrium

Here we illustrate the possibility of applying the generator equations to follow changes in haplotype frequencies, rather than changes in allele frequencies and linkage disequilibria. This shows that the former approach is easier to set up but yields more complicated expressions for genetic evolution.

Let $x^{(\mathbf{k})}_{\ell} = \delta_{\ell\mathbf{k}}$ be the haplotype content of haplotype $\ell$ for haplotype $\mathbf{k}$, where $\delta_{\ell\mathbf{k}}$ is Kronecker's delta. For instance, for two biallelic loci, $x^{(\mathrm{ab})}_{\mathrm{ab}} = 1$, which means that the haplotype ab is present, whereas $x^{(\mathrm{ab})}_{\mathrm{Ab}} = 0$, which means that the haplotype ab is absent. The frequency $p_{\mathbf{k}}$ of haplotype $\mathbf{k}$ is the average haplotype content: $p_{\mathbf{k}} = \sum_{\ell\in\mathbb{H}} p_\ell x^{(\mathbf{k})}_{\ell} = \bar{x}^{(\mathbf{k})}$ (since $x^{(\mathbf{k})}_{\ell}$ is always 0 except when $\ell = \mathbf{k}$, in which case it is 1). So, let us form the vector $\mathbf{x}_\ell$ of haplotype content of haplotype $\ell$, which lists the haplotype content $x^{(\mathbf{k})}_{\ell}$ of haplotype $\ell$ for all haplotypes $\mathbf{k}$, excepting one since haplotype frequencies add up to one so we only need to track all haplotype frequencies excepting an arbitrarily chosen one. For instance, for two biallelic loci, leaving haplotype ab out, the haplotype content of haplotype $\mathbf{k}$ is

$$\mathbf{x}_{\mathbf{k}} = (x^{(\mathrm{aB})}_{\mathbf{k}}, x^{(\mathrm{Ab})}_{\mathbf{k}}, x^{(\mathrm{AB})}_{\mathbf{k}})^\intercal, \tag{S45}$$

so the haplotype content of haplotype AB is

$$\mathbf{x}_{\mathrm{AB}} = (x^{(\mathrm{aB})}_{\mathrm{AB}}, x^{(\mathrm{Ab})}_{\mathrm{AB}}, x^{(\mathrm{AB})}_{\mathrm{AB}})^\intercal = (0,0,1)^\intercal. \tag{S46}$$

The average vector $\bar{\mathbf{x}} = \sum_{\ell\in\mathbb{H}} p_\ell \mathbf{x}_\ell$ of haplotype content lists the haplotype frequencies $\bar{x}^{(\mathbf{k})}$ for all haplotypes $\mathbf{k}$, except the one left out. For instance, for two biallelic loci,

$$\bar{\mathbf{x}} = (\bar{x}^{(\mathrm{aB})}, \bar{x}^{(\mathrm{Ab})}, \bar{x}^{(\mathrm{AB})})^\intercal = (p_{\mathrm{aB}}, p_{\mathrm{Ab}}, p_{\mathrm{AB}})^\intercal. \tag{S47}$$

This vector has 3 entries as in example 5 and lists all the genetic variables we need to track (since $p_{\mathrm{ab}} = 1 - p_{\mathrm{aB}} - p_{\mathrm{Ab}} - p_{\mathrm{AB}}$).

Thus, applying the generator equations to the vector of haplotype content $\mathbf{x}_\ell$, we have that the change in haplotype frequencies for arbitrarily many loci with arbitrarily many alleles, under arbitrary linkage disequilibria and recombination frequencies, is in general given by

$$\Delta\bar{\mathbf{x}} = \mathbf{T}_{\mathbf{x}}\beta_{\mathbf{x}} + \mathbf{u}_{\mathbf{x}} + \zeta_{\mathbf{x}} - \bar{\mathbf{x}} = \mathbf{P}'_{\mathbf{x}}\beta'_{\mathbf{x}} + \zeta_{\mathbf{x}} - \bar{\mathbf{x}}. \tag{S48}$$

We now derive $\mathbf{T}_{\mathbf{x}}$ and $\beta_{\mathbf{x}}$ for the case of two biallelic loci for the vector of haplotype content (S45), so tracking the frequencies of haplotypes aB, Ab, and AB. This is equivalent to case considered in example 4 in the main text but there we follow allele frequencies and linkage disequilibrium.

We have that $\mathbf{T}_{\mathbf{x}} = \mathrm{cov}[\mathbf{x}',\mathbf{x}] = \mathbf{H}_{\mathbf{x}}\mathbf{P}_{\mathbf{x}}$, where $\mathbf{x}'$ is the expected haplotype content among offspring conditional on parent haplotype content. The covariance matrix of haplotype content is

$$\mathbf{P}_{\mathbf{x}} = \mathrm{cov}[\mathbf{x},\mathbf{x}] = \sum_{\mathbf{k}\in\mathbb{H}} p_{\mathbf{k}}(\mathbf{x}_{\mathbf{k}} - \bar{\mathbf{x}})(\mathbf{x}_{\mathbf{k}} - \bar{\mathbf{x}})^\intercal = \begin{pmatrix} \bar{x}^{(\mathrm{aB})}(1-\bar{x}^{(\mathrm{aB})}) & -\bar{x}^{(\mathrm{aB})}\bar{x}^{(\mathrm{Ab})} & -\bar{x}^{(\mathrm{aB})}\bar{x}^{(\mathrm{AB})} \\ -\bar{x}^{(\mathrm{Ab})}\bar{x}^{(\mathrm{aB})} & \bar{x}^{(\mathrm{Ab})}(1-\bar{x}^{(\mathrm{Ab})}) & -\bar{x}^{(\mathrm{Ab})}\bar{x}^{(\mathrm{AB})} \\ -\bar{x}^{(\mathrm{AB})}\bar{x}^{(\mathrm{aB})} & -\bar{x}^{(\mathrm{AB})}\bar{x}^{(\mathrm{Ab})} & \bar{x}^{(\mathrm{AB})}(1-\bar{x}^{(\mathrm{AB})}) \end{pmatrix}. \tag{S49}$$

Notice that while the matrix in eq. (123) becomes diagonal without linkage disequilibrium, the matrix in (S49) may not.

We now compute the expected offspring haplotype contents, $\mathbf{x}'$. Analogously to eq. 125, the expected haplotype content for haplotype $\ell$ among offspring of parental haplotype $\mathbf{k}$ is

$$x'^{(\ell)}(\mathbf{x}_\mathbf{k}) = \sum_{\mathbf{l}\in\mathbb{H}} p_\mathbf{l} \sum_{\mathbf{n}\in\mathbb{H}} R_{\mathbf{kln}} x_\mathbf{n}^{(\ell)}, \tag{S50}$$

where $R_{\mathbf{kln}}$ is the probability that genotype $\mathbf{kl}$ produces gametes with haplotype $\mathbf{n}$, as given in eq. 126, originally from Nagylaki (eq. 8.9 of 1992). Doing the indicated calculations, the vector $\mathbf{x}'$ takes the values

$$\begin{aligned}
\mathbf{x}'(\mathbf{x}_\mathrm{ab}) &= \frac{1}{2}\left(\bar{x}^{(\mathrm{aB})} + r\bar{x}^{(\mathrm{AB})}, \bar{x}^{(\mathrm{Ab})} + r\bar{x}^{(\mathrm{AB})}, (1-r)\bar{x}^{(\mathrm{AB})}\right)^\intercal \\
\mathbf{x}'(\mathbf{x}_\mathrm{aB}) &= \frac{1}{2}\left(1 + \bar{x}^{(\mathrm{aB})} - r\bar{x}^{(\mathrm{Ab})}, (1-r)\bar{x}^{(\mathrm{Ab})}, r\bar{x}^{(\mathrm{Ab})} + \bar{x}^{(\mathrm{AB})}\right)^\intercal \\
\mathbf{x}'(\mathbf{x}_\mathrm{Ab}) &= \frac{1}{2}\left((1-r)\bar{x}^{(\mathrm{aB})}, 1 - r\bar{x}^{(\mathrm{aB})} + \bar{x}^{(\mathrm{Ab})}, r\bar{x}^{(\mathrm{aB})} + \bar{x}^{(\mathrm{AB})}\right)^\intercal \\
\mathbf{x}'(\mathbf{x}_\mathrm{AB}) &= \frac{1}{2}\left(\bar{x}^{(\mathrm{aB})} + r\bar{x}^{(\mathrm{ab})}, \bar{x}^{(\mathrm{Ab})} + r\bar{x}^{(\mathrm{ab})}, 1 + \bar{x}^{(\mathrm{AB})} - r\bar{x}^{(\mathrm{ab})}\right)^\intercal.
\end{aligned} \tag{S51}$$

Hence, the mean offspring haplotype content is

$$\begin{aligned}
\zeta_\mathbf{x} = \mathrm{E}[\mathbf{x}'] &= \sum_{\mathbf{k}\in\mathbb{H}} p_\mathbf{k}\mathbf{x}'(\mathbf{x}_\mathbf{k}) \\
&= \begin{pmatrix}
\bar{x}^{(\mathrm{aB})} + r\bar{x}^{(\mathrm{AB})}\left(1 - \bar{x}^{(\mathrm{Ab})} - \bar{x}^{(\mathrm{AB})}\right) - r\bar{x}^{(\mathrm{aB})}\left(\bar{x}^{(\mathrm{Ab})} + \bar{x}^{(\mathrm{AB})}\right) \\
\bar{x}^{(\mathrm{Ab})} + r\bar{x}^{(\mathrm{AB})}\left(1 - \bar{x}^{(\mathrm{aB})} - \bar{x}^{(\mathrm{AB})}\right) - r\bar{x}^{(\mathrm{Ab})}\left(\bar{x}^{(\mathrm{aB})} + \bar{x}^{(\mathrm{AB})}\right) \\
\bar{x}^{(\mathrm{AB})} + r\bar{x}^{(\mathrm{AB})}\left(1 - \bar{x}^{(\mathrm{Ab})} - \bar{x}^{(\mathrm{AB})}\right) - r\bar{x}^{(\mathrm{aB})}\left(\bar{x}^{(\mathrm{Ab})} + \bar{x}^{(\mathrm{AB})}\right)
\end{pmatrix}.
\end{aligned} \tag{S52}$$

Pure transmission bias in haplotype content is then

$$\zeta_\mathbf{x} - \bar{\mathbf{x}} = \begin{pmatrix}
r\bar{x}^{(\mathrm{AB})}\left(1 - \bar{x}^{(\mathrm{Ab})} - \bar{x}^{(\mathrm{AB})}\right) - r\bar{x}^{(\mathrm{aB})}\left(\bar{x}^{(\mathrm{Ab})} + \bar{x}^{(\mathrm{AB})}\right) \\
r\bar{x}^{(\mathrm{AB})}\left(1 - \bar{x}^{(\mathrm{aB})} - \bar{x}^{(\mathrm{AB})}\right) - r\bar{x}^{(\mathrm{Ab})}\left(\bar{x}^{(\mathrm{aB})} + \bar{x}^{(\mathrm{AB})}\right) \\
r\bar{x}^{(\mathrm{AB})}\left(1 - \bar{x}^{(\mathrm{Ab})} - \bar{x}^{(\mathrm{AB})}\right) - r\bar{x}^{(\mathrm{aB})}\left(\bar{x}^{(\mathrm{Ab})} + \bar{x}^{(\mathrm{AB})}\right)
\end{pmatrix}. \tag{S53}$$

These expressions for transmission bias are much more complicated and difficult to interpret than those in eqs. (129-130).

Then, the transmission matrix of haplotype content is

$$\mathbf{T}_\mathbf{x} = \mathrm{cov}[\mathbf{x}', \mathbf{x}] = \sum_{\mathbf{k}\in\mathbb{H}} p_\mathbf{k}(\mathbf{x}'(\mathbf{x}_\mathbf{k}) - \mathrm{E}[\mathbf{x}'])(\mathbf{x}_\mathbf{k} - \bar{\mathbf{x}})^\intercal = \mathbf{H}_\mathbf{x}\mathbf{P}_\mathbf{x}, \tag{S54}$$

which yields an expression that is too complicated to fit on the page, but the heredity matrix is

$$\mathbf{H}_\mathbf{x} = \mathbf{T}_\mathbf{x}\mathbf{P}_\mathbf{x}^{-1} = \frac{1}{2}\begin{pmatrix}
1 - r(\bar{x}^{(\mathrm{Ab})} + \bar{x}^{(\mathrm{AB})}) & -r(\bar{x}^{(\mathrm{aB})} + \bar{x}^{(\mathrm{AB})}) & r(\bar{x}^{(\mathrm{ab})} - \bar{x}^{(\mathrm{AB})}) \\
-r(\bar{x}^{(\mathrm{Ab})} + \bar{x}^{(\mathrm{AB})}) & 1 - r(\bar{x}^{(\mathrm{aB})} + \bar{x}^{(\mathrm{AB})}) & r(\bar{x}^{(\mathrm{ab})} - \bar{x}^{(\mathrm{AB})}) \\
r(\bar{x}^{(\mathrm{Ab})} + \bar{x}^{(\mathrm{AB})}) & r(\bar{x}^{(\mathrm{aB})} + \bar{x}^{(\mathrm{AB})}) & 1 - r(\bar{x}^{(\mathrm{ab})} - \bar{x}^{(\mathrm{AB})})
\end{pmatrix}. \tag{S55}$$

These expressions are much more complicated than those in eqs. (131-132).

From (3), we have that the residual of regressing expected offspring haplotype on parent haplotype is $\eta_{\mathbf{x}_\mathbf{k}} = \mathbf{x}'_\mathbf{k} - \zeta_\mathbf{x} - \mathbf{H}_\mathbf{x}(\mathbf{x}_\mathbf{k} - \bar{\mathbf{x}})$. Doing this calculation yields that $\eta_{\mathbf{x}_\mathbf{k}} = \mathbf{0}$ for all $\mathbf{k} \in \mathbb{H}$, so the expected offspring haplotype is exactly given by its linear regression on parent haplotype content. Hence, from (5), there cannot be selection response of haplotype content from non-linear inheritance, that is, $\mathbf{u}_\mathbf{x} = \mathbf{0}$.

The selection regression coefficient of haplotype content is

$$\beta_\mathbf{x} = \mathbf{P}_\mathbf{x}^{-1}\mathrm{cov}[\mathbf{x}, w] = \mathbf{P}_\mathbf{x}^{-1}\sum_{\mathbf{k}\in\mathbb{H}} p_\mathbf{k}(\mathbf{x}_\mathbf{k} - \bar{\mathbf{x}})(w_\mathbf{k} - 1),$$

where $w_\mathbf{k} = \sum_{\mathbf{l}\in\mathbb{H}} p_\mathbf{l} w_\mathbf{kl}$ is the fitness of haplotype $\mathbf{k}$ and $w_\mathbf{kl}$ is the fitness of genotype $\mathbf{kl}$, for $\mathbf{k} \in \mathbb{H}$ and $\mathbf{l} \in \mathbb{H}$. Doing this calculation yields

$$\beta_\mathbf{x} = \begin{pmatrix} w_\mathrm{aB} - w_\mathrm{ab} \\ w_\mathrm{Ab} - w_\mathrm{ab} \\ w_\mathrm{AB} - w_\mathrm{ab} \end{pmatrix}. \tag{S56}$$

Mean absolute fitness is

$$\bar{W} = \sum_{\mathbf{k},\mathbf{l}\in\mathbb{H}} p_{\mathbf{k}} p_{\mathbf{l}} W_{\mathbf{kl}}.$$

Then, assuming that absolute fitness $W_{\mathbf{kl}}$ is independent of haplotype frequency $\bar{\mathbf{x}}$ and that fitness is haplotype-symmetric $W_{\mathbf{kl}} = W_{\mathbf{lk}}$, it can be checked that the selection regression coefficient of haplotype content is

$$\beta_{\mathbf{x}} = \frac{1}{2}\frac{1}{\bar{W}}\frac{\mathrm{d}\bar{W}}{\mathrm{d}\bar{\mathbf{x}}}. \tag{S57}$$

Numerical solution of this implementation yields the same results as in the implementation in example 4 (Fig. S2).

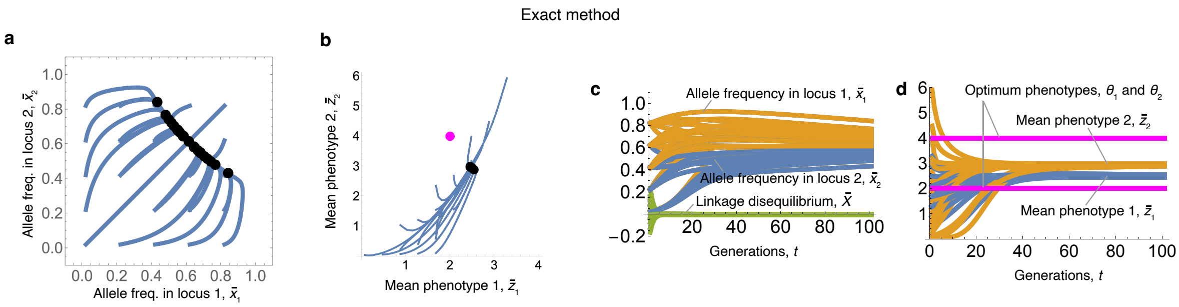


Figure S2: Example 4 continued: The same results are obtained by tracking haplotype frequencies rather than allele frequencies and linkage disequilbrium. Compare with Fig. 6a-d. Relative to that figure, there are some differences in the trajectories due to the different initial conditions arising from the different implementation.

#### S9.4 Example 5: evo-devo dynamics of one phenotype influenced by one biallelic locus under explicit development, without weakening of selection with age

A basic aspect of development is that it takes time, so individuals may not have enough time to develop optimal phenotypes even if all else is in place. These and other features are illustrated here, where we consider an example with explicit development for a single phenotype influenced by a single biallelic locus. Specifically, the scenario is the same as in examples 2-3, but where the phenotype is constructed via developmental dynamics rather than being given by a genotype-phenotype map.

##### Exact method

We first use the exact method. As in examples 2-3, $x_k = x_k^{(1\mathrm{A})}$ is the gene content for allele A in haplotype $k \in \{\mathrm{a},\mathrm{A}\}$. Let the phenotype of an individual of age $a+1 \geq 2$ and genotype $kl$ be

$$z_{a+1,kl} = g(z_{akl}, x_k, x_l) = [1 + y(x_k, x_l)]z_{akl} \tag{S58a}$$
$$= [1 + y(x_k, x_l)]^a z_1, \tag{S58b}$$

with constant initial condition $z_{1kl} = z_1$ and, following Eq. 84, growth rate

$$y(x_k, x_l) = c(1+d)(x_k + x_l) - 2cdx_kx_l. \tag{S58c}$$

It is straightforward to remove the assumption of a constant initial condition provided one specifies the dependence of $z_1$ on haplotype content and other factors. Let the absolute fitness of genotype $kl$ depend on the phenotype at the last age only; specifically, $W_{kl} = \exp[-(z_{N_\mathrm{a}kl} - \theta)^2/\sigma^2]$. One might expect that because fitness depends on the phenotype at the last age only and there is no age structure, the consideration explicit development has no evolutionary effect. However, as we show below, this is incorrect.

As the phenotype is influenced by a single biallelic locus, change in allele frequency is still given by $\Delta\bar{x} = T_x\beta_x$, where $T_x = \frac{1}{2}\bar{x}(1-\bar{x})$ and $\beta_x = w_\mathrm{A} - w_\mathrm{a}$ as derived in example 3. The difference here is in the developmental process, which alters the form of $w_\mathrm{A} - w_\mathrm{a}$ as well as the admissible evolutionary path that the mean phenotype and phenotypic variance can take. Although it is not necessary for the method, the developmental dynamics (S58a) are simple enough that we can solve them to obtain the analytical expression of the genotype-phenotype map giving the mean phenotype at the final age:

$$\begin{aligned}\bar{z}_{N_\mathrm{a}} &= \sum_{k,l\in\mathbb{H}} p_k p_l z_{N_\mathrm{a}kl} \\ &= z_1\left\{(1-\bar{x})^2 + 2^{N_\mathrm{a}-1}\left[1+c(1+d)\right]\bar{x}(1-\bar{x}) + (1+2c)^{N_\mathrm{a}-1}\bar{x}^2\right\}.\end{aligned} \tag{S59}$$

The fact that this expression for the mean phenotype is different from that in our example with implicit development (Eq. 85) immediately suggests that the evolutionary dynamics will be different even if selection (specifically, the fitness function) is the same. Similarly, the expression for the phenotype variance is more complicated than found in the implicit development example, so I do not write it here, but is also a function of allele frequency, dominance coefficient, developmentally initial phenotype, and final age.

Numerical solutions of Eq. (101a) subject to the developmental constraint (S59) are in Fig. S3a-o. The mean phenotype and phenotype variance evolve to different levels relative to the implicit development example 3 (Fig. 3a-o). In particular, the mean phenotype is less able to approach a high optimum as there may not be enough developmental time to develop such a phenotype (Figs. 3b and S3b), no disruptive selection emerges with overdominance (Figs. 3l and S3l), and phenotypic variation transiently increases but eventually vanishes with stabilising selection (Figs. 3h and S3h).

#### Approximated method

We now use the approximated method. As transmission of haplotype content is here unbiased, allele frequency change is to first order

$$\Delta\bar{x} \approx T_x \left.\frac{\mathrm{d}w}{\mathrm{d}x_k}\right|_{x_k=x_l=\bar{x}}, \tag{S60}$$

where the total selection gradient of the haplotype is

$$\left.\frac{\mathrm{d}w}{\mathrm{d}x_k}\right|_{x_k=x_l=\bar{x}} = \left.\left(\frac{\mathrm{d}z_{N_\mathrm{a}kl}}{\mathrm{d}x_k}\frac{\partial w}{\partial z_{N_\mathrm{a}kl}}\right)\right|_{x_k=x_l=\bar{x}}$$

given that fitness depends on the phenotype at the last age only.

From Eq. (71), the total haplotypic effect on the phenotype at age $a$ is

$$\left.\frac{\mathrm{d}z_{akl}}{\mathrm{d}x_k}\right|_{x_k=x_l=\bar{x}} = \left.\frac{\partial\mathbf{z}_{kl}^\intercal}{\partial x_k}\frac{\mathrm{d}z_{akl}}{\mathrm{d}\mathbf{z}_{kl}}\right|_{x_k=x_l=\bar{x}} \tag{S61}$$

where, from Eq. (72), the direct haplotypic effect on the phenotype at all ages is

$$\frac{\partial\mathbf{z}_{kl}^\intercal}{\partial x_k} = \begin{pmatrix}0 & \frac{\partial z_{2kl}}{\partial x_k} & \cdots & \frac{\partial z_{N_\mathrm{a}kl}}{\partial x_k}\end{pmatrix}$$

and the total effect of the phenotype at any age on the phenotype at age $a$ is

$$\frac{\mathrm{d}z_{akl}}{\mathrm{d}\mathbf{z}_{kl}} = \begin{pmatrix}\frac{\mathrm{d}z_{akl}}{\mathrm{d}z_{1kl}} & \cdots & \frac{\mathrm{d}z_{akl}}{\mathrm{d}z_{N_\mathrm{a}kl}}\end{pmatrix}^\intercal.$$

Thus, taking the matrix multiplication indicated in (S61), the total haplotypic effect on the phenotype at age $a$ is

$$\left.\frac{\mathrm{d}z_{akl}}{\mathrm{d}x_k}\right|_{x_k=x_l=\bar{x}} = \left.\sum_{j=1}^{N_\mathrm{a}}\frac{\partial z_{jkl}}{\partial x_k}\frac{\mathrm{d}z_{akl}}{\mathrm{d}z_{jkl}}\right|_{x_k=x_l=\bar{x}}.$$

Since the direct effect of the haplotype on the phenotype at the first age is zero because of our assumption of constant phenotype at the initial age, the first term in the sum is zero yielding

$$\left.\frac{\mathrm{d}z_{akl}}{\mathrm{d}x_k}\right|_{x_k=x_l=\bar{x}} = \left.\sum_{j=2}^{N_\mathrm{a}}\frac{\partial z_{jkl}}{\partial x_k}\frac{\mathrm{d}z_{akl}}{\mathrm{d}z_{jkl}}\right|_{x_k=x_l=\bar{x}}.$$

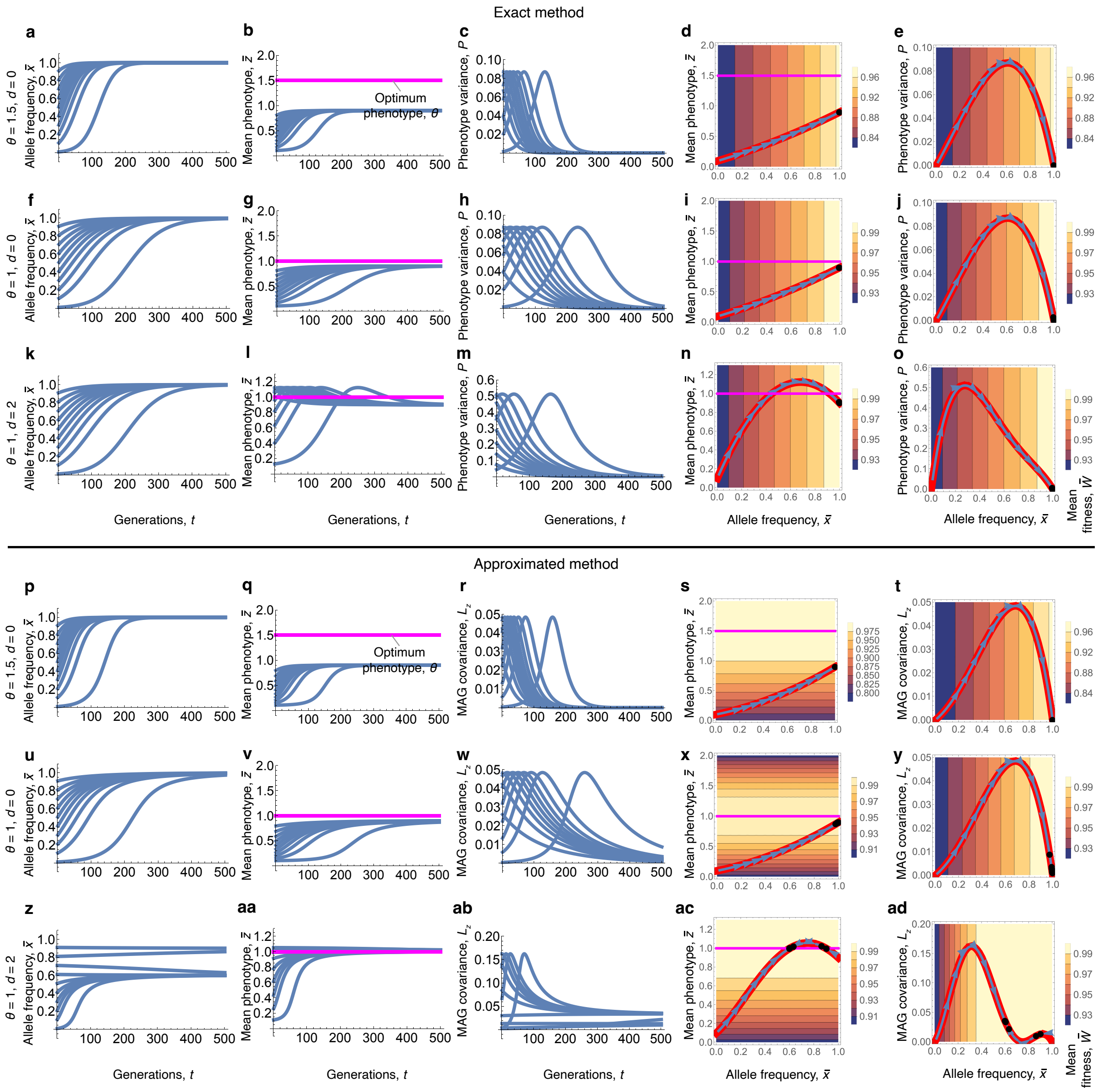


Figure S3: Example 5: one phenotype and one biallelic locus under explicit development. **a-o,** Exact method. As in Fig. 3a-o but with developmental map (S58a) and fitness $W_{kl} = \exp[-(z_{N_\text{a}kl} - \theta)^2/\sigma^2]$. Parameter values: $N_\text{a} = 3$, $z_1 = 0.1$, $\sigma^2 = 10$, and $c = 1$. In **d,i,n,** the red lines are the mean phenotype $\bar{z}$ as a function allele frequency $\bar{x}$. In **e,j,o,** the red lines are the phenotypic variance $P$ as a function of allele frequency $\bar{x}$. **p-ad,** Approximated method. As in panels a-o but using the first-order approximation. The approximation is fairly accurate, although less so with overdominance.

From Eq. (12) of González-Forero (2024), the total effects of the phenotype on itself are

$$\frac{\mathrm{d}z_{akl}}{\mathrm{d}z_{jkl}} = \begin{cases} \displaystyle\prod_{i=j}^{a-1} \frac{\partial z_{i+1,kl}}{\partial z_{ikl}} & \text{if } j < a \\ 1 & \text{if } j = a \\ 0 & \text{if } j > a, \end{cases}$$

where the latter case equals zero because changing the phenotype at a later age does not alter the phenotype at a previous age, and the middle case equals one because the derivative of the phenotype with respect to itself at the same age is one. These two latter cases imply that the total effects of the haplotype on the phenotype reduces further to

$$\begin{aligned}\left.\frac{\mathrm{d}z_{akl}}{\mathrm{d}x_k}\right|_{x_k=x_l=\bar{x}} &= \sum_{j=2}^{a} \left.\frac{\partial z_{jkl}}{\partial x_k}\frac{\mathrm{d}z_{akl}}{\mathrm{d}z_{jkl}}\right|_{x_k=x_l=\bar{x}} \\ &= \sum_{j=2}^{a-1} \left.\frac{\partial z_{jkl}}{\partial x_k}\frac{\mathrm{d}z_{akl}}{\mathrm{d}z_{jkl}}\right|_{x_k=x_l=\bar{x}} + \left.\frac{\partial z_{akl}}{\partial x_k}\right|_{x_k=x_l=\bar{x}}.\end{aligned}$$

Now, using the developmental map (S58a), the direct haplotypic effect for $a+1 \geq 2$ is

$$\begin{aligned}\left.\frac{\partial z_{a+1,kl}}{\partial x_k}\right|_{x_k=x_l=\bar{x}} &= Dz_{akl}|_{x_k=x_l=\bar{x}} \\ &= D(1+y^\circ)^{a-1}z_1,\end{aligned}$$

where $D = c(1+d) - 2cd\bar{x}$ and $y^\circ = y(\bar{x},\bar{x}) = 2c(1+d)\bar{x} - 2cd\bar{x}^2$. The direct phenotypic effect for $a+1 \geq 2$ is

$$\left.\frac{\partial z_{a+1,kl}}{\partial z_{akl}}\right|_{x_k=x_l=\bar{x}} = 1+y^\circ.$$

Hence, the total haplotypic effect for $a \leq 2$ is

$$\left.\frac{\mathrm{d}z_{akl}}{\mathrm{d}x_k}\right|_{x_k=x_l=\bar{x}} = \begin{cases} 0 & \text{if } a=1 \\ Dz_1 & \text{if } a=2, \end{cases}$$

and for $a > 2$ it is

$$\begin{aligned}\left.\frac{\mathrm{d}z_{akl}}{\mathrm{d}x_k}\right|_{x_k=x_l=\bar{x}} &= \sum_{j=2}^{a-1} \frac{\partial z_{jkl}}{\partial x_k} \prod_{i=j}^{a-1} \left.\frac{\partial z_{i+1,kl}}{\partial z_{ikl}}\right|_{x_k=x_l=\bar{x}} \\ &\quad + D(1+y^\circ)^{a-2}z_1 \\ &= \sum_{j=2}^{a-1} D(1+y^\circ)^{j-2}z_1 \prod_{i=j}^{a-1}(1+y^\circ) \\ &\quad + D(1+y^\circ)^{a-2}z_1 \\ &= (a-1)Dz_1(1+y^\circ)^{a-2}.\end{aligned}$$

Hence, allele frequency equilibria can occur either because an allele is fixed (so $T_x = 0$), the optimum phenotype is reached (so the selection gradient on the phenotype is zero), or the total haplotypic effect vanishes ($D = 0$), which requires overdominance if $d > 0$. Specifically, $D = 0$ occurs when $\bar{x}^* = (1+d)/(2d)$, which is $\bar{x}^* = 0.75$ if $d = 2$.

To analyse the role of constraints on phenotypic evolution, we can calculate the MAG covariance of the phenotype at the final age, which from (57) reduces here to

$$L_{z_{N_a}} = 2T_x \left.\left(\frac{\mathrm{d}z_{N_a kl}}{\mathrm{d}x_k}\right)^2\right|_{x_k=x_l=\bar{x}}, \tag{S62a}$$

as in eq. (112). Similarly, letting $\mathbf{m}_{kl} = (z_{N_a kl}, x_k)^\intercal$, the MAG covariance matrix of the phenotype at the final age and the haplotype is given by

$$\begin{aligned}\mathbf{L_m} &\equiv \mathrm{cov}[\mathbf{b}_{\hat{\mathbf{m}}_k}, \mathbf{b}_{\mathbf{m}_k}] \\ &= \left.\begin{pmatrix} 2\left(\frac{\mathrm{d}z_{N_a kl}}{\mathrm{d}x_k}\right)^2 T_x & 2\frac{\mathrm{d}z_{N_a kl}}{\mathrm{d}x_k}T_x \\ T_x\frac{\mathrm{d}z_{N_a kl}}{\mathrm{d}x_k} & T_x \end{pmatrix}\right|_{x_k=x_l=\bar{x}},\end{aligned} \tag{S62b}$$

as in eq. (115b).

As in example 3, let the absolute fitness be $W(z_{N_a kl}) = \exp[-(z_{N_a kl} - \theta)^2/\sigma^2]$. Numerical solutions of this example using the approximated method (S60) are similar to those using the exact method (Fig. S3p-ad). When $d = 0$, the allele fixes, but the phenotype optimum is not achieved because there is not enough developmental time for the phenotype to grow ($N_a = 3$ is too small; Fig. S3p-y). When $d = 2$, there is disruptive selection on the genes (but not on the phenotype), such that different internal allele frequency equilibria are achieved depending on the initial allele frequencies, and at such equilibria the mean phenotype is at the optimum phenotype (Fig. S3a-ad).

#### S9.5 Example 6: evo-devo dynamics of one phenotype influenced by one biallelic locus under explicit development, with weakening of selection with age

Another basic aspect of development taking time is that as individuals' age advances, selection weakens due to individuals dying and having fewer reproductive opportunities left (Medawar, 1952; Hamilton, 1966). Thus, for our last example, we modify the previous one to allow for the weakening of selection with age, again under explicit development of a single phenotype influenced by a single biallelic locus. To do this, we still consider non-overlapping generations, but use a fitness function that weighs survival and fertility with decreasing values, where the weights have been previously derived for age-structured demographies under a constant population size (Hamilton, 1966; Caswell, 1978; Caswell and Shyu, 2017). In particular, rather than selection occurring at a single age as in the previous example, in this example selection occurs at every age. Results show that the evolution is substantially slower relative to the previous example without age structure and selection at one age only.

##### Exact method

As there is still a single biallelic locus as in examples 3 and 5, allele frequency change continues to be given by $\Delta\bar{x} = T_x\beta_x$, with $T_x = \frac{1}{2}\bar{x}(1-\bar{x})$ and $\beta_x = w_{\mathrm{A}} - w_{\mathrm{a}}$. The developmental map is also the same as in example 5. The difference here is in fitness, where we consider weakening of selection with age.

To do this, let the absolute fitness of genotype $kl$ be

$$W_{kl} = \frac{1}{\tau}\sum_{a=1}^{N_{\mathrm{a}}}(\phi_a f_{akl} + \pi_a s_{akl}), \tag{S63}$$

where for an individual of age $a$ and genotype $kl$, their fertility is $f_{akl}$ and their probability of surviving from age $a$ to $a+1$ is $s_{akl}$ (this fitness function is a modification of eq. 5 of González-Forero, 2024 to allow for diploid genotypes of any frequency, which is in turn a fitness function previously derived for age-structured populations at carrying capacity; Hamilton, 1966; Caswell, 1978; Caswell and Shyu, 2017). The force of selection on fertility at age $a$ is

$$\phi_a = \sum_{k,l\in\mathbb{H}} p_k p_l \ell_{akl}, \tag{S64a}$$

the force of selection on survival at age $a$ is

$$\pi_a = \sum_{k,l\in\mathbb{H}} p_k p_l \frac{1}{s_{akl}} \sum_{j=a+1}^{N_{\mathrm{a}}} \ell_{jkl} f_{jkl}, \tag{S64b}$$

generation time is

$$\tau = \sum_{k,l\in\mathbb{H}} p_k p_l \sum_{a=1}^{N_{\mathrm{a}}} a\ell_{akl} f_{akl}, \tag{S64c}$$

and the survivorship to age $a$ for genotype $kl$ is

$$\ell_{akl} = \prod_{j=1}^{a-1} s_{jkl}. \tag{S64d}$$

For comparability with previous examples, let fertility be

$$f_{akl} = \exp[-(z_{akl} - \theta_{\mathrm{fa}})^2/\sigma^2_{\mathrm{fa}}] \tag{S65a}$$

and survival be

$$s_{akl} = \exp[-(z_{akl} - \theta_{\mathrm{sa}})^2/\sigma^2_{\mathrm{sa}}], \tag{S65b}$$

with possibly different fertility and survival optima and selection strengths at every age. The forces of selection and generation time depend on haplotype frequency $p_k$ and so on allele frequency. Consequently, $W_{kl}$ is frequency-dependent and the selection regression coefficient $\beta_x$ is no longer a selection gradient so mean fitness may decrease.

Numerical solutions of Eq. (101a) subject to the developmental constraint (S59) under fitness (S63) are in Fig. S4a-o. Fitness valleys are crossed (Fig. S4d,i,n) due to frequency dependent selection, since $\beta_x$ is no longer a selection gradient. Results are similar to those in example 5 without weakening of selection with age, although evolution is much slower (see horizontal axes in Fig. S4a-c,f-h,k-m).

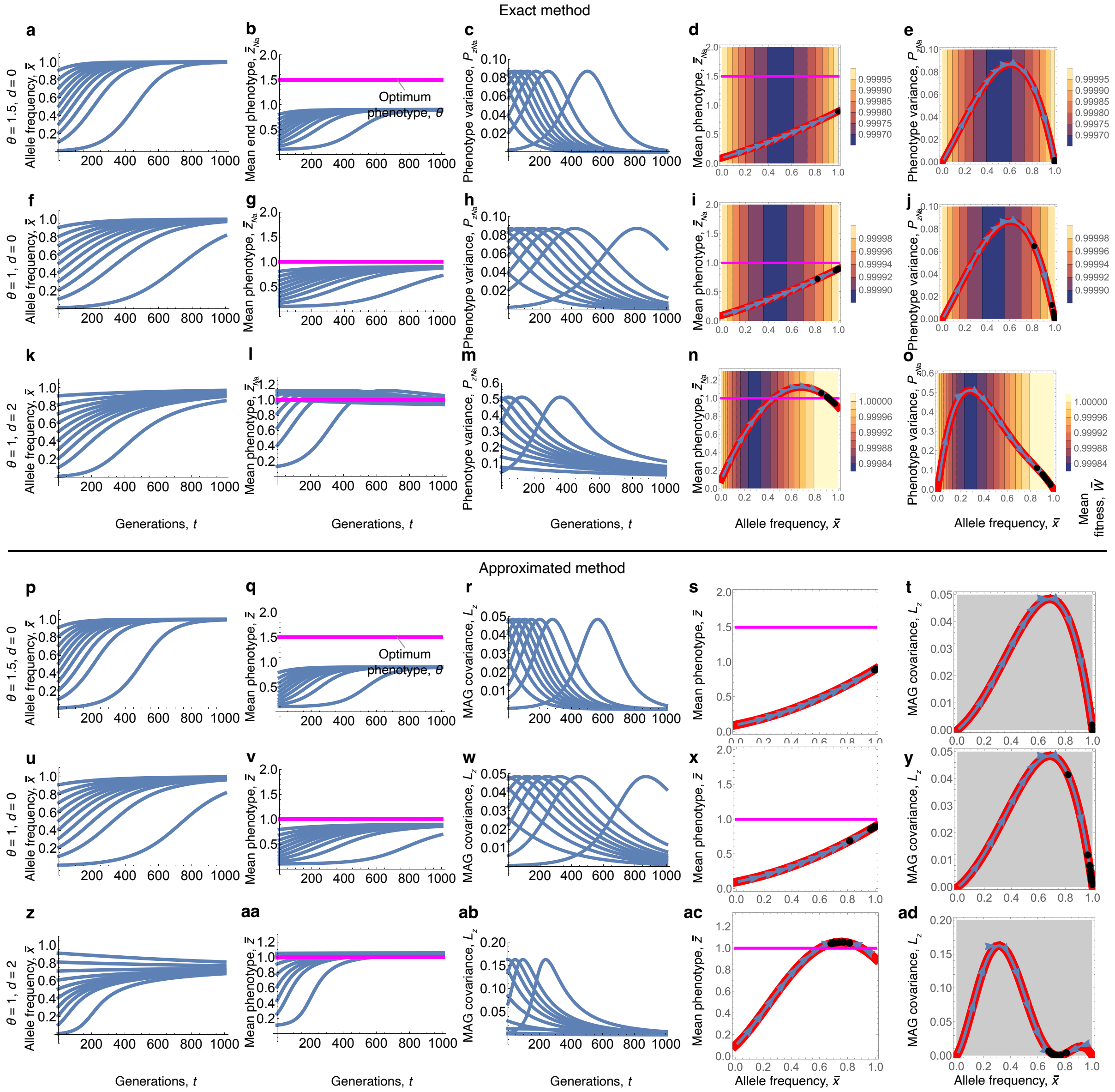


Figure S4: Example 6: one phenotype and one biallelic locus under explicit development and weakening of selection with age. **a-o,** Exact approach. As in Fig. S3 but with fitness (S63) and most plots now refer to the phenotype at the final age, $z_{N_a}$. Parameter values: $N_a = 3$, $z_1 = 0.1$, $\theta_{sa} = \theta_{fa} = \theta$, and $\sigma^2_{sa} = \sigma^2_{fa} = 10$ for all $a \in \{1, \ldots, N_a\}$. Specifically, there is selection for the same phenotypic optimum and of the same strength at every age. In **d,i,n,** the red lines are the mean phenotype $\bar{z}$ as a function allele frequency $\bar{x}$. In **e,j,o,** the red lines are the phenotype variance $P$ as a function of allele frequency $\bar{x}$. **p-ad,** Approximated approach. As in **a-o** but using the first-order approximation. The approximation is fairly accurate, although less so with overdominance. Evolution is much slower than in the example without age structure and with selection at one age only (e.g., compare the the horizontal axes in panels **a-c** with Fig. S3**a-c**).

#### Approximated method

We now use the approximated method.

As transmission of haplotype content is here unbiased, allele frequency change is to first order

$$\Delta \bar{x} \approx T_x \left. \frac{\mathrm{d}w}{\mathrm{d}x_k} \right|_{x_k = x_l = \bar{x}},$$

where the total selection gradient of the haplotype is

$$\left.\frac{\mathrm{d}w}{\mathrm{d}x_k}\right|_{x_k=x_l=\bar{x}} = \left.\left(\frac{\mathrm{d}\mathbf{z}_{kl}^{\intercal}}{\mathrm{d}x_k}\frac{\partial w}{\partial \mathbf{z}_{kl}}\right)\right|_{x_k=x_l=\bar{x}} = \left.\left(\sum_{a=1}^{N_\mathrm{a}}\frac{\mathrm{d}z_{akl}}{\mathrm{d}x_k}\frac{\partial w}{\partial z_{akl}}\right)\right|_{x_k=x_l=\bar{x}}$$

given that here fitness depends on the phenotype at all ages.

Hence, in addition to the allele frequency equilibria found in example 5, there are other equilibria possible where the mean phenotype is not at the optimum at every age but is at values that trade-off persistent negative and positive phenotypic selection at different ages with weights given by the total haplotypic effects. Other possibilities would exist if the total haplotypic effect had different signs at different ages, but this is not the case in this simple example.

Then, the selection gradient of the phenotype at the $a$-th age is

$$\left.\frac{\partial w}{\partial z_{akl}}\right|_{x_k=x_l=\bar{x}} = \frac{1}{\tau}\left.\left(\phi_a\frac{\partial f_{akl}}{\partial z_{akl}} + \pi_a\frac{\partial s_{akl}}{\partial z_{akl}}\right)\right|_{x_k=x_l=\bar{x}},$$

where the phenotypic effects on fertility and survival are

$$\left.\frac{\partial f_{akl}}{\partial z_{akl}}\right|_{x_k=x_l=\bar{x}} = 2f_a^{\circ}\frac{\theta_{\mathrm{fa}}-z_a^{\circ}}{\sigma_{\mathrm{fa}}^2}$$

$$\left.\frac{\partial s_{akl}}{\partial z_{akl}}\right|_{x_k=x_l=\bar{x}} = 2s_a^{\circ}\frac{\theta_{\mathrm{sa}}-z_a^{\circ}}{\sigma_{\mathrm{sa}}^2},$$

and where $\circ$ indicates evaluation at the mean haplotype content.

Following eq. 5 of González-Forero, 2024, let the absolute fitness of genotype $kl$ be

$$W(\mathbf{z}_{kl},\bar{\mathbf{z}}) = \frac{1}{\tau(\bar{\mathbf{z}})}\sum_{a=1}^{N_\mathrm{a}}\left[\phi_a(\bar{\mathbf{z}})f_a(z_{akl}) + \pi_a(\bar{\mathbf{z}})s_a(z_{akl})\right], \tag{S66}$$

where the lifetime phenotype of genotype $kl$ is $\mathbf{z}_{kl} = (z_{1kl}, z_{2kl}, \ldots, z_{N_\mathrm{a}kl})^{\intercal}$ and the average lifetime phenotype $\bar{\mathbf{z}} = (\bar{z}_1, \bar{z}_2, \ldots, \bar{z}_{N_\mathrm{a}})^{\intercal}$ is to first order given by the solution of (S58a) evaluated at $x_k = x_l = \bar{x}$:

$$\bar{z}_{a+1} \approx g(\bar{z}_a, \bar{x}, \bar{x}), \tag{S67}$$

with initial condition $\bar{z}_1 = z_1$. The force of selection on fertility at age $a$ is

$$\phi_a(\bar{\mathbf{z}}) = \ell_a(\bar{\mathbf{z}}), \tag{S68a}$$

the force of selection on survival at age $a$ is

$$\pi_a(\bar{\mathbf{z}}) = \frac{1}{s_a(\bar{z}_a)}\sum_{j=a+1}^{N_\mathrm{a}}\ell_j(\bar{\mathbf{z}})f_j(\bar{z}_j), \tag{S68b}$$

generation time is

$$\tau(\bar{\mathbf{z}}) = \sum_{a=1}^{N_\mathrm{a}} a\ell_a(\bar{\mathbf{z}})f_a(\bar{z}_a), \tag{S68c}$$

and the survivorship to age $a$ for genotype $kl$ is

$$\ell_a(\mathbf{z}_{kl}) = \prod_{j=1}^{a-1}s_j(z_{jkl}). \tag{S68d}$$

We let fertility be

$$f_a(z_{akl}) = \exp[-(z_{akl}-\theta_{\mathrm{fa}})^2/\sigma_{\mathrm{fa}}^2] \tag{S69a}$$

and survival be

$$s_a(z_{akl}) = \exp[-(z_{akl}-\theta_{\mathrm{sa}})^2/\sigma_{\mathrm{sa}}^2], \tag{S69b}$$

To first order, mean absolute fitness is $W(\bar{\mathbf{z}}, \bar{\mathbf{z}}) = 1$, so absolute and relative fitness are the same to first order. The MAG covariances are still given by (S62).

Numerical solutions of this approximated approach for this model are in Fig. S4p-ad. The approximation mostly recovers the qualitative results of the exact approach. Mean fitness decrease is not seen in this approximation because mean fitness is always one to first order in this example (i.e., here $\bar{W} \approx W(\bar{\mathbf{z}}, \bar{\mathbf{z}}) = 1$). However, the trajectories are similar.

### References


Caswell, H. (1978). A general formula for the sensitivity of population growth rate to changes in life history parameters. *Theor. Popul. Biol.*, **14**, 215–230.

Caswell, H. and Shyu, E. (2017). *Senescence, selection gradients and mortality*, chapter 4, pages 56–82. Cambridge Univ. Press, Cambridge, UK.

Frank, S.A. (1997). The Price equation, Fisher's fundamental theorem, kin selection, and causal analysis. *Evolution*, **51**(6), 1712–1729.

González-Forero, M. (2024). A mathematical framework for evo-devo dynamics. *Theor. Popul. Biol.*, **155**, 24–50.

Hamilton, W.D. (1966). The moulding of senescence by natural selection. *J. Theor. Biol.*, **12**, 12–45.

Lande, R. (1979). Quantitative genetic analysis of multivariate evolution applied to brain: body size allometry. *Evolution*, **34**, 402–416.

Lande, R. and Arnold, S.J. (1983). The measurement of selection on correlated characters. *Evolution*, **37**, 1210–1226.

Lewontin, R.C. and Kojima, K. (1960). The evolutionary dynamics of complex polymorphisms. *Evolution*, **14**, 458–472.

Medawar, P.B. (1952). *An unsolved problem of biology*. H. K. Lewis, London, UK.

Nagylaki, T. (1992). *Theoretical Population Genetics*. Springer-Verlag, Berlin, Germany.

Rice, S.H. (2004). *Evolutionary Theory*. Sinauer, Sunderland, MA, USA.
