## Supplementary material for "A mathematical synthesis of genetics, development, and evolution": Computer code

### Computer Code for: A mathematical synthesis of genetics, development, and evolution

Mauricio González-Forero
Konrad Lorenz Institute for Evolution and Cognition Research, Klosterneuburg
A-3400, Austria

This file contains the computer code used to do routine derivations and generate the figures in the main text and supplementary information. This code was prepared in Mathematica 13.3.1.0 and it is made available under a Creative Commons Attribution licence (CC BY NC).

### ComputerCode Contents

©1988–2008 Wolfram Research, Inc. All rights reserved.

#### Example 1: Quantitative genetics model

In[ ]:=
```
Clear["Global`*"]
```

In[ ]:=
```
(*This is absolute fitness for phenotype z1,
with optimum θ1 and selection weakness σ2*)
W[z1_, θ1_, σ2_] := Exp[-(z1 - θ1)^2/σ2]

(*This is the normal distribution for
 phenotype z1 with mean z1b and covariance matrix Pd*)
Pfun[z1_, z1b_, Pd_] := (2 π)^(-1/2) Det[Pd]^(-1/2)
   Exp[-1/2 Transpose[({{z1}} - {{z1b}})].Inverse[Pd].({{z1}} - {{z1b}})];

(*This extracts the probability
 density for z1 with mean z1b and variance P11*)
P[z1_, z1b_, P11_] := Simplify[Pfun[z1, z1b, {{P11}}][[1, 1]]]
```

In[ ]:=
```
(*This calculates mean fitness*)
Wbars[z1b_, θ1_, σ2_, P11_] = Assuming[P11 > 0 && σ2 > 0,
   Integrate[W[z1, θ1, σ2] × P[z1, z1b, P11], {z1, -∞, ∞}]];

Wbar[z1b_, θ1_, σ2_, P11_] :=
 Assuming[P11 > 0 && σ2 > 0, Simplify[Wbars[z1b, θ1, σ2, P11]]]
```

In[ ]:=
```
Wbar[z1b, θ1, σ2, P11]
```

Out[ ]=

$$\frac{e^{-\frac{(z1b-\theta1)^2}{2\,P11+\sigma2}}}{\sqrt{1+\frac{2\,P11}{\sigma2}}}$$

In[ ]:=
```
(*This is relative fitness*)
Simplify[W[Z1, θ1, σ2] / Wbar[Z1b, θ1, σ2, P11]]
```

Out[ ]=

$$e^{-\frac{(Z1-\theta1)^2}{\sigma2}+\frac{(Z1b-\theta1)^2}{2\,P11+\sigma2}}\sqrt{1+\frac{2\,P11}{\sigma2}}$$

In[ ]:=
```
(*This is the average individual selection gradient*)
B[z1b_, θ1_, σ2_, P11_] =
  Simplify[Assuming[1/(2 P11) + 1/σ2 > 0, Integrate[D[W[z1, θ1, σ2], z1] ×
        P[z1, z1b, P11], {z1, -∞, ∞}]] / Wbar[z1b, θ1, σ2, P11]];
```

In[ ]:=
```
B[z1b, θ1, σ2, P11]
```

Out[ ]=

$$-\frac{2\,(z1b-\theta1)}{2\,P11+\sigma2}$$

```
In[ ]:= (*This is for the first entry of the Hessian*)
        H11[z1b_, θ1_, σ2_, P11_] =
          Simplify[Assuming[1/(2 P11) + 1/σ2 > 0, Integrate[Simplify[D[W[z1, θ1, σ2], z1, z1]] ×
               P[z1, z1b, P11], {z1, -∞, ∞}]] / Wbar[z1b, θ1, σ2, P11]];

In[ ]:= H11[z1b, θ1, σ2, P11]
Out[ ]=
```

$$-\frac{2\left(2\ P11-2\ (z1b-\theta1)^{2}+\sigma2\right)}{(2\ P11+\sigma2)^{2}}$$

```
In[ ]:= (*This is the dynamic equation of ΔP*)
         DGToSolve[z1b_, θ1_, σ2_, P11_] :=
          Simplify[P11 (H11[z1b, θ1, σ2, P11] - B[z1b, θ1, σ2, P11] × B[z1b, θ1, σ2, P11]) P11]

In[ ]:= (*This calculates the numerical solutions of the evolutionary dynamics of
          the mean phenotype and phenotype variance for many initial conditions*)

        θ1 = .75;
        σ2 = .1;

        tend = 500;

        z1init = {0.01, 1, .1}; (*To sample initial conditions for z bar*)
        P11init = {0.001, .02, .005} ; (*To sample initial conditions for P*)

        InitialConditions =
          Table[{z1b[0, i, j] = i, P11[0, i, j] = j}, {i, z1init[[1]], z1init[[2]], z1init[[3]]},
           {j, P11init[[1]], P11init[[2]], P11init[[3]]}];

        Table[Table[{z1b[t + 1, i, j] =
              z1b[t, i, j] + (P11[t, i, j] × B[z1b[t, i, j], θ1, σ2, P11[t, i, j]]),
             P11[t + 1, i, j] = P11[t, i, j] + DGToSolve[z1b[t, i, j], θ1, σ2, P11[t, i, j]]},
           {t, 0, tend}], {i, z1init[[1]], z1init[[2]], z1init[[3]]},
          {j, P11init[[1]], P11init[[2]], P11init[[3]]}];
```

In[ ]:=
```
(*This plots the numerical solutions over time*)

lThickness = 0.02; (*Line thickness*)
Show[Table[ListLinePlot[{Table[z1b[t, i, j], {t, 0, tend}]},
    PlotRange → {0, 1}, TicksStyle → Large, PlotStyle → Thickness[lThickness]],
   {i, z1init[[1]], z1init[[2]], z1init[[3]]}, {j, P11init[[1]], P11init[[2]], P11init[[3]]}],
  ListLinePlot[Table[{t, θ1}, {t, 0, tend}],
   PlotStyle → {Thickness[lThickness], Magenta}]]
Export[StringJoin[ToString[NotebookDirectory[]], "Fig.-1a", ".pdf"], %];
Show[Table[ListLinePlot[{Table[P11[t, i, j], {t, 0, tend}]},
    PlotRange → {0, .02}, TicksStyle → Large, PlotStyle → Thickness[lThickness]],
   {i, z1init[[1]], z1init[[2]], z1init[[3]]}, {j, P11init[[1]], P11init[[2]], P11init[[3]]}]]
Export[StringJoin[ToString[NotebookDirectory[]], "Fig.-1b", ".pdf"], %];
```

Out[ ]=

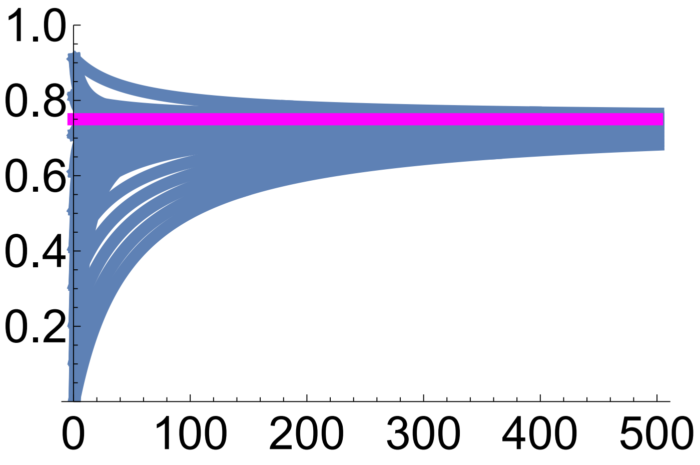

Out[ ]=

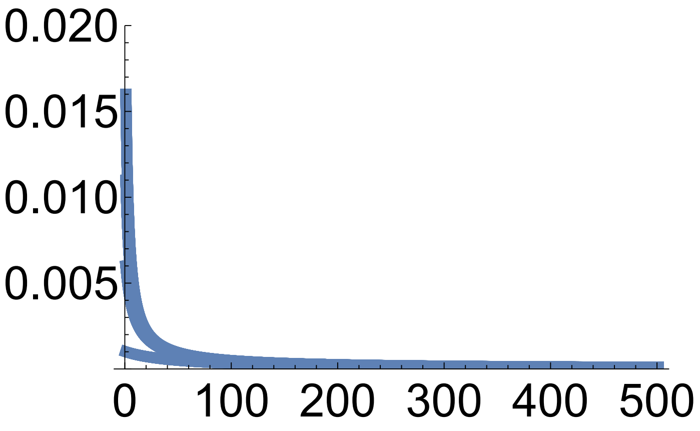

In[ ]:=
```
(*This plots the numerical solutions in the
 space of mean phenotype and phenotype variance*)

lThickness = 0.02; (*Line thickness*)

p0 = Table[ListLinePlot[Table[{z1b[t, i, j], P11[t, i, j]}, {t, 0, tend}],
    PlotRange → {{0, 1}, {0, 0.02}}, PlotStyle → {Thickness[lThickness]}], {i,
    z1init[[1]], z1init[[2]], z1init[[3]]}, {j, P11init[[1]], P11init[[2]], P11init[[3]]}];

plot = Show[ContourPlot[Wbar[z1b, θ1, σ2, 0.01], {z1b, 0, 1}, {P11, 0, .02},
   FrameStyle → Large, PlotLegends → Automatic, LabelStyle → Large],
  p0 /. Line[x_] :→ {Arrowheads[Table[{.1, .2}, {3}]], Arrow[x]},
  ListLinePlot[Table[{θ1, P}, {P, 0, 0.02, 0.01}],
   PlotStyle → {Thickness[lThickness], Magenta}]]

plot = Rasterize[plot, "Image", ImageResolution → 300];

Export[StringJoin[ToString[NotebookDirectory[]], "Fig.-1c", ".pdf"], plot];
```

Out[ ]=

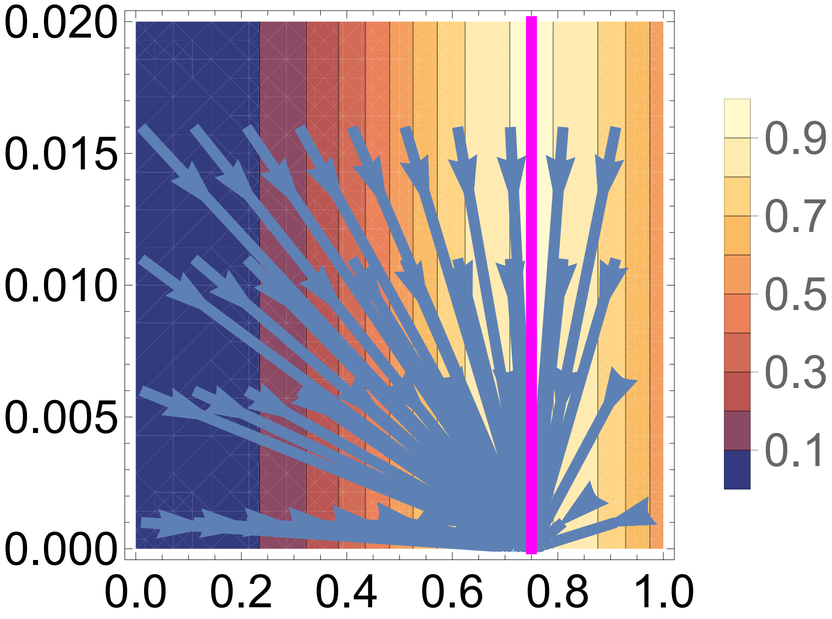

In[ ]:=

```
(*This plots the fitness landscape at phenotype variance P=0.01*)

Show[Plot3D[Wbar[Z1b, θ1, σ2, 0.01], {xb, 0, 1}, {Z1b, 0, 1},
  AxesLabel → None, AxesStyle → Large, PlotStyle → Yellow,
  PlotRange → {0, 1.1}, ViewPoint → {1.3, -1.3, 1.5}]]
Export[StringJoin[ToString[NotebookDirectory[]], "Fig.-1d", ".pdf"], %];
```

Out[ ]=

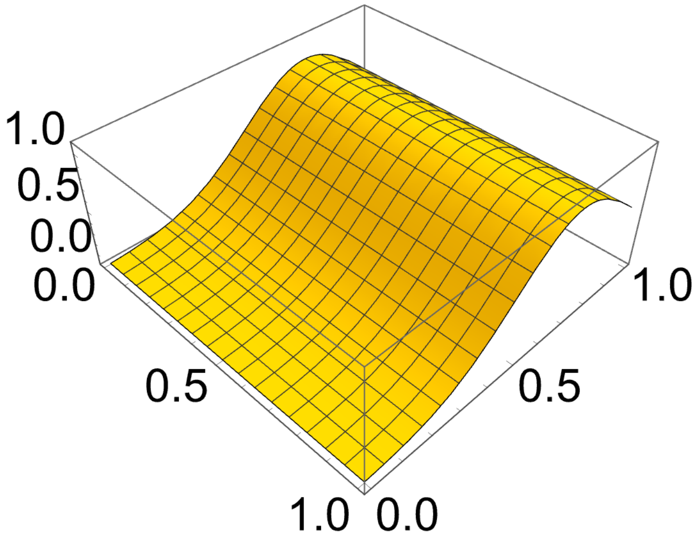

#### Example 2: Coefficients of generator equations under random mating

In[ ]:=
```
Clear["Global`*"]
```

In[ ]:=
```
(*This lists gene content, haplotype frequencies,
genotype frequencies, and phenotypes*)

xa = 0; xA = 1;
pa = (1 - xb); pA = xb;
paa = pa^2; pAa = 2 pa pA; pAA = pA^2;

zaa = 0;
zAa = c (1 + d);
zAA = 2 c;

(*This is Px=var[x], the allelic variance*)
Px[xvar_] = Simplify[Sum[p[i] (x[i] - xb)^2, {i, 1, 2}] /. {p[1] → pa, p[2] → pA} /.
      {x[1] → xa, x[2] → xA}] /. {xb → xvar};
Px[xb]

(*This is the mean phenotype*)
zb = Simplify[Sum[p[i] × p[j] × z[i, j], {i, 1, 2}, {j, 1, 2}] /.
    {p[1] → pa, p[2] → pA, z[1, 1] → zaa, z[1, 2] → zAa, z[2, 1] → zAa, z[2, 2] → zAA}]

(*This is to plot the mean phenotype as a function of allele frequency*)
zbfun[xvar_, dvar_] = zb /. {xb → xvar, d → dvar} ;
```

Out[ ]=

$-((-1 + xb)\, xb)$

Out[ ]=

$2\, c\, xb\, (1 + d - d\, xb)$

In[ ]:=
```
zb /. d → 0
```

Out[ ]=

$2\, c\, xb$

```
(*This is genotype content of genotype kl*)
y[k_, l_] :=
 If[k == 1 && l == 1, 0, If[(k == 1 && l == 2) || (k == 2 && l == 1), 1, If[k == 2 && l == 2, 2]]]
(*y[k_,l_]:=If[k==1&&l==1,0,If[(k==1&&l==2)||(k==2&&l==1),1,If[k==2&&l==2,2]]]*)

(*This is the average genotype content*)
yb =
  Simplify[Sum[p[i] × p[j] × y[i, j], {i, 1, 2}, {j, 1, 2}] /. {p[1] → pa, p[2] → pA}];

(*This is xi and D*)
xi[xvar_, dvar_] := FullSimplify[Sum[p[i] × p[j] × z[i, j], {i, 1, 2}, {j, 1, 2}] /.
     {p[1] → pa, p[2] → pA} /. {z[1, 1] → zaa, z[1, 2] → zAa,
     z[2, 1] → zAa, z[2, 2] → zAA}] /. {xb → xvar, d → dvar}

Dd[xvar_, dvar_] :=
 FullSimplify[Sum[p[i] × p[j] (z[i, j] - zb) (y[i, j] - yb), {i, 1, 2}, {j, 1, 2}] /
       Sum[p[i] × p[j] (y[i, j] - yb)^2, {i, 1, 2}, {j, 1, 2}] /.
      {p[1] → pa, p[2] → pA} /. {z[1, 1] → zaa, z[1, 2] → zAa, z[2, 1] → zAa,
      z[2, 2] → zAA} /. {x[1] → xa, x[2] → xA}] /. {xb → xvar, d → dvar}

yb
Out[ ]=
2 xb

In[ ]:= xi[xb, d] == zb
Simplify[zb == 2 c (1 + d) xb - 2 c d xb^2]
Simplify[Dd[xb, d] == c (1 + d) - 2 c d xb]
xi[xb, 0]
Dd[xb, 0]
Out[ ]=
True
Out[ ]=
True
Out[ ]=
True
Out[ ]=
2 c xb
Out[ ]=
c

In[ ]:= (*This checks that D = 1/2 dzb/dxb*)
Simplify[Dd[xb, d] == 1/2 D[zb, xb]]
Out[ ]=
True
```

In[ ]:=
```
(*This is P*)
P[xvar_, dvar_] :=
 FullSimplify[Sum[p[i] × p[j] (z[i, j] - zb)^2, {i, 1, 2}, {j, 1, 2}] /.
     {p[1] → pa, p[2] → pA} /. {z[1, 1] → zaa, z[1, 2] → zAa,
     z[2, 1] → zAa, z[2, 2] → zAA}] /. {xb → xvar, d → dvar}

Simplify[P[xb, d] == 2 Px[xb] Dd[xb, d]^2 + (2 Px[xb] c d)^2]
Simplify[P[xb, 0]]
```

Out[ ]=
True

Out[ ]=
$-2\, c^2\, (-1 + xb)\, xb$

In[ ]:=
```
(*This enters the mean phenotype in offspring of a given parental phenotype*)
```

$$zaap = paa\ zaa + pAa\left(\frac{1}{2}\ zAa + \frac{1}{2}\ zaa\right) + pAA\ zAa;$$

$$zAap = paa\left(\frac{1}{2}\ zAa + \frac{1}{2}\ zaa\right) + pAa\left(\frac{1}{4}\ zaa + \frac{1}{2}\ zAa + \frac{1}{4}\ zAA\right) + pAA\left(\frac{1}{2}\ zAa + \frac{1}{2}\ zAA\right);$$

$$zAAp = paa\ zAa + pAa\left(\frac{1}{2}\ zAa + \frac{1}{2}\ zAA\right) + pAA\ zAA;$$

```
zp[k_, l_] := If[k == 1 && l == 1, zaap,
  If[(k == 1 && l == 2) || (k == 2 && l == 1), zAap, If[k == 2 && l == 2, zAAp]]]
```

In[ ]:=
```
Simplify[zaap]
Simplify[zAap]
Simplify[zAAp]
Simplify[zAAp == c (1 + d + xb (1 - d))]
```

Out[ ]=
$c\,(1 + d)\, xb$

Out[ ]=
$\frac{1}{2}\, c\,(1 + d + 2\, xb)$

Out[ ]=
$c\,(1 + d + xb - d\, xb)$

Out[ ]=
True

```
In[•]:= (*This is T*)
T[xvar_, dvar_] :=
  FullSimplify[Sum[p[i] × p[j] (zp[i, j] - zb) (z[i, j] - zb), {i, 1, 2}, {j, 1, 2}] /.
      {p[1] → pa, p[2] → pA} /. {z[1, 1] → zaa, z[1, 2] → zAa,
      z[2, 1] → zAa, z[2, 2] → zAA, zp[1, 1] → zaap, zp[1, 2] → zAap,
      zp[2, 1] → zAap, zp[2, 2] → zAAp}] /. {xb → xvar, d → dvar}

Simplify[T[xb, d] == Px[xb] Dd[xb, d]^2]
T[xb, 0]

Out[•]=
True

Out[•]=
-c^2 (-1 + xb) xb

In[•]:= (*This is zeta and H*)
zeta[xvar_, dvar_] :=
  FullSimplify[Sum[p[i] × p[j] × zp[i, j], {i, 1, 2}, {j, 1, 2}] /.
      {p[1] → pa, p[2] → pA} /. {zp[1, 1] → zaap, zp[1, 2] → zAap,
      zp[2, 1] → zAap, zp[2, 2] → zAAp}] /. {xb → xvar, d → dvar}

H[xvar_, dvar_] :=
  FullSimplify[Sum[p[i] × p[j] (zp[i, j] - zb) (z[i, j] - zb), {i, 1, 2}, {j, 1, 2}] /
       Sum[p[i] × p[j] (z[i, j] - zb)^2, {i, 1, 2}, {j, 1, 2}] /.
      {p[1] → pa, p[2] → pA} /. {z[1, 1] → zaa, z[1, 2] → zAa,
      z[2, 1] → zAa, z[2, 2] → zAA, zp[1, 1] → zaap, zp[1, 2] → zAap,
      zp[2, 1] → zAap, zp[2, 2] → zAAp}] /. {xb → xvar, d → dvar}

Simplify[zeta[xb, d] == zb]
Simplify[H[xb, d] == Px[xb]/P[xb, d] Dd[xb, d]^2]
Simplify[H[xb, 0]]

Out[•]=
True

Out[•]=
True

Out[•]=
1/2
```

In[ ]:=
```
(*This is P'=var[z']*)
Pp[xvar_, dvar_] :=
 FullSimplify[Sum[p[i] × p[j] (zp[i, j] - zb)^2, {i, 1, 2}, {j, 1, 2}] /.
     {p[1] → pa, p[2] → pA} /. {zp[1, 1] → zaap, zp[1, 2] → zAap,
     zp[2, 1] → zAap, zp[2, 2] → zAAp}] /. {xb → xvar, d → dvar}

Simplify[Pp[xb, d] == 1/2 Px[xb] Dd[xb, d]^2]
Simplify[Pp[xb, 0]]
```

Out[ ]=

True

Out[ ]=

$$-\frac{1}{2} c^2 (-1 + xb) xb$$

In[ ]:=
```
(*This checks that, with d=0,
offspring phenotype is perfectly predicted by the parent-
 offspring linear regression,
which happens if the equality holds (eq. 18 in main text)*)

Simplify[Pp[xb, d] == H[xb, d]^2 P[xb, d]]
Simplify[Pp[xb, d] == H[xb, d]^2 P[xb, d] /. d → 0]
```

Out[ ]=

$$\frac{c\, d\, (-1 + xb)\, xb\, (-1 + d\, (-1 + 2\, xb))}{1 + d\, (2 - 4\, xb) + d^2 \left(1 - 2\, xb + 2\, xb^2\right)} == 0$$

Out[ ]=

True

In[ ]:=
```
zb
```

Out[ ]=

$$2\, c\, xb\, (1 + d - d\, xb)$$

In[ ]:=
```
(*This is eta, which shows they are all zero if d=0*)
etaaa = zaap - zeta[xb, d] - H[xb, d] (zaa - zb);
etaAa = zAap - zeta[xb, d] - H[xb, d] (zAa - zb);
etaAA = zAAp - zeta[xb, d] - H[xb, d] (zAA - zb);

Simplify[{etaaa, etaAa, etaAA}]
Simplify[{etaaa, etaAa, etaAA} /. d → 0]
```

Out[ ]=

$$\left\{\frac{c\, (-1 + d)\, d\, xb^2\, (-1 + d\, (-1 + 2\, xb))}{1 + d\, (2 - 4\, xb) + d^2 \left(1 - 2\, xb + 2\, xb^2\right)},\right.$$
$$\left. -\frac{c\, d\, (-1 + xb)\, xb\, (-1 + d\, (-1 + 2\, xb))}{1 + d\, (2 - 4\, xb) + d^2 \left(1 - 2\, xb + 2\, xb^2\right)},\ -\frac{c\, d\, (1 + d)\, (-1 + xb)^2\, (-1 + d\, (-1 + 2\, xb))}{1 + d\, (2 - 4\, xb) + d^2 \left(1 - 2\, xb + 2\, xb^2\right)}\right\}$$

Out[ ]=

{0, 0, 0}

```
(*This is breeding value for genotype kl*)
a[k_, l_] := xi[xb, d] + Dd[xb, d] (y[k, l] - yb);

(*This is the mean breeding value and
 a check that it equals the mean phenotype*)

ab =
  Simplify[Sum[p[k] × p[l] × a[k, l], {k, 1, 2}, {l, 1, 2}] /. {p[1] → pa, p[2] → pA}];

ab == zb

(*This checks that xi and zeta equals the mean phenotype. The
 latter entails that there is no pure transmission bias*)
FullSimplify[xi[xb, d] == zeta[xb, d] == zb]
```

Out[ ]=

True

Out[ ]=

True

```
(*This checks that assumption I of the Lande equation does not
  hold: that is z' is not E[a|z]=xi[xb,d]+Dd[xb,d](y[k,l]-yb)*)
FullSimplify[
 Table[zp[k, l] == xi[xb, d] + 1/2 Dd[xb, d] (y[k, l] - yb), {k, 1, 2}, {l, 1, 2}]]
```

Out[ ]=

{{True, True}, {True, True}}

In[ ]:=

```
(*This is G, the additive genetic variance*)
G[xvar_, dvar_] :=
 FullSimplify[Sum[p[k] × p[l] (a[k, l] - zb)^2, {k, 1, 2}, {l, 1, 2}] /.
     {p[1] → pa, p[2] → pA} /. {x[1] → xa, x[2] → xA}] /. {xb → xvar, d → dvar}

G[xb, d] == 2 Px[xb] Dd[xb, d]^2
G[xb, d] == 2 T[xb, d]
G[xb, 0]
```

Out[ ]=

True

Out[ ]=

True

Out[ ]=

$-2 c^2 (-1 + xb) xb$

In[ ]:=
```
(*This is heritability*)
h2[xb_, d_] := FullSimplify[G[xb, d] / P[xb, d]]

Simplify[h2[xb, d] == 2 Px[xb] / P[xb, d] Dd[xb, d]^2]
Simplify[h2[xb, d] == 2 H[xb, d]]
h2[xb, 0]
```

Out[ ]=
```
True
```

Out[ ]=
```
True
```

Out[ ]=
```
1
```

In[ ]:=
```
(*Some evaluations*)
Simplify[P[xb, d] /. d → 0]
G[xb, d] /. {d → 0}
T[xb, d] /. {d → 0}
```

Out[ ]=

$-2 c^2 (-1 + xb) xb$

Out[ ]=

$-2 c^2 (-1 + xb) xb$

Out[ ]=

$-c^2 (-1 + xb) xb$

```
(*This plots the coefficients vs allele frequency for the indicated d*)

c = 1;
d = 2;

Plot1FileName = If[d == -1, "a", If[d == -0.8, "b", If[d == 0, "c",
      If[d == 0.8, "d", If[d == 1, "e", If[d == 1.2, "f", If[d == 2, "g"]]]]]]];
Plot2FileName = If[d == -1, "h", If[d == -0.8, "i", If[d == 0, "j",
      If[d == 0.8, "k", If[d == 1, "l", If[d == 1.2, "m", If[d == 2, "n"]]]]]]];
Plot3FileName = If[d == -1, "o", If[d == -0.8, "p", If[d == 0, "q",
      If[d == 0.8, "r", If[d == 1, "s", If[d == 1.2, "t", If[d == 2, "u"]]]]]]];
Plot4FileName = If[d == -1, "v", If[d == -0.8, "w", If[d == 0, "x",
      If[d == 0.8, "y", If[d == 1, "z", If[d == 1.2, "aa", If[d == 2, "ab"]]]]]]];

lThickness = .03;
lDashing = .01;

(*Plot genotype-phenotype map*)
BarChart[{zaa, zAa, zAA}, Frame → True,
 FrameTicks → {{Automatic, None}, {{{1, "aa"}, {2, "Aa"}, {3, "AA"}}, None}},
 FrameTicksStyle → Large, PlotRange → {{.5, 3.5}, {0, 3.1}}]
Export[
  StringJoin[ToString[NotebookDirectory[]], "Fig.2", Plot1FileName, ".pdf"], %];

(*Plot zb and D for any chosen d*)
Plot[{zbfun[xb, d], Dd[xb, d]}, {xb, 0, 1},
 PlotRange → {-1, 3.1}, PlotStyle → {{Thickness[lThickness]},
   {Dashing[lDashing], Thickness[lThickness]}}, TicksStyle → Large]
Export[
  StringJoin[ToString[NotebookDirectory[]], "Fig.2", Plot2FileName, ".pdf"], %];

(*Plot G and T for any chosen d*)
Plot[{G[xb, d], T[xb, d], P[xb, d]},
 {xb, 0, 1}, PlotRange → {-0.01, 1.6}, PlotStyle →
  {{Thickness[lThickness]}, {Dashing[lDashing], Thickness[lThickness]},
   {Dashing[lDashing], Thickness[lThickness]}}, TicksStyle → Large]
Export[
  StringJoin[ToString[NotebookDirectory[]], "Fig.2", Plot3FileName, ".pdf"], %];

(*Plot heritability and H for any chosen d*)
Plot[{h2[xb, d], H[xb, d]}, {xb, 0, 1},
 PlotRange → {-.01, 1.1}, PlotStyle → {{Thickness[lThickness]},
   {Dashing[lDashing], Thickness[lThickness]}}, TicksStyle → Large]
Export[
  StringJoin[ToString[NotebookDirectory[]], "Fig.2", Plot4FileName, ".pdf"], %];
```

Out[ ]=

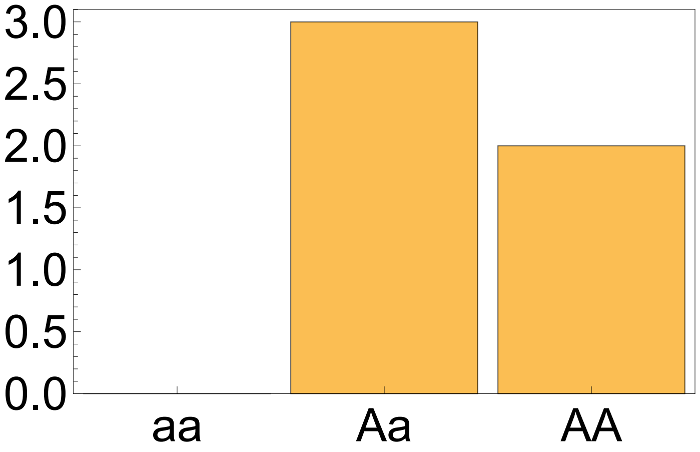
3.0
2.5
2.0
1.5
1.0
0.5
0.0
aa
Aa
AA

Out[ ]=

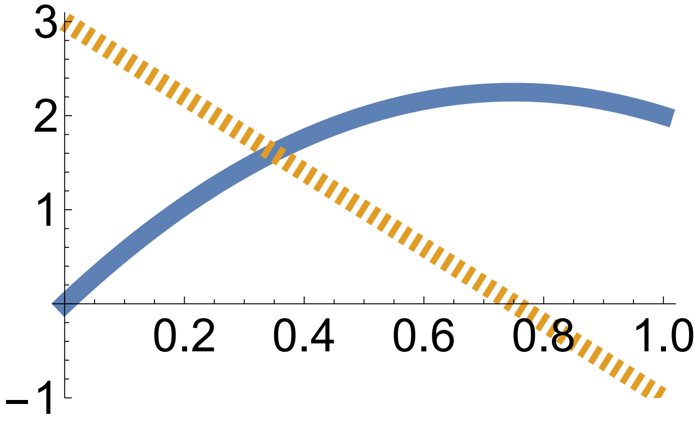
3
2
1
0.2
0.4
0.6
0.8
1.0
−1

Out[ ]=

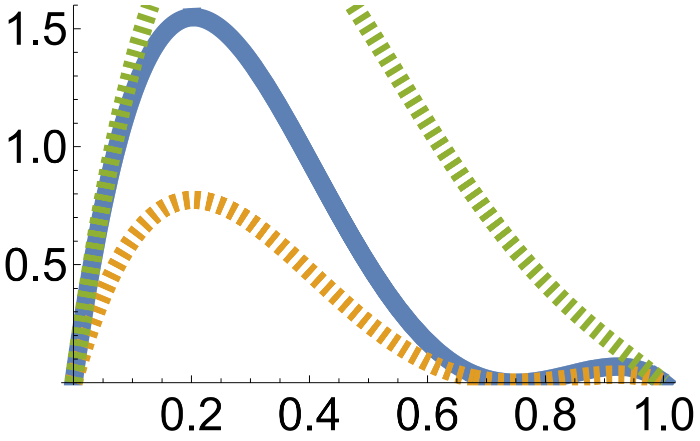
1.5
1.0
0.5
0.2
0.4
0.6
0.8
1.0

Out[ ]=

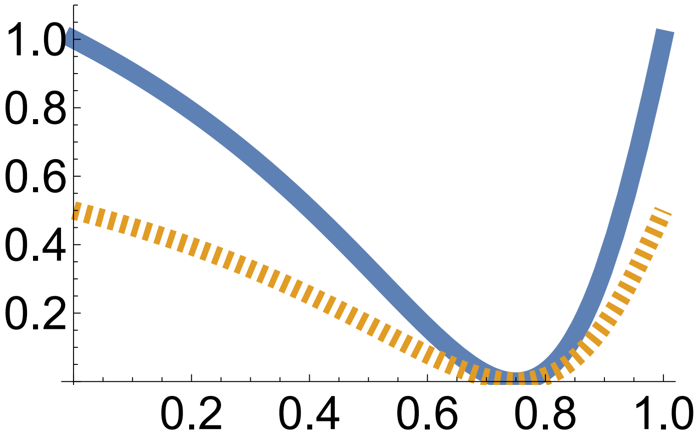

#### Example 2 continued: coefficients of generator equations under selfing, see SI section S9.1

In[ ]:=
```
Clear["Global`*"]
```

In[ ]:=
```
(*This lists gene content, haplotype frequencies,
genotype frequencies, and phenotypes*)

xa = 0; xA = 1;
pa = (1 - xb); pA = xb;
paa = pa^2; pAa = 2 pa pA; pAA = pA^2;

zaa = 0;
zAa = c (1 + d);
zAA = 2 c;
```

In[ ]:=
```
(*This is Px=var[x], the allelic variance*)
Px[xvar_] = Simplify[Sum[p[i] (x[i] - xb)^2, {i, 1, 2}] /. {p[1] → pa, p[2] → pA} /.
      {x[1] → xa, x[2] → xA}] /. {xb → xvar};
Px[xb]

(*This is the mean phenotype*)
zb = Simplify[Sum[p[i] × p[j] × z[i, j], {i, 1, 2}, {j, 1, 2}] /.
    {p[1] → pa, p[2] → pA, z[1, 1] → zaa, z[1, 2] → zAa, z[2, 1] → zAa, z[2, 2] → zAA}]

(*This is to plot the mean phenotype as a function of allele frequency*)
zbfun[xvar_, dvar_] = zb /. {xb → xvar, d → dvar} ;
```

Out[ ]=

$-((-1 + xb)\ xb)$

Out[ ]=

$2\ c\ xb\ (1 + d - d\ xb)$

In[ ]:=
```
(*This is genotype content of genotype kl*)
y[k_, l_] :=
 If[k == 1 && l == 1, 0, If[(k == 1 && l == 2) || (k == 2 && l == 1), 1, If[k == 2 && l == 2, 2]]]

(*This is the average genotype content*)
yb =
  Simplify[Sum[p[i] × p[j] × y[i, j], {i, 1, 2}, {j, 1, 2}] /. {p[1] → pa, p[2] → pA}];

(*This is xi and D*)
xi[xvar_, dvar_] := FullSimplify[Sum[p[i] × p[j] × z[i, j], {i, 1, 2}, {j, 1, 2}] /.
     {p[1] → pa, p[2] → pA} /. {z[1, 1] → zaa, z[1, 2] → zAa,
     z[2, 1] → zAa, z[2, 2] → zAA}] /. {xb → xvar, d → dvar}

Dd[xvar_, dvar_] :=
 FullSimplify[(Sum[p[i] × p[j] (z[i, j] - zb) (y[i, j] - yb), {i, 1, 2}, {j, 1, 2}]) /
              (Sum[p[i] × p[j] (y[i, j] - yb)^2, {i, 1, 2}, {j, 1, 2}]) /.
      {p[1] → pa, p[2] → pA} /. {z[1, 1] → zaa, z[1, 2] → zAa, z[2, 1] → zAa,
      z[2, 2] → zAA} /. {x[1] → xa, x[2] → xA}] /. {xb → xvar, d → dvar}

yb
```

Out[ ]=

$2\,xb$

In[ ]:=
```
(*This is P*)
P[xvar_, dvar_] :=
 FullSimplify[Sum[p[i] × p[j] (z[i, j] - zb)^2, {i, 1, 2}, {j, 1, 2}] /.
     {p[1] → pa, p[2] → pA} /. {z[1, 1] → zaa, z[1, 2] → zAa,
     z[2, 1] → zAa, z[2, 2] → zAA}] /. {xb → xvar, d → dvar}

Simplify[P[xb, d] == 2 Px[xb] Dd[xb, d]^2 + (2 Px[xb] c d)^2]
Simplify[P[xb, 0]]
```

Out[ ]=

True

Out[ ]=

$-2\,c^2\,(-1 + xb)\,xb$

In[ ]:=
```
(*This enters the mean phenotype in offspring
 of a given parental phenotype under selfing*)

zaap = zaa; zAap = 1/4 zaa + 1/2 zAa + 1/4 zAA; zAAp = zAA;

zp[k_, l_] := If[k == 1 && l == 1, zaap,
  If[(k == 1 && l == 2) || (k == 2 && l == 1), zAap, If[k == 2 && l == 2, zAAp]]]
```

In[ ]:=
```
Simplify[zaap]
Simplify[zAap]
Simplify[zAAp]
```

Out[ ]=

0

Out[ ]=

$\frac{1}{2}$ c (2 + d)

Out[ ]=

2 c

In[ ]:=
```
(*This is T*)
T[xvar_, dvar_] :=
 FullSimplify[Sum[p[i] × p[j] (zp[i, j] - zb) (z[i, j] - zb), {i, 1, 2}, {j, 1, 2}] /.
     {p[1] → pa, p[2] → pA} /. {z[1, 1] → zaa, z[1, 2] → zAa,
     z[2, 1] → zAa, z[2, 2] → zAA, zp[1, 1] → zaap, zp[1, 2] → zAap,
     zp[2, 1] → zAap, zp[2, 2] → zAAp}] /. {xb → xvar, d → dvar}

(*Some evaluations*)
T[xb, d]
T[xb, 0]
T[.99, 1.2]
```

Out[ ]=

$-c^2 (-1 + xb)\, xb\, (2 + d (3 + d - 6\, xb + 2\, d (-1 + xb)\, xb))$

Out[ ]=

$-2\, c^2 (-1 + xb)\, xb$

Out[ ]=

$-0.00115347\, c^2$

In[∘]:=
```
(*This is zeta and H*)
zeta[xvar_, dvar_] :=
 FullSimplify[Sum[p[i] × p[j] × zp[i, j], {i, 1, 2}, {j, 1, 2}] /.
     {p[1] → pa, p[2] → pA} /. {zp[1, 1] → zaap, zp[1, 2] → zAap,
     zp[2, 1] → zAap, zp[2, 2] → zAAp}] /. {xb → xvar, d → dvar}

H[xvar_, dvar_] :=
 FullSimplify[Sum[p[i] × p[j] (zp[i, j] - zb) (z[i, j] - zb), {i, 1, 2}, {j, 1, 2}] /
      Sum[p[i] × p[j] (z[i, j] - zb)^2, {i, 1, 2}, {j, 1, 2}] /.
     {p[1] → pa, p[2] → pA} /. {z[1, 1] → zaa, z[1, 2] → zAa,
     z[2, 1] → zAa, z[2, 2] → zAA, zp[1, 1] → zaap, zp[1, 2] → zAap,
     zp[2, 1] → zAap, zp[2, 2] → zAAp}] /. {xb → xvar, d → dvar}

(*Some evaluations*)
H[xb, d]
H[.99, 1.2]
```

Out[∘]=

$$\frac{2 + d\,(3 + d - 6\,xb + 2\,d\,(-1 + xb)\,xb)}{2 + 2\,d\,(2 + d - 4\,xb + 2\,d\,(-1 + xb)\,xb)}$$

Out[∘]=

-0.97929

In[∘]:=
```
(*This is pure transmission bias*)
Simplify[zeta[xb, d] - zb]
```

Out[∘]=

$c\,d\,(-1 + xb)\,xb$

In[∘]:=
```
(*This is eta, which shows they are all zero if d=0*)
etaaa = zaap - zeta[xb, d] - H[xb, d] (zaa - zb);
etaAa = zAap - zeta[xb, d] - H[xb, d] (zAa - zb);
etaAA = zAAp - zeta[xb, d] - H[xb, d] (zAA - zb);

Simplify[{etaaa, etaAa, etaAA}]
Simplify[{etaaa, etaAa, etaAA} /. d → 0]
```

Out[∘]=

$$\left\{-\frac{c\,(-1 + d)\,d\,xb^2}{1 + d\,(2 - 4\,xb) + d^2\,(1 - 2\,xb + 2\,xb^2)},\ \frac{c\,d\,(-1 + xb)\,xb}{1 + d\,(2 - 4\,xb) + d^2\,(1 - 2\,xb + 2\,xb^2)},\ \frac{c\,d\,(1 + d)\,(-1 + xb)^2}{1 + d\,(2 - 4\,xb) + d^2\,(1 - 2\,xb + 2\,xb^2)}\right\}$$

Out[∘]=

{0, 0, 0}

In[ ]:=
```
(*This is breeding value for genotype kl*)
a[k_, l_] := xi[xb, d] + Dd[xb, d] (y[k, l] - yb);

(*This is the mean breeding value*)
ab =
  Simplify[Sum[p[k] × p[l] × a[k, l], {k, 1, 2}, {l, 1, 2}] /. {p[1] → pa, p[2] → pA}];

(*This checks that mean breeding values equals the mean phenotype*)

ab == zb
```
Out[ ]=
```
True
```
In[ ]:=
```
(*This checks that assumption I of the Lande equation does not
  hold: that is z' is not E[a|z]=xi[xb,d]+Dd[xb,d](y[k,l]-yb)*)
FullSimplify[
 Table[zp[k, l] == xi[xb, d] + Dd[xb, d] (y[k, l] - yb), {k, 1, 2}, {l, 1, 2}]]
```
Out[ ]=
```
{{c d xb == 0, 2 c d xb == c d}, {2 c d xb == c d, c d xb == c d}}
```
In[ ]:=
```
(*This is G, the additive genetic variance*)
G[xvar_, dvar_] :=
 FullSimplify[Sum[p[k] × p[l] (a[k, l] - zb)^2, {k, 1, 2}, {l, 1, 2}] /.
     {p[1] → pa, p[2] → pA} /. {x[1] → xa, x[2] → xA}] /. {xb → xvar, d → dvar}

G[xb, d] == 2 Px[xb] Dd[xb, d]^2
G[xb, 0]
```
Out[ ]=
```
True
```
Out[ ]=
```
-2 c^2 (-1 + xb) xb
```

In[ ]:=
```
(*This is heritability*)
h2[xb_, d_] := FullSimplify[G[xb, d]/P[xb, d]]

Simplify[h2[xb, d] == 2 Px[xb]/P[xb, d] Dd[xb, d]^2]
Simplify[h2[xb, d] == H[xb, d]]
h2[xb, 0]
```

Out[ ]=
True

Out[ ]=
$$\frac{d\,(1+d-2\,xb-6\,d\,xb+6\,d\,xb^2)}{1+d\,(2-4\,xb)+d^2\,(1-2\,xb+2\,xb^2)} == 0$$

Out[ ]=
1

In[ ]:=
```
(*This plots the coefficients vs allele frequency for the indicated d*)

c = 1;
d = 2;

Plot1FileName = If[d == -1, "a", If[d == -0.8, "b", If[d == 0, "c",
      If[d == 0.8, "d", If[d == 1, "e", If[d == 1.2, "f", If[d == 2, "g"]]]]]]];
Plot2FileName = If[d == -1, "h", If[d == -0.8, "i", If[d == 0, "j",
      If[d == 0.8, "k", If[d == 1, "l", If[d == 1.2, "m", If[d == 2, "n"]]]]]]];
Plot3FileName = If[d == -1, "o", If[d == -0.8, "p", If[d == 0, "q",
      If[d == 0.8, "r", If[d == 1, "s", If[d == 1.2, "t", If[d == 2, "u"]]]]]]];
Plot4FileName = If[d == -1, "v", If[d == -0.8, "w", If[d == 0, "x",
      If[d == 0.8, "y", If[d == 1, "z", If[d == 1.2, "aa", If[d == 2, "ab"]]]]]]];

lThickness = .03;
lDashing = .01;

(*Plot genotype-phenotype map*)
BarChart[{zaa, zAa, zAA}, Frame → True,
 FrameTicks → {{Automatic, None}, {{{1, "aa"}, {2, "Aa"}, {3, "AA"}}, None}},
 FrameTicksStyle → Large, PlotRange → {{.5, 3.5}, {0, 3.1}}]
Export[StringJoin[ToString[NotebookDirectory[]],
   "Fig.S4", Plot1FileName, ".pdf"], %];

(*Plot zb and D for any chosen d*)
Plot[{zbfun[xb, d], Dd[xb, d]}, {xb, 0, 1},
 PlotRange → {-1, 3.1}, PlotStyle → {{Thickness[lThickness]},
   {Dashing[lDashing], Thickness[lThickness]}}, TicksStyle → Large]
Export[StringJoin[ToString[NotebookDirectory[]],
   "Fig.S4", Plot2FileName, ".pdf"], %];

(*Plot G and T for any chosen d*)
Plot[{G[xb, d], T[xb, d], P[xb, d]},
 {xb, 0, 1}, PlotRange → {-0.1, 1.6}, PlotStyle →
  {{Thickness[lThickness]}, {Dashing[lDashing], Thickness[lThickness]},
   {Dashing[lDashing], Thickness[lThickness]}}, TicksStyle → Large]
Export[StringJoin[ToString[NotebookDirectory[]],
   "Fig.S4", Plot3FileName, ".pdf"], %];

(*Plot heritability and H for any chosen d*)
Plot[{h2[xb, d], H[xb, d]}, {xb, 0, 1},
 PlotRange → {-1.1, 2.1}, PlotStyle → {{Thickness[lThickness]},
   {Dashing[lDashing], Thickness[lThickness]}}, TicksStyle → Large]
Export[
  StringJoin[ToString[NotebookDirectory[]], "Fig.S4", Plot4FileName, ".pdf"], %];
```

Out[ ]=

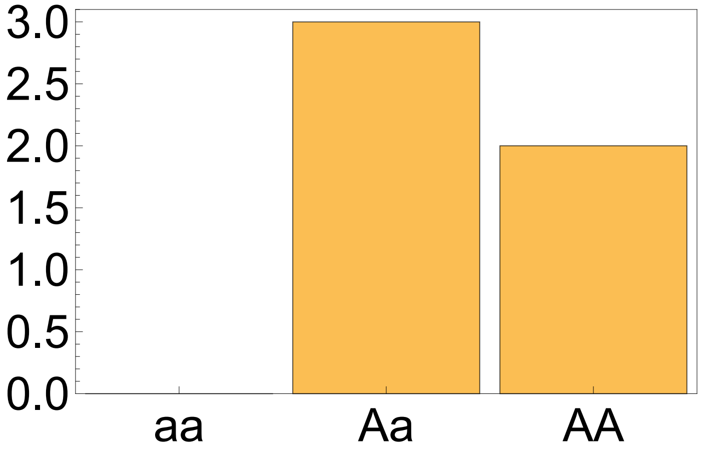
3.0
2.5
2.0
1.5
1.0
0.5
0.0
aa
Aa
AA

Out[ ]=

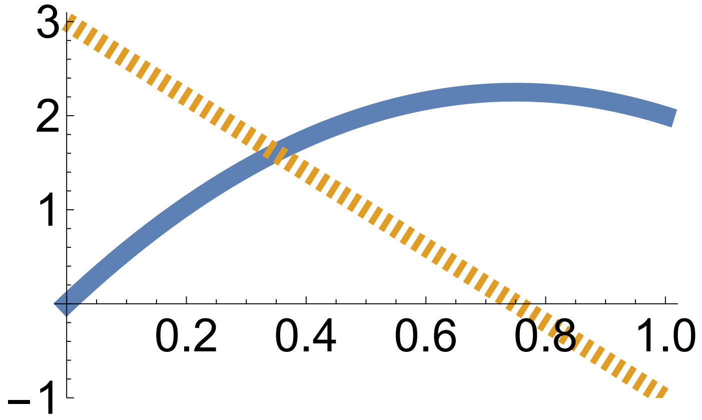
3
2
1
−1
0.2
0.4
0.6
0.8
1.0

Out[ ]=

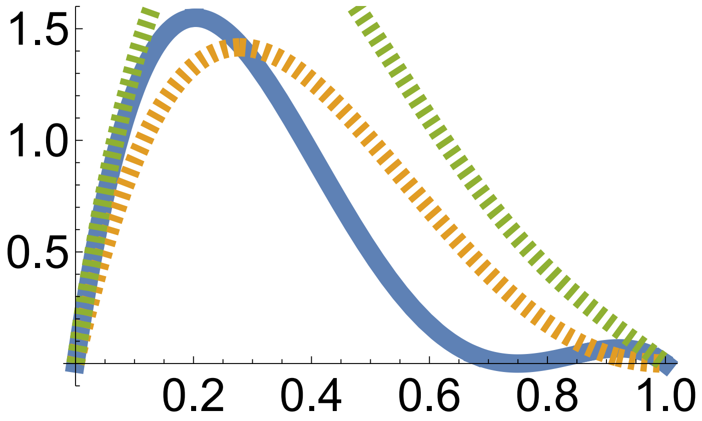
1.5
1.0
0.5
0.2
0.4
0.6
0.8
1.0

Out[ ]=

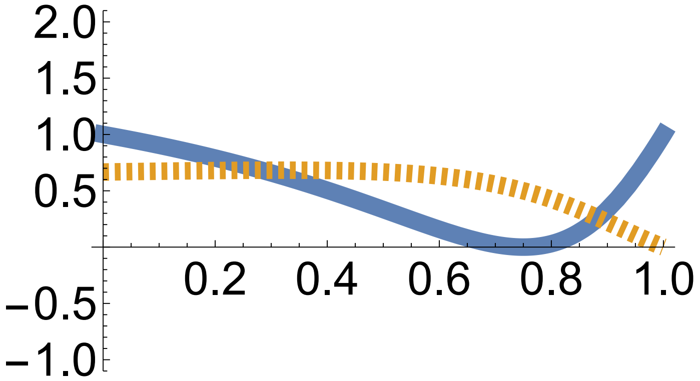

#### Example 3 exact evo-devo dynamics: one bialellic locus, one phenotype

In[ ]:=
```
Clear["Global`*"]
```

In[ ]:=
```
(*This lists the gene content of haplotypes,
and of average offspring haplotypes*)
xa = 0; xA = 1;
xap = pa xa + pA (1/2 xA + 1/2 xa) ; xAp = pa (1/2 xA + 1/2 xa) + pA xA;

(*This lists the mean phenotype in offspring,
in the population, and the phenotypes for a given genotype*)
zaap = paa zaa + pAa (1/2 zAa + 1/2 zaa) + pAA zAa;
zAap = paa (1/2 zAa + 1/2 zaa) + pAa (1/4 zaa + 1/2 zAa + 1/4 zAA) + pAA (1/2 zAa + 1/2 zAA);
zAAp = paa zAa + pAa (1/2 zAa + 1/2 zAA) + pAA zAA;
zb = paa zaa + pAa zAa + pAA zAA;
zaa = 0; zAa = c (1 + d); zAA = 2 c;

(*This lists the frequency of haplotypes and of genotypes*)
pa = (1 - xb);
pA = xb;
paa = (1 - xb)^2;
pAa = 2 xb (1 - xb); pAA = xb^2;
```

In[ ]:=
```
{zaa, zAa, zAA}
```

Out[ ]=
```
{0, c (1 + d), 2 c}
```

In[∘ ]:=
```
(*This is Tx*)
Tx[xvar_] = FullSimplify[
   Sum[p[i] (xp[i] - xb) (x[i] - xb), {i, 1, 2}] /. {p[1] → pa, p[2] → pA} /.
     {x[1] → xa, x[2] → xA} /. {xp[1] → xap, xp[2] → xAp}] /. {xb → xvar}

(*This is Px = var[x]*)
Px[xvar_] =
 FullSimplify[Sum[p[i] (x[i] - xb)^2, {i, 1, 2}] /. {p[1] → pa, p[2] → pA} /.
     {x[1] → xa, x[2] → xA} /. {xp[1] → xap, xp[2] → xAp}] /. {xb → xvar}

(*This is Px' = var[x']*)
Pxp[xvar_] =
 FullSimplify[Sum[p[i] (xp[i] - xb)^2, {i, 1, 2}] /. {p[1] → pa, p[2] → pA} /.
     {x[1] → xa, x[2] → xA} /. {xp[1] → xap, xp[2] → xAp}] /. {xb → xvar}
```

Out[∘ ]=

$$-\frac{1}{2}\ (-1 + \text{xvar})\ \text{xvar}$$

Out[∘ ]=

$$-((-1 + \text{xvar})\ \text{xvar})$$

Out[∘ ]=

$$-\frac{1}{4}\ (-1 + \text{xvar})\ \text{xvar}$$

In[∘ ]:=
```
(*This is zetax*)
zetax[xvar_] :=
 FullSimplify[Sum[p[i] × xp[i], {i, 1, 2}] /. {p[1] → pa, p[2] → pA} /.
    {xp[1] → xap, xp[2] → xAp}] /. {xb → xvar}
```

In[∘ ]:=
```
zetax[xb]
```

Out[∘ ]=

xb

In[∘ ]:=
```
(*This is Hx*)
Hx = Tx[xb] / Px[xb]
```

Out[∘ ]=

$$\frac{1}{2}$$

In[∘ ]:=
```
(*This is etax*)
etaa = xap - zetax[xb] - Hx (xa - xb)
etaA = Simplify[xAp - zetax[xb] - Hx (xA - xb)]
```

Out[∘ ]=

0

Out[∘ ]=

0

```
In[•]:= (*This is cov[w,x]*)
covwx[xvar_] := Simplify[
   Sum[p[i] (x[i] - xb) (wh[i, xb, θ, σ] - 1), {i, 1, 2}] /. {x[1] → xa, x[2] → xA} /.
    {p[1] → pa, p[2] → pA}] /. {xb → xvar}

(*This is betax*)
betax[xb_] := covwx[xb] / Px[xb]

betax[xb]
Out[•]=
-wh[1, xb, θ, σ] + wh[2, xb, θ, σ]

In[•]:= (*This is cov[w,x']*)
covwxp[xvar_] := Simplify[Sum[p[i] (xp[i] - xb) (wh[i, xb, θ, σ] - 1), {i, 1, 2}] /.
     {xp[1] → xap, xp[2] → xAp} /. {p[1] → pa, p[2] → pA}] /. {xb → xvar}

(*This is betax'*)
betaxp[xb_] := covwxp[xb] / Pxp[xb]

betaxp[xb]
Out[•]=
-2 (wh[1, xb, θ, σ] - wh[2, xb, θ, σ])

In[•]:= (*This is the adaptability of the haplotype*)
Ad[xb_] = (2 betax[xb]) Tx[xb] (2 betax[xb])

(*This is the respondability of the haplotype*)
Res[xb_] = betaxp[xb] × Pxp[xb] × betaxp[xb]
Out[•]=
-2 (-1 + xb) xb (-wh[1, xb, θ, σ] + wh[2, xb, θ, σ])^2
Out[•]=
-((-1 + xb) xb (wh[1, xb, θ, σ] - wh[2, xb, θ, σ])^2)

In[•]:= (*This is mean absolute fitness*)
Wb[xvar_, θ_, σ_] :=
 FullSimplify[Sum[p[i] × p[j] × W[i, j, θ, σ], {i, 1, 2}, {j, 1, 2}] /.
    {p[1] → pa, p[2] → pA}] /. {xb → xvar}

(*This is the absolute fitness of haplotype i*)
Wh[i_, θ_, σ_] := Sum[p[j] × W[i, j, θ, σ], {j, 1, 2}] /. {p[1] → pa, p[2] → pA}

(*This checks that the selection gradient of mean haplotype is 2(WA-Wa)*)
Simplify[
 D[Wb[xb, θ, σ], xb] == 2 (Wh[2, θ, σ] - Wh[1, θ, σ]) /. {W[2, 1, θ, σ] → W[1, 2, θ, σ]}]
Out[•]=
True
```

In[•]:=
```
(*This specifies fitness for genotype ij*)
W[i_, j_, θ_, σ_] :=
 Exp[-(z[i, j] - θ)^2/σ^2] /. {z[1, 1] → zaa, z[1, 2] → zAa, z[2, 1] → zAa, z[2, 2] → zAA}
w[i_, j_, xb_, θ_, σ_] :=
 W[i, j, θ, σ] / Wb[xb, θ, σ] /. {z[1] → zaa, z[2] → zAa, z[3] → zAA}

FullSimplify[Sum[p[i] × p[j] × w[i, j, xb, θ, σ], {i, 1, 2}, {j, 1, 2}] /.
   {z[1] → zaa, z[2] → zAa, z[3] → zAA} /. {p[1] → pa, p[2] → pA}]

(*This is the relative fitness for haplotype i*)
wh[i_, xb_, θ_, σ_] :=
 Sum[p[j] × w[i, j, xb, θ, σ], {j, 1, 2}] /. {p[1] → pa, p[2] → pA}
```

Out[•]=
```
1
```

In[•]:=
```
(*This is P*)
P[xvar_] := FullSimplify[
   Sum[p[i] × p[j] (z[i, j] - zb)^2, {i, 1, 2}, {j, 1, 2}] /. {p[1] → pa, p[2] → pA} /.
    {z[1, 1] → zaa, z[1, 2] → zAa, z[2, 1] → zAa, z[2, 2] → zAA}] /. {xb → xvar}

Dd[xb_, d_] := c (1 + d (1 - 2 xb))
Simplify[P[xb] == 2 Px[xb] Dd[xb, d]^2 + (2 Px[xb] c d)^2]
```

Out[•]=
```
True
```

In[•]:=
```
(*This checks that ΔP takes the form given in equation 103*)

(*This is the change in additive genetic variance*)
Simplify[D[2 Px[xb[t]] Dd[xb[t], d]^2, t] ==
  2 Dd[xb[t], d] xb′[t] (Dd[xb[t], d] (1 - 2 xb[t]) - 4 c d Px[xb[t]])]

(*This is the change in dominance genetic variance*)
Simplify[D[(2 Px[xb[t]] c d)^2, t] == 8 Px[xb[t]] (c d )^2 (1 - 2 xb[t]) xb′[t]]
```

Out[•]=
```
True
```

Out[•]=
```
True
```

In[•]:=
```
(*This checks that offspring haplotype content is perfectly
 predicted from parent haplotype content; eq. 18 in main text*)

Pxp[xb] == Hx^2 Px[xb]
```

Out[•]=
```
True
```

```
In[ ]:= (*This runs the numerical solutions for the parameter values indicated*)

c = 1;
d = 2;

θ = 1;
σ = Sqrt[10];

Plot1FileName =
  If[θ == 1.5 && d == 0, "a", If[θ == 1 && d == 0, "f", If[θ == 1 && d == 2, "k"]]];
Plot2FileName =
  If[θ == 1.5 && d == 0, "b", If[θ == 1 && d == 0, "g", If[θ == 1 && d == 2, "l"]]];
Plot3FileName =
  If[θ == 1.5 && d == 0, "c", If[θ == 1 && d == 0, "h", If[θ == 1 && d == 2, "m"]]];

tend = 200;

x1binit = {0.01, 1, .1};
(*This is to sample the initial conditions for allele frequency*)

InitialConditions =
  Table[{x1bSol[0, i] = i}, {i, x1binit[[1]], x1binit[[2]], x1binit[[3]]}];

Table[
  Table[{x1bSol[t + 1, i] = x1bSol[t, i] + (Tx[x1bSol[t, i]] × betax[x1bSol[t, i]])},
   {t, 0, tend}], {i, x1binit[[1]], x1binit[[2]], x1binit[[3]]}];
```

```
(*This makes plots of allele frequency, mean phenotype,
phenotype variance, and mean absolute fitness over time*)

yAxisLimitz[d_] := If[d ≥ 2, 2.5, 2]
yAxisLimitP[d_] := If[d ≥ 2, 2.5, .6]

zbfun[xvar_] = zb /. {xb → xvar} ;(*To plot the mean phenotype*)

lThickness = 0.02; (*Line thickness*)

(*This is allele frequency over time*)
Show[
 Table[ListLinePlot[{Table[x1bSol[t, i], {t, 0, tend}]}, PlotRange → {-0.1, 1.1},
   TicksStyle → Large, PlotStyle → Thickness[lThickness]],
  {i, x1binit⟦1⟧, x1binit⟦2⟧, x1binit⟦3⟧}]]
Export[
  StringJoin[ToString[NotebookDirectory[]], "Fig.3", Plot1FileName, ".pdf"], %];

(*This is mean phenotype over time*)
Show[Table[
  ListLinePlot[{Table[zbfun[x1bSol[t, i]], {t, 0, tend}], Table[θ, {t, 0, tend}]},
   PlotRange → {-0.1, yAxisLimitz[d]}, TicksStyle → Large,
   PlotStyle → {Thickness[lThickness], {Magenta, Thickness[lThickness]}}],
  {i, x1binit⟦1⟧, x1binit⟦2⟧, x1binit⟦3⟧}]]
Export[
  StringJoin[ToString[NotebookDirectory[]], "Fig.3", Plot2FileName, ".pdf"], %];

(*This is phenotype variance over time*)
Show[Table[ListLinePlot[{Table[P[x1bSol[t, i]], {t, 0, tend}]},
   PlotRange → {-0.1 yAxisLimitP[d], yAxisLimitP[d]}, TicksStyle → Large,
   PlotStyle → Thickness[lThickness]], {i, x1binit⟦1⟧, x1binit⟦2⟧, x1binit⟦3⟧}]]
Export[
  StringJoin[ToString[NotebookDirectory[]], "Fig.3", Plot3FileName, ".pdf"], %];

(*This is mean fitness over time*)
Show[Table[ListLinePlot[{Table[Wb[x1bSol[t, i], θ, σ], {t, 0, tend}]},
   PlotRange → {-0.1, 1.1}, TicksStyle → Large,
   PlotStyle → Thickness[lThickness]], {i, x1binit⟦1⟧, x1binit⟦2⟧, x1binit⟦3⟧}]]
```

Out[ ]=

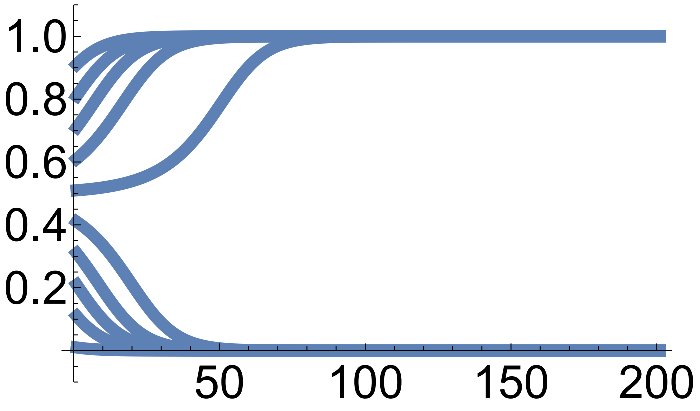

Out[ ]=

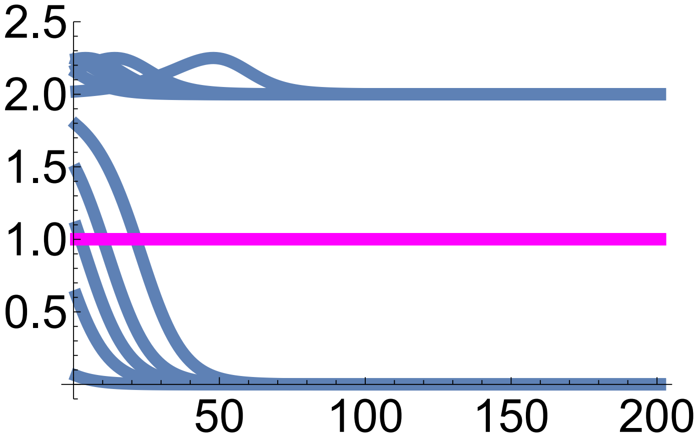

Out[ ]=

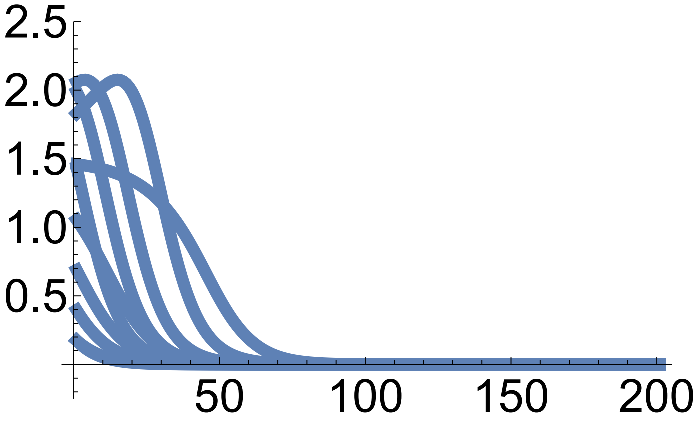

Out[ ]=

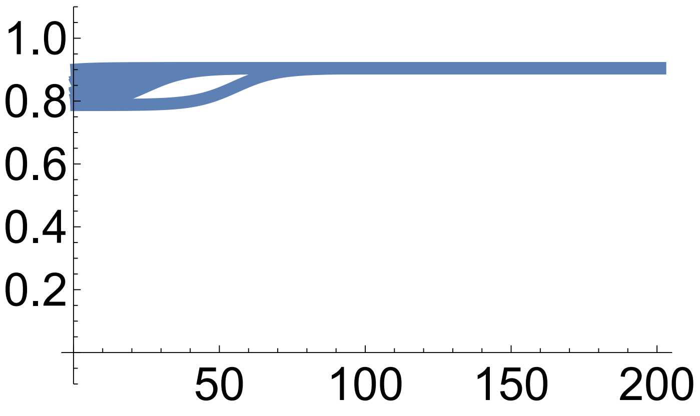

In[ ]:=
```
(*This plots the mean phenotype vs allele frequency
 over the total fitness landscape of the mean haplotype*)
```

```
Plot4FileName =
  If[θ == 1.5 && d == 0, "d", If[θ == 1 && d == 0, "i", If[θ == 1 && d == 2, "n"]]];
Plot5FileName =
  If[θ == 1.5 && d == 0, "e", If[θ == 1 && d == 0, "j", If[θ == 1 && d == 2, "o"]]];

pointSize = 0.05;

(*These are the end points of the
 trajectories of allele frequency and mean phenotype*)
endPoints = Table[{x1bSol[tend, i], zbfun[x1bSol[tend, i]]}, {i, x1binit⟦1⟧,
    x1binit⟦2⟧, x1binit⟦3⟧}, {j, x1binit⟦1⟧, x1binit⟦2⟧, x1binit⟦3⟧}];

(*These are the trajectories of mean phenotype vs allele frequency*)
p0 = Table[ListLinePlot[Table[{x1bSol[t, i], zbfun[x1bSol[t, i]]}, {t, 0, tend}],
    PlotRange → {{0, 1}, {0, yAxisLimitz[d]}}, PlotStyle →
     {Thickness[lThickness]}], {i, x1binit⟦1⟧, x1binit⟦2⟧, x1binit⟦3⟧}];

(*This superimposes the above plots with a
 fitness contour and with the admissible manifold*)
plot =  Show[ContourPlot[Wb[xb, θ, σ], {xb, 0, 1}, {zb, 0, yAxisLimitz[d]},
   FrameStyle → Large, PlotLegends → Automatic, LabelStyle → Large,
   Epilog → {PointSize[pointSize], Point[Flatten[endPoints, 1]]}],
  Plot[zbfun[x1b], {x1b, 0, 1}, PlotStyle → {Red, Thickness[3 lThickness]}],
  p0 /. Line[x_] :→ {Arrowheads[Table[{.1, .2}, {3}]], Arrow[x]}, ListLinePlot[
   Table[{xb, θ}, {xb, 0, 1}], PlotStyle → {Magenta, Thickness[lThickness]}]]

(*This rasterises the image to remove visual artifacts*)
plot = Rasterize[plot, "Image", ImageResolution → 300];

(*This saves the image*)
Export[StringJoin[ToString[NotebookDirectory[]],
   "Fig.3", Plot4FileName, ".pdf"], plot];

(*These are the end points of the trajectories
 of allele frequency and phenotype variance*)
endPointsxP = Table[{x1bSol[tend, i], P[x1bSol[tend, i]]}, {i, x1binit⟦1⟧,
    x1binit⟦2⟧, x1binit⟦3⟧}, {j, x1binit⟦1⟧, x1binit⟦2⟧, x1binit⟦3⟧}];

(*These are the trajectories of phenotype variance vs allele frequency*)
p0 = Table[ListLinePlot[Table[{x1bSol[t, i], P[x1bSol[t, i]]}, {t, 0, tend}],
    PlotRange → {{0, 1}, {0, yAxisLimitP[d]}}, PlotStyle →
     {Thickness[lThickness]}], {i, x1binit⟦1⟧, x1binit⟦2⟧, x1binit⟦3⟧}];

(*This superimposes the above plots with a
 fitness contour and with the admissible manifold*)
```

```
plot = Show[ContourPlot[Wb[xb, θ, σ], {xb, 0, 1}, {P, 0, yAxisLimitP[d]},
    FrameStyle → Large, PlotLegends → Automatic, LabelStyle → Large,
    Epilog → {PointSize[pointSize], Point[Flatten[endPointsxP, 1]]}],
  Plot[P[x1b], {x1b, 0, 1}, PlotStyle → {Red, Thickness[3 lThickness]}],
  p0 /. Line[x_] :→ {Arrowheads[Table[{.1, .2}, {3}]], Arrow[x]}]

(*This rasterises the image to remove visual artifacts*)
plot = Rasterize[plot, "Image", ImageResolution → 300];

(*This saves the image*)
Export[StringJoin[ToString[NotebookDirectory[]],
    "Fig.3", Plot5FileName, ".pdf"], plot];
```

Out[ ]=

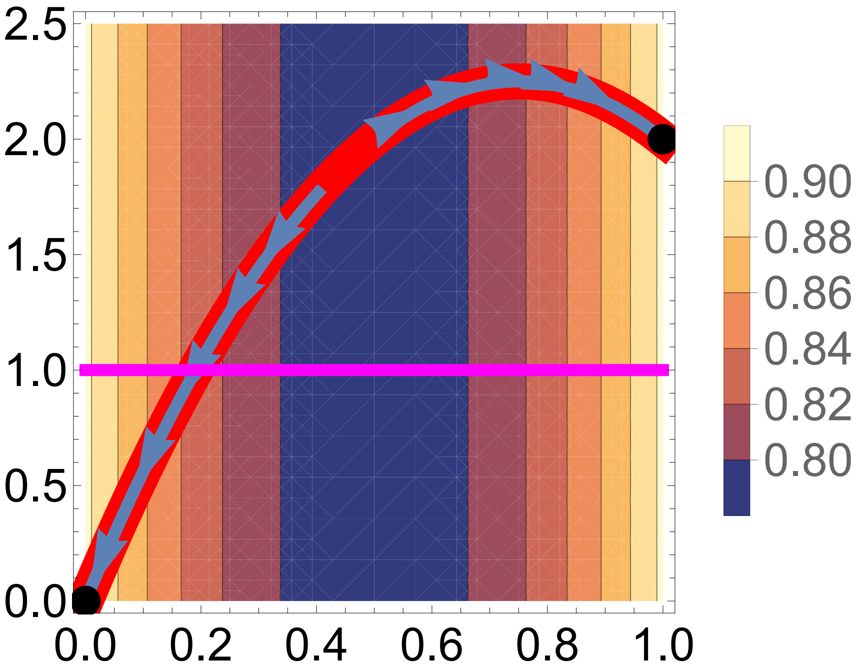

Out[ ]=

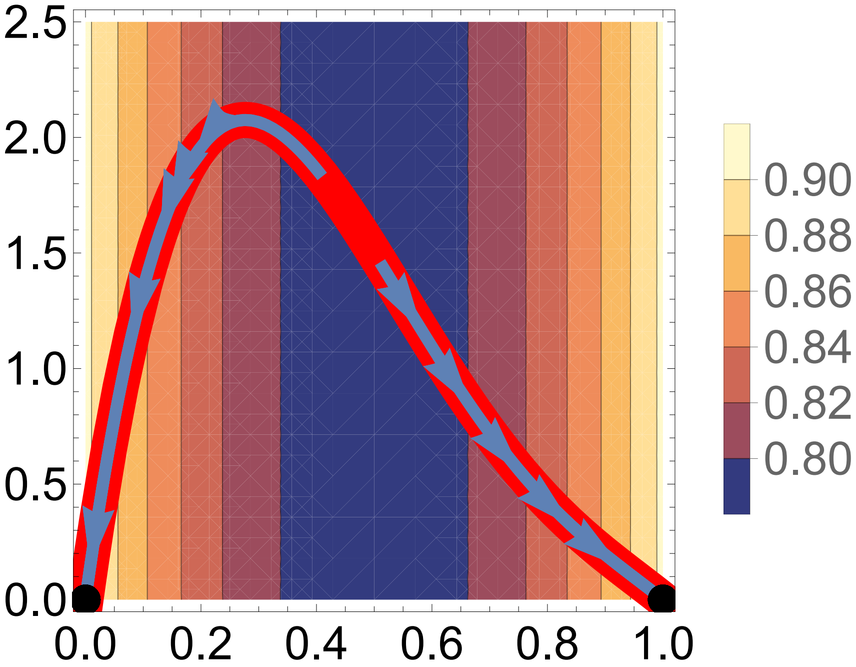

#### Example 3 continued: see SI section S9.2

In[ ]:=
```
Clear[d]

(*This defines the vector of phenotype and haplotype,
also for offspring, and the mean of this vector*)
m[i_, j_] := {{z[i, j]}, {x[i]}}
mp[i_, j_] := {{zp[i, j]}, {xp[i]}}
mb = {{zb}, {xb}};

(*This is the variance of this vector*)
Pm = FullSimplify[
   Sum[p[i] × p[j] (m[i, j] - mb).Transpose[(m[i, j] - mb)], {i, 1, 2}, {j, 1, 2}] /.
      {p[1] → pa, p[2] → pA} /. {x[1] → xa, x[2] → xA} /.
    {z[1, 1] → zaa, z[1, 2] → zAa, z[2, 1] → zAa, z[2, 2] → zAA}];

(*This is mean relative fitness*)
wb = FullSimplify[
   Sum[p[i] × p[j] × w[i, j], {i, 1, 2}, {j, 1, 2}] /. {p[1] → pa, p[2] → pA} /.
      {x[1] → xa, x[2] → xA} /. {z[1, 1] → zaa, z[1, 2] → zAa,
      z[2, 1] → zAa, z[2, 2] → zAA} /. w[2, 1] → w[1, 2]];

(*This is the covariance vector of this vector with fitness*)
covmw =
  FullSimplify[Sum[p[i] × p[j] (m[i, j] - mb) (w[i, j] - wb), {i, 1, 2}, {j, 1, 2}] /.
       {p[1] → pa, p[2] → pA} /. {x[1] → xa, x[2] → xA} /. {z[1, 1] → zaa,
      z[1, 2] → zAa, z[2, 1] → zAa, z[2, 2] → zAA} /. w[2, 1] → w[1, 2]];

(*This is the mean m prime*)
mpb = FullSimplify[
   Sum[p[i] × p[j] × mp[i, j], {i, 1, 2}, {j, 1, 2}] /. {p[1] → pa, p[2] → pA} /.
       {x[1] → xa, x[2] → xA} /. {z[1, 1] → zaa, z[1, 2] → zAa,
       z[2, 1] → zAa, z[2, 2] → zAA} /. {zp[1, 1] → zaap, zp[1, 2] → zAap,
      zp[2, 1] → zAap, zp[2, 2] → zAAp} /. {xp[1] → xap, xp[2] → xAp}];

(*This is the transmission matrix of this vector*)
Tm = FullSimplify[
   Sum[p[i] × p[j] (mp[i, j] - mpb).Transpose[(m[i, j] - mb)], {i, 1, 2}, {j, 1, 2}] /.
        {p[1] → pa, p[2] → pA} /. {x[1] → xa, x[2] → xA} /.
      {z[1, 1] → zaa, z[1, 2] → zAa, z[2, 1] → zAa, z[2, 2] → zAA} /.
     {zp[1, 1] → zaap, zp[1, 2] → zAap, zp[2, 1] → zAap, zp[2, 2] → zAAp} /.
    {xp[1] → xap, xp[2] → xAp}];
```

In[∘]:=
```
(*This checks the form of Tm*)
Simplify[Tm == Px[xb] {{Dd[xb, d]^2, Hx Dd[xb, d]}, {Hx Dd[xb, d], Hx}}]

MatrixForm[{{Px Dd^2, Tx Dd}, {Tx Dd, Tx}}]
```

Out[∘]=

True

Out[∘]//MatrixForm=

$$\begin{pmatrix} Dd^2\,Px & Dd\,Tx \\ Dd\,Tx & Tx \end{pmatrix}$$

In[∘]:=
```
(*This finds the determinant and
 eigenvalues of Tm and plots the eigenvalues*)
Simplify[Det[Px {{Dd^2, Hx Dd}, {Hx Dd, Hx}}] == Px^2 Dd^2 / 4]

Simplify[Eigenvalues[Px {{Dd^2, Hx Dd}, {Hx Dd, Hx}}]]

Plot3D[{-1/4 (-1 - 2 Dd^2 + Sqrt[1 + 4 Dd^4]) Px, 1/4 (1 + 2 Dd^2 + Sqrt[1 + 4 Dd^4]) Px},
 {Dd, -1, 1}, {Px, 0, 1/4}]
```

Out[∘]=

True

Out[∘]=

$$\left\{-\frac{1}{4}\left(-1-2\,Dd^2+\sqrt{1+4\,Dd^4}\right)Px,\ \frac{1}{4}\left(1+2\,Dd^2+\sqrt{1+4\,Dd^4}\right)Px\right\}$$

Out[∘]=

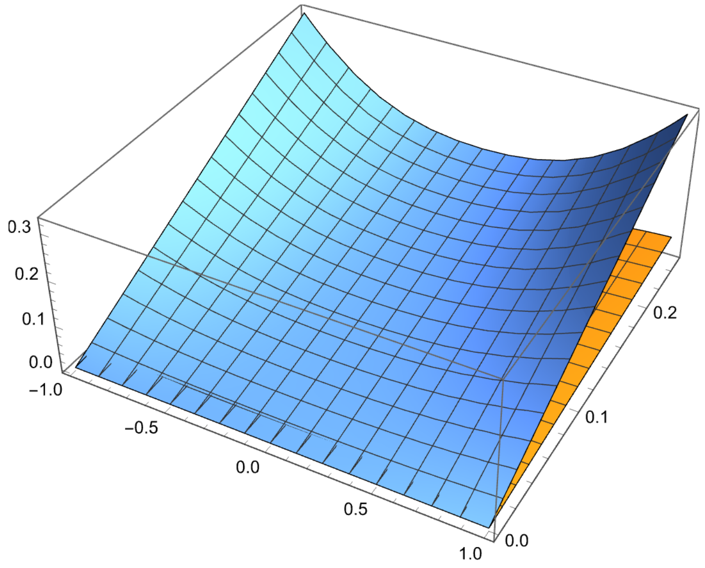

In[ ]:=
```
(*This checks the form of Pm*)
Simplify[Pm == {{P[xb], Px[xb] × Dd[xb, d]}, {Px[xb] × Dd[xb, d], Px[xb]}}]

MatrixForm[{{P, Px Dd}, {Px Dd, Px}}]

Simplify[Det[{{2 Px Dd^2 + (2 Px c d)^2, Px Dd}, {Px Dd, Px}}]]
```

Out[ ]=

True

Out[ ]//MatrixForm=

$$\begin{pmatrix} P & Dd\ Px \\ Dd\ Px & Px \end{pmatrix}$$

Out[ ]=

$$Px^2\ \left(Dd^2 + 4\ d^2\ Px\right)$$

In[ ]:=
```
(*This is Hm*)

Hm = Simplify[Tm.Inverse[Pm]];
MatrixForm[Hm]
```

$$\text{Simplify}\left[Hm == \left\{\left\{\frac{Dd[xb, d]^2}{2\ c^2\ \left((1+d)^2 - 4\ d\ xb\right)}, \frac{2\ d^2\ Px[xb] \times Dd[xb, d]}{\left((1+d)^2 - 4\ d\ xb\right)}\right\}, \{0, Hx\}\right\}\right]$$

```
MatrixForm[Hm /. d → 0]
```

Out[ ]//MatrixForm=

$$\begin{pmatrix} \frac{(-1+d\ (-1+2\ xb))^2}{2\ \left(1+d^2+d\ (2-4\ xb)\right)} & \frac{2\ d^2\ (-1+xb)\ xb\ (-1+d\ (-1+2\ xb))}{1+d^2+d\ (2-4\ xb)} \\ 0 & \frac{1}{2} \end{pmatrix}$$

Out[ ]=

True

Out[ ]//MatrixForm=

$$\begin{pmatrix} \frac{1}{2} & 0 \\ 0 & \frac{1}{2} \end{pmatrix}$$

In[ ]:= (*This plots the two top entries of Hm showing they can be negative*)

$$\text{Manipulate}\left[\text{Plot3D}\left[\left\{\frac{(1+d\,(1-2\,xb))^2}{2\,\left((1+d)^2-4\,d\,xb\right)},\ -\frac{2\,c\,d^2\,(1+d\,(1-2\,xb))\,(-1+xb)\,xb}{(1+d)^2-4\,d\,xb}\right\},\ \{xb,0,1\},\{d,-2,2\}\right],\{c,1,2\}\right]$$

Out[ ]=

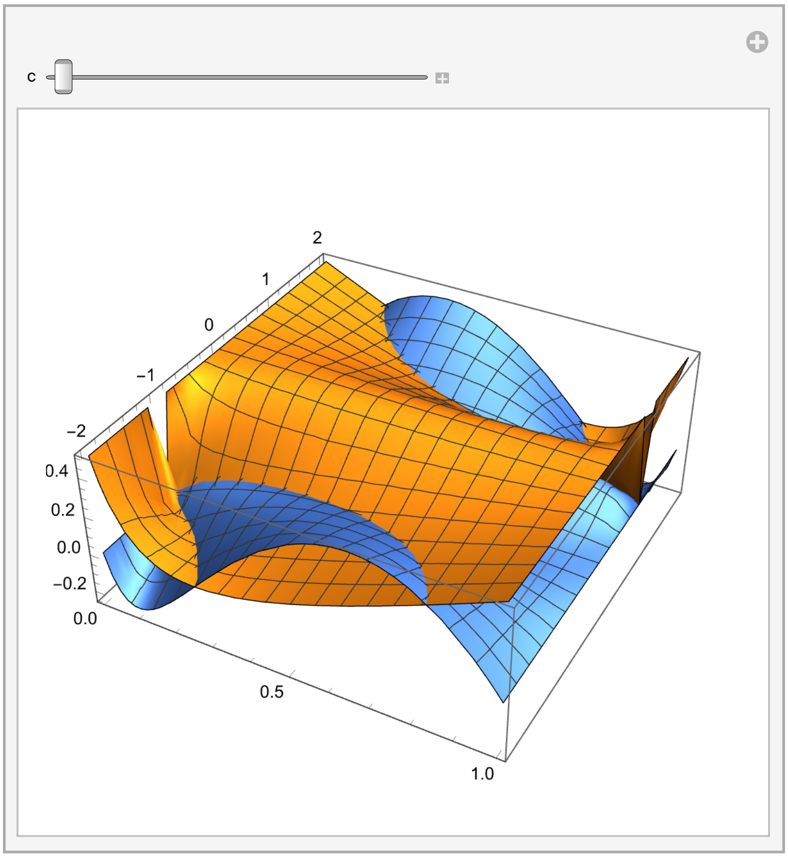

In[ ]:= (*This is βm*)

```
betam = FullSimplify[Inverse[Pm].covmw];
MatrixForm[betam]
MatrixForm[betam /. d → 0]
```

Out[ ]//MatrixForm=

$$\begin{pmatrix} \frac{(1+d)\,(-1+xb)\,w[1,1]+(1+d-2\,xb)\,w[1,2]-(-1+d)\,xb\,w[2,2]}{(1+d)^2-4\,d\,xb} \\ \frac{2\,d\,(-1+xb)\,xb\,((-1+d)\,w[1,1]+2\,w[1,2]-(1+d)\,w[2,2])}{(1+d)^2-4\,d\,xb} \end{pmatrix}$$

Out[ ]//MatrixForm=

$$\begin{pmatrix} (-1+xb)\,w[1,1]+(1-2\,xb)\,w[1,2]+xb\,w[2,2] \\ 0 \end{pmatrix}$$

In[ ]:=

```
(*This is cov[w,x']*)
covwx[xvar_] :=
 Simplify[Sum[p[i] × p[j] (x[i] - xb) (w[i, j] - 1), {i, 1, 2}, {j, 1, 2}] /. w[2, 1] →
       w[1, 2] /. {x[1] → xa, x[2] → xA} /. {p[1] → pa, p[2] → pA}] /. {xb → xvar}

(*This is betax'*)
betax[xb_] := covwx[xb] / Px[xb]

(*This checks that the top entry of betam is betax if d=0 and c=1*)
Simplify[
 betax[xb] == betam[[1, 1]] /. {w[1, 1] → w[1, 1, xb, θ, σ], w[1, 2] → w[1, 2, xb, θ, σ],
    w[2, 1] → w[2, 1, xb, θ, σ], w[2, 2] → w[2, 2, xb, θ, σ]} /. {d → 0, c → 1}]
```

Out[ ]=

```
True
```

In[ ]:=

```
(*This is cov[z,w]*)
covzw =
  FullSimplify[Sum[p[i] × p[j] (z[i, j] - zb) (w[i, j] - wb), {i, 1, 2}, {j, 1, 2}] /.
      {p[1] → pa, p[2] → pA} /. {x[1] → xa, x[2] → xA} /. {z[1, 1] → zaa,
     z[1, 2] → zAa, z[2, 1] → zAa, z[2, 2] → zAA} /. w[2, 1] → w[1, 2]];

(*This is beta*)
beta = FullSimplify[covzw / P[xb]];

(*This checks that the top entry of betam is beta if d=0*)
Simplify[
 beta == betam[[1, 1]] /. {w[1, 1] → w[1, 1, xb, θ, σ], w[1, 2] → w[1, 2, xb, θ, σ],
    w[2, 1] → w[2, 1, xb, θ, σ], w[2, 2] → w[2, 2, xb, θ, σ]} /. d → 0]
```

Out[ ]=

```
True
```

In[ ]:=

```
(*This checks that betax=D betam[[1,1]]+betam[[2,1]]*)
Simplify[betax[xb] == Dd[xb, d] × betam[[1, 1]] + betam[[2, 1]]]
```

Out[ ]=

```
True
```

In[•]:=
```
(*This is cov[m',w]*)

covmpw =
  FullSimplify[Sum[p[i] × p[j] (mp[i, j] - mb) (w[i, j] - wb), {i, 1, 2}, {j, 1, 2}] /.
        {p[1] → pa, p[2] → pA} /. {x[1] → xa, x[2] → xA} /. {z[1, 1] → zaa,
       z[1, 2] → zAa, z[2, 1] → zAa, z[2, 2] → zAA} /. w[2, 1] → w[1, 2] /.
     {zp[1, 1] → zaap, zp[1, 2] → zAap, zp[2, 1] → zAap, zp[2, 2] → zAAp} /.
    {xp[1] → xap, xp[2] → xAp}];
MatrixForm[covmpw]
```

Out[•]//MatrixForm=

$$\begin{pmatrix} (-1+xb)\ xb\ (-1+d\ (-1+2\ xb))\ ((-1+xb)\ w[1,1]+(1-2\ xb)\ w[1,2]+xb\ w[2,2]) \\ -\frac{1}{2}\ (-1+xb)\ xb\ ((-1+xb)\ w[1,1]+(1-2\ xb)\ w[1,2]+xb\ w[2,2]) \end{pmatrix}$$

In[•]:=
```
Simplify[covmpw == Px[xb] {{Dd[xb, d]}, {Hx}} betax[xb] /.
  {w[1, 1] → w[1, 1, xb, θ, σ], w[1, 2] → w[1, 2, xb, θ, σ],
   w[2, 1] → w[2, 1, xb, θ, σ], w[2, 2] → w[2, 2, xb, θ, σ]}]
```

Out[•]=
True

In[•]:=
```
(*This is um*)

um = FullSimplify[covmpw - Hm.covmw];

MatrixForm[um]

MatrixForm[um /. d → 0]

(*This checks the form of um given in the SI*)
Simplify[
 um[[1, 1]] == -d (Px[xb]^2 Dd[xb, d])/((1 + d)^2 - 4 d xb) ((-1 + d) w[1, 1] + 2 w[1, 2] - (1 + d) w[2, 2])]
```

Out[•]//MatrixForm=

$$\begin{pmatrix} \frac{d\ (-1+xb)^2\ xb^2\ (-1+d\ (-1+2\ xb))\ ((-1+d)\ w[1,1]+2\ w[1,2]-(1+d)\ w[2,2])}{(1+d)^2-4\ d\ xb} \\ 0 \end{pmatrix}$$

Out[•]//MatrixForm=

$$\begin{pmatrix} 0 \\ 0 \end{pmatrix}$$

Out[•]=
True

In[∘]:=
```
(*This is Pm'=cov[m',m']*)

Pmp = FullSimplify[
   Sum[p[i] × p[j] × mp[i, j].Transpose[(mp[i, j] - mb)], {i, 1, 2}, {j, 1, 2}] /.
         {p[1] → pa, p[2] → pA} /. {x[1] → xa, x[2] → xA} /.
       {z[1, 1] → zaa, z[1, 2] → zAa, z[2, 1] → zAa, z[2, 2] → zAA} /.
      {zp[1, 1] → zaap, zp[1, 2] → zAap, zp[2, 1] → zAap, zp[2, 2] → zAAp} /.
     {xp[1] → xap, xp[2] → xAp}];
MatrixForm[Pmp]

(*This checks the form of Pm given in the SI*)
Simplify[Pmp == 1/2 Px[xb] {{Dd[xb, d]^2, Hx Dd[xb, d]}, {Hx Dd[xb, d], Hx}}]
```

Out[∘]//MatrixForm=

$$\begin{pmatrix} -\frac{1}{2}\ (-1+\text{xb})\ \text{xb}\ (1+\text{d}-2\ \text{d}\ \text{xb})^2 & \frac{1}{4}\ (-1+\text{xb})\ \text{xb}\ (-1+\text{d}\ (-1+2\ \text{xb})) \\ \frac{1}{4}\ (-1+\text{xb})\ \text{xb}\ (-1+\text{d}\ (-1+2\ \text{xb})) & -\frac{1}{4}\ (-1+\text{xb})\ \text{xb} \end{pmatrix}$$

Out[∘]=

True

In[∘]:=
```
(*This finds the determinant of Pmp,
showing that it is invertible if D and Px are non-zero*)
Det[1/2 Px {{Dd^2, 1/2 Dd}, {1/2 Dd, 1/2}}]
```

Out[∘]=

$$\frac{\text{Dd}^2\ \text{Px}^2}{16}$$

In[∘]:=
```
(*This is betam'*)

betamp = Simplify[Inverse[Pmp].covmpw];
MatrixForm[betamp]

Simplify[
 betamp[[1, 1]] == 2 (Sum[p[j] × w[2, j], {j, 1, 2}] - Sum[p[j] × w[1, j], {j, 1, 2}]) / Dd[xb, d] /.
   {w[2, 1] → w[1, 2]} /. {p[1] → pa, p[2] → pA}]
```

Out[∘]//MatrixForm=

$$\begin{pmatrix} -\frac{2\ ((-1+\text{xb})\ w[1,1]+(1-2\ \text{xb})\ w[1,2]+\text{xb}\ w[2,2])}{-1+\text{d}\ (-1+2\ \text{xb})} \\ 0 \end{pmatrix}$$

Out[∘]=

True

In[∘]:=
```
(*This checks that the dot product of betam and betamp can be negative,
so they can be at an angle greater than 90 degrees*)

FullSimplify[Transpose[betamp].betam]

Manipulate[Plot3D[
  -((2 ((-1 + xb) w11 + (1 - 2 xb) w12 + xb w22) ((1 + d) (-1 + xb) w11 + (1 + d - 2 xb) w12 -
        (-1 + d) xb w22)) / (c^2 ((1 + d)^2 - 4 d xb) (-1 + d (-1 + 2 xb)))),
  {xb, 0, 1}, {d, -2, 2}, PlotRange → {0, 1}], {{w11, 0.01}, 0, 1},
 {{w12, 0.12}, 0, 1}, {{w22, 0.1}, 0, 1},
 {c, 1, 2}]
```

Out[∘]=
```
{{-((2 ((-1 + xb) w[1, 1] + (1 - 2 xb) w[1, 2] + xb w[2, 2])
      ((1 + d) (-1 + xb) w[1, 1] + (1 + d - 2 xb) w[1, 2] - (-1 + d) xb w[2, 2])) /
    (((1 + d)^2 - 4 d xb) (-1 + d (-1 + 2 xb))))}}
```

Out[∘]=

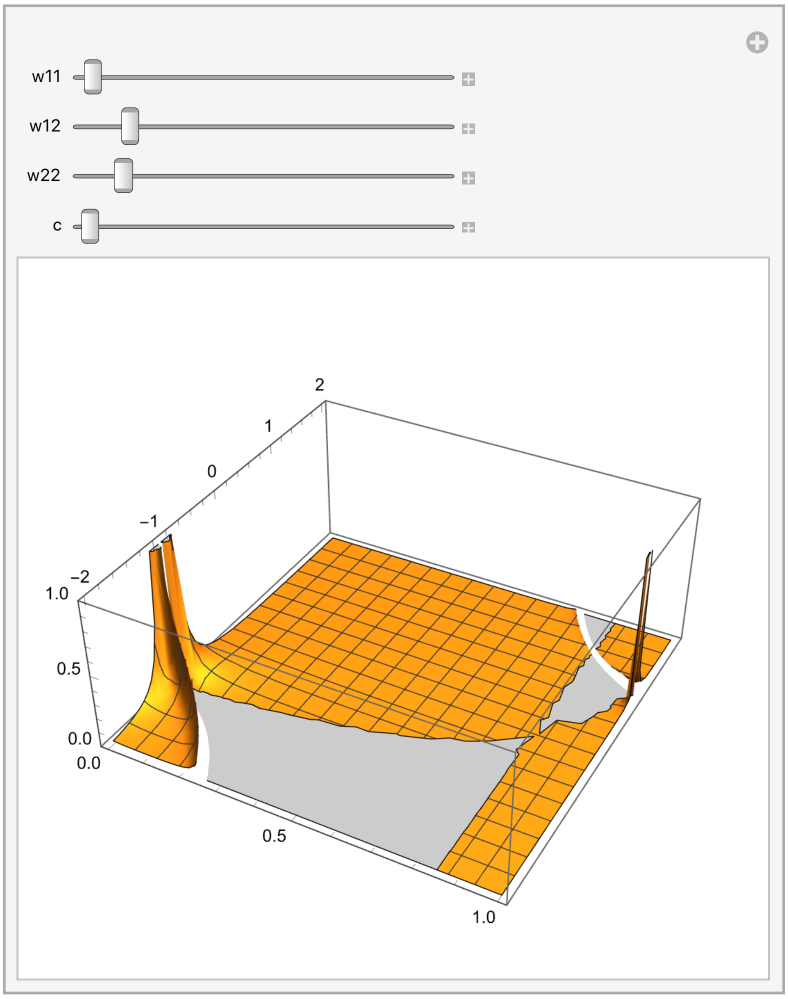

#### Example 3 approximated evo-devo dynamics: one bialellic locus, one phenotype

In[ ]:=
```
Clear["Global`*"]
```

```
(*This defines the genotype-
 phenotype map and the Tx matrix previously calculated*)
z[xk_, xl_] := c (1 + d) (xk + xl) - 2 c d xk xl
Tx[xb_] := 1/2 xb (1 - xb)

(*This defines the indicated derivatives and
 the mechanistic additive genetic covariance Lz*)
dwdx[xb_] := dzdx[xb] × dwdz[z[xb, xb]]

(*This is ∂w/∂z evaluated at the mean haplotype content*)
dwdz[zb_] := 2/σ^2 (θ - zb)

(*This dz/dx evaluated at the mean haplotype content*)
dzdx[xb_] := c (1 + d (1 - 2 xb))

(*This is the MAG covariance of the phenotype*)
Lz[xb_] := 2 dzdx[xb]^2 Tx[xb]
```

In[ ]:=
```
(*This checks that D=dz/dx*)
Simplify[c (1 + d (1 - 2 xb)) == D[z[xk, xl], xk] /. {xl → xb}]

(*This checks that the approximated
 mean phenotype equals the exact mean phenotype*)
Simplify[z[xb, xb] == 2 c xb (1 + d - d xb)]
```

Out[ ]=
```
True
```

Out[ ]=
```
True
```

In[ ]:=
```
(*This finds the allele frequency under which
 the average effect of allelic substitution is zero*)
Solve[c (1 + d) - 2 c d xb == 0, xb]

(*This finds the allele frequency under which the mean phenotype is optimum*)
Simplify[Solve[z[xb, xb] == θ, xb]]

(*This uses L'Hopital's rule to find the allele
  frequency under which the mean phenotype is optimum if d=0 *)

FullSimplify[D[c + c d - Sqrt[c (c (1 + d)^2 - 2 d θ)], d] / D[2 c d, d]] /. d → 0
```

Out[ ]=

$$\left\{\left\{xb \to \frac{1+d}{2\,d}\right\}\right\}$$

Out[ ]=

$$\left\{\left\{xb \to \frac{c + c\,d - \sqrt{c\,\left(c\,(1+d)^2 - 2\,d\,\theta\right)}}{2\,c\,d}\right\}, \left\{xb \to \frac{c + c\,d + \sqrt{c\,\left(c\,(1+d)^2 - 2\,d\,\theta\right)}}{2\,c\,d}\right\}\right\}$$

Out[ ]=

$$\frac{1}{2} + \frac{-c + \theta}{2\,\sqrt{c^2}}$$

In[ ]:=
```
(*This is Lm*)
Lm[xb_] = {{2 dzdx[xb]^2 Tx[xb], 2 dzdx[xb] × Tx[xb]}, {dzdx[xb] × Tx[xb], Tx[xb]}};

(*These are the eigenvalues of Lm*)
Eigenvalues[{{2 dzdx^2 Tx, 2 dzdx Tx}, {dzdx Tx, Tx}}]
```

Out[ ]=

$$\left\{0,\ \left(1 + 2\,dzdx^2\right)\,Tx\right\}$$

In[ ]:=
```
(*This checks that Lm is asymmetric*)
Simplify[Transpose[Lm[xb]] == Lm[xb]]
Simplify[Transpose[Lm[xb]] == Lm[xb] /. d → 0]
```

Out[ ]=

$$\left\{\left\{0, -\frac{1}{2}\,c\,(-1+xb)\,xb\,(-1+d\,(-1+2\,xb))\right\}, \left\{\frac{1}{2}\,c\,(-1+xb)\,xb\,(-1+d\,(-1+2\,xb)),\ 0\right\}\right\} == \{\{0, 0\}, \{0, 0\}\}$$

Out[ ]=

$$\left\{\left\{0, \frac{1}{2}\,c\,(-1+xb)\,xb\right\}, \left\{-\frac{1}{2}\,c\,(-1+xb)\,xb,\ 0\right\}\right\} == \{\{0, 0\}, \{0, 0\}\}$$

In[ ]:=
```
(*This is Lm is general*)
LmG = {{2 dzdx^2 Tx, 2 dzdx Tx}, {dzdx Tx, Tx}};

(*This is the symmetric part of Lm in general*)
LmsG = 1/2 (LmG + Transpose[LmG]);

(*These are the eigenvalues of the symmetric part of Lm*)
Simplify[Eigenvalues[LmsG]]
```

Out[ ]=

$$\left\{-\frac{1}{2}\left(-1-2\,\text{dzdx}^2+\sqrt{1+5\,\text{dzdx}^2+4\,\text{dzdx}^4}\right)\text{Tx},\ \frac{1}{2}\left(1+2\,\text{dzdx}^2+\sqrt{1+5\,\text{dzdx}^2+4\,\text{dzdx}^4}\right)\text{Tx}\right\}$$

In[ ]:=
```
(*This plots the eigenvalues of LmsG/Tx,
showing that one of them can be negative*)

Plot[{-1/2 (-1 - 2 dzdx^2 + Sqrt[1 + 5 dzdx^2 + 4 dzdx^4]),
  1/2 (1 + 2 dzdx^2 + Sqrt[1 + 5 dzdx^2 + 4 dzdx^4])}, {dzdx, -1, 1}]
```

Out[ ]=

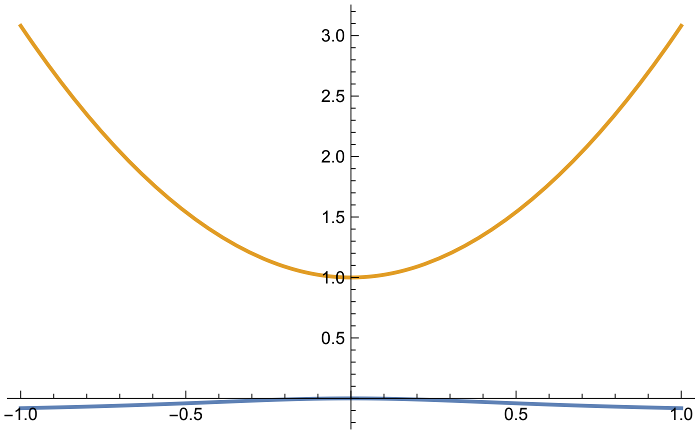

```
(*This is fitness*)
W[z_, θ_, σ_] := Exp[-(z - θ)^2 / σ^2]

(*This runs the numerical solutions for the indicated parameter values*)

c = 1;
d = 2;

θ = 1;
σ = Sqrt[10];

Plot1FileName =
  If[θ == 1.5 && d == 0, "p", If[θ == 1 && d == 0, "u", If[θ == 1 && d == 2, "z"]]];
Plot2FileName =
  If[θ == 1.5 && d == 0, "q", If[θ == 1 && d == 0, "v", If[θ == 1 && d == 2, "aa"]]];
Plot3FileName =
  If[θ == 1.5 && d == 0, "r", If[θ == 1 && d == 0, "w", If[θ == 1 && d == 2, "ab"]]];

tend = 100;

x1init = {0.01, 1, .1};

InitialConditions = Table[{x1b[0, i] = i}, {i, x1init[[1]], x1init[[2]], x1init[[3]]}];

Table[Table[{x1b[t + 1, i] = x1b[t, i] + (Tx[x1b[t, i]] × dwdx[x1b[t, i]])},
   {t, 0, tend}], {i, x1init[[1]], x1init[[2]], x1init[[3]]}];
```

```
(*This plots allele frequency, mean phenotype,
Lz, and mean fitness over time*)

(*These are the axes limits*)
yAxisLimitz[d_] := If[d ≥ 2, 2.5, 2]
yAxisLimitP[d_] := If[d ≥ 2, 0.9, .3]

lThickness = 0.02; (*Line thickness*)

(*This is allele frequency over time*)
Show[
 Table[ListLinePlot[{Table[x1b[t, i], {t, 0, tend}]}, PlotRange → {-0.1, 1.1},
   TicksStyle → Large, PlotStyle → Thickness[lThickness]],
  {i, x1init⟦1⟧, x1init⟦2⟧, x1init⟦3⟧}]]
Export[
  StringJoin[ToString[NotebookDirectory[]], "Fig.3", Plot1FileName, ".pdf"], %];

(*This is mean phenotype over time*)
Show[Table[ListLinePlot[
   {Table[z[x1b[t, i], x1b[t, i]], {t, 0, tend}], Table[θ, {t, 0, tend}]},
   PlotRange → {-0.1, yAxisLimitz[d]}, TicksStyle → Large,
   PlotStyle → {Thickness[lThickness], {Magenta, Thickness[lThickness]}}],
  {i, x1init⟦1⟧, x1init⟦2⟧, x1init⟦3⟧}]]
Export[
  StringJoin[ToString[NotebookDirectory[]], "Fig.3", Plot2FileName, ".pdf"], %];

(*This is Lz over time*)
Show[Table[ListLinePlot[{Table[Lz[x1b[t, i]], {t, 0, tend}]},
   PlotRange → {-0.1 yAxisLimitP[d], yAxisLimitP[d]}, TicksStyle → Large,
   PlotStyle → Thickness[lThickness]], {i, x1init⟦1⟧, x1init⟦2⟧, x1init⟦3⟧}]]
Export[
  StringJoin[ToString[NotebookDirectory[]], "Fig.3", Plot3FileName, ".pdf"], %];

(*This is mean fitness over time*)
Show[Table[ListLinePlot[{Table[W[z[x1b[t, i], x1b[t, i]], θ, σ], {t, 0, tend}]},
   PlotRange → {-0.1, 1.1}, TicksStyle → Large,
   PlotStyle → Thickness[lThickness]], {i, x1init⟦1⟧, x1init⟦2⟧, x1init⟦3⟧}]]
```

Out[ ]=

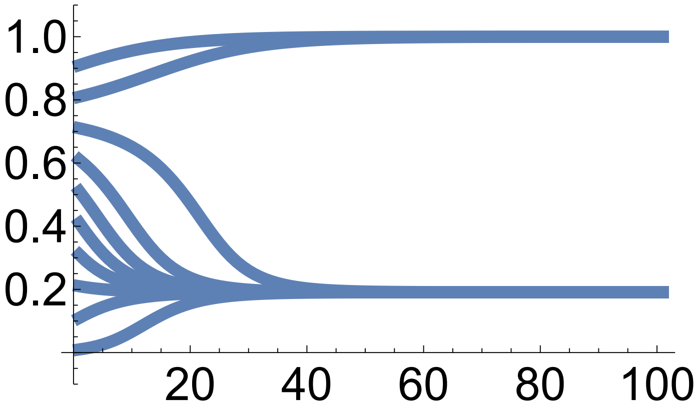

Out[ ]=

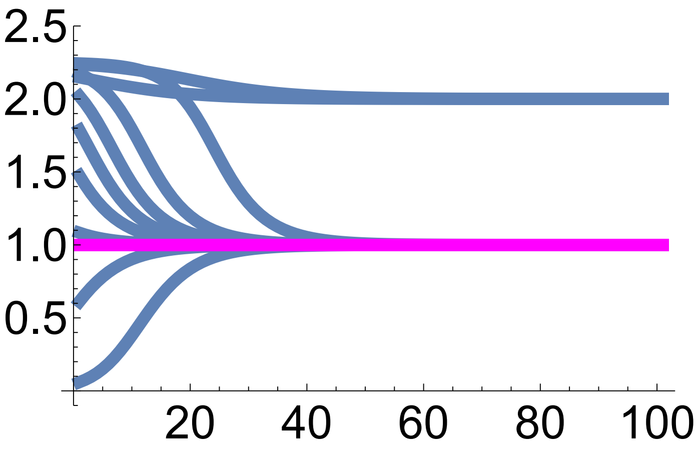

Out[ ]=

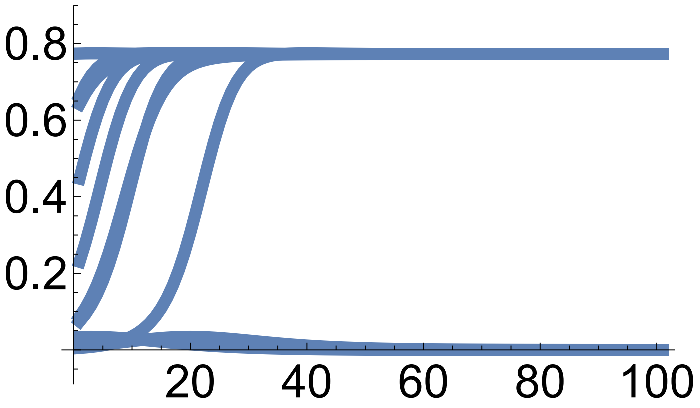

Out[ ]=

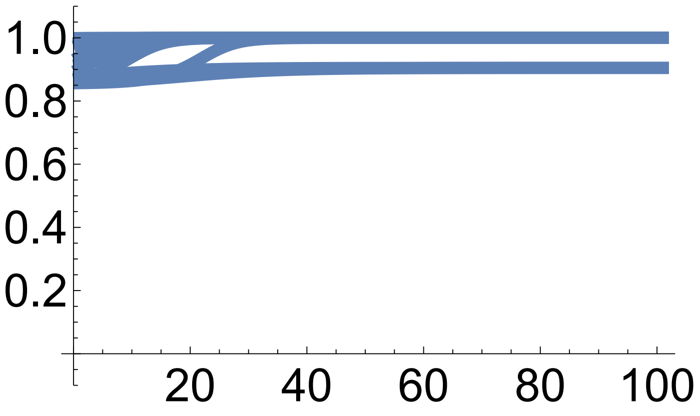

In[ ]:=
```
(*This plots the trajectories of mean phenotypes and Lz vs allele frequency*)
```

```
pointSize = 0.05;

Plot4FileName =
  If[θ == 1.5 && d == 0, "s", If[θ == 1 && d == 0, "x", If[θ == 1 && d == 2, "ac"]]];
Plot5FileName =
  If[θ == 1.5 && d == 0, "t", If[θ == 1 && d == 0, "y", If[θ == 1 && d == 2, "ad"]]];

(*These are the end points for the trajectories
 of allele frequencies and mean phenotypes*)
endPoints = Table[{x1b[tend, i], z[x1b[tend, i], x1b[tend, i]]},
   {i, x1init[[1]], x1init[[2]], x1init[[3]]}];

(*This plots the trajectories of mean phenotype vs allele frequency*)
p0 = Table[ListLinePlot[Table[{x1b[t, i], z[x1b[t, i], x1b[t, i]]}, {t, 0, tend}],
    PlotRange → {{0, 1}, {0, yAxisLimitz[d]}},
    PlotStyle → {Thickness[lThickness]}], {i, x1init[[1]], x1init[[2]], x1init[[3]]}];

(*This superimposes a contour of mean fitness and the final points*)
plot =  Show[ContourPlot[W[zb, θ, σ], {xb, 0, 1},
   {zb, 0, yAxisLimitz[d]}, FrameStyle → Large, PlotLegends → Automatic,
   LabelStyle → Large, Epilog → {PointSize[pointSize], Point[endPoints]}],
  Plot[z[x1b, x1b], {x1b, 0, 1}, PlotStyle → {Red, Thickness[3 lThickness]}],
  p0 /. Line[x_] :→ {Arrowheads[Table[{.1, .2}, {3}]], Arrow[x]}, ListLinePlot[
   Table[{xb, θ}, {xb, 0, 1}], PlotStyle → {Magenta, Thickness[lThickness]}]]

plot = Rasterize[plot, "Image", ImageResolution → 300];

Export[StringJoin[ToString[NotebookDirectory[]],
   "Fig.3", Plot4FileName, ".pdf"], plot];

(*These are the end points of the trajectories of
 allele frequency and MAG covariance of the phenotype *)
endPointsxL =
  Table[{x1b[tend, i], Lz[x1b[tend, i]]}, {i, x1init[[1]], x1init[[2]], x1init[[3]]}];

(*This plots the trajectories of the MAG covariance vs allele frequency*)
p0 = Table[ListLinePlot[Table[{x1b[t, i], Lz[x1b[t, i]]}, {t, 0, tend}],
    PlotRange → {{0, 1}, {0, yAxisLimitP[d]}},
    PlotStyle → {Thickness[lThickness]}], {i, x1init[[1]], x1init[[2]], x1init[[3]]}];

(*This superimposes a countour of mean fitness and the end points*)
plot = Show[ContourPlot[W[z[xb, xb], θ, σ], {xb, 0, 1},
   {P, 0, yAxisLimitP[d]}, FrameStyle → Large, PlotLegends → Automatic,
   LabelStyle → Large, Epilog → {PointSize[pointSize], Point[endPointsxL]}],
  Plot[Lz[x1b], {x1b, 0, 1}, PlotStyle → {Red, Thickness[3 lThickness]}],
  p0 /. Line[x_] :→ {Arrowheads[Table[{.1, .2}, {3}]], Arrow[x]}]
```

```
plot = Rasterize[plot, "Image", ImageResolution → 300];

Export[StringJoin[ToString[NotebookDirectory[]],
   "Fig.3", Plot5FileName, ".pdf"], plot];
```

Out[ ]=

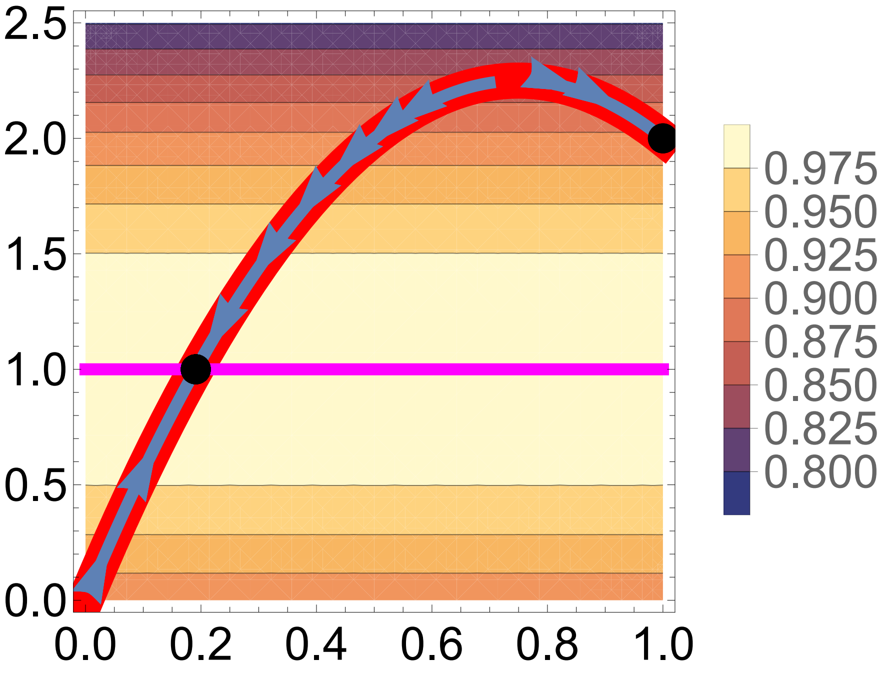

Out[ ]=

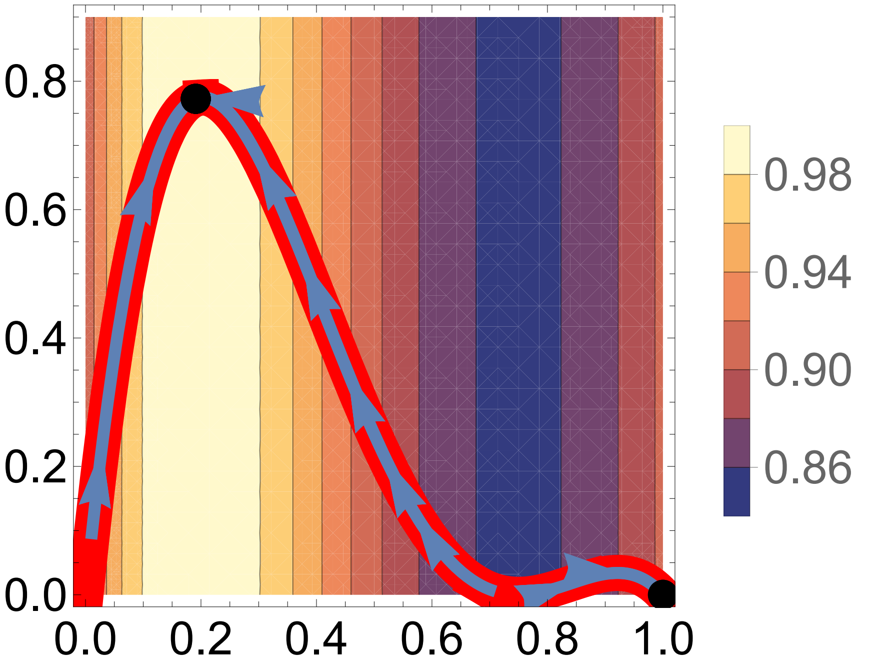

#### Fig. 4

In[ ]:=
```
(*This runs the numerical solutions*)
Clear[d, θ]

d = 0;
θ = 1.5;

(*This defines the change in m as Lm dw/dz,
to be used for numerical solutions,
where Lm is singular due to the coupling of gene content and phenotype*)
Deltam[xbvar_, zbvar_] = Lm[xb].{{dwdz[zb]}, {0}} /. {xb → xbvar, zb → zbvar};

tend = 100;

x1init = {0.01, 1, .1};

(*This specifies the initial conditions given by the original genotype-
 phenotype map plus random noise*)
InitialConditions =
  Table[{x1b[0, i] = i, zb[0, i] = z[x1b[0, i], x1b[0, i]] + RandomReal[{-.5, .5}]},
   {i, x1init⟦1⟧, x1init⟦2⟧, x1init⟦3⟧}];

(*Numerical solution*)
Table[Table[{x1b[t + 1, i] = x1b[t, i] + Deltam[x1b[t, i], zb[t, i]]⟦2, 1⟧,
     zb[t + 1, i] = zb[t, i] + Deltam[x1b[t, i], zb[t, i]]⟦1, 1⟧},
   {t, 0, tend}], {i, x1init⟦1⟧, x1init⟦2⟧, x1init⟦3⟧}];
```

```
(*This plots the resulting trajectories over time*)

yAxisLimitz[d_] := If[d ≥ 2, 2.5, 2]
yAxisLimitP[d_] := If[d ≥ 2, 0.9, .3]

lThickness = 0.02; (*Line thickness*)

(*This is allele frequency over time*)
Show[
 Table[ListLinePlot[{Table[x1b[t, i], {t, 0, tend}]}, PlotRange → {-0.1, 1.1},
   TicksStyle → Large, PlotStyle → Thickness[lThickness]],
  {i, x1init⟦1⟧, x1init⟦2⟧, x1init⟦3⟧}]]
Export[StringJoin[ToString[NotebookDirectory[]], "Fig.S1a", ".pdf"], %];

(*This is mean phenotype over time*)
Show[Table[ListLinePlot[{Table[zb[t, i], {t, 0, tend}], Table[θ, {t, 0, tend}]},
   PlotRange → {-0.1, yAxisLimitz[d]}, TicksStyle → Large,
   PlotStyle → {Thickness[lThickness], {Magenta, Thickness[lThickness]}}],
  {i, x1init⟦1⟧, x1init⟦2⟧, x1init⟦3⟧}]]
Export[StringJoin[ToString[NotebookDirectory[]], "Fig.S1b", ".pdf"], %];

(*This is MAG covariance over time*)
Show[Table[ListLinePlot[{Table[Lz[x1b[t, i]], {t, 0, tend}]},
   PlotRange → {-0.1 yAxisLimitP[d], yAxisLimitP[d]}, TicksStyle → Large,
   PlotStyle → Thickness[lThickness]], {i, x1init⟦1⟧, x1init⟦2⟧, x1init⟦3⟧}]]
Export[StringJoin[ToString[NotebookDirectory[]], "Fig.S1c", ".pdf"], %];
```

Out[ ]=

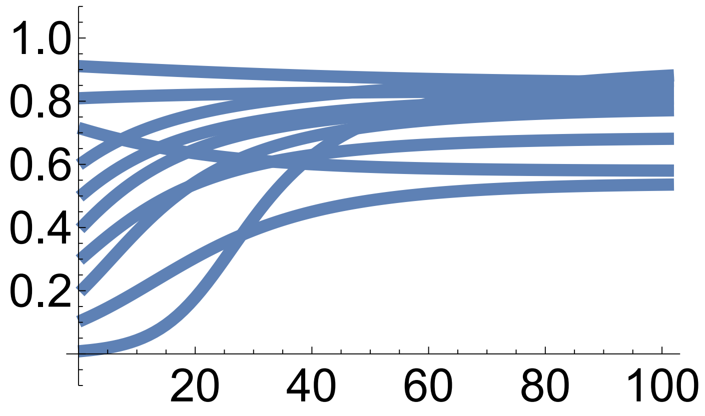

Out[ ]=

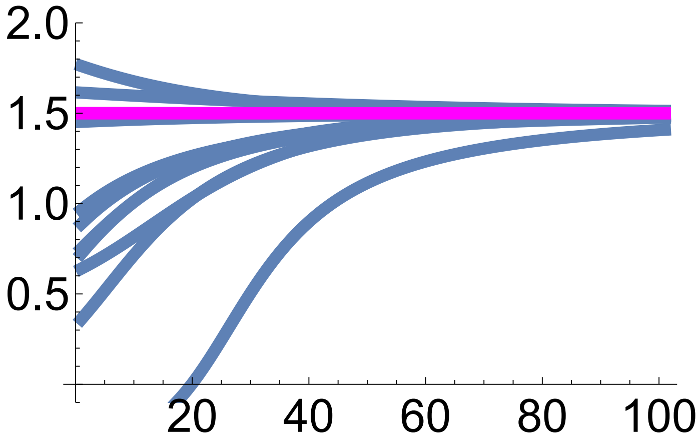

Out[ ]=

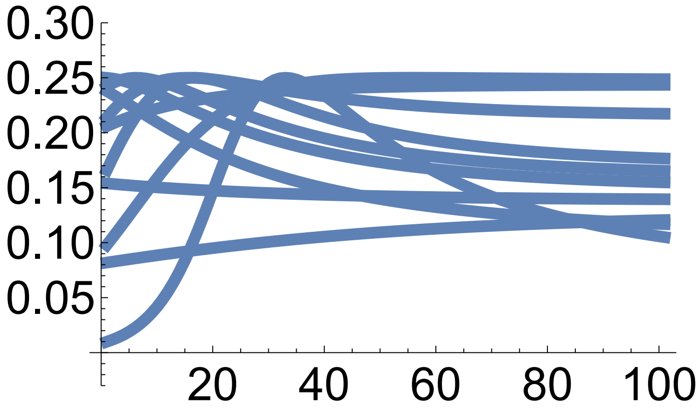

```
In[•]:= (*This plots the resulting phase diagrams*)

pointSize = 0.05;

(*This is for mean phenotype versus allele frequency*)

endPoints =
  Table[{x1b[tend, i], zb[tend, i]}, {i, x1init⟦1⟧, x1init⟦2⟧, x1init⟦3⟧}];

p0 = Table[ListLinePlot[Table[{x1b[t, i], zb[t, i]}, {t, 0, tend}],
    PlotRange → {{0, 1}, {0, yAxisLimitz[d]}},
    PlotStyle → {Thickness[lThickness]}], {i, x1init⟦1⟧, x1init⟦2⟧, x1init⟦3⟧}];

plot =  Show[ContourPlot[W[zb, θ, σ], {xb, 0, 1},
   {zb, 0, yAxisLimitz[d]}, FrameStyle → Large, PlotLegends → Automatic,
   LabelStyle → Large, Epilog → {PointSize[pointSize], Point[endPoints]}],
  Plot[z[x1b, x1b], {x1b, 0, 1}, PlotStyle → {Red, Thickness[3 lThickness]}],
  p0 /. Line[x_] :→ {Arrowheads[Table[{.1, .2}, {3}]], Arrow[x]}, ListLinePlot[
   Table[{xb, θ}, {xb, 0, 1}], PlotStyle → {Magenta, Thickness[lThickness]}]]

plot = Rasterize[plot, "Image", ImageResolution → 300];

Export[StringJoin[ToString[NotebookDirectory[]], "Fig.S1d", ".pdf"], plot];

(*This is for MAG covariance vs allele frequency*)

endPointsxL =
  Table[{x1b[tend, i], Lz[x1b[tend, i]]}, {i, x1init⟦1⟧, x1init⟦2⟧, x1init⟦3⟧}];

p0 = Table[ListLinePlot[Table[{x1b[t, i], Lz[x1b[t, i]]}, {t, 0, tend}],
    PlotRange → {{0, 1}, {0, yAxisLimitP[d]}},
    PlotStyle → {Thickness[lThickness]}], {i, x1init⟦1⟧, x1init⟦2⟧, x1init⟦3⟧}];

plot = Show[ContourPlot[W[z[xb, xb], θ, σ](*Wbz[zb,h,θ,σ]*), {xb, 0, 1},
   {P, 0, yAxisLimitP[d]}, FrameStyle → Large, PlotLegends → Automatic,
   LabelStyle → Large, Epilog → {PointSize[pointSize], Point[endPointsxL]}],
  Plot[Lz[x1b], {x1b, 0, 1}, PlotStyle → {Red, Thickness[3 lThickness]}],
  p0 /. Line[x_] :→ {Arrowheads[Table[{.1, .2}, {3}]], Arrow[x]}]

plot = Rasterize[plot, "Image", ImageResolution → 300];

Export[StringJoin[ToString[NotebookDirectory[]], "Fig.S1e", ".pdf"], plot];
```

Out[ ]=

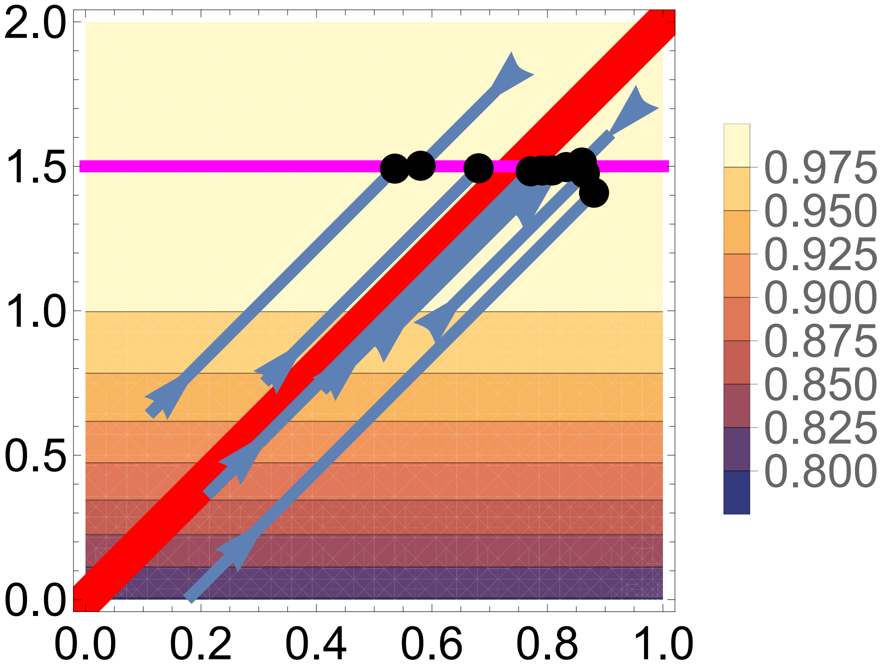
2.0
1.5
1.0
0.5
0.0
0.0
0.2
0.4
0.6
0.8
1.0
0.975
0.950
0.925
0.900
0.875
0.850
0.825
0.800

Out[ ]=

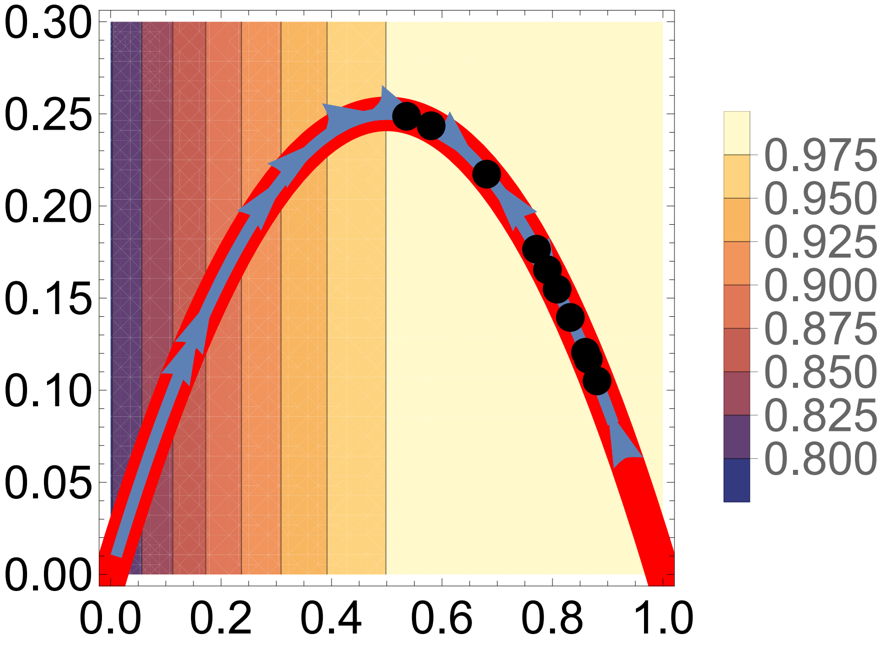
0.30
0.25
0.20
0.15
0.10
0.05
0.00
0.0
0.2
0.4
0.6
0.8
1.0
0.975
0.950
0.925
0.900
0.875
0.850
0.825
0.800

#### Example 4 exact evo-devo dynamics: two biallelic loci, two phenotypes

In[101]:=
```
Clear["Global`*"]
```

In[102]:=
```
(*This defines the gene content for locus i and allele j in haplotype k1k2*)
x[i_, j_, k1_, k2_] :=
 KroneckerDelta[ToString[j], ToString[If[i == 1, k1, If[i == 2, k2]]]]

(*This defines haplotype frequency*)
p[i_, j_, Xb_] := xb[1, i] × xb[2, j] + covx12[i, j, Xb]

(*This defines allele frequency*)
xb[1, a] = 1 - x1b;
xb[1, A] = x1b;
xb[2, b] = 1 - x2b;
xb[2, B] = x2b;

(*This defines gene content covariance between loci*)
covx12[a, b, Xb_] = Xb;
covx12[a, B, Xb_] = -Xb;
covx12[A, b, Xb_] = -Xb;
covx12[A, B, Xb_] = Xb;
```

In[112]:=
```
(*This defines xvec[i,j], which is vector of haplotype content,
listing gene content in both loci and the
 haplotype content deviation for haplotype ij*)
xvec[i_, j_] :=
  {{x[1, A, i, j]}, {x[2, B, i, j]}, {(x[1, A, i, j] - x1b) (x[2, B, i, j] - x2b)}};

(*This is the vector of mean haplotype content*)
xbvec = {{x1b}, {x2b}, {Xb}};

(*This writes the vector of mean haplotype
 content as a function of the evolving traits*)
xbFun[x1bvar_, x2bvar_, Xbvar_] =
  {{x1b}, {x2b}, {Xb}} /. {x1b → x1bvar, x2b → x2bvar, Xb → Xbvar};
```

In[ ]:=
```
Table[MatrixForm[xvec[i, j]], {i, {a, A}}, {j, {b, B}}]
```

Out[ ]=

$$\left\{\left\{\begin{pmatrix} 0 \\ 0 \\ x1b\ x2b \end{pmatrix}, \begin{pmatrix} 0 \\ 1 \\ -x1b\ (1 - x2b) \end{pmatrix}\right\}, \left\{\begin{pmatrix} 1 \\ 0 \\ -((1 - x1b)\ x2b) \end{pmatrix}, \begin{pmatrix} 1 \\ 1 \\ (1 - x1b)\ (1 - x2b) \end{pmatrix}\right\}\right\}$$

In[115]:=

```
(*This is Px, that is, cov[x,x], the covariance matrix of haplotype content*)

Px = Simplify[Sum[p[i, j, Xb] (xvec[i, j] - xbvec).Transpose[(xvec[i, j] - xbvec)],
   {i, {a, A}}, {j, {b, B}}]];
MatrixForm[Px]
PxFun[x1bvar_, x2bvar_, Xbvar_] = Px /. {x1b → x1bvar, x2b → x2bvar, Xb → Xbvar};
```

Out[115]//MatrixForm=

$$\begin{pmatrix} -((-1+x1b)\ x1b) & Xb & Xb - 2\ x1b\ Xb \\ Xb & -((-1+x2b)\ x2b) & Xb - 2\ x2b\ Xb \\ Xb - 2\ x1b\ Xb & Xb - 2\ x2b\ Xb & x1b^2\ (-1+x2b)\ x2b - Xb\ (-1+2\ x2b+Xb) + x1b\ \big(x2b - \end{pmatrix}$$

In[117]:=

```
(*This is the haplotype content deviation for the two loci,
X, its mean, and variance*)

X[k1_, k2_] := (x[1, A, k1, k2] - x1b) (x[2, B, k1, k2] - x2b)

Simplify[Sum[p[i, j, Xb] × X[i, j], {i, {a, A}}, {j, {b, B}}]]

varX = Simplify[Sum[p[i, j, Xb] (X[i, j] - Xb)^2, {i, {a, A}}, {j, {b, B}}]]

(*This checks that the bottom right entry of Px is var[X]*)
Simplify[Px[[3, 3]] == varX]
```

Out[118]=

Xb

Out[119]=

$$x1b^2\ (-1+x2b)\ x2b - Xb\ (-1+2\ x2b+Xb) + x1b\ \left(x2b - x2b^2 - 2\ Xb + 4\ x2b\ Xb\right)$$

Out[120]=

True

In[121]:=

```
(*Probability that genotype k1k2 l1l2 produces gametes with haplotype n1n2*)

R[k1_, k2_, l1_, l2_, n1_, n2_] := 1/2 KroneckerDelta[ToString[k1], ToString[n1]]
    ((1 - r) KroneckerDelta[ToString[k2], ToString[n2]] +
      r KroneckerDelta[ToString[l2], ToString[n2]]) +
   1/2 KroneckerDelta[ToString[l1], ToString[n1]]
    ((1 - r) KroneckerDelta[ToString[l2], ToString[n2]] +
      r KroneckerDelta[ToString[k2], ToString[n2]])
```

In[122]:=

```
(*This is the x' for gene content denoted here as xp[i,j,k1,k2]:
 the expected gene content in locus i for allele
  j among offspring haplotypes of haplotype k1k2
*)

xp[i_, j_, k1_, k2_, Xb_] :=
 Sum[p[l1, l2, Xb] × Sum[R[k1, k2, l1, l2, m1, m2] × x[i, j, m1, m2],
    {m1, {a, A}}, {m2, {b, B}}], {l1, {a, A}}, {l2, {b, B}}]

MatrixForm[Table[
  Simplify[{xp[1, A, k1, k2, Xb], xp[2, B, k1, k2, Xb]}], {k1, {a, A}}, {k2, {b, B}}]]
```

Out[123]//MatrixForm=

$$\begin{pmatrix} \begin{pmatrix} \frac{x1b}{2} \\ \frac{x2b}{2} \end{pmatrix} & \begin{pmatrix} \frac{x1b}{2} \\ \frac{1+x2b}{2} \end{pmatrix} \\ \begin{pmatrix} \frac{1+x1b}{2} \\ \frac{x2b}{2} \end{pmatrix} & \begin{pmatrix} \frac{1+x1b}{2} \\ \frac{1+x2b}{2} \end{pmatrix} \end{pmatrix}$$

In[124]:=

```
(*This is the X' for haplotype deviation denoted here as Xp[k1,k2,Xb]:
 the expected haplotype gene content deviation between
  loci among gametes produced by parental haplotype k1k2
*)

Xp[k1_, k2_, Xb_] :=
 Simplify[Sum[p[l1, l2, Xb] × Sum[R[k1, k2, l1, l2, m1, m2] × X[m1, m2],
     {m1, {a, A}}, {m2, {b, B}}], {l1, {a, A}}, {l2, {b, B}}]]

MatrixForm[Table[Simplify[Xp[k1, k2, Xb]], {k1, {a, A}}, {k2, {b, B}}]]
```

Out[125]//MatrixForm=

$$\begin{pmatrix} -\frac{1}{2}(-1+r)(x1b\,x2b+Xb) & -\frac{1}{2}(-1+r)(x1b(-1+x2b)+Xb) \\ -\frac{1}{2}(-1+r)((-1+x1b)\,x2b+Xb) & -\frac{1}{2}(-1+r)(1+x1b(-1+x2b)-x2b+Xb) \end{pmatrix}$$

In[ ]:=
```
(*This checks that the third entries of vectors in eq. 128 are correct*)
Simplify[Xp[a, b, Xb] == 1/2 (1 - r) p[A, B, Xb]]
Simplify[Xp[a, B, Xb] == -1/2 (1 - r) p[A, b, Xb]]
Simplify[Xp[A, b, Xb] == -1/2 (1 - r) p[a, B, Xb]]
Simplify[Xp[A, B, Xb] == 1/2 (1 - r) p[a, b, Xb]]
```
Out[ ]=
```
True
```
Out[ ]=
```
True
```
Out[ ]=
```
True
```
Out[ ]=
```
True
```
In[126]:=
```
(*This is the vector x' in the main text, denoted here as xpvec[k1,k2]:
 vector listing the expected gene content in both loci and haplotype content
  deviation between loci among offspring haplotypes of haplotype k1k2*)
xpvec[k1_, k2_, Xb_] :=
  {{xp[1, A, k1, k2, Xb]}, {xp[2, B, k1, k2, Xb]}, {Xp[k1, k2, Xb]}};

(*This is zetax = E[x']*)
zetax = Simplify[Sum[p[k1, k2, Xb] × xpvec[k1, k2, Xb], {k1, {a, A}}, {k2, {b, B}}]]
```
Out[127]=
```
{{x1b}, {x2b}, {Xb - r Xb}}
```
In[128]:=
```
(*This is Px', that is, cov[x',x']*)
Pxp[x1bvar_, x2bvar_, Xbvar_] =
  Simplify[Sum[p[k1, k2, Xb] (xpvec[k1, k2, Xb] - zetax).
       Transpose[(xpvec[k1, k2, Xb] - zetax)], {k1, {a, A}},
     {k2, {b, B}}]] /. {x1b → x1bvar, x2b → x2bvar, Xb → Xbvar};
MatrixForm[Pxp[x1b, x2b, Xb]]
```
Out[129]//MatrixForm=

$$\begin{pmatrix} -\frac{1}{4}(-1+x1b)\,x1b & \frac{Xb}{4} & \frac{1}{4}(-1\cdot \\ \frac{Xb}{4} & -\frac{1}{4}(-1+x2b)\,x2b & \frac{1}{4}(-1\cdot \\ \frac{1}{4}(-1+r)(-1+2\,x1b)\,Xb & \frac{1}{4}(-1+r)(-1+2\,x2b)\,Xb & \frac{1}{4}(-1+r)^2\left(x1b^2(-1+x2b)\,x2b-Xb\,( \right. \end{pmatrix}$$

In[130]:=

```
(*This is Tx*)
Tx[x1bvar_, x2bvar_, Xbvar_] = Simplify[
    Sum[p[k1, k2, Xb] (xpvec[k1, k2, Xb] - zetax).Transpose[(xvec[k1, k2] - xbvec)],
     {k1, {a, A}}, {k2, {b, B}}]] /. {x1b → x1bvar, x2b → x2bvar, Xb → Xbvar};
MatrixForm[Tx[x1b, x2b, Xb]]

(*This is Hx*)
Hx = Simplify[Tx[x1b, x2b, Xb].Inverse[Px]];
MatrixForm[Hx]
```

Out[131]//MatrixForm=

$$\begin{pmatrix} -\frac{1}{2}(-1+x1b)\,x1b & \frac{Xb}{2} & \frac{1}{2} \\ \frac{Xb}{2} & -\frac{1}{2}(-1+x2b)\,x2b & \frac{1}{2} \\ \frac{1}{2}(-1+r)(-1+2\,x1b)\,Xb & \frac{1}{2}(-1+r)(-1+2\,x2b)\,Xb & -\frac{1}{2}(-1+r)\left(x1b^2(-1+x2b)\,x2b-Xb\,(\right. \end{pmatrix}$$

Out[133]//MatrixForm=

$$\begin{pmatrix} \frac{1}{2} & 0 & 0 \\ 0 & \frac{1}{2} & 0 \\ 0 & 0 & \frac{1-r}{2} \end{pmatrix}$$

In[ ]:=

```
(*This checks that the bottom right entry of Tx is 1/2(1-r)var[X]*)
Simplify[Tx[x1b, x2b, Xb][[3, 3]] == 1/2 (1 - r) varX]
```

Out[ ]=

```
True
```

In[134]:=

```
(*This checks that the determinant of Tx is
 proportional to the product of haplotype frequencies*)

(*This writes haplotype frequencies as functions
 of allele frequencies and linkage disequilibrium*)
pp[i_, j_, x1bb_, x2bb_, Xbb_] := p[i, j, Xb] /. {x1b → x1bb, x2b → x2bb, Xb → Xbb}

(*This is the determinant of Tx*)
DetTx = FullSimplify[Det[Tx[x1b, x2b, Xb]]];

(*This divides the determinant of Tx by the product of haplotype frequencies,
yielding the specified proportionality constant*)
Simplify[DetTx / (pp[a, b, x1b, x2b, Xb] ×
    pp[a, B, x1b, x2b, Xb] × pp[A, b, x1b, x2b, Xb] × pp[A, B, x1b, x2b, Xb])]
```

Out[136]=

$$\frac{1-r}{8}$$

In[137]:=

```
(*This specifies the condition that all
 haplotype frequencies are between zero and one*)
Conditions[x1b_, x2b_, Xb_] :=
 0 ≤ pp[a, b, x1b, x2b, Xb] ≤ 1 && 0 ≤ pp[a, B, x1b, x2b, Xb] ≤ 1 &&
  0 ≤ pp[A, b, x1b, x2b, Xb] ≤ 1 && 0 ≤ pp[A, B, x1b, x2b, Xb] ≤ 1

(*These are the eigenvalues of Tx subject to the constraint
 that haplotype frequencies are between zero and one*)
EigC[x1b_, x2b_, Xb_, rr_] = If[Conditions[x1b, x2b, Xb],
   Eigenvalues[Tx[x1b, x2b, Xb]] /. r → rr, {NaN, NaN, NaN}];

Manipulate[Plot3D[
  {EigC[x1b, x2b, Xb, r]⟦1⟧, EigC[x1b, x2b, Xb, r]⟦2⟧, EigC[x1b, x2b, Xb, r]⟦3⟧},
  {x1b, 0, 1}, {x2b, 0, 1}], {{Xb, 0}, -1, 1}, {{r, 0.5}, 0, 1 / 2}]
```

Out[139]=

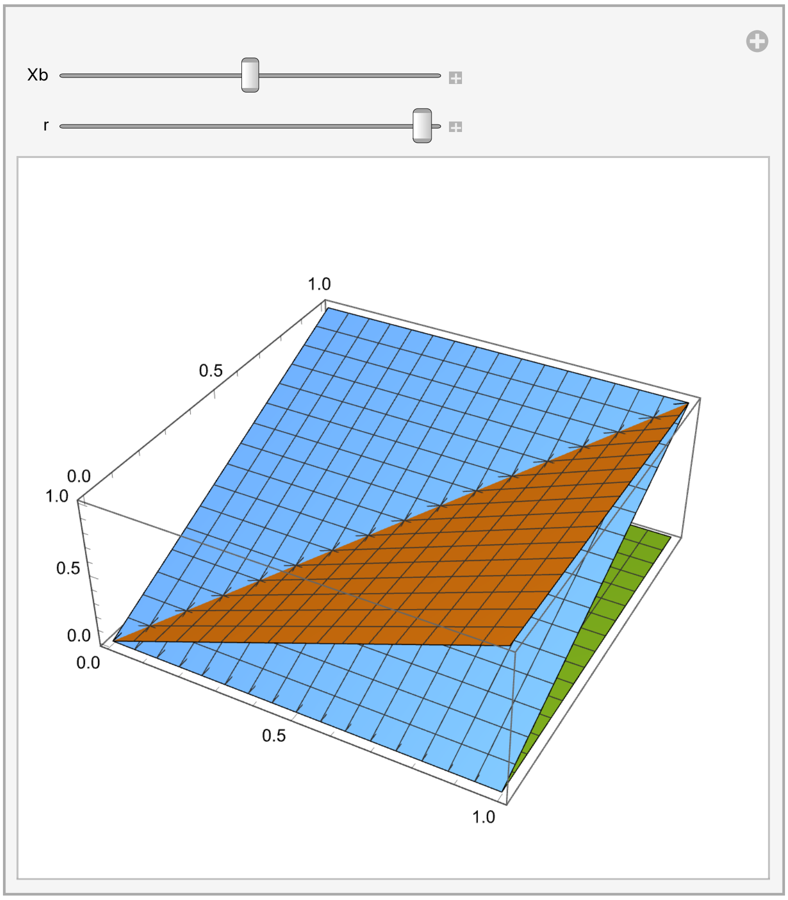

In[140]:=

```
(*These are the eigenvalues of the symmetric part
 of Tx subject to the constraint that haplotype frequencies
 are between zero and one, showing they are non-negative*)

EigCS[x1b_, x2b_, Xb_, rr_] = If[Conditions[x1b, x2b, Xb],
   Eigenvalues[1/2 (Tx[x1b, x2b, Xb] + Transpose[Tx[x1b, x2b, Xb]])] /. r → rr,
   {NaN, NaN, NaN}];

Manipulate[Plot3D[
  {EigCS[x1b, x2b, Xb, r][[1]], EigCS[x1b, x2b, Xb, r][[2]], EigCS[x1b, x2b, Xb, r][[3]]},
  {x1b, 0, 1}, {x2b, 0, 1}], {{Xb, 0.1}, -1, 1}, {{r, 0.4}, 0, 1 / 2}]
```

Out[141]=

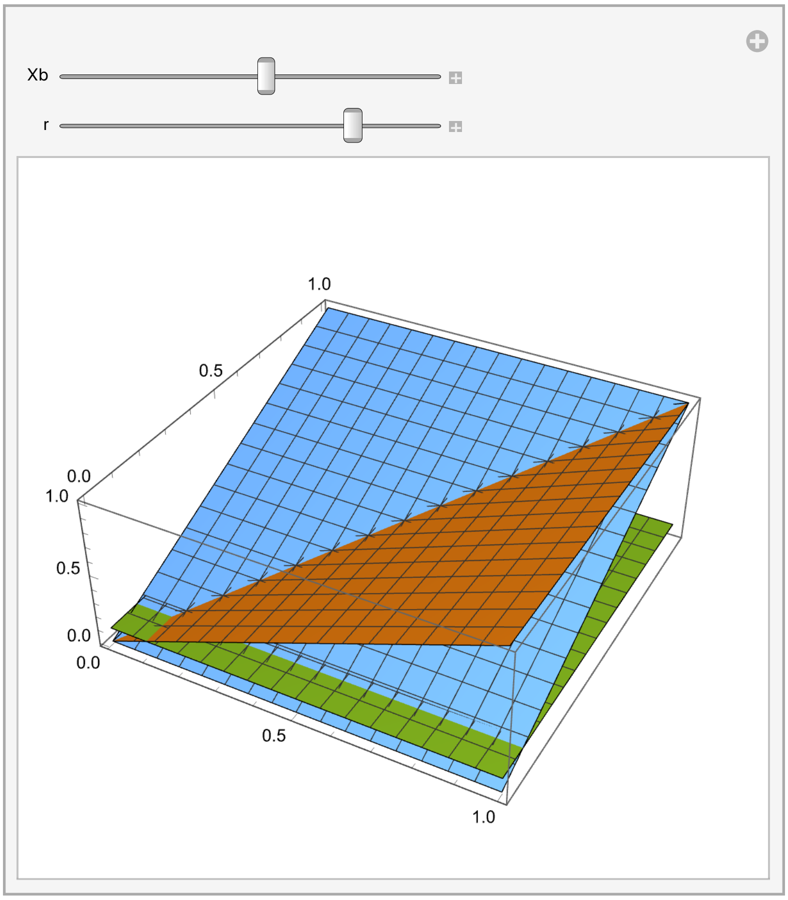

In[ ]:=

```
(*This checks that parent-
 offspring residuals of haplotype content are uncorrelated,
from eq. 18 in main text*)

Simplify[Pxp[x1b, x2b, Xb] == Hx.Px.Hx]
```

Out[ ]=

```
True
```

In[ ]:=
```
(*This is etax*)
eta[k1_, k2_, Xb_] := xpvec[k1, k2, Xb] - zetax - Hx.(xvec[k1, k2] - xbvec);

MatrixForm[Table[Simplify[eta[k1, k2, Xb]], {k1, {a, A}}, {k2, {b, B}}]]
```
Out[ ]//MatrixForm=
```
((0,0,0) (0,0,0)
 (0,0,0) (0,0,0))
```

In[ ]:=
```
(*This is βx in general*)
betaxGen =
  Simplify[Inverse[Px].Sum[p[k1, k2, Xb] (xvec[k1, k2] - xbvec) (wG[k1, k2] - 1),
    {k1, {a, A}}, {k2, {b, B}}]];

MatrixForm[betaxGen]
```
Out[ ]//MatrixForm=
```
( (-1 + x2b) wG[a, b] - x2b wG[a, B] + wG[A, b] - x2b wG[A, b] + x2b wG[A, B]
  (-1 + x1b) wG[a, b] - (-1 + x1b) wG[a, B] + x1b (-wG[A, b] + wG[A, B])
  wG[a, b] - wG[a, B] - wG[A, b] + wG[A, B] )
```

In[ ]:=
```
(*This checks that βx is that presented in eq. 134*)
Simplify[wA - wa == betaxGen[[1, 1]] /.
  {wA → x2b wG[A, B] + (1 - x2b) wG[A, b], wa → x2b wG[a, B] + (1 - x2b) wG[a, b]}]

Simplify[wB - wb == betaxGen[[2, 1]] /.
  {wB → x1b wG[A, B] + (1 - x1b) wG[a, B], wb → x1b wG[A, b] + (1 - x1b) wG[a, b]}]

Simplify[wcou - wrep == betaxGen[[3, 1]] /.
  {wcou → wG[a, b] + wG[A, B], wrep → wG[a, B] + wG[A, b]}]
```
Out[ ]=
```
True
```
Out[ ]=
```
True
```
Out[ ]=
```
True
```

In[ ]:=
```
(*This is the selection pointer βx' in general*)
betaxpGen = FullSimplify[Inverse[Hx].betaxGen];

MatrixForm[betaxpGen]
```
Out[ ]//MatrixForm=
```
( 2 ((-1 + x2b) wG[a, b] + wG[A, b] - x2b (wG[a, B] + wG[A, b] - wG[A, B]))
  2 ((-1 + x1b) wG[a, b] + wG[a, B] - x1b (wG[a, B] + wG[A, b] - wG[A, B]))
  - 2 (wG[a,b]-wG[a,B]-wG[A,b]+wG[A,B]) / (-1+r) )
```

In[ ]:=
```
(*This is mean absolute fitness*)
WbGen = Simplify[Sum[p[k1, k2, Xb] × p[l1, l2, Xb] × WG[k1, k2, l1, l2],
    {k1, {a, A}}, {k2, {b, B}}, {l1, {a, A}}, {l2, {b, B}}]]
```

Out[ ]=
```
(1 + x1b (-1 + x2b) - x2b + Xb)^2 WG[a, b, a, b] -
 (1 + x1b (-1 + x2b) - x2b + Xb) ((-1 + x1b) x2b + Xb) WG[a, b, a, B] -
 (x1b (-1 + x2b) + Xb) (1 + x1b (-1 + x2b) - x2b + Xb) WG[a, b, A, b] +
 (1 + x1b (-1 + x2b) - x2b + Xb) (x1b x2b + Xb) WG[a, b, A, B] -
 (1 + x1b (-1 + x2b) - x2b + Xb) ((-1 + x1b) x2b + Xb) WG[a, B, a, b] +
 ((-1 + x1b) x2b + Xb)^2 WG[a, B, a, B] +
 (x1b (-1 + x2b) + Xb) ((-1 + x1b) x2b + Xb) WG[a, B, A, b] -
 ((-1 + x1b) x2b + Xb) (x1b x2b + Xb) WG[a, B, A, B] -
 (x1b (-1 + x2b) + Xb) (1 + x1b (-1 + x2b) - x2b + Xb) WG[A, b, a, b] +
 (x1b (-1 + x2b) + Xb) ((-1 + x1b) x2b + Xb) WG[A, b, a, B] +
 (x1b (-1 + x2b) + Xb)^2 WG[A, b, A, b] -
 (x1b (-1 + x2b) + Xb) (x1b x2b + Xb) WG[A, b, A, B] +
 (1 + x1b (-1 + x2b) - x2b + Xb) (x1b x2b + Xb) WG[A, B, a, b] -
 ((-1 + x1b) x2b + Xb) (x1b x2b + Xb) WG[A, B, a, B] -
 (x1b (-1 + x2b) + Xb) (x1b x2b + Xb) WG[A, B, A, b] + (x1b x2b + Xb)^2 WG[A, B, A, B]
```

In[ ]:=
```
(*This proves that betax==1/2 1/Wb dWb/dxb assuming that Wij is symmetric, Wij==Wji,
and is independent of allele frequency and of linkage disequilibrium*)

Simplify[
 {betaxGen[[1, 1]] == 1/2 1/WbGen D[WbGen, x1b], betaxGen[[2, 1]] == 1/2 1/WbGen D[WbGen, x2b],
    betaxGen[[3, 1]] == 1/2 1/WbGen D[WbGen, Xb]} /.
   {wG[a, b] → WG[a, b] / WbGen, wG[a, B] → WG[a, B] / WbGen,
    wG[A, b] → WG[A, b] / WbGen, wG[A, B] → WG[A, B] / WbGen} /.
  {WG[a, b] → Sum[p[k1, k2, Xb] × WG[a, b, k1, k2], {k1, {a, A}}, {k2, {b, B}}],
   WG[a, B] → Sum[p[k1, k2, Xb] × WG[a, B, k1, k2], {k1, {a, A}}, {k2, {b, B}}],
   WG[A, b] → Sum[p[k1, k2, Xb] × WG[A, b, k1, k2], {k1, {a, A}}, {k2, {b, B}}],
   WG[A, B] → Sum[p[k1, k2, Xb] × WG[A, B, k1, k2], {k1, {a, A}}, {k2, {b, B}}]} /.
 {WG[a, B, a, b] → WG[a, b, a, B], WG[A, b, a, b] → WG[a, b, A, b],
  WG[A, B, a, b] → WG[a, b, A, B], WG[A, b, a, B] → WG[a, B, A, b],
  WG[A, B, a, B] → WG[a, B, A, B], WG[A, B, A, b] → WG[A, b, A, B]}]
```

Out[ ]=
```
{True, True, True}
```

#### Genotype-phenotype map

In[∘]:=
```
(*This is the m-th phenotype of genotype k1k2l1l2*)
z[m_, k1_, k2_, l1_, l2_] :=
 cb (1 + e[m]) (y[1, A, k1, k2, l1, l2, m] + y[2, B, k1, k2, l1, l2, m]) -
  2 cb e[m] × y[1, A, k1, k2, l1, l2, m] × y[2, B, k1, k2, l1, l2, m]

(*This is the contribution of locus i with allele j to the m-th phenotype*)
y[i_, j_, k1_, k2_, l1_, l2_, m_] :=
 c[i] (1 + d[i, m]) (x[i, j, k1, k2] + x[i, j, l1, l2]) -
  2 c[i] × d[i, m] × x[i, j, k1, k2] × x[i, j, l1, l2]

(*This is the vector of the two phenotypes*)
zvec[k1_, k2_, l1_, l2_] := {{z[1, k1, k2, l1, l2]}, {z[2, k1, k2, l1, l2]}};
```

In[∘]:=
```
(*This is the mean phenotype*)
zbvec[x1bvar_, x2bvar_, Xbvar_] =
  Simplify[Sum[p[k1, k2, Xb] × p[l1, l2, Xb] × zvec[k1, k2, l1, l2],
     {k1, {a, A}}, {k2, {b, B}}, {l1, {a, A}}, {l2, {b, B}}]] /.
   {x1b → x1bvar, x2b → x2bvar, Xb → Xbvar};

MatrixForm[zbvec[x1b, x2b, Xb]]
```

Out[∘]//MatrixForm=

$$\begin{pmatrix} -2\,cb\,\left(x1b^2\,c[1]\times d[1,1]\,\left(1+\left(1+4\,x2b^2\,c[2]\times d[2,1]-4\,x2b\,c[2]\,(1+d[2,1])\right)\,e[1]\right)- \\ -2\,cb\,\left(x1b^2\,c[1]\times d[1,2]\,\left(1+\left(1+4\,x2b^2\,c[2]\times d[2,2]-4\,x2b\,c[2]\,(1+d[2,2])\right)\,e[2]\right)- \end{pmatrix}$$

In[∘]:=
```
(*Mean phenotypes evaluated at the
 chosen dominance and epistasis coefficients*)
Simplify[MatrixForm[zbvec[x1b, x2b, Xb] /. {cb → 1, c[1] → 1, c[2] → 1,
    d[1, 1] → 0, d[1, 2] → 0, d[2, 1] → 0, d[2, 2] → 0, e[1] → 0, e[2] → -1}]]

(*Phenotypes of all genotypes evaluated at
 the chosen dominance and epistasis coefficients*)
Simplify[
 Table[MatrixForm[zvec[k1, k2, l1, l2] /. {cb → 1, c[1] → 1, c[2] → 1, d[1, 1] → 0,
     d[1, 2] → 0, d[2, 1] → 0, d[2, 2] → 0, e[1] → 0, e[2] → -1}],
  {k1, {a, A}}, {k2, {b, B}}, {l1, {a, A}}, {l2, {b, B}}]]
```

Out[∘]//MatrixForm=

$$\begin{pmatrix} 2\,(x1b+x2b) \\ 4\,(2\,x1b\,x2b+Xb) \end{pmatrix}$$

Out[∘]=

$$\left\{\left\{\left\{\left\{\begin{pmatrix}0\\0\end{pmatrix},\begin{pmatrix}1\\0\end{pmatrix}\right\},\left\{\begin{pmatrix}1\\0\end{pmatrix},\begin{pmatrix}2\\2\end{pmatrix}\right\}\right\},\left\{\left\{\begin{pmatrix}1\\0\end{pmatrix},\begin{pmatrix}2\\0\end{pmatrix}\right\},\left\{\begin{pmatrix}2\\2\end{pmatrix},\begin{pmatrix}3\\4\end{pmatrix}\right\}\right\}\right\},\right.$$
$$\left.\left\{\left\{\left\{\begin{pmatrix}1\\0\end{pmatrix},\begin{pmatrix}2\\2\end{pmatrix}\right\},\left\{\begin{pmatrix}2\\0\end{pmatrix},\begin{pmatrix}3\\4\end{pmatrix}\right\}\right\},\left\{\left\{\begin{pmatrix}2\\2\end{pmatrix},\begin{pmatrix}3\\4\end{pmatrix}\right\},\left\{\begin{pmatrix}3\\4\end{pmatrix},\begin{pmatrix}4\\8\end{pmatrix}\right\}\right\}\right\}\right\}$$

```
In[•]:= (*This gives the phenotypes for the specified genotypes*)

        MatrixForm[zvec[A, B, A, B] /. {cb → 1, c[1] → 1, c[2] → 1,
           d[1, 1] → 0, d[1, 2] → 0, d[2, 1] → 0, d[2, 2] → 0, e[1] → 0, e[2] → -1}]
        MatrixForm[zvec[A, B, A, b] /. {cb → 1, c[1] → 1, c[2] → 1,
           d[1, 1] → 0, d[1, 2] → 0, d[2, 1] → 0, d[2, 2] → 0, e[1] → 0, e[2] → -1}]
        MatrixForm[zvec[A, b, A, B] /. {cb → 1, c[1] → 1, c[2] → 1,
           d[1, 1] → 0, d[1, 2] → 0, d[2, 1] → 0, d[2, 2] → 0, e[1] → 0, e[2] → -1}]
        MatrixForm[zvec[A, B, a, B] /. {cb → 1, c[1] → 1, c[2] → 1,
           d[1, 1] → 0, d[1, 2] → 0, d[2, 1] → 0, d[2, 2] → 0, e[1] → 0, e[2] → -1}]
```

Out[•]//MatrixForm=

$$\begin{pmatrix} 4 \\ 8 \end{pmatrix}$$

Out[•]//MatrixForm=

$$\begin{pmatrix} 3 \\ 4 \end{pmatrix}$$

Out[•]//MatrixForm=

$$\begin{pmatrix} 3 \\ 4 \end{pmatrix}$$

Out[•]//MatrixForm=

$$\begin{pmatrix} 3 \\ 4 \end{pmatrix}$$

In[•]:=

```
(*This plots the phenotypes that can be
 developed under the chosen dominance and epistasis*)

(*This makes a list of the possible phenotypes*)
PossiblePhenotypes =
  Flatten[Table[{zvec[k1, k2, l1, l2]⟦1, 1⟧, zvec[k1, k2, l1, l2]⟦2, 1⟧} /.
      {cb → 1, c[1] → 1, c[2] → 1, d[1, 1] → 0, d[1, 2] → 0,
       d[2, 1] → 0, d[2, 2] → 0, e[1] → 0, e[2] → -1},
     {k1, {a, A}}, {k2, {b, B}}, {l1, {a, A}}, {l2, {b, B}}], 3];

(*This plots them*)
plot = ListPlot[PossiblePhenotypes, AspectRatio → 1, PlotStyle → PointSize[.04],
  AxesStyle → Large, PlotRange → {{-0.1, 4.5}, {-0.2, 8.5}}, Epilog →
   Table[Text[Style[Count[PossiblePhenotypes, PossiblePhenotypes⟦i⟧], Large],
     PossiblePhenotypes⟦i⟧ + {.2, 0.2}], {i, 1, Length[PossiblePhenotypes]}]]
(*Rasterize the plot to eliminate vectorization artifacts*)
plot = Rasterize[plot, "Image", ImageResolution → 300];
Export[StringJoin[ToString[NotebookDirectory[]], "Fig.4a", ".pdf"], plot];
```

Out[•]=

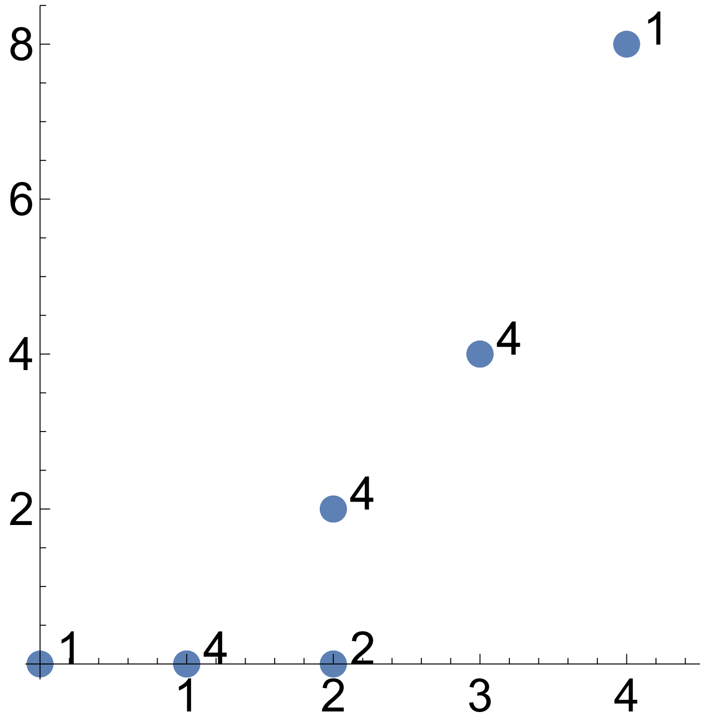

In[•]:=

```
(*This plots the phenotype distribution and phenotype mean for
 a sample of individuals with gene content drawn from a Bernoulli
 distribution with mean given by a chosen allele frequency*)

PopSize = 1000;(*population size*)
ATargetFreq = 0.9;
(*frequency of allele A to be used in Bernoulli distribution*)
BTargetFreq = 0.9;
(*frequency of allele B to be used in Bernoulli distribution*)
```

```
(*This samples the genotypes*)
GenotypeSample = Table[{RandomVariate[BernoulliDistribution[ATargetFreq]],
    RandomVariate[BernoulliDistribution[BTargetFreq]],
    RandomVariate[BernoulliDistribution[ATargetFreq]],
    RandomVariate[BernoulliDistribution[BTargetFreq]]}, {i, 1, PopSize}];

(*This forms the haplotypes*)
Haplotypes = Join[Table[GenotypeSample[[i, 1 ;; 2]], {i, 1, PopSize}],
   Table[GenotypeSample[[i, 3 ;; 4]], {i, 1, PopSize}]];

(*This calculates and prints the haplotype frequencies,
to check they are indeed always between zero and one*)
abFreq = N[Count[Haplotypes, {0, 0}] / Length[Haplotypes]]
aBFreq = N[Count[Haplotypes, {0, 1}] / Length[Haplotypes]]
AbFreq = N[Count[Haplotypes, {1, 0}] / Length[Haplotypes]]
ABFreq = N[Count[Haplotypes, {1, 1}] / Length[Haplotypes]]

(*This lists the phenotypes of the individuals sampled*)
ActualPhenotypes =
  Table[IndividualPhenotype[i] = Flatten[zvec[If[GenotypeSample[[i, 1]] == 0, a, A],
       If[GenotypeSample[[i, 2]] == 0, b, B], If[GenotypeSample[[i, 3]] == 0, a, A],
       If[GenotypeSample[[i, 4]] == 0, b, B]]] /.
     {cb → 1, c[1] → 1, c[2] → 1, d[1, 1] → 0, d[1, 2] → 0, d[2, 1] → 0,
      d[2, 2] → 0, e[1] → 0, e[2] → -1}, {i, 1, PopSize}];

(*This gives the allele frequencies among the sampled individuals*)
AFreq = Sum[Haplotypes[[i, 1]], {i, 1, Length[Haplotypes]}] / Length[Haplotypes];
BFreq = Sum[Haplotypes[[i, 2]], {i, 1, Length[Haplotypes]}] / Length[Haplotypes];

(*This gives the haplotype content deviation for the sampled individuals*)
Xs = Table[X[i] = (Haplotypes[[i, 1]] - AFreq) (Haplotypes[[i, 2]] - BFreq),
   {i, 1, Length[Haplotypes]}];

(*This lists the haplotype contents of the sampled individuals*)
HaplotypeContents =
  Table[Join[Haplotypes[[i]], {Xs[[i]]}], {i, 1, Length[Haplotypes]}];

(*This finds the mean haplotype content*)
MeanHaplotypeContent =
  {Sum[HaplotypeContents[[i, 1]], {i, 1, Length[Haplotypes]}] / Length[Haplotypes],
```

```
    Sum[HaplotypeContents[[i, 2]], {i, 1, Length[Haplotypes]}]
    ---------------------------------------------------------- ,
                     Length[Haplotypes]
    Sum[HaplotypeContents[[i, 3]], {i, 1, Length[Haplotypes]}]
    ---------------------------------------------------------- };
                     Length[Haplotypes]

(*This plots the sampled phenotypes and the
 resulting allele frequencies and linkage disequilibrium*)
plot =
 Show[ListPlot[ActualPhenotypes, AspectRatio → 1, PlotStyle → PointSize[.04],
   AxesStyle → Large, PlotRange → {{-0.1, 4.5}, {-0.2, 8.5}},
   Epilog → {Text[Style[StringJoin[{"LD=", ToString[
          Round[N[MeanHaplotypeContent[[3]]], 0.001]]}], Large], {1.5, 7}],
     Table[Text[Style[Count[ActualPhenotypes, PossiblePhenotypes[[i]]], Large],
       PossiblePhenotypes[[i]] + {.2, 0.2}], {i, 1, Length[PossiblePhenotypes]}]}],
  ListPlot[{zbvec[MeanHaplotypeContent[[1]], MeanHaplotypeContent[[2]],
       MeanHaplotypeContent[[3]]][[1 ;; 2, 1]] /. {cb → 1, c[1] → 1, c[2] → 1,
      d[1, 1] → 0, d[1, 2] → 0, d[2, 1] → 0, d[2, 2] → 0, e[1] → 0, e[2] → -1}},
   PlotStyle → {Red, PointSize[.04]}]]
(*Rasterize the plot to eliminate vectorization artifacts*)
plot = Rasterize[plot, "Image", ImageResolution → 300];
Export[StringJoin[ToString[NotebookDirectory[]], "Fig.4c", ".pA=",
   ToString[ATargetFreq], ".pB=", ToString[BTargetFreq], ".pdf"], plot];
```

Out[ ]=

```
0.015
```

Out[ ]=

```
0.0865
```

Out[ ]=

```
0.082
```

Out[ ]=

```
0.8165
```

Out[ ]=

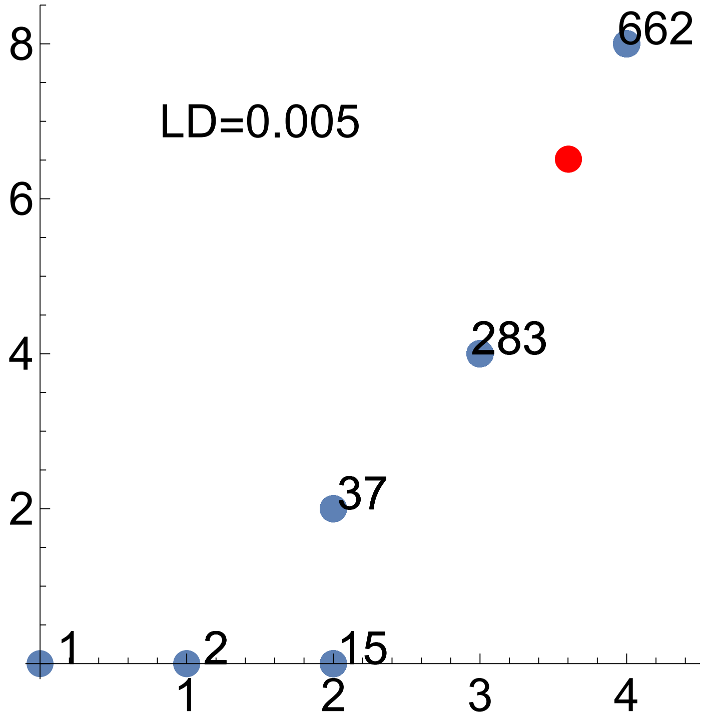

In[ ]:=

```
(*This plots the admissible manifold,
which is the set of points where allele frequency can occur*)

(*Somehow, the plot is unreliable when done using ParametricPlot,
failing to plot regions that should be plotted. So
 it is done here by sampling every point discretely*)

Resol = 0.009; (*Set the resolution; I lower value means higher resolution,
so more points to sample and a heavier file*)

Points = {{0, 0}};(*Starting point that belongs in the manifold;
it belongs because both alleles A and B can be absent*)

(*Build a list with each point in the region by appending points for a
 large combination of allele; this is for linkage diequilibrium = 0.1*)
Table[
  If[Conditions[x1b, x2b, Xb], Points = Append[Points, {zbvec[x1b, x2b, Xb][[1, 1]],
        zbvec[x1b, x2b, Xb][[2, 1]]} /. {cb → 1, c[1] → 1, c[2] → 1, d[1, 1] → 0,
        d[1, 2] → 0, d[2, 1] → 0, d[2, 2] → 0, e[1] → 0, e[2] → -1}]] /.
   Xb → 0.1, {x1b, 0, 1, Resol}, {x2b, 0, 1, Resol}];

(*Make the plot*)
plot = ListPlot[Points, AspectRatio → 1, PlotStyle → {Red, PointSize[Medium]},
  PlotRange → {{0, 4}, {0, 8}}, TicksStyle → Large]
(*Rasterize the plot to eliminate vectorization artifacts*)
plot = Rasterize[plot, "Image", ImageResolution → 300];
Export[
```

```
  StringJoin[ToString[NotebookDirectory[]], "Fig.4d.Xb=0.1", ".pdf"], plot];

(*Do the same for linkage diequilibrium = 0*)
Points = {{0, 0}};
Table[
 If[Conditions[x1b, x2b, Xb], Points = Append[Points, {zbvec[x1b, x2b, Xb][[1, 1]],
       zbvec[x1b, x2b, Xb][[2, 1]]} /. {cb → 1, c[1] → 1, c[2] → 1, d[1, 1] → 0,
       d[1, 2] → 0, d[2, 1] → 0, d[2, 2] → 0, e[1] → 0, e[2] → -1}]] /.
  Xb → 0, {x1b, 0, 1, Resol}, {x2b, 0, 1, Resol}];
plot = ListPlot[Points, AspectRatio → 1, PlotStyle → {Red, PointSize[Medium]},
  PlotRange → {{0, 4}, {0, 8}}, TicksStyle → Large]
(*Rasterize the plot to eliminate vectorization artifacts*)
plot = Rasterize[plot, "Image", ImageResolution → 300];
Export[StringJoin[ToString[NotebookDirectory[]], "Fig.4d.Xb=0", ".pdf"], plot];

(*Do the same for linkage diequilibrium = -0.1*)
Points = {{0, 0}};
Table[
 If[Conditions[x1b, x2b, Xb], Points = Append[Points, {zbvec[x1b, x2b, Xb][[1, 1]],
       zbvec[x1b, x2b, Xb][[2, 1]]} /. {cb → 1, c[1] → 1, c[2] → 1, d[1, 1] → 0,
       d[1, 2] → 0, d[2, 1] → 0, d[2, 2] → 0, e[1] → 0, e[2] → -1}]] /.
  Xb → -0.1, {x1b, 0, 1, Resol}, {x2b, 0, 1, Resol}];
plot = ListPlot[Points, AspectRatio → 1, PlotStyle → {Red, PointSize[Medium]},
  PlotRange → {{0, 4}, {0, 8}}, TicksStyle → Large]
(*Rasterize the plot to eliminate vectorization artifacts*)
plot = Rasterize[plot, "Image", ImageResolution → 300];
Export[
  StringJoin[ToString[NotebookDirectory[]], "Fig.4d.Xb=-0.1", ".pdf"], plot];
```

Out[ ]=

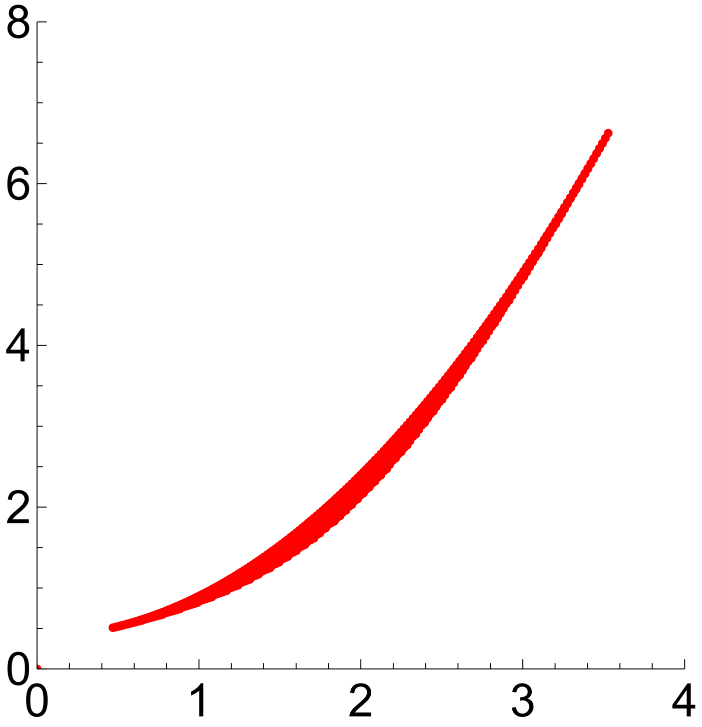

Out[ ]=

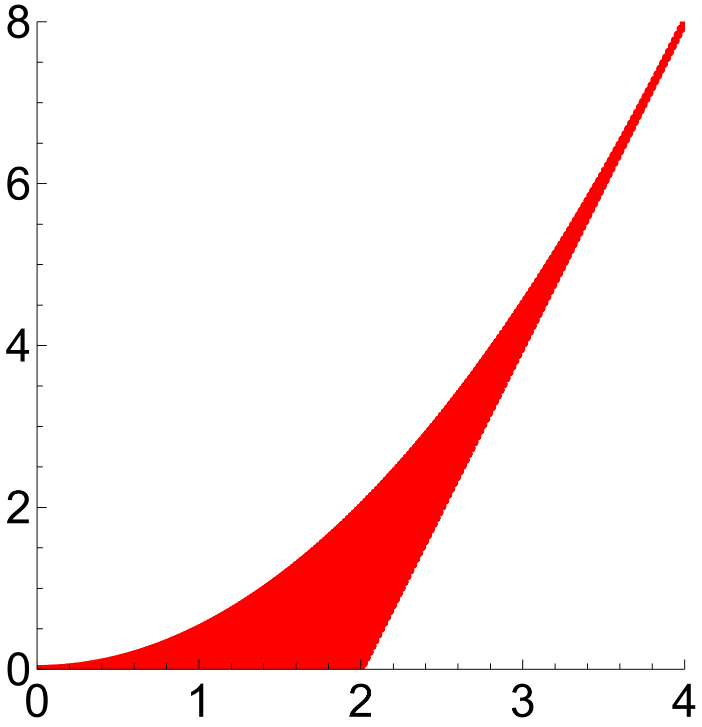

Out[ ]=

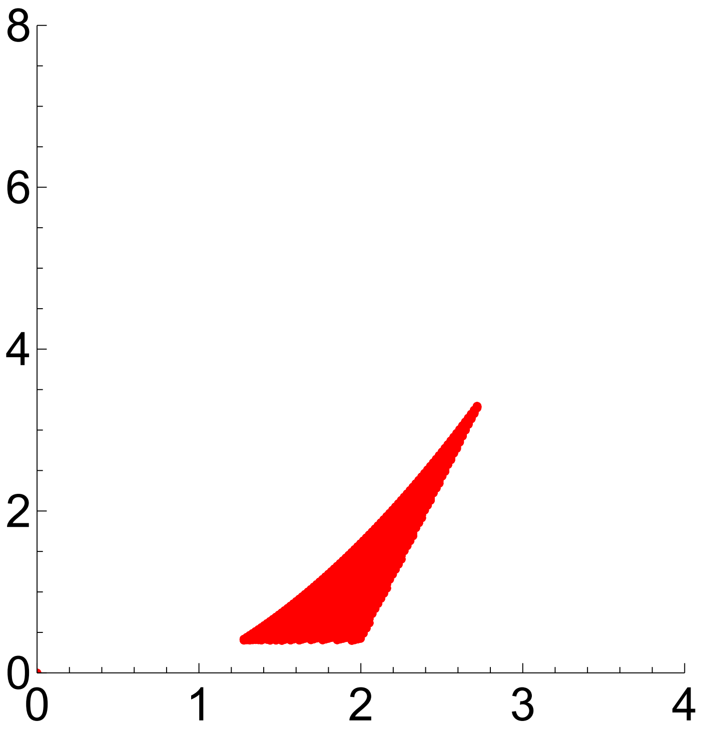

In[ ]:=
```
(*This is P at the chosen dominance and epistasis coefficients*)
P[x1bvar_, x2bvar_, Xbvar_] = Simplify[
    Sum[p[k1, k2, Xb] × p[l1, l2, Xb] (zvec[k1, k2, l1, l2] - zbvec[x1b, x2b, Xb]).
        Transpose[(zvec[k1, k2, l1, l2] - zbvec[x1b, x2b, Xb])], {k1, {a, A}},
      {k2, {b, B}}, {l1, {a, A}}, {l2, {b, B}}] /. {cb → 1, c[1] → 1, c[2] → 1,
      d[1, 1] → 0, d[1, 2] → 0, d[2, 1] → 0, d[2, 2] → 0, e[1] → 0, e[2] → -1}] /.
   {x1b → x1bvar, x2b → x2bvar, Xb → Xbvar};
MatrixForm[P[x1b, x2b, Xb]]

(*This checks that the var[z2] is as written in the main text, eq. 141b*)
Simplify[P[x1b, x2b, Xb][[2, 2]] ==
  8 (2 x1b x2b (1 + x1b + x2b - 3 x1b x2b) + Xb (1 + 2 x1b + 2 x2b - 4 x1b x2b))]
```

Out[ ]//MatrixForm=

$$\begin{pmatrix} -2\left(-x1b + x1b^2 - x2b + x2b^2 - 2\,Xb\right) & -8\left(x1b^2\,x2b + x1b\,(-2 + x2b)\,x2b - \right. \\ -8\left(x1b^2\,x2b + x1b\,(-2 + x2b)\,x2b - Xb\right) & 8\left(2\,x1b^2\,(1 - 3\,x2b)\,x2b + Xb + 2\,x2b\,Xb + 2\,x1b\,(x2b + \right. \end{pmatrix}$$

Out[ ]=

True

#### Evolution

```
In[ ]:= (*This is fitness of genotype kl*)
W[k1_, k2_, l1_, l2_, θ1_, σ1_, θ2_, σ2_] :=
  Exp[-(z[1, k1, k2, l1, l2] - θ1)^2/σ1^2 - (z[2, k1, k2, l1, l2] - θ2)^2/σ2^2]

(*This is mean fitness*)
Wb[Xb_, θ1_, σ1_, θ2_, σ2_] :=
  Simplify[Sum[p[k1, k2, Xb] × p[l1, l2, Xb] × W[k1, k2, l1, l2, θ1, σ1, θ2, σ2],
    {k1, {a, A}}, {k2, {b, B}}, {l1, {a, A}}, {l2, {b, B}}]]

WbFun[x1bvar_, x2bvar_, Xbvar_] =
   Wb[Xb, θ1, σ1, θ2, σ2] /. {x1b → x1bvar, x2b → x2bvar, Xb → Xbvar};

(*This is relative fitness of genotype kl*)
w[k1_, k2_, l1_, l2_, θ1_, σ1_, θ2_, σ2_, Xb_] :=
  W[k1, k2, l1, l2, θ1, σ1, θ2, σ2] / Wb[Xb, θ1, σ1, θ2, σ2]

(*This is the relative fitness of haplotype k*)
wG[k1_, k2_, θ1_, σ1_, θ2_, σ2_, Xb_] := Simplify[Sum[
    p[l1, l2, Xb] × w[k1, k2, l1, l2, θ1, σ1, θ2, σ2, Xb], {l1, {a, A}}, {l2, {b, B}}]]

(*This runs the numerical solutions*)

r = 0.5;

cb = 1;
c[1] = 1;
c[2] = 1;
d[1, 1] = 0;
d[1, 2] = 0;
d[2, 1] = 0;
d[2, 2] = 0;
e[1] = 0;
e[2] = -1;

θ1 = 2;
θ2 = 4;
σ1 = Sqrt[10];
σ2 = Sqrt[40];

tend = 100;

(*This is Δx bar to be used for numerical solutions*)
Deltax[x1bvar_, x2bvar_, Xbvar_] =
   Tx[x1b, x2b, Xb].betaxGen + zetax - xbFun[x1b, x2b, Xb] /.
```

```
{wG[a, b] → wG[a, b, θ1, σ1, θ2, σ2, Xb], wG[a, B] → wG[a, B, θ1, σ1, θ2, σ2, Xb],
 wG[A, b] → wG[A, b, θ1, σ1, θ2, σ2, Xb], wG[A, B] →
  wG[A, B, θ1, σ1, θ2, σ2, Xb]} /. {x1b → x1bvar, x2b → x2bvar, Xb → Xbvar};

(*This defines the range of values from which the initial
 allele frequencies and linkage disequilibrium are sampled;
later it is narrowed down to those where haplotype
 frequencies are between zero and one*)
x1init = {0.01, 1, .2};
x2init = {0.01, 1, .2};
Xbinit = {-0.99, 1, .1};

(*If the initial allele frequencies and linkage disequilibrium for which
 haplotype frequences are between zero and one, the solution is run*)
Table[If[Conditions[i, j, k], {x1bSol[0, i, j, k] = i,
    x2bSol[0, i, j, k] = j, XbSol[0, i, j, k] = k, xbvecSol[0, i, j, k] =
     {{x1bSol[0, i, j, k]}, {x2bSol[0, i, j, k]}, {XbSol[0, i, j, k]}},
    Table[{xbvecSol[t + 1, i, j, k] = xbvecSol[t, i, j, k] +
        Deltax[x1bSol[t, i, j, k], x2bSol[t, i, j, k], XbSol[t, i, j, k]],
      x1bSol[t + 1, i, j, k] = xbvecSol[t + 1, i, j, k]〚1, 1〛,
      x2bSol[t + 1, i, j, k] = xbvecSol[t + 1, i, j, k]〚2, 1〛,
      XbSol[t + 1, i, j, k] = xbvecSol[t + 1, i, j, k]〚3, 1〛}, {t, 0, tend}]}],
  {i, x1init〚1〛, x1init〚2〛, x1init〚3〛}, {j, x2init〚1〛,
   x2init〚2〛, x2init〚3〛},
  {k, Xbinit〚1〛, Xbinit〚2〛, Xbinit〚3〛}];
```

In[ ]:=
```
(*This counts the number of initial conditions that meet the
 criterion that haplotype frequencies are between zero and one*)
Counter = 0;
Table[If[Conditions[i, j, k], {Counter = Counter + 1;
    ActualInitialConditions[Counter] = {i, j, k}}],
  {i, x1init〚1〛, x1init〚2〛, x1init〚3〛}, {j, x2init〚1〛, x2init〚2〛, x2init〚3〛},
  {k, Xbinit〚1〛, Xbinit〚2〛, Xbinit〚3〛}];
CounterEnd = Counter
```

Out[ ]=
```
40
```

In[ ]:=
```
(*This makes a few plots*)

(*This computes the final points to be plotted*)
pointSize = 0.05;
endPointsxP = Table[{x1bSol[tend, ActualInitialConditions[Counter]〚1〛,
     ActualInitialConditions[Counter]〚2〛, ActualInitialConditions[Counter]〚3〛],
    x2bSol[tend, ActualInitialConditions[Counter]〚1〛,
     ActualInitialConditions[Counter]〚2〛,
     ActualInitialConditions[Counter]〚3〛]}, {Counter, 1, CounterEnd}];
```

```
(*Set the axes limits*)
yAxisLimitz := 1.1
yAxisLimitP := .6

lThickness = 0.02; (*Line thickness*)

(*This plots the change of allele frequencies
 for both loci on top of the mean fitness contour*)
plot = Show[ContourPlot[Wb[0, θ1, σ1, θ2, σ2], {x1b, 0, 1}, {x2b, 0, 1},
   FrameStyle → Large, PlotLegends → Automatic, LabelStyle → Large,
   TicksStyle → Large, PlotRange → {{-0.1, 1.1}, {-.1, 1.1}},
   Epilog → {PointSize[pointSize], Point[endPointsxP]}],
  Table[ListLinePlot[Table[{x1bSol[t, ActualInitialConditions[Counter][[1]],
       ActualInitialConditions[Counter][[2]],
       ActualInitialConditions[Counter][[3]]], x2bSol[t,
       ActualInitialConditions[Counter][[1]], ActualInitialConditions[Counter][[
        2]], ActualInitialConditions[Counter][[3]]]}, {t, 0, tend}],
    PlotStyle → Thickness[lThickness]], {Counter, 1, CounterEnd}]]
(*Rasterize the plot to eliminate vectorization artifacts*)
plot = Rasterize[plot, "Image", ImageResolution → 300];
Export[StringJoin[ToString[NotebookDirectory[]], "Fig.5a", ".pdf"], plot];

(*This shows the change in allele frequencies
 in both loci and in linkage disquilibrium over time*)
Show[Table[ListLinePlot[{Table[x1bSol[t,
      ActualInitialConditions[Counter][[1]], ActualInitialConditions[Counter][[2]],
      ActualInitialConditions[Counter][[3]]], {t, 0, tend}],
    Table[x2bSol[t, ActualInitialConditions[Counter][[1]],
      ActualInitialConditions[Counter][[2]],
      ActualInitialConditions[Counter][[3]]], {t, 0, tend}],
    Table[XbSol[t, ActualInitialConditions[Counter][[1]],
      ActualInitialConditions[Counter][[2]],
      ActualInitialConditions[Counter][[3]]], {t, 0, tend}]},
   PlotRange → {-0.2, yAxisLimitz}, TicksStyle → Large,
   PlotStyle → Thickness[lThickness]],
  {Counter, 1, CounterEnd}]]
Export[StringJoin[ToString[NotebookDirectory[]], "Fig.5c", ".pdf"], %];

(*This plots the change in mean absolute fitness over time*)
Show[
 Table[ListLinePlot[{Table[WbFun[x1bSol[t, ActualInitialConditions[Counter][[1]],
       ActualInitialConditions[Counter][[2]],
       ActualInitialConditions[Counter][[3]]], x2bSol[t,
       ActualInitialConditions[Counter][[1]], ActualInitialConditions[Counter][[
        2]], ActualInitialConditions[Counter][[3]]], XbSol[t,
       ActualInitialConditions[Counter][[1]], ActualInitialConditions[Counter][[
        2]], ActualInitialConditions[Counter][[3]]]], {t, 0, tend}]},
```

```
    PlotRange → {0, yAxisLimitz}, TicksStyle → Large,
    PlotStyle → Thickness[lThickness]],
  {Counter, 1, CounterEnd}]]
Export[StringJoin[ToString[NotebookDirectory[]], "Fig.5e", ".pdf"], %];
```

Out[ ]=

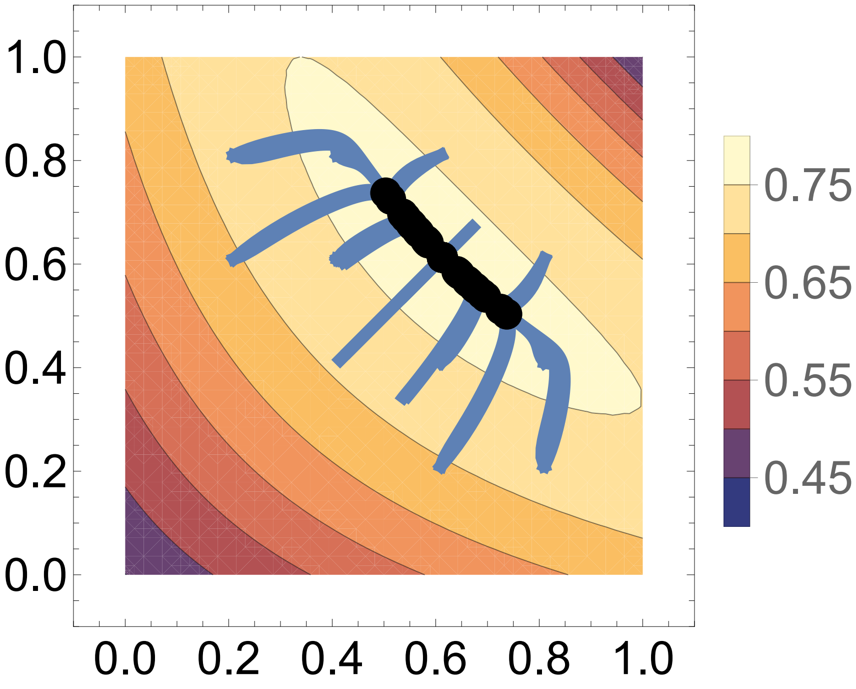

Out[ ]=

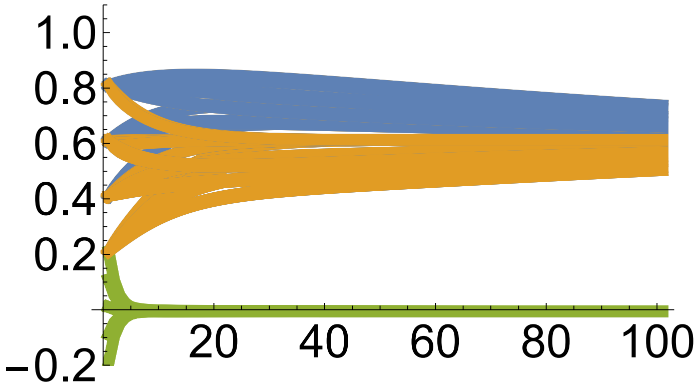

Out[ ]=

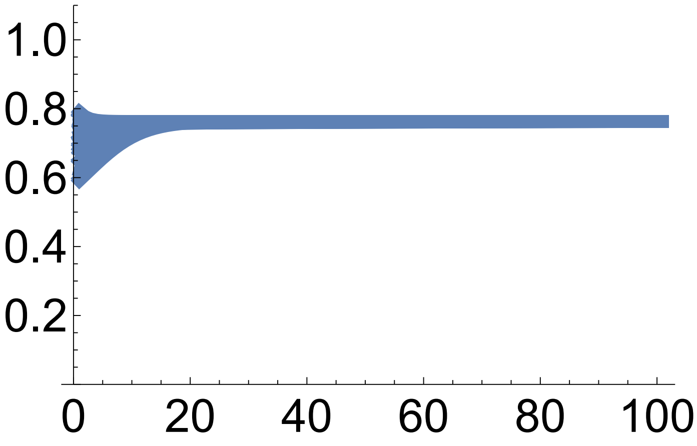

```
In[]:= (*This plots haplotype frequencies over time*)
Show[
 Table[ListLinePlot[{Table[(1 - x1bSol[t, ActualInitialConditions[Counter]〚1〛,
          ActualInitialConditions[Counter]〚2〛,
          ActualInitialConditions[Counter]〚3〛]) (1 - x2bSol[t,
          ActualInitialConditions[Counter]〚1〛, ActualInitialConditions[
            Counter]〚2〛, ActualInitialConditions[Counter]〚3〛]) +
      XbSol[t, ActualInitialConditions[Counter]〚1〛,
       ActualInitialConditions[Counter]〚2〛,
       ActualInitialConditions[Counter]〚3〛], {t, 0, tend}],
    Table[x1bSol[t, ActualInitialConditions[Counter]〚1〛,
        ActualInitialConditions[Counter]〚2〛,
        ActualInitialConditions[Counter]〚3〛]
       (1 - x2bSol[t, ActualInitialConditions[Counter]〚1〛,
          ActualInitialConditions[Counter]〚2〛,
          ActualInitialConditions[Counter]〚3〛]) -
      XbSol[t, ActualInitialConditions[Counter]〚1〛,
       ActualInitialConditions[Counter]〚2〛,
       ActualInitialConditions[Counter]〚3〛], {t, 0, tend}], Table[
     (1 - x1bSol[t, ActualInitialConditions[Counter]〚1〛, ActualInitialConditions[
            Counter]〚2〛, ActualInitialConditions[Counter]〚3〛])
       x2bSol[t, ActualInitialConditions[Counter]〚1〛, ActualInitialConditions[
          Counter]〚2〛, ActualInitialConditions[Counter]〚3〛] -
      XbSol[t, ActualInitialConditions[Counter]〚1〛,
       ActualInitialConditions[Counter]〚2〛,
       ActualInitialConditions[Counter]〚3〛], {t, 0, tend}],
    Table[x1bSol[t, ActualInitialConditions[Counter]〚1〛,
        ActualInitialConditions[Counter]〚2〛,
        ActualInitialConditions[Counter]〚3〛] ×
       x2bSol[t, ActualInitialConditions[Counter]〚1〛,
        ActualInitialConditions[Counter]〚2〛,
        ActualInitialConditions[Counter]〚3〛] +
      XbSol[t, ActualInitialConditions[Counter]〚1〛,
       ActualInitialConditions[Counter]〚2〛,
       ActualInitialConditions[Counter]〚3〛], {t, 0, tend}]},
   PlotRange → {-0.1, yAxisLimitz}, TicksStyle → Large,
   PlotStyle →
    Thickness[lThickness]],
  {Counter, 1, CounterEnd}]]
```

Out[ ]=

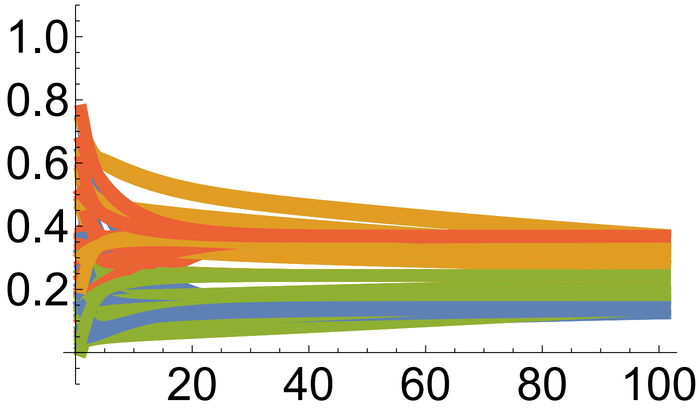

In[ ]:=
```
(*This plots mean z1 (blue) and mean z2 (orange) over time*)
Show[
 Table[ListLinePlot[{Table[zbvec[x1bSol[t, ActualInitialConditions[Counter]〚1〛,
        ActualInitialConditions[Counter]〚2〛,
        ActualInitialConditions[Counter]〚3〛], x2bSol[t,
        ActualInitialConditions[Counter]〚1〛, ActualInitialConditions[Counter]〚
         2〛, ActualInitialConditions[Counter]〚3〛], XbSol[t,
        ActualInitialConditions[Counter]〚1〛, ActualInitialConditions[Counter]〚
         2〛, ActualInitialConditions[Counter]〚3〛]]〚1, 1〛, {t, 0, tend}],
    Table[zbvec[x1bSol[t, ActualInitialConditions[Counter]〚1〛,
        ActualInitialConditions[Counter]〚2〛,
        ActualInitialConditions[Counter]〚3〛], x2bSol[t,
        ActualInitialConditions[Counter]〚1〛, ActualInitialConditions[Counter]〚
         2〛, ActualInitialConditions[Counter]〚3〛], XbSol[t,
        ActualInitialConditions[Counter]〚1〛, ActualInitialConditions[Counter]〚
         2〛, ActualInitialConditions[Counter]〚3〛]]〚2, 1〛, {t, 0, tend}],
    Table[θ1, {t, 0, tend}], Table[θ2, {t, 0, tend}]}, PlotRange →
    {0, 6}, TicksStyle → Large,
   PlotStyle → {Thickness[lThickness], Thickness[lThickness],
     {Magenta, Thickness[lThickness]},
     {Magenta, Thickness[lThickness]}}], {Counter, 1, CounterEnd}]]
Export[StringJoin[ToString[NotebookDirectory[]], "Fig.5d", ".pdf"], %];
```

Out[ ]=

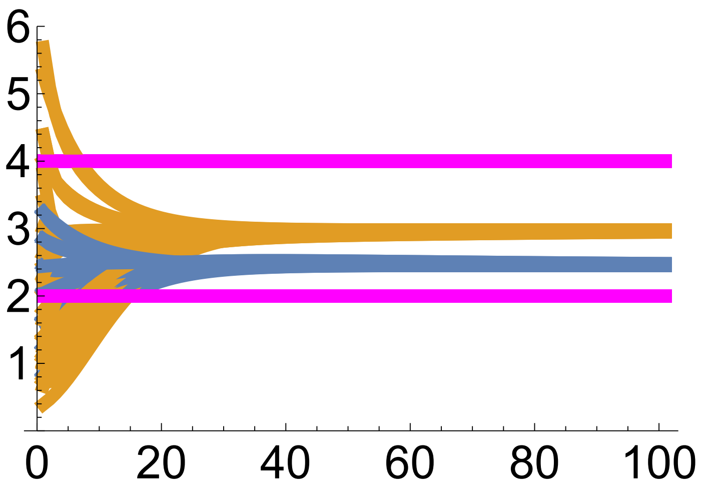

In[ ]:=
```
(*This makes another series of plots*)

(*This computes the final points for the plots of the covariance evolution*)
endPointszP = Table[{zbvec[x1bSol[tend, ActualInitialConditions[Counter]〚1〛,
       ActualInitialConditions[Counter]〚2〛,
       ActualInitialConditions[Counter]〚3〛], x2bSol[tend,
       ActualInitialConditions[Counter]〚1〛, ActualInitialConditions[Counter]〚
        2〛, ActualInitialConditions[Counter]〚3〛],
      XbSol[tend, ActualInitialConditions[Counter]〚1〛,
       ActualInitialConditions[Counter]〚2〛,
```

```
        ActualInitialConditions[Counter]⟦3⟧]]⟦1, 1⟧,
     zbvec[x1bSol[tend, ActualInitialConditions[Counter]⟦1⟧,
        ActualInitialConditions[Counter]⟦2⟧,
        ActualInitialConditions[Counter]⟦3⟧],
       x2bSol[tend, ActualInitialConditions[Counter]⟦1⟧,
        ActualInitialConditions[Counter]⟦2⟧,
        ActualInitialConditions[Counter]⟦3⟧],
       XbSol[tend, ActualInitialConditions[Counter]⟦1⟧,
        ActualInitialConditions[Counter]⟦2⟧,
        ActualInitialConditions[Counter]⟦3⟧]]⟦2, 1⟧}, {Counter, 1, CounterEnd}];

(*This plots mean z1 vs mean z2 with
 admissible developmental manifold in red*)

(*First, determine the initial and final
 admissible manifolds for a particular run*)
Table[Table[{PointsSol[t] = {{0, 0}};,
    Table[If[Conditions[x1b, x2b, Xb], PointsSol[t] = Append[PointsSol[t],
          {zbvec[x1b, x2b, Xb]⟦1, 1⟧, zbvec[x1b, x2b, Xb]⟦2, 1⟧}];] /.
        Xb → XbSol[t, ActualInitialConditions[Counter]⟦1⟧,
          ActualInitialConditions[Counter]⟦2⟧,
          ActualInitialConditions[Counter]⟦3⟧],
       {x1b, 0, 1, Resol}, {x2b, 0, 1, Resol}];},
   {t, {0, tend}}], {Counter, {CounterEnd}}];

(*Then, do the plot*)
plot = Show[Table[Table[{
      ListPlot[PointsSol[t], AspectRatio → 1,
       PlotStyle → {Red, PointSize[Medium]}, PlotRange → {{0, 4}, {0, 8}},
       TicksStyle → Large, Epilog → {{PointSize[pointSize], Point[endPointszP]},
         {PointSize[pointSize], Magenta, Point[{θ1, θ2}]}}]},
     {t, {0, tend}}], {Counter, {CounterEnd}}], Table[ListLinePlot[
     {Table[{zbvec[x1bSol[t, ActualInitialConditions[Counter]⟦1⟧,
           ActualInitialConditions[Counter]⟦2⟧,
           ActualInitialConditions[Counter]⟦3⟧], x2bSol[t,
           ActualInitialConditions[Counter]⟦1⟧, ActualInitialConditions[
             Counter]⟦2⟧, ActualInitialConditions[Counter]⟦3⟧],
          XbSol[t, ActualInitialConditions[Counter]⟦1⟧, ActualInitialConditions[
             Counter]⟦2⟧, ActualInitialConditions[Counter]⟦3⟧]]⟦1, 1⟧,
        zbvec[x1bSol[t, ActualInitialConditions[Counter]⟦1⟧,
           ActualInitialConditions[Counter]⟦2⟧,
           ActualInitialConditions[Counter]⟦3⟧], x2bSol[t,
           ActualInitialConditions[Counter]⟦1⟧, ActualInitialConditions[
             Counter]⟦2⟧, ActualInitialConditions[Counter]⟦3⟧],
          XbSol[t, ActualInitialConditions[Counter]⟦1⟧, ActualInitialConditions[
             Counter]⟦2⟧, ActualInitialConditions[Counter]⟦3⟧]]⟦2, 1⟧},
```

```
        {t, 0, tend}]}, PlotRange → {{-0.1, 4.1}, {-0.1, 6.1}},
     PlotStyle → Thickness[0.5 lThickness]], {Counter,
     1,
     CounterEnd}]]
(*Rasterize the plot to eliminate vectorization artifacts*)
plot = Rasterize[plot, "Image", ImageResolution → 300];
Export[StringJoin[ToString[NotebookDirectory[]], "Fig.5b", ".pdf"], plot];

(*This plots, over time, the variance of phenotype 1 (cov[z1,z1], blue),
the variance of phenotype 2 (cov[z2,z2], orange),
and the covariance of phenotype 1 and 2 (cov[z1,z2], green)*)
Show[Table[ListLinePlot[{Table[
      P[x1bSol[t, ActualInitialConditions[Counter]〚1〛, ActualInitialConditions[
            Counter]〚2〛, ActualInitialConditions[Counter]〚3〛],
        x2bSol[t, ActualInitialConditions[Counter]〚1〛, ActualInitialConditions[
            Counter]〚2〛, ActualInitialConditions[Counter]〚3〛], XbSol[t,
         ActualInitialConditions[Counter]〚1〛, ActualInitialConditions[Counter]〚
          2〛, ActualInitialConditions[Counter]〚3〛]]〚1, 1〛, {t, 0, tend}],
     Table[P[x1bSol[t, ActualInitialConditions[Counter]〚1〛,
         ActualInitialConditions[Counter]〚2〛,
         ActualInitialConditions[Counter]〚3〛], x2bSol[t,
         ActualInitialConditions[Counter]〚1〛, ActualInitialConditions[Counter]〚
          2〛, ActualInitialConditions[Counter]〚3〛], XbSol[t,
         ActualInitialConditions[Counter]〚1〛, ActualInitialConditions[Counter]〚
          2〛, ActualInitialConditions[Counter]〚3〛]]〚2, 2〛, {t, 0, tend}],
     Table[P[x1bSol[t, ActualInitialConditions[Counter]〚1〛,
         ActualInitialConditions[Counter]〚2〛,
         ActualInitialConditions[Counter]〚3〛], x2bSol[t,
         ActualInitialConditions[Counter]〚1〛, ActualInitialConditions[Counter]〚
          2〛, ActualInitialConditions[Counter]〚3〛],
        XbSol[t, ActualInitialConditions[Counter]〚1〛,
         ActualInitialConditions[Counter]〚2〛,
         ActualInitialConditions[Counter]〚3〛]]〚2, 1〛, {t, 0, tend}]},
    PlotRange → {0, 10.5}, TicksStyle → Large, PlotStyle →
     Thickness[lThickness]],
   {Counter, 1, CounterEnd}]]
Export[StringJoin[ToString[NotebookDirectory[]], "Fig.5h", ".pdf"], %];
```

Out[◦]=

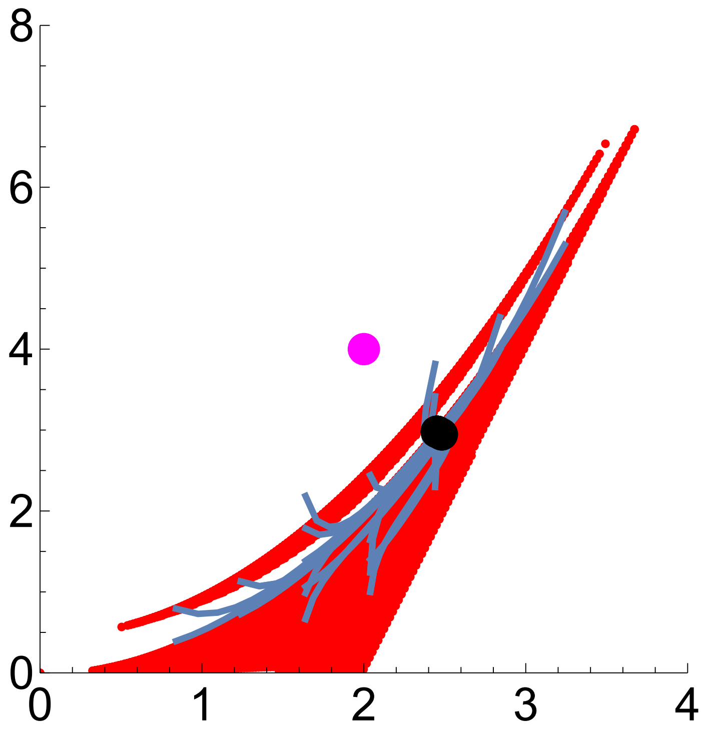

Out[◦]=

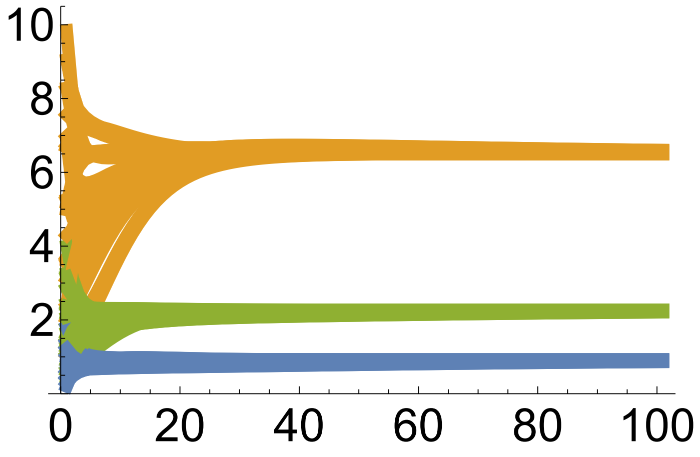

In[◦]:=

```
(*This plots, over time,
the allelic variance in locus 1 for allele A (cov[x1,x1], blue),
the allelic variance in locus 2 for allele B (cov[x2,x2], orange),
and the variance in haplotype content
 deviation between alleles A and B (cov[X,X], green)*)

Show[
 Table[ListLinePlot[{Table[PxFun[x1bSol[t, ActualInitialConditions[Counter]⟦1⟧,
        ActualInitialConditions[Counter]⟦2⟧,
        ActualInitialConditions[Counter]⟦3⟧], x2bSol[t,
        ActualInitialConditions[Counter]⟦1⟧, ActualInitialConditions[Counter]⟦
         2⟧, ActualInitialConditions[Counter]⟦3⟧], XbSol[t,
        ActualInitialConditions[Counter]⟦1⟧, ActualInitialConditions[Counter]⟦
         2⟧, ActualInitialConditions[Counter]⟦3⟧]]⟦1, 1⟧, {t, 0, tend}],
```

```
      Table[PxFun[x1bSol[t, ActualInitialConditions[Counter]⟦1⟧,
          ActualInitialConditions[Counter]⟦2⟧,
          ActualInitialConditions[Counter]⟦3⟧], x2bSol[t,
          ActualInitialConditions[Counter]⟦1⟧, ActualInitialConditions[Counter]⟦
           2⟧, ActualInitialConditions[Counter]⟦3⟧], XbSol[t,
          ActualInitialConditions[Counter]⟦1⟧, ActualInitialConditions[Counter]⟦
           2⟧, ActualInitialConditions[Counter]⟦3⟧]]⟦2, 2⟧, {t, 0, tend}],
      Table[PxFun[x1bSol[t, ActualInitialConditions[Counter]⟦1⟧,
          ActualInitialConditions[Counter]⟦2⟧,
          ActualInitialConditions[Counter]⟦3⟧], x2bSol[t,
          ActualInitialConditions[Counter]⟦1⟧, ActualInitialConditions[Counter]⟦
           2⟧, ActualInitialConditions[Counter]⟦3⟧], XbSol[t,
          ActualInitialConditions[Counter]⟦1⟧, ActualInitialConditions[Counter]⟦
           2⟧, ActualInitialConditions[Counter]⟦3⟧]]⟦3, 3⟧, {t, 0, tend}]},
     PlotRange → {0, .3}, TicksStyle → Large, PlotStyle →
      Thickness[lThickness]],
    {Counter, 1, CounterEnd}]]
Export[StringJoin[ToString[NotebookDirectory[]], "Fig.5g", ".pdf"], %];

(*This plots, over time, the covariance between allele A and haplotype
 content deviation(cov[x1,X], blue) and the covariance between
 allele B and haplotype content deviation (cov[x2,X], orange)*)
Show[
 Table[ListLinePlot[{Table[PxFun[x1bSol[t, ActualInitialConditions[Counter]⟦1⟧,
          ActualInitialConditions[Counter]⟦2⟧,
          ActualInitialConditions[Counter]⟦3⟧], x2bSol[t,
          ActualInitialConditions[Counter]⟦1⟧, ActualInitialConditions[Counter]⟦
           2⟧, ActualInitialConditions[Counter]⟦3⟧], XbSol[t,
          ActualInitialConditions[Counter]⟦1⟧, ActualInitialConditions[Counter]⟦
           2⟧, ActualInitialConditions[Counter]⟦3⟧]]⟦1, 3⟧, {t, 0, tend}],
      Table[PxFun[x1bSol[t, ActualInitialConditions[Counter]⟦1⟧,
          ActualInitialConditions[Counter]⟦2⟧,
          ActualInitialConditions[Counter]⟦3⟧], x2bSol[t,
          ActualInitialConditions[Counter]⟦1⟧, ActualInitialConditions[Counter]⟦
           2⟧, ActualInitialConditions[Counter]⟦3⟧], XbSol[t,
          ActualInitialConditions[Counter]⟦1⟧, ActualInitialConditions[Counter]⟦
           2⟧, ActualInitialConditions[Counter]⟦3⟧]]⟦2, 3⟧, {t, 0, tend}]},
     PlotRange → {-0.1, .1}, TicksStyle → Large, PlotStyle →
      Thickness[lThickness]],
    {Counter, 1, CounterEnd}]]
Export[StringJoin[ToString[NotebookDirectory[]], "Fig.5f", ".pdf"], %];
```

Out[ ]=

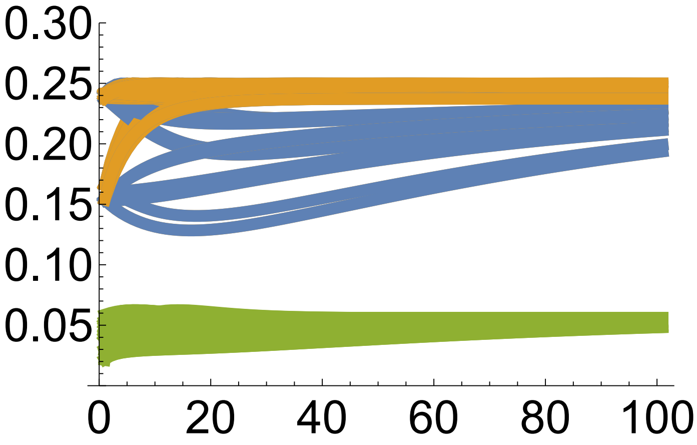
0.30
0.25
0.20
0.15
0.10
0.05
0
20
40
60
80
100

Out[ ]=

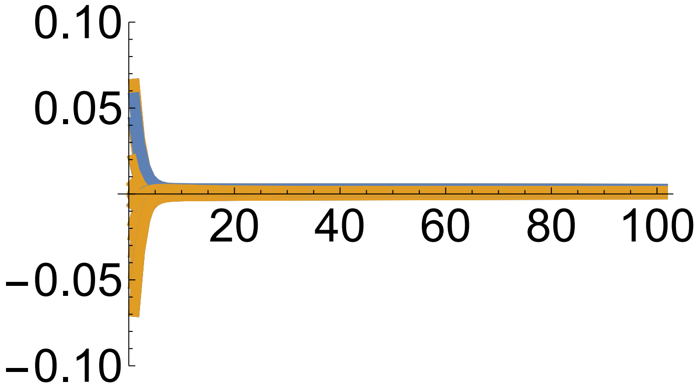
0.10
0.05
20
40
60
80
100
-0.05
-0.10

In[ ]:=
```
(*This plots, over time, the covariance between offspring haplotype
 content deviation and parent gene content for A (cov[X',x1], blue)
 and the covariance between offspring haplotype content
 deviation and parent gene content for B (cov[X',x2], orange)*)
Show[
 Table[ListLinePlot[{Table[Tx[x1bSol[t, ActualInitialConditions[Counter][[1]],
        ActualInitialConditions[Counter][[2]],
        ActualInitialConditions[Counter][[3]]], x2bSol[t,
        ActualInitialConditions[Counter][[1]], ActualInitialConditions[Counter][[
         2]], ActualInitialConditions[Counter][[3]]], XbSol[t,
        ActualInitialConditions[Counter][[1]], ActualInitialConditions[Counter][[
         2]], ActualInitialConditions[Counter][[3]]]][[3, 1]], {t, 0, tend}],
    Table[Tx[x1bSol[t, ActualInitialConditions[Counter][[1]],
        ActualInitialConditions[Counter][[2]],
        ActualInitialConditions[Counter][[3]]], x2bSol[t,
        ActualInitialConditions[Counter][[1]], ActualInitialConditions[Counter][[
         2]], ActualInitialConditions[Counter][[3]]], XbSol[t,
        ActualInitialConditions[Counter][[1]], ActualInitialConditions[Counter][[
         2]], ActualInitialConditions[Counter][[3]]]][[3, 2]], {t, 0, tend}]},
   PlotRange → {-0.1, .1}, TicksStyle → Large, PlotStyle →
    Thickness[lThickness]],
  {Counter, 1, CounterEnd}]]
```

Out[ ]=

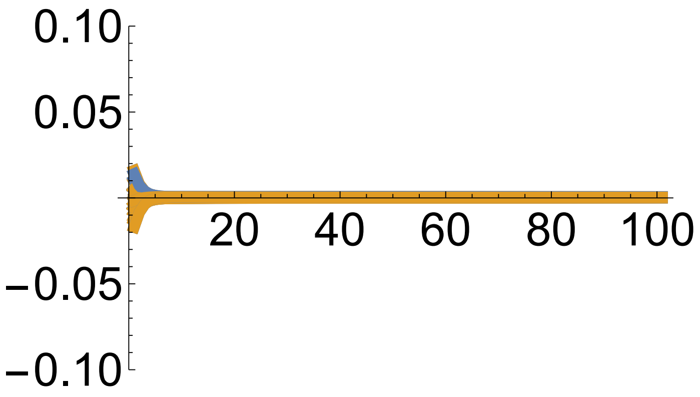

#### Example 4 approximated evo-devo dynamics: two biallelic loci, two phenotypes

In[142]:=

```
Clear["Global`*"]
```

In[143]:=

```
(*This is the frequency of haplotype ij*)
p[i_, j_, Xb_] := xb[1, i] × xb[2, j] + covx12[i, j, Xb]

(*This is the allele frequencies*)
xb[1, a] = 1 - x1b;
xb[1, A] = x1b;
xb[2, b] = 1 - x2b;
xb[2, B] = x2b;

(*This is the covariance in gene content between loci*)
covx12[a, b, Xb_] = Xb;
covx12[a, B, Xb_] = -Xb;
covx12[A, b, Xb_] = -Xb;
covx12[A, B, Xb_] = Xb;
```

In[152]:=

```
(*This is the genotype-phenotype map*)
z[m_, y1_, y2_] := cb (1 + e[m]) (y1 + y2) - 2 cb e[m] y1 y2
y[i_, m_] := c[i] (1 + d[i, m]) (xk[i] + xl[i]) - 2 c[i] × d[i, m] × xk[i] × xl[i]

(*These are the phenotypes evaluated at the mean haplotype
 content and at the chosen dominance and epistasis coefficients*)
zb[x1bvar_, x2bvar_] =
 FullSimplify[Table[z[m, y[1, m], y[2, m]] /. {cb → 1, c[1] → 1, c[2] → 1,
        d[1, 1] → 0, d[1, 2] → 0, d[2, 1] → 0, d[2, 2] → 0, e[1] → 0, e[2] → -1} /.
      {xk[1] → x1b, xk[2] → x2b} /. {xl[1] → x1b, xl[2] → x2b},
    {m, 1, 2}]] /. {x1b → x1bvar, x2b → x2bvar}
```

Out[154]=

```
{2 (x1bvar + x2bvar), 8 x1bvar x2bvar}
```

In[155]:=

```
(*The following confirms this by using the
 exact definition of z and evaluating it at the x bars*)
zExact[m_, k1_, k2_, l1_, l2_] :=
 cb (1 + e[m]) (yExact[1, A, k1, k2, l1, l2, m] + yExact[2, B, k1, k2, l1, l2, m]) -
  2 cb e[m] × yExact[1, A, k1, k2, l1, l2, m] × yExact[2, B, k1, k2, l1, l2, m]

yExact[i_, j_, k1_, k2_, l1_, l2_, m_] :=
 c[i] (1 + d[i, m]) (x[i, j, k1, k2] + x[i, j, l1, l2]) -
  2 c[i] × d[i, m] × x[i, j, k1, k2] × x[i, j, l1, l2]

Table[zExact[m, k1, k2, l1, l2] /. {xk[1] → x1b, xk[2] → x2b} /.
     {xl[1] → x1b, xl[2] → x2b}, {m, 1, 2}] /. {cb → 1, c[1] → 1, c[2] → 1,
    d[1, 1] → 0, d[1, 2] → 0, d[2, 1] → 0, d[2, 2] → 0, e[1] → 0, e[2] → -1} /.
 {x[1, A, k1, k2] → x1b, x[1, A, l1, l2] → x1b,
  x[2, B, k1, k2] → x2b, x[2, B, l1, l2] → x2b}
```

Out[157]=

$\{2\,x1b + 2\,x2b,\ 8\,x1b\,x2b\}$

In[158]:=

```
(*This is dz^\intercal/dx and checks that it is singular if x1b==
 x2b or e1==e2*)

dzdxSing =
  Table[Simplify[D[z[m, y[1, m], y[2, m]], xk[k]] /. {cb → 1, c[1] → 1, c[2] → 1,
        d[1, m] → 0, d[2, m] → 0} /. {xk[1] → x1b, xl[1] → x1b,
       xk[2] → x2b, xl[2] → x2b}], {k, 1, 3}, {m, 1, 2}].{{a}, {b}};
Reduce[{dzdxSing[[1, 1]] == 0, dzdxSing[[2, 1]] == 0}, {a, b}]
```

Out[159]=

$$(e[1] == e[2]\ \&\&\ x1b - x2b \neq 0\ \&\&\ b == -a)\ ||$$
$$((x1b - x2b)\,(e[1] - e[2]) \neq 0\ \&\&\ a == 0\ \&\&\ b == 0)\ ||\ \left(x1b == x2b\ \&\&\ -1 - e[2] + 4\,x2b\,e[2] \neq 0\ \&\&\ b == \frac{a + a\,e[1] - 4\,a\,x2b\,e[1]}{-1 - e[2] + 4\,x2b\,e[2]}\ \&\&\ e[1] - e[2] \neq 0\right)\ ||$$
$$(e[1] == e[2]\ \&\&\ x1b == x2b\ \&\&\ -1 - e[2] + 4\,x2b\,e[2] \neq 0\ \&\&\ b == -a)\ ||$$
$$\left(e[1] == e[2]\ \&\&\ e[2] \neq 0\ \&\&\ x2b == \frac{1 + e[2]}{4\,e[2]}\ \&\&\ x1b == x2b\right)\ ||$$
$$\left(e[2] \neq 0\ \&\&\ x2b == \frac{1 + e[2]}{4\,e[2]}\ \&\&\ x1b == x2b\ \&\&\ a == 0\ \&\&\ e[1] - e[2] \neq 0\right)$$

In[160]:=

```
(*This is dz^\intercal/dx for the particular
 dominance and epistasis coefficients used*)

dzdx[x1var_, x2var_] =
  Table[Simplify[D[z[m, y[1, m], y[2, m]], xk[k]] /. {xk[1] → x1b, xl[1] → x1b,
        xk[2] → x2b, xl[2] → x2b} /. {d[1, m] → 0, d[2, m] → 0} /.
      {cb → 1, c[1] → 1, c[2] → 1}], {k, 1, 3}, {m, 1, 2}] /.
   {e[1] → e1, e[2] → e2} /. {x1b → x1var, x2b → x2var};

MatrixForm[dzdx[x1b, x2b]]
MatrixForm[dzdx[x1b, x2b] /. {e1 → 0, e2 → -1}]
```

Out[161]//MatrixForm=

$$\begin{pmatrix} 1+e1-4\,e1\,x2b & 1+e2-4\,e2\,x2b \\ 1+e1-4\,e1\,x1b & 1+e2-4\,e2\,x1b \\ 0 & 0 \end{pmatrix}$$

Out[162]//MatrixForm=

$$\begin{pmatrix} 1 & 4\,x2b \\ 1 & 4\,x1b \\ 0 & 0 \end{pmatrix}$$

In[163]:=

```
(*This is fitness*)
W[z1_, z2_] := Exp[-(z1 - θ1)^2/σ1^2 - (z2 - θ2)^2/σ2^2]

(*These are the direct and total selection gradients*)
dwdz[z1b_, z2b_] = 2 {{1/σ1^2 (θ1 - z1b)}, {1/σ2^2 (θ2 - z2b)}};

dwdx[x1b_, x2b_] = dzdx[x1b, x2b].dwdz[zb[x1b, x2b][[1]], zb[x1b, x2b][[2]]] ;

(*This the total selection gradient
 of the haplotype at the epistasis chosen*)
dzdx[x1b, x2b].dwdz[z1b, z2b] /. {e1 → 0, e2 → -1}
```

Out[166]=

$$\left\{\left\{\frac{2\,(-z1b+\theta1)}{\sigma1^2}+\frac{8\,x2b\,(-z2b+\theta2)}{\sigma2^2}\right\},\left\{\frac{2\,(-z1b+\theta1)}{\sigma1^2}+\frac{8\,x1b\,(-z2b+\theta2)}{\sigma2^2}\right\},\{0\}\right\}$$

In[ ]:=
```
(*These are the equilibria arising from setting
 the total selection gradient of the haplotype to zero*)
Simplify[Reduce[{2 (-z1b + θ1)/σ1^2 + 8 x2b (-z2b + θ2)/σ2^2 == 0,
   2 (-z1b + θ1)/σ1^2 + 8 x1b (-z2b + θ2)/σ2^2 == 0}, {z1b, z2b}]]
```

Out[ ]=
```
(z1b == θ1 && σ1 σ2 ≠ 0 && ((x1b == 0 && x2b == 0) || (z2b == θ2 && x1b ≠ x2b))) ||
 (x1b == x2b && z2b == θ2 + ((-z1b + θ1) σ2^2)/(4 x2b σ1^2) && x2b σ1 ≠ 0 && σ2 ≠ 0)
```

In[167]:=
```
(*This is pure transmission bias, zetax - xb, as previously found*)
Deltaxvec[Xb_] = (0
                  0
                  -r Xb);

(*This is Tx as previously found*)
Tx[x1b_, x2b_, Xb_] = (-1/2 (-1 + x1b) x1b                 Xb/2
                       Xb/2                               -1/2 (-1 + x2b) x2b
                       1/2 (-1 + r) (-1 + 2 x1b) Xb  1/2 (-1 + r) (-1 + 2 x2b) Xb  -1/2 (-1 + r) (x[illegible]

(*This is the MAG covariance matrix of the phenotype, Lz*)
Lz[x1b_, x2b_, Xb_] := 2 Transpose[dzdx[x1b, x2b]].Tx[x1b, x2b, Xb].dzdx[x1b, x2b]
```

In[ ]:=
```
(*This is the determinant of Lz for the cs and ds used,
which shows that the determinant is proportional to(e1-e2)^2 (x1b-x2b)^2*)
Simplify[Det[Lz[x1b, x2b, Xb]]]
```

Out[ ]=
```
16 (e1 - e2)^2 (x1b - x2b)^2 (-x1b (-1 + x2b) x2b + x1b^2 (-1 + x2b) x2b - Xb^2)
```

In[ ]:=
```
(*This checks that Lz is symmetric*)
Simplify[Lz[x1b, x2b, Xb] == Transpose[Lz[x1b, x2b, Xb]]]
```

Out[ ]=
```
True
```

In[ ]:=
```
(*This is Lz for the coefficients used*)
Simplify[MatrixForm[Lz[x1b, x2b, Xb] /. {e1 → 0, e2 → -1}]]

(*This checks that the offdiagonal entry is as given in the main text*)
Simplify[
 Lz[x1b, x2b, Xb][[1, 2]] == 8 x1b x2b + 4 (x1b + x2b) (Xb - x1b x2b) /. {e1 → 0, e2 → -1}]
```

Out[ ]//MatrixForm=
```
(x1b - x1b^2 + x2b - x2b^2 + 2 Xb                              -4 x1b^2 x2b + 4 x2b Xb + x1b (8 x2b - 4 x2b[illegible]
 -4 x1b^2 x2b + 4 x2b Xb + x1b (8 x2b - 4 x2b^2 + 4 Xb)        16 x1b x2b (x1b + x2b - 2 x1b x2b + 2 [illegible]
```

Out[ ]=
```
True
```

In[170]:=

```
(*These are the haplotype frequencies as functions
 of allele frequencies and linkage disequilibrium*)
pp[i_, j_, x1bb_, x2bb_, Xbb_] := p[i, j, Xb] /. {x1b → x1bb, x2b → x2bb, Xb → Xbb}

(*This is the constraint that haplotype
 frequencies are between zero and one*)
Conditions[x1b_, x2b_, Xb_] :=
 0 ≤ pp[a, b, x1b, x2b, Xb] ≤ 1 && 0 ≤ pp[a, B, x1b, x2b, Xb] ≤ 1 &&
  0 ≤ pp[A, b, x1b, x2b, Xb] ≤ 1 && 0 ≤ pp[A, B, x1b, x2b, Xb] ≤ 1

(*These are the eigenvalues of Lz subject to the constraint
 that haplotype frequencies are between zero and one*)

EigC[x1b_, x2b_, Xb_] = If[Conditions[x1b, x2b, Xb],
   Eigenvalues[Lz[x1b, x2b, Xb] /. {e1 → 0, e2 → -1}], {NaN, NaN, NaN}];

Manipulate[Plot3D[{EigC[x1b, x2b, Xb][[1]], EigC[x1b, x2b, Xb][[2]]},
  {x1b, 0, 1}, {x2b, 0, 1}], {{Xb, 0}, -1, 1}]
```

Out[173]=

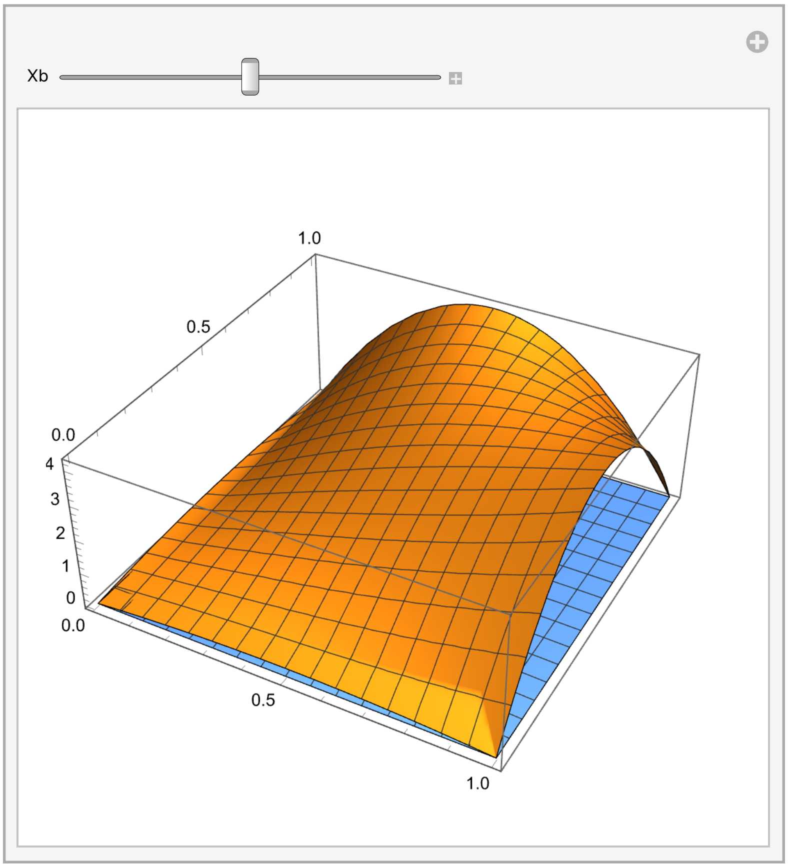

In[174]:=

```
(*This is Lm*)
Lm[x1bvar_, x2bvar_, Xbvar_] :=
 Am.Tx[x1b, x2b, Xb].BT /. {x1b → x1bvar, x2b → x2bvar, Xb → Xbvar}

(*This is A*)
Am := Join[2 Transpose[dzdx[x1b, x2b]], IdentityMatrix[3]]

(*This is B transpose*)
BT := Join[dzdx[x1b, x2b], IdentityMatrix[3], 2]
```

In[ ]:=

```
(*This checks that Lm is singular*)
Det[Lm[x1b, x2b, Xb]]
```

Out[ ]=

```
0
```

In[ ]:=

```
(*This checks that Lm is asymmetric*)
Simplify[Lm[x1b, x2b, Xb] == Transpose[Lm[x1b, x2b, Xb]] /. {e1 → 0, e2 → -1}]
```

Out[ ]=

$$\left\{\left\{0, 0, \frac{1}{2}\left(x1b - x1b^2 + Xb\right), \frac{1}{2}\left(x2b - x2b^2 + Xb\right), -((1+r)(-1+x1b+x2b)\,Xb)\right\},\right.$$
$$\{0, 0, 2\,x1b\,(x2b - x1b\,x2b + Xb), 2\,x2b\,(x1b - x1b\,x2b + Xb), -2\,(1+r)\,(-x2b + x1b\,(-1+4\,x2b))\,Xb\},$$
$$\left\{\frac{1}{2}\left(-x1b + x1b^2 - Xb\right), -2\,x1b\,(x2b - x1b\,x2b + Xb), 0, 0, \frac{1}{2}\,r\,(1-2\,x1b)\,Xb\right\},$$
$$\left\{\frac{1}{2}\left(-x2b + x2b^2 - Xb\right), -2\,x2b\,(x1b - x1b\,x2b + Xb), 0, 0, \frac{1}{2}\,r\,(1-2\,x2b)\,Xb\right\},$$
$$\left.\left\{(1+r)(-1+x1b+x2b)\,Xb, 2\,(1+r)\,(-x2b + x1b\,(-1+4\,x2b))\,Xb, \frac{1}{2}\,r\,(-1+2\,x1b)\,Xb, \frac{1}{2}\,r\,(-1+2\,x2b)\,Xb, 0\right\}\right\} ==$$
$$\{\{0, 0, 0, 0, 0\}, \{0, 0, 0, 0, 0\}, \{0, 0, 0, 0, 0\}, \{0, 0, 0, 0, 0\}, \{0, 0, 0, 0, 0\}\}$$

In[177]:=

```
(*These are the eigenvalues of Lm subject to the constraint
 that haplotype frequencies are between zero and one*)

EigCLm[x1b_, x2b_, Xb_, rr_] = If[Conditions[x1b, x2b, Xb], Eigenvalues[
    Lm[x1b, x2b, Xb] /. {e1 → 0, e2 → -1, r → rr}], {NaN, NaN, NaN, NaN, NaN}];
```

In[ ]:=
```
Manipulate[EigCLm[x1b, x2b, Xb, r], {{x1b, 0.138}, 0, 1},
 {{x2b, 0.246}, 0, 1}, {{Xb, 0.025}, -1, 1}, {{r, 0.355}, 0, 0.5}]
```

Out[ ]=

x1b

x2b

Xb

r

$\{0.636504, 0.0700729, 0.0088189, -6.01253\times10^{-17}, 9.55402\times10^{-19}\}$

In[178]:=
```
Manipulate[Plot3D[{EigCLm[x1b, x2b, Xb, r][[1]],
   EigCLm[x1b, x2b, Xb, r][[2]], EigCLm[x1b, x2b, Xb, r][[3]],
   EigCLm[x1b, x2b, Xb, r][[4]], EigCLm[x1b, x2b, Xb, r][[5]]},
  {x1b, 0, 1}, {x2b, 0, 1}], {{Xb, 0}, -1, 1}, {{r, 0.5}, 0, 0.5}]
```

Out[178]=

Xb

r

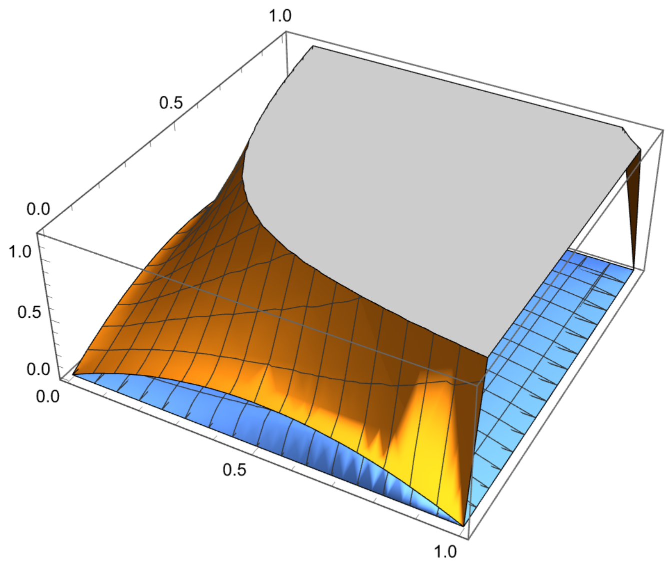

In[179]:=

```
(*These are the eigenvalues of the symmetric part of Lm subject to the
 constraint that haplotype frequencies are between zero and one*)

EigCSLm[x1b_, x2b_, Xb_, rr_] =
  If[0 ≤ pp[a, b, x1b, x2b, Xb] ≤ 1 && 0 ≤ pp[a, B, x1b, x2b, Xb] ≤ 1 &&
    0 ≤ pp[A, b, x1b, x2b, Xb] ≤ 1 && 0 ≤ pp[A, B, x1b, x2b, Xb] ≤ 1,
   Eigenvalues[1/2 (Lm[x1b, x2b, Xb] + Transpose[Lm[x1b, x2b, Xb]]) /.
     {e1 → 0, e2 → -1, r → rr}], {NaN, NaN, NaN, NaN, NaN}];
```

In[ ]:=

```
Manipulate[EigCSLm[x1b, x2b, Xb, r], {{x1b, 0.138}, 0, 1},
 {{x2b, 0.246}, 0, 1}, {{Xb, 0.025}, -1, 1}, {{r, 0.355}, 0, 0.5}]
```

Out[ ]=

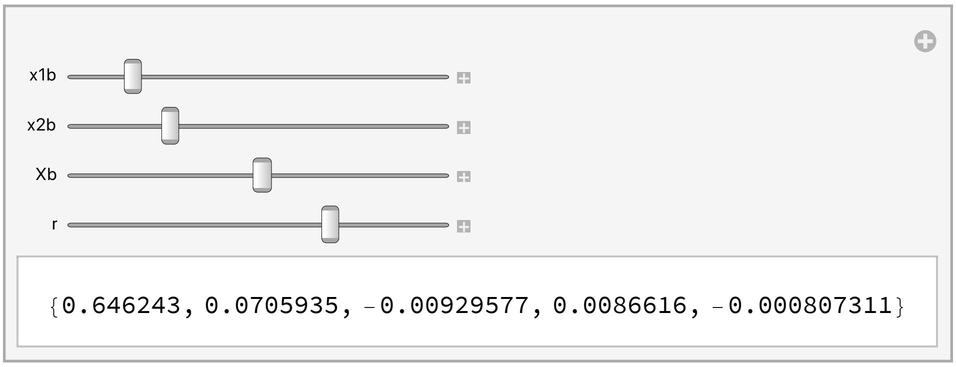

In[180]:=

```
Manipulate[Plot3D[{EigCSLm[x1b, x2b, Xb, r][[1]],
    EigCSLm[x1b, x2b, Xb, r][[2]], EigCSLm[x1b, x2b, Xb, r][[3]],
    EigCSLm[x1b, x2b, Xb, r][[4]], EigCSLm[x1b, x2b, Xb, r][[5]]},
   {x1b, 0, 1}, {x2b, 0, 1}], {{Xb, 0.025}, -1, 1}, {{r, 0.355}, 0, 0.5}]
```

Out[180]=

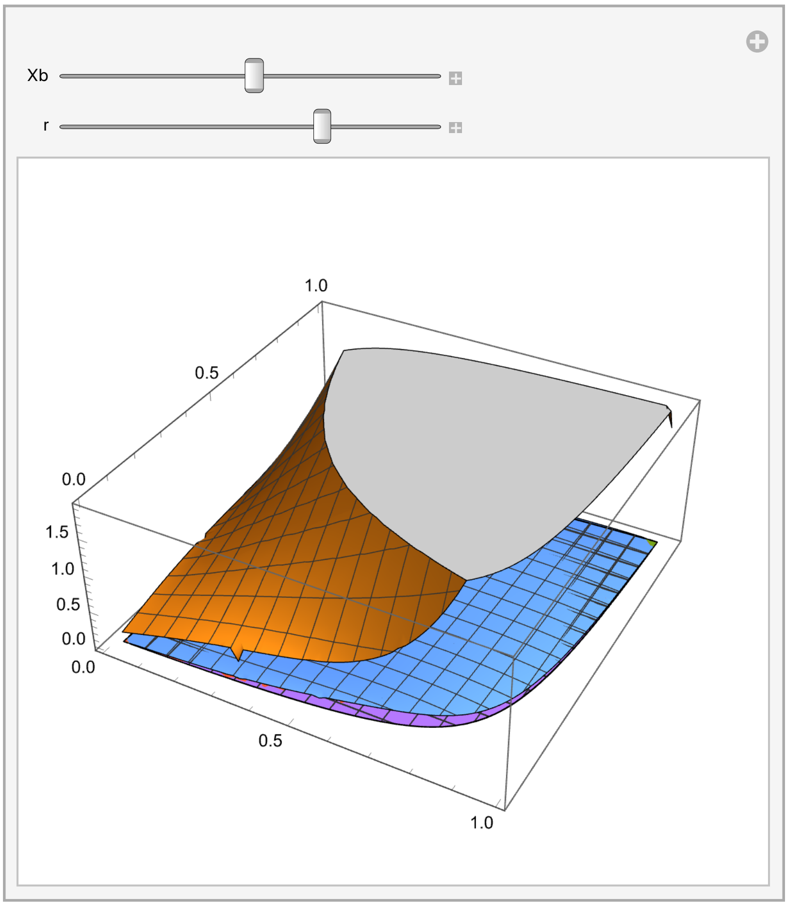

```
In[ ]:= (*These are the eigenvalues and eigenvectors of Lz for the
         chosen epistasis coefficients, written as function of allele
         frequencies and linkage disequilibrium to be used in plots*)
        Eigen[x1var_, x2var_, Xbvar_] =
          Simplify[Eigensystem[Lz[x1b, x2b, Xb] /. {e1 → 0, e2 → -1}]] /.
           {x1b → x1var, x2b → x2var, Xb → Xbvar};
        EigenValues[x1b_, x2b_, Xb_] = Eigen[x1b, x2b, Xb][[1]];
        EigenVectors[x1b_, x2b_, Xb_] = Eigen[x1b, x2b, Xb][[2]];

        (*These are the angles between the
         eigenvectors of Lz and the selection gradient*)
        Angle1[x1var_, x2var_, Xbvar_] = VectorAngle[EigenVectors[x1b, x2b, Xb][[1]],
            Flatten[dwdz[zb[x1b, x2b][[1]], zb[x1b, x2b][[2]]]]] /.
           {x1b → x1var, x2b → x2var, Xb → Xbvar};
        Angle2[x1var_, x2var_, Xbvar_] = VectorAngle[EigenVectors[x1b, x2b, Xb][[2]],
            Flatten[dwdz[zb[x1b, x2b][[1]], zb[x1b, x2b][[2]]]]] /.
           {x1b → x1var, x2b → x2var, Xb → Xbvar};

        (*This is Px = cov[x,x], as previously found*)
                                   ( -((-1 + x1b) x1b)          Xb
        Px[x1b_, x2b_, Xb_] =      (        Xb           -((-1 + x2b) x2b)
                                   (   Xb - 2 x1b Xb       Xb - 2 x2b Xb     x1b^2 (-1 + x2b) x2b - Xb (-1 +
```

In[ ]:=
```
(*This runs the numerical solutions*)

r = 0.5;

cb = 1;
c[1] = 1;
c[2] = 1;
d[1, 1] = 0;
d[1, 2] = 0;
d[2, 1] = 0;
d[2, 2] = 0;
e1 = 0;
e2 = -1;

θ1 = 2;
θ2 = 4;
σ1 = Sqrt[10];
σ2 = Sqrt[40];

tend = 100;

(*Initial conditions must be such that
 initial gamete frequencies are between zero and one*)
x1init = {0.01, 1, .2};
x2init = {0.01, 1, .2};
Xbinit = {-0.99, 1, .1};

(*pSol[i_,j_,x1bvar_,x2bvar_,Xbvar_]:=
  p[i,j,Xb]/.{x1b→x1bvar,x2b→x2bvar,Xb→Xbvar};*)

(*If the initial allele frequencies and linkage disequilibrium for which
 haplotype frequences are between zero and one, the solution is run*)
Table[If[Conditions[i, j, k], {x1bSol[0, i, j, k] = i,
    x2bSol[0, i, j, k] = j, XbSol[0, i, j, k] = k, xbvecSol[0, i, j, k] =
     {{x1bSol[0, i, j, k]}, {x2bSol[0, i, j, k]}, {XbSol[0, i, j, k]}},
    Table[{xbvecSol[t + 1, i, j, k] = xbvecSol[t, i, j, k] +
        Tx[x1bSol[t, i, j, k], x2bSol[t, i, j, k], XbSol[t, i, j, k]].dwdx[
          x1bSol[t, i, j, k], x2bSol[t, i, j, k]] + Deltaxvec[XbSol[t, i, j, k]],
      x1bSol[t + 1, i, j, k] = xbvecSol[t + 1, i, j, k]⟦1, 1⟧,
      x2bSol[t + 1, i, j, k] = xbvecSol[t + 1, i, j, k]⟦2, 1⟧,
      XbSol[t + 1, i, j, k] = xbvecSol[t + 1, i, j, k]⟦3, 1⟧}, {t, 0, tend}]}],
  {i, x1init⟦1⟧, x1init⟦2⟧, x1init⟦3⟧},
  {j, x2init⟦1⟧,
   x2init⟦2⟧, x2init⟦3⟧},
  {k, Xbinit⟦1⟧, Xbinit⟦2⟧, Xbinit⟦3⟧}];
```

In[ ]:=
```
(*This counts the number of initial conditions that meet the
 criterion that the haplotype frequencies are between 0 and 1*)
Counter = 0;
Table[If[Conditions[i, j, k], {Counter = Counter + 1;
    ActualInitialConditions[Counter] = {i, j, k}}],
  {i, x1init⟦1⟧, x1init⟦2⟧, x1init⟦3⟧}, {j, x2init⟦1⟧, x2init⟦2⟧, x2init⟦3⟧},
  {k, Xbinit⟦1⟧, Xbinit⟦2⟧, Xbinit⟦3⟧}];
CounterEnd = Counter
```

Out[ ]=
```
40
```

In[ ]:=
```
(*This calculates the final points for the plots*)
pointSize = 0.05;

endPointsxP = Table[{x1bSol[tend, ActualInitialConditions[Counter]⟦1⟧,
     ActualInitialConditions[Counter]⟦2⟧, ActualInitialConditions[Counter]⟦3⟧],
    x2bSol[tend, ActualInitialConditions[Counter]⟦1⟧,
     ActualInitialConditions[Counter]⟦2⟧,
     ActualInitialConditions[Counter]⟦3⟧]}, {Counter, 1, CounterEnd}];
```

In[ ]:=
```
yAxisLimitz := 1.1
yAxisLimitP := .6

lThickness = 0.02; (*Line thickness*)

(*This plots the change of allele frequencies
 for both loci on top of the mean fitness contour*)
plot = Show[ContourPlot[W[zb[x1b, x2b]⟦1⟧, zb[x1b, x2b]⟦2⟧],
   {x1b, 0, 1}, {x2b, 0, 1}, FrameStyle → Large, PlotLegends → Automatic,
   LabelStyle → Large, TicksStyle → Large, PlotRange → {{-0.1, 1.1}, {-.1, 1.1}},
   Epilog → {PointSize[pointSize], Point[endPointsxP]}],
  Table[ListLinePlot[Table[{x1bSol[t, ActualInitialConditions[Counter]⟦1⟧,
       ActualInitialConditions[Counter]⟦2⟧,
       ActualInitialConditions[Counter]⟦3⟧], x2bSol[t,
       ActualInitialConditions[Counter]⟦1⟧, ActualInitialConditions[Counter]⟦
        2⟧, ActualInitialConditions[Counter]⟦3⟧]}, {t, 0, tend}],
    PlotStyle → Thickness[lThickness]], {Counter, 1, CounterEnd}]]
(*Rasterize the plot to eliminate vectorization artifacts*)
plot = Rasterize[plot, "Image", ImageResolution → 300];
Export[StringJoin[ToString[NotebookDirectory[]], "Fig.5i", ".pdf"], plot];

(*This shows the change in allele frequencies
 in both loci and in linkage disquilibrium over time*)
Show[Table[ListLinePlot[{Table[x1bSol[t,
      ActualInitialConditions[Counter]⟦1⟧, ActualInitialConditions[Counter]⟦2⟧,
      ActualInitialConditions[Counter]⟦3⟧], {t, 0, tend}],
    Table[x2bSol[t, ActualInitialConditions[Counter]⟦1⟧,
```

```
      ActualInitialConditions[Counter]⟦2⟧,
      ActualInitialConditions[Counter]⟦3⟧], {t, 0, tend}],
    Table[XbSol[t, ActualInitialConditions[Counter]⟦1⟧,
      ActualInitialConditions[Counter]⟦2⟧,
      ActualInitialConditions[Counter]⟦3⟧], {t, 0, tend}]},
   PlotRange → {-0.2, yAxisLimitz}, TicksStyle → Large,
   PlotStyle → Thickness[lThickness]],
  {Counter, 1, CounterEnd}]]
Export[StringJoin[ToString[NotebookDirectory[]], "Fig.5k", ".pdf"], %];

(*This plots the change in mean absolute fitness over time*)
Show[
 Table[ListLinePlot[{Table[W[(zb[x1bSol[t, ActualInitialConditions[Counter]⟦1⟧,
         ActualInitialConditions[Counter]⟦2⟧,
         ActualInitialConditions[Counter]⟦3⟧], x2bSol[t,
         ActualInitialConditions[Counter]⟦1⟧, ActualInitialConditions[
           Counter]⟦2⟧, ActualInitialConditions[Counter]⟦3⟧]])⟦1⟧,
      (zb[x1bSol[t, ActualInitialConditions[Counter]⟦1⟧,
         ActualInitialConditions[Counter]⟦2⟧,
         ActualInitialConditions[Counter]⟦3⟧], x2bSol[t,
         ActualInitialConditions[Counter]⟦1⟧, ActualInitialConditions[
           Counter]⟦2⟧, ActualInitialConditions[Counter]⟦3⟧]])⟦2⟧],
     {t, 0, tend}]}, PlotRange → {0, yAxisLimitz}, TicksStyle → Large,
   PlotStyle → Thickness[lThickness]], {Counter, 1,
   CounterEnd}]]
Export[StringJoin[ToString[NotebookDirectory[]], "Fig.5m", ".pdf"], %];
```

Out[ ]=

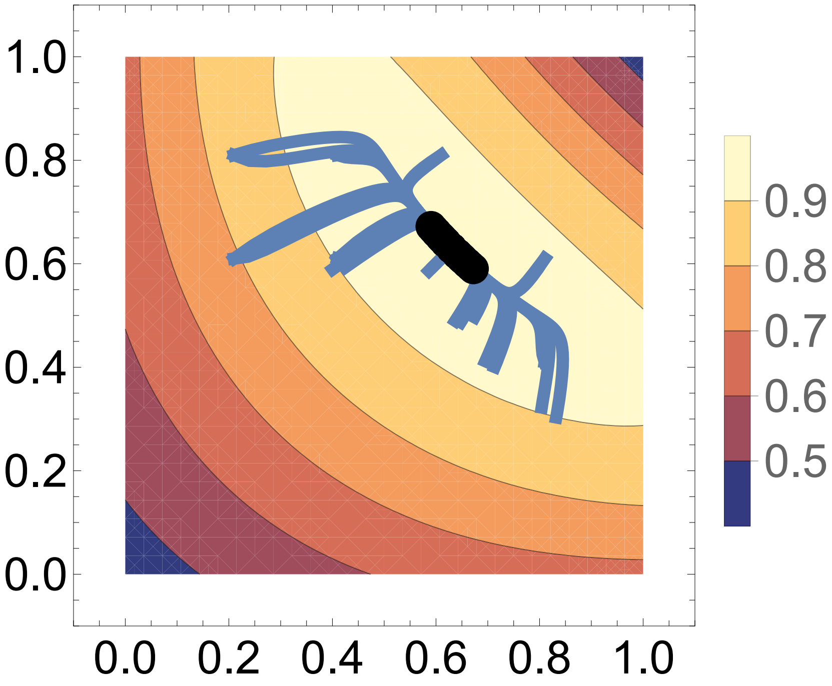

Out[ ]=

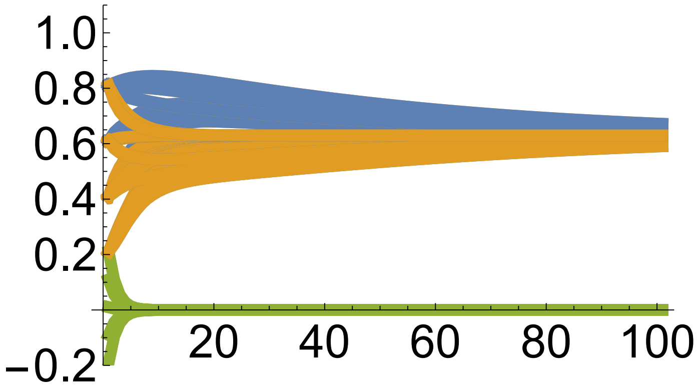

Out[ ]=

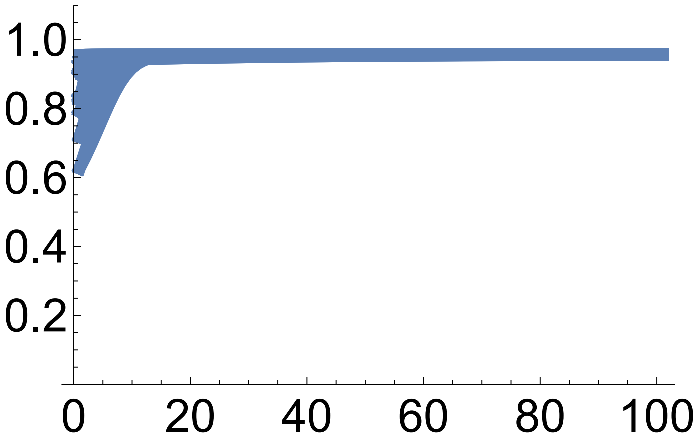

In[ ]:=
```
(*This plots mean z1 (blue) and mean z2 (orange) over time*)
Show[
 Table[ListLinePlot[{Table[zb[x1bSol[t, ActualInitialConditions[Counter][[1]],
        ActualInitialConditions[Counter][[2]],
        ActualInitialConditions[Counter][[3]]], x2bSol[t,
        ActualInitialConditions[Counter][[1]], ActualInitialConditions[Counter][[
         2]], ActualInitialConditions[Counter][[3]]]][[1]], {t, 0, tend}],
    Table[zb[x1bSol[t, ActualInitialConditions[Counter][[1]],
        ActualInitialConditions[Counter][[2]],
        ActualInitialConditions[Counter][[3]]], x2bSol[t,
        ActualInitialConditions[Counter][[1]], ActualInitialConditions[Counter][[
         2]], ActualInitialConditions[Counter][[3]]]][[2]], {t, 0, tend}],
    Table[θ1, {t, 0, tend}], Table[θ2, {t, 0, tend}]}, PlotRange →
    {0, 6}, TicksStyle → Large,
   PlotStyle → {Thickness[lThickness], Thickness[lThickness],
     {Magenta, Thickness[lThickness]},
     {Magenta, Thickness[lThickness]}}], {Counter, 1, CounterEnd}]]
Export[StringJoin[ToString[NotebookDirectory[]], "Fig.5l", ".pdf"], %];
```

Out[ ]=

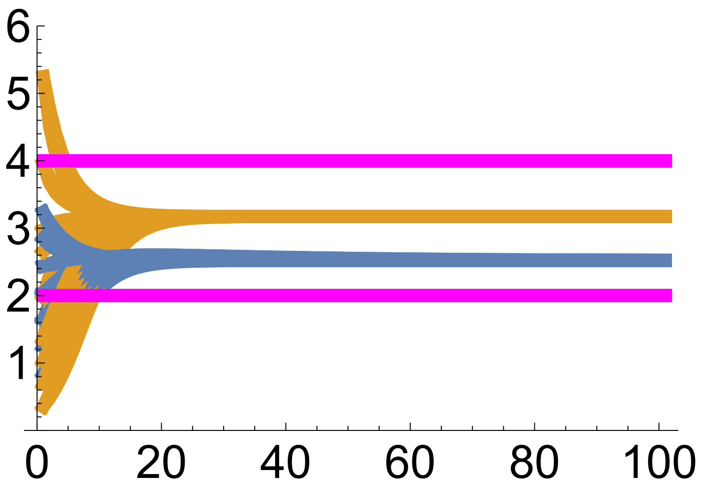

In[ ]:=
```
(*This computes the final points for the variance plots*)

endPointszP = Table[{zb[x1bSol[tend, ActualInitialConditions[Counter][[1]],
       ActualInitialConditions[Counter][[2]],
       ActualInitialConditions[Counter][[3]]], x2bSol[tend,
       ActualInitialConditions[Counter][[1]], ActualInitialConditions[Counter][[
        2]], ActualInitialConditions[Counter][[3]]]][[1]],
    zb[x1bSol[tend, ActualInitialConditions[Counter][[1]],
       ActualInitialConditions[Counter][[2]],
       ActualInitialConditions[Counter][[3]]],
      x2bSol[tend, ActualInitialConditions[Counter][[1]],
       ActualInitialConditions[Counter][[2]],
       ActualInitialConditions[Counter][[3]]]][[2]]}, {Counter, 1, CounterEnd}];

(*This plots mean z1 vs mean z2 with
 admissible developmental manifold in red*)
```

```
(*First, determine the initial and final
 admissible manifolds for a particular run*)
Resol = 0.009;
Table[Table[{PointsSol[t] = {{0, 0}};,
    Table[If[Conditions[x1b, x2b, Xb], PointsSol[t] =
          Append[PointsSol[t], {zb[x1b, x2b]⟦1⟧, zb[x1b, x2b]⟦2⟧}];] /.
       Xb → XbSol[t, ActualInitialConditions[Counter]⟦1⟧,
         ActualInitialConditions[Counter]⟦2⟧,
         ActualInitialConditions[Counter]⟦3⟧],
      {x1b, 0, 1, Resol}, {x2b, 0, 1, Resol}];},
   {t, {0, tend}}], {Counter, {CounterEnd}}];

(*Then, do the plot*)
plot = Show[ContourPlot[W[z1b, z2b], {z1b, 0, 4},
   {z2b, 0, 8}, FrameStyle → Large, PlotLegends → Automatic,
   LabelStyle → Large, TicksStyle → Large, PlotRange → {{0, 4}, {0, 8}},
   Epilog → {{PointSize[pointSize], Point[endPointszP]},
     {PointSize[pointSize], Magenta, Point[{θ1, θ2}]}}],
  Table[Table[{
     ListPlot[PointsSol[t], AspectRatio → 1, PlotStyle → {Red, PointSize[Medium]},
      PlotRange → {{0, 4}, {0, 8}}, TicksStyle → Large]},
    {t, {0, tend}}], {Counter, {CounterEnd}}],
  Table[ListLinePlot[{Table[{zb[x1bSol[t, ActualInitialConditions[Counter]⟦1⟧,
          ActualInitialConditions[Counter]⟦2⟧,
          ActualInitialConditions[Counter]⟦3⟧], x2bSol[t,
          ActualInitialConditions[Counter]⟦1⟧, ActualInitialConditions[
            Counter]⟦2⟧, ActualInitialConditions[Counter]⟦3⟧]]⟦1⟧,
       zb[x1bSol[t, ActualInitialConditions[Counter]⟦1⟧,
          ActualInitialConditions[Counter]⟦2⟧,
          ActualInitialConditions[Counter]⟦3⟧], x2bSol[t,
          ActualInitialConditions[Counter]⟦1⟧, ActualInitialConditions[
            Counter]⟦2⟧, ActualInitialConditions[Counter]⟦3⟧]]⟦2⟧},
      {t, 0, tend}]}, PlotRange → {{-0.1, 4.1}, {-0.1, 6.1}},
    PlotStyle → Thickness[0.5 lThickness]], {Counter,
    1,
    CounterEnd}]]
(*Rasterize the plot to eliminate vectorization artifacts*)
plot = Rasterize[plot, "Image", ImageResolution → 300];
Export[StringJoin[ToString[NotebookDirectory[]], "Fig.5j", ".pdf"], plot];

(*This plots, over time, the MAG covariance of phenotype 1 (blue),
the MAG covariance of phenotype 2 (orange),
and the MAG covariance of phenotype 1 and 2 (green)*)
Show[
 Table[ListLinePlot[{Table[Lz[x1bSol[t, ActualInitialConditions[Counter]⟦1⟧,
        ActualInitialConditions[Counter]⟦2⟧,
```

```
ActualInitialConditions[Counter][[3]], x2bSol[t,
ActualInitialConditions[Counter][[1]], ActualInitialConditions[Counter][[
 2]], ActualInitialConditions[Counter][[3]]], XbSol[t,
ActualInitialConditions[Counter][[1]], ActualInitialConditions[Counter][[
 2]], ActualInitialConditions[Counter][[3]]]][[1, 1]], {t, 0, tend}],
Table[Lz[x1bSol[t, ActualInitialConditions[Counter][[1]],
  ActualInitialConditions[Counter][[2]],
  ActualInitialConditions[Counter][[3]], x2bSol[t,
  ActualInitialConditions[Counter][[1]], ActualInitialConditions[Counter][[
   2]], ActualInitialConditions[Counter][[3]]], XbSol[t,
  ActualInitialConditions[Counter][[1]], ActualInitialConditions[Counter][[
   2]], ActualInitialConditions[Counter][[3]]]][[2, 2]], {t, 0, tend}],
Table[Lz[x1bSol[t, ActualInitialConditions[Counter][[1]],
  ActualInitialConditions[Counter][[2]],
  ActualInitialConditions[Counter][[3]], x2bSol[t,
  ActualInitialConditions[Counter][[1]], ActualInitialConditions[Counter][[
   2]], ActualInitialConditions[Counter][[3]]], XbSol[t,
  ActualInitialConditions[Counter][[1]], ActualInitialConditions[Counter][[
   2]], ActualInitialConditions[Counter][[3]]]][[2, 1]], {t, 0, tend}]},
 PlotRange → {0, 6}, TicksStyle → Large, PlotStyle →
  Thickness[lThickness]],
 {Counter, 1, CounterEnd}]]
Export[StringJoin[ToString[NotebookDirectory[]], "Fig.5p", ".pdf"], %];
```

Out[ ]=

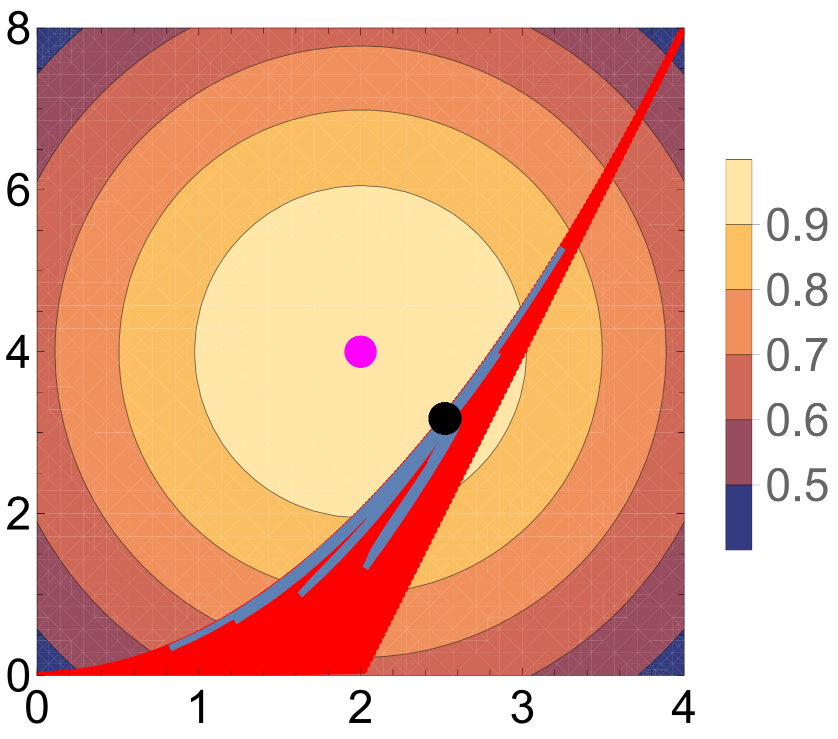

Out[ ]=

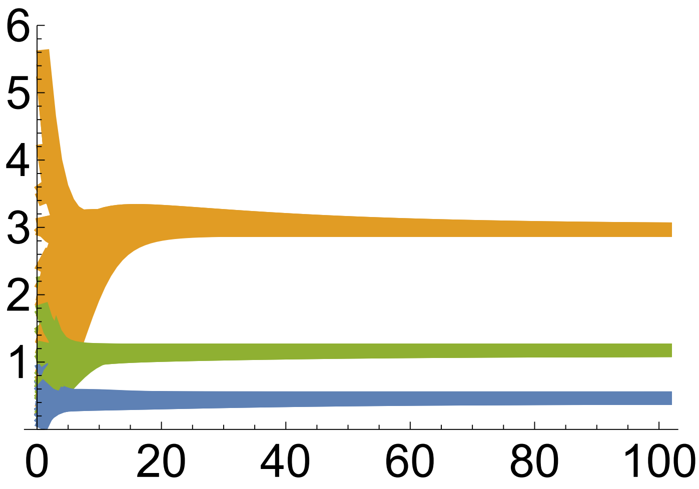

In[ ]:=
```
(*This plots, over time, the allelic variance in locus 1 for allele A (blue),
the allelic variance in locus 2 for allele B (orange),
and the variance in the squared deviation
 in gene content between alleles A and B (green)*)

Show[
 Table[ListLinePlot[{Table[Px[x1bSol[t, ActualInitialConditions[Counter][[1]],
        ActualInitialConditions[Counter][[2]],
        ActualInitialConditions[Counter][[3]]], x2bSol[t,
        ActualInitialConditions[Counter][[1]], ActualInitialConditions[Counter][[
         2]], ActualInitialConditions[Counter][[3]]], XbSol[t,
        ActualInitialConditions[Counter][[1]], ActualInitialConditions[Counter][[
         2]], ActualInitialConditions[Counter][[3]]]][[1, 1]], {t, 0, tend}],
    Table[Px[x1bSol[t, ActualInitialConditions[Counter][[1]],
        ActualInitialConditions[Counter][[2]],
        ActualInitialConditions[Counter][[3]]], x2bSol[t,
        ActualInitialConditions[Counter][[1]], ActualInitialConditions[Counter][[
         2]], ActualInitialConditions[Counter][[3]]], XbSol[t,
        ActualInitialConditions[Counter][[1]], ActualInitialConditions[Counter][[
         2]], ActualInitialConditions[Counter][[3]]]][[2, 2]], {t, 0, tend}],
    Table[Px[x1bSol[t, ActualInitialConditions[Counter][[1]],
        ActualInitialConditions[Counter][[2]],
        ActualInitialConditions[Counter][[3]]], x2bSol[t,
        ActualInitialConditions[Counter][[1]], ActualInitialConditions[Counter][[
         2]], ActualInitialConditions[Counter][[3]]], XbSol[t,
        ActualInitialConditions[Counter][[1]], ActualInitialConditions[Counter][[
         2]], ActualInitialConditions[Counter][[3]]]][[3, 3]], {t, 0, tend}]},
   PlotRange → {0, .3}, TicksStyle → Large, PlotStyle →
    Thickness[lThickness]],
  {Counter, 1, CounterEnd}]]
Export[StringJoin[ToString[NotebookDirectory[]], "Fig.5o", ".pdf"], %];

(*This plots, over time, the covariance between allele A
```

```
 and squared deviation in gene content (blue) and the covariance
 between allele B and squared deviation in gene content (orange)*)
Show[
 Table[ListLinePlot[{Table[Px[x1bSol[t, ActualInitialConditions[Counter]⟦1⟧,
        ActualInitialConditions[Counter]⟦2⟧,
        ActualInitialConditions[Counter]⟦3⟧], x2bSol[t,
        ActualInitialConditions[Counter]⟦1⟧, ActualInitialConditions[Counter]⟦
         2⟧, ActualInitialConditions[Counter]⟦3⟧], XbSol[t,
        ActualInitialConditions[Counter]⟦1⟧, ActualInitialConditions[Counter]⟦
         2⟧, ActualInitialConditions[Counter]⟦3⟧]]⟦1, 3⟧, {t, 0, tend}],
    Table[Px[x1bSol[t, ActualInitialConditions[Counter]⟦1⟧,
        ActualInitialConditions[Counter]⟦2⟧,
        ActualInitialConditions[Counter]⟦3⟧], x2bSol[t,
        ActualInitialConditions[Counter]⟦1⟧, ActualInitialConditions[Counter]⟦
         2⟧, ActualInitialConditions[Counter]⟦3⟧], XbSol[t,
        ActualInitialConditions[Counter]⟦1⟧, ActualInitialConditions[Counter]⟦
         2⟧, ActualInitialConditions[Counter]⟦3⟧]]⟦2, 3⟧, {t, 0, tend}]},
   PlotRange → {-0.1, .1}, TicksStyle → Large, PlotStyle →
    Thickness[lThickness]],
  {Counter, 1, CounterEnd}]]
Export[StringJoin[ToString[NotebookDirectory[]], "Fig.5n", ".pdf"], %];
```

Out[ ]=

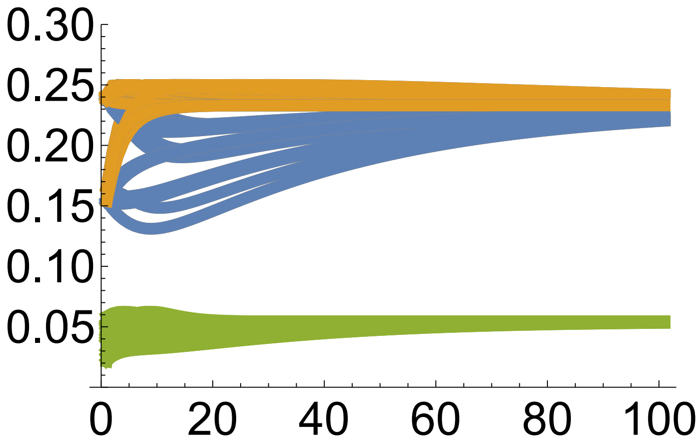

Out[ ]=

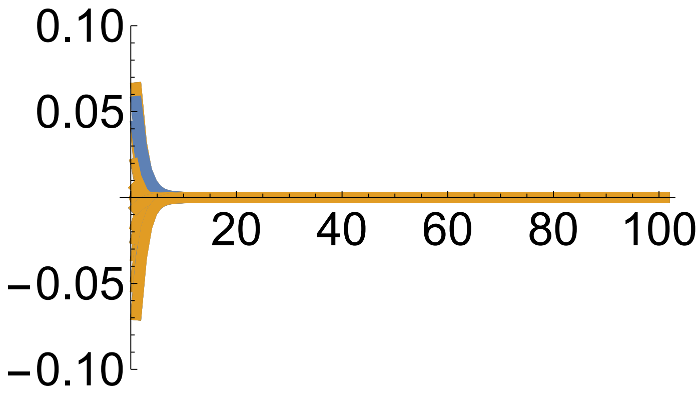

In[ ]:=
```
(*This plots, over time,
the covariance between offspring squared deviation in gene content and parent
 gene content for A (blue) and the covariance between offspring squared
 deviation in gene content and parent gene content for B (orange)*)
Show[
 Table[ListLinePlot[{Table[Tx[x1bSol[t, ActualInitialConditions[Counter][[1]],
       ActualInitialConditions[Counter][[2]],
       ActualInitialConditions[Counter][[3]]], x2bSol[t,
       ActualInitialConditions[Counter][[1]], ActualInitialConditions[Counter][[
        2]], ActualInitialConditions[Counter][[3]]], XbSol[t,
       ActualInitialConditions[Counter][[1]], ActualInitialConditions[Counter][[
        2]], ActualInitialConditions[Counter][[3]]]][[3, 1]], {t, 0, tend}],
    Table[Tx[x1bSol[t, ActualInitialConditions[Counter][[1]],
       ActualInitialConditions[Counter][[2]],
       ActualInitialConditions[Counter][[3]]], x2bSol[t,
       ActualInitialConditions[Counter][[1]], ActualInitialConditions[Counter][[
        2]], ActualInitialConditions[Counter][[3]]], XbSol[t,
       ActualInitialConditions[Counter][[1]], ActualInitialConditions[Counter][[
        2]], ActualInitialConditions[Counter][[3]]]][[3, 2]], {t, 0, tend}]},
   PlotRange → {-0.1, .1}, TicksStyle → Large, PlotStyle →
    Thickness[lThickness]],
  {Counter, 1, CounterEnd}]]
```

Out[ ]=

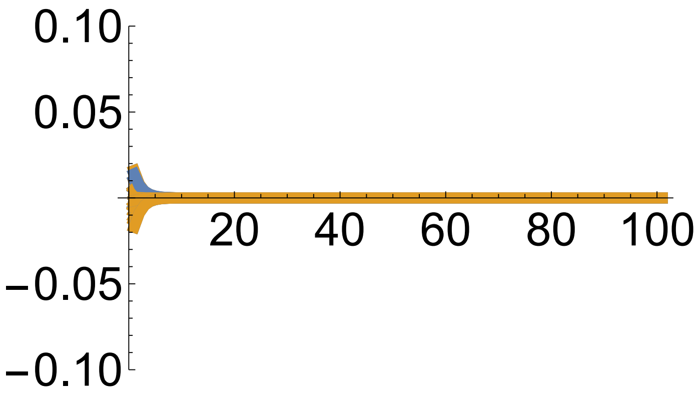

In[ ]:=
```
(*This plots the eigenvalues of Lz*)
Show[Table[ListLinePlot[
   {Table[EigenValues[x1bSol[t, ActualInitialConditions[Counter][[1]],
       ActualInitialConditions[Counter][[2]],
       ActualInitialConditions[Counter][[3]]], x2bSol[t,
       ActualInitialConditions[Counter][[1]], ActualInitialConditions[Counter][[
        2]], ActualInitialConditions[Counter][[3]]], XbSol[t,
       ActualInitialConditions[Counter][[1]], ActualInitialConditions[Counter][[
        2]], ActualInitialConditions[Counter][[3]]]][[1]], {t, 0, tend}],
    Table[EigenValues[x1bSol[t, ActualInitialConditions[Counter][[1]],
       ActualInitialConditions[Counter][[2]],
       ActualInitialConditions[Counter][[3]]], x2bSol[t,
       ActualInitialConditions[Counter][[1]], ActualInitialConditions[Counter][[
```

```
          2]], ActualInitialConditions[Counter][[3]]], XbSol[t,
        ActualInitialConditions[Counter][[1]], ActualInitialConditions[Counter][[
          2]], ActualInitialConditions[Counter][[3]]]][[2]], {t, 0, tend}]},
    PlotRange → {-0.01, 7}, TicksStyle → Large, PlotStyle →
     Thickness[lThickness]],
   {Counter, 1, CounterEnd}]]

Export[StringJoin[ToString[NotebookDirectory[]], "Fig.5q", ".pdf"], %];

(*This plots the angle between the
 eigenvalues of Lz and the selection gradient*)
Show[Table[
  ListLinePlot[{Table[180/π Angle1[x1bSol[t, ActualInitialConditions[Counter][[1]],
        ActualInitialConditions[Counter][[2]],
        ActualInitialConditions[Counter][[3]]], x2bSol[t,
        ActualInitialConditions[Counter][[1]], ActualInitialConditions[Counter][[
          2]], ActualInitialConditions[Counter][[3]]], XbSol[t,
        ActualInitialConditions[Counter][[1]], ActualInitialConditions[Counter][[
          2]], ActualInitialConditions[Counter][[3]]]], {t, 0, tend}],
     Table[180/π Angle2[x1bSol[t, ActualInitialConditions[Counter][[1]],
        ActualInitialConditions[Counter][[2]],
        ActualInitialConditions[Counter][[3]]], x2bSol[t,
        ActualInitialConditions[Counter][[1]], ActualInitialConditions[Counter][[
          2]], ActualInitialConditions[Counter][[3]]], XbSol[t,
        ActualInitialConditions[Counter][[1]], ActualInitialConditions[Counter][[
          2]], ActualInitialConditions[Counter][[3]]]], {t, 0, tend}]},
    PlotRange → {-0.2, 125}, TicksStyle → Large, PlotStyle →
     Thickness[lThickness]],
   {Counter, 1, CounterEnd}]]

Export[StringJoin[ToString[NotebookDirectory[]], "Fig.5r", ".pdf"], %];
```

Out[ ]=

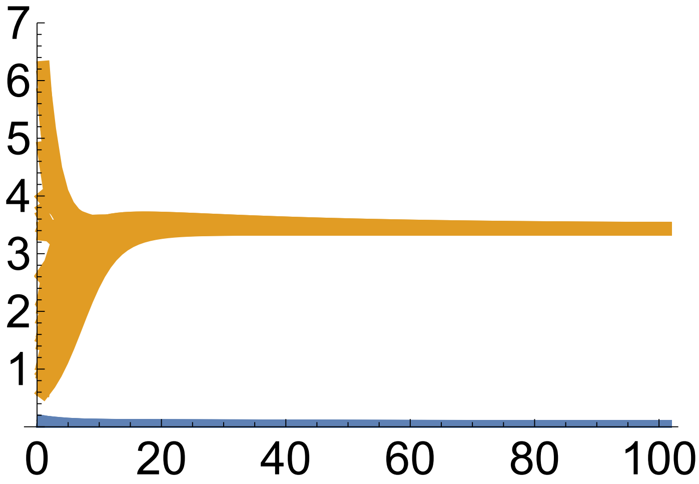

Out[ ]=

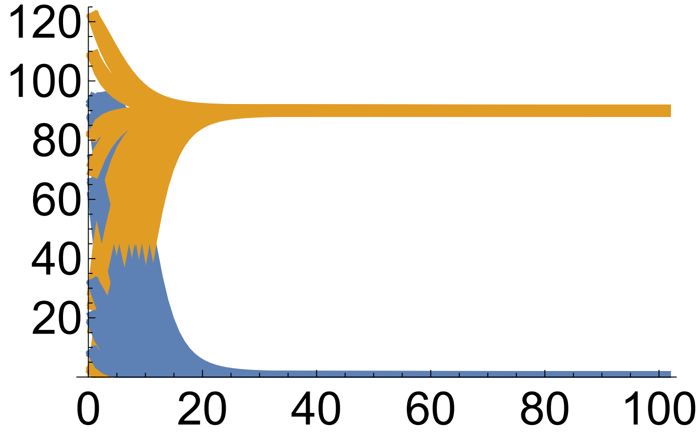

#### Example 4 continued: haplotype frequency implementation

In[ ]:=
```
Clear["Global`*"]
```

In[ ]:=
```
(*This defines the content of haplotype i in haplotype k*)
x[i_, k_] := KroneckerDelta[ToString[i], ToString[k]]

(*This defines haplotype frequency*)
p[i_] := xb[i]
```

In[ ]:=
```
(*This defines xvec[i,j], which is vector of haplotype content*)
xvec[i_] := {{x[i, aB]}, {x[i, Ab]}, {x[i, AB]}};

(*This is the vector of mean haplotype content*)
xbvec = {{xb[aB]}, {xb[Ab]}, {xb[AB]}};

(*This writes the vector of mean haplotype
 content as a function of the evolving traits*)
xbFun[xaBbvar_, xAbbvar_, xABbvar_] = {{xb[aB]}, {xb[Ab]}, {xb[AB]}} /.
   {xb[aB] → xaBbvar, xb[Ab] → xAbbvar, xb[AB] → xABbvar};
```

In[ ]:=
```
Table[MatrixForm[xvec[i]], {i, {aB, Ab, AB}}]
```

Out[ ]=

$$\left\{\begin{pmatrix}1\\0\\0\end{pmatrix}, \begin{pmatrix}0\\1\\0\end{pmatrix}, \begin{pmatrix}0\\0\\1\end{pmatrix}\right\}$$

In[ ]:=
```
(*This is Px, that is, cov[x,x], the covariance matrix of haplotype content*)

Px = Simplify[Sum[p[i] (xvec[i] - xbvec).Transpose[(xvec[i] - xbvec)],
      {i, {ab, aB, Ab, AB}}] /. {xb[ab] → 1 - xb[aB] - xb[Ab] - xb[AB]}] /.
   {xb[aB] → xaBb, xb[Ab] → xAbb, xb[AB] → xABb};
MatrixForm[Px]
PxFun[xaBbvar_, xAbbvar_, xABbvar_] =
   Px /. {xaBb → xaBbvar, xAbb → xAbbvar, xABb → xABbvar};
```

Out[ ]//MatrixForm=

$$\begin{pmatrix} -((-1+\text{xaBb})\ \text{xaBb}) & -\text{xaBb xAbb} & -\text{xaBb xABb} \\ -\text{xaBb xAbb} & -((-1+\text{xAbb})\ \text{xAbb}) & -\text{xAbb xABb} \\ -\text{xaBb xABb} & -\text{xAbb xABb} & -((-1+\text{xABb})\ \text{xABb}) \end{pmatrix}$$

In[ ]:=
```
(*Probability that genotype k1k2 l1l2 produces gametes with haplotype n1n2*)

R[k1_, k2_, l1_, l2_, n1_, n2_] := 1/2 KroneckerDelta[ToString[k1], ToString[n1]]
   ((1 - r) KroneckerDelta[ToString[k2], ToString[n2]] +
     r KroneckerDelta[ToString[l2], ToString[n2]]) +
  1/2 KroneckerDelta[ToString[l1], ToString[n1]]
   ((1 - r) KroneckerDelta[ToString[l2], ToString[n2]] +
     r KroneckerDelta[ToString[k2], ToString[n2]])
```

In[ ]:=
```
(*This is x' for gene content, denoted xp[k,i]:
 expected content of haplotype i among offspring haplotype k*)

xp[k_, i_] :=
 Sum[p[l] × Sum[R[StringPart[ToString[k], 1], StringPart[ToString[k], 2],
       StringPart[ToString[l], 1], StringPart[ToString[l], 2],
       StringPart[ToString[n], 1], StringPart[ToString[n], 2]] × x[n, i],
     {n, {ab, aB, Ab, AB}}], {l, {ab, aB, Ab, AB}}] /.
   {xb[ab] → 1 - xb[aB] - xb[Ab] - xb[AB]}

(*xpvec[k]: vector listing the average
  haplotype content among offspring of haplotype k*)
xpvec[k_] := {{xp[k, aB]}, {xp[k, Ab]}, {xp[k, AB]}};

MatrixForm[Table[Simplify[xpvec[k]], {k, {aB, Ab, AB}}]]
```

Out[ ]//MatrixForm=

$$\begin{pmatrix} \left(\frac{1}{2}\left(1 + xb[aB] - r\, xb[Ab]\right)\right) & \left(-\frac{1}{2}(-1+r)\, xb[Ab]\right) \\ \left(-\frac{1}{2}(-1+r)\, xb[aB]\right) & \left(\frac{1}{2}\left(1 - r\, xb[aB] + xb[Ab]\right)\right) \\ \left(\frac{1}{2}\left(-\left((-1+r)\, xb[aB]\right) - r\left(-1 + xb[Ab] + xb[AB]\right)\right)\right) & \left(\frac{1}{2}\left(r - r\, xb[aB] + xb[Ab] - r\, xb[Ab] \cdots\right.\right. \end{pmatrix}$$

In[ ]:=
```
(*This is the vector x'(xk) for a chosen parental haplotype k*)
MatrixForm[Simplify[xpvec[AB]]]
```

Out[ ]//MatrixForm=

$$\begin{pmatrix} \frac{1}{2}\left(-\left((-1+r)\, xb[aB]\right) - r\left(-1 + xb[Ab] + xb[AB]\right)\right) \\ \frac{1}{2}\left(r - r\, xb[aB] + xb[Ab] - r\, xb[Ab] - r\, xb[AB]\right) \\ \frac{1}{2}\left(1 - r + r\, xb[aB] + r\, xb[Ab] + xb[AB] + r\, xb[AB]\right) \end{pmatrix}$$

In[ ]:=
```
(*This is zetax = E[x']*)
zetax = Simplify[Sum[p[k] × xpvec[k], {k, {ab, aB, Ab, AB}}] /.
     {xb[ab] → 1 - xb[aB] - xb[Ab] - xb[AB]} /.
    {xb[aB] → xaBb, xb[Ab] → xAbb, xb[AB] → xABb}];
MatrixForm[zetax]
MatrixForm[zetax - xbFun[xaBb, xAbb, xABb]]
```

Out[ ]//MatrixForm=

$$\begin{pmatrix} xaBb - r\, xABb\,(-1 + xAbb + xABb) - r\, xaBb\,(xAbb + xABb) \\ xAbb - r\, xABb\,(-1 + xaBb + xABb) - r\, xAbb\,(xaBb + xABb) \\ xABb + r\, xABb\,(-1 + xAbb + xABb) + r\, xaBb\,(xAbb + xABb) \end{pmatrix}$$

Out[ ]//MatrixForm=

$$\begin{pmatrix} -r\, xABb\,(-1 + xAbb + xABb) - r\, xaBb\,(xAbb + xABb) \\ -r\, xABb\,(-1 + xaBb + xABb) - r\, xAbb\,(xaBb + xABb) \\ r\, xABb\,(-1 + xAbb + xABb) + r\, xaBb\,(xAbb + xABb) \end{pmatrix}$$

In[∘]:=
```
(*This is Px', that is, cov[x',x']*)
Pxp[xaBbvar_, xAbbvar_, xABbvar_] =
  FullSimplify[Sum[p[k] (xpvec[k] - zetax).Transpose[(xpvec[k] - zetax)],
      {k, {ab, aB, Ab, AB}}] /. {xb[ab] → 1 - xb[aB] - xb[Ab] - xb[AB]} /.
     {xb[aB] → xaBbvar, xb[Ab] → xAbbvar, xb[AB] → xABbvar} /.
    {xaBb → xaBbvar, xAbb → xAbbvar, xABb → xABbvar}];
MatrixForm[Pxp[xaBb, xAbb, xABb]]
```

Out[∘]//MatrixForm=

$$\begin{pmatrix} \frac{1}{4}\left(-r^2\, xABb\,(-1+xAbb+xABb)\left((1-2\,xABb)^2+xAbb\,(-1+4\,xABb)\right)+xaBb^2\,(- \right. \\ \frac{1}{4}\left(-r\, xaBb^2\,(xAbb+xABb)\,(-2+r\,(-1+4\,xAbb+4\,xABb))-r\,xABb\,(-1+xAbb+xABb \right. \\ \frac{1}{4}\left(-xaBb\,xABb+r\left(xaBb\,(xAbb+xABb)-2\,xaBb^2\,(xAbb+xABb)+xABb\,(-1+xAbb+xABb)\,(-1 \right.\right. \end{pmatrix}$$

In[∘]:=
```
(*This is Tx*)
Tx[xaBbvar_, xAbbvar_, xABbvar_] =
  Simplify[Sum[p[k] (xpvec[k] - zetax).Transpose[(xvec[k] - xb)],
      {k, {ab, aB, Ab, AB}}] /. {xb[ab] → 1 - xb[aB] - xb[Ab] - xb[AB]} /.
     {xb[aB] → xaBbvar, xb[Ab] → xAbbvar, xb[AB] → xABbvar} /.
    {xaBb → xaBbvar, xAbb → xAbbvar, xABb → xABbvar}];
MatrixForm[Tx[xaBb, xAbb, xABb]]

(*This is Hx*)
Hx = Simplify[Tx[xaBb, xAbb, xABb].Inverse[Px]];
MatrixForm[Hx]
```

Out[∘]//MatrixForm=

$$\begin{pmatrix} \frac{1}{2}\,xaBb\,(1+2\,r\,(-1+xABb)\,xABb+r\,xAbb\,(-1+2\,xABb)+xaBb\,(-1+2\,r\,(xAbb+xABb))) & r\,x[illegible] \\ r\,xaBb\,xABb\,(-1+xaBb+xABb)+\frac{1}{2}\,xaBb\,xAbb\,(-1+r\,(-1+2\,xaBb+2\,xABb)) & \frac{1}{2}\,xA[illegible] \\ -\frac{1}{2}\,xaBb\,(xABb+2\,r\,xABb\,(-1+xaBb+xABb)+r\,xAbb\,(-1+2\,xaBb+2\,xABb)) & -\frac{1}{2}\,[illegible] \end{pmatrix}$$

Out[∘]//MatrixForm=

$$\begin{pmatrix} \frac{1}{2}\,(1-r\,(xAbb+xABb)) & -\frac{1}{2}\,r\,(xaBb+xABb) & -\frac{1}{2}\,r\,(-1+xaBb+xAbb+2\,xABb) \\ -\frac{1}{2}\,r\,(xAbb+xABb) & \frac{1}{2}\,(1-r\,(xaBb+xABb)) & -\frac{1}{2}\,r\,(-1+xaBb+xAbb+2\,xABb) \\ \frac{1}{2}\,r\,(xAbb+xABb) & \frac{1}{2}\,r\,(xaBb+xABb) & \frac{1}{2}\,(1+r\,(-1+xaBb+xAbb+2\,xABb)) \end{pmatrix}$$

In[∘]:=
```
(*This is etax*)
eta[k_] :=
  xpvec[k] - zetax - Hx.(xvec[k] - xbvec) /. {xb[ab] → 1 - xb[aB] - xb[Ab] - xb[AB]} /.
   {xb[aB] → xaBb, xb[Ab] → xAbb, xb[AB] → xABb};

MatrixForm[Table[Simplify[eta[k]], {k, {ab, aB, Ab, AB}}]]
```

Out[∘]//MatrixForm=

$$\begin{pmatrix} (0) & (0) & (0) \\ (0) & (0) & (0) \\ (0) & (0) & (0) \\ (0) & (0) & (0) \end{pmatrix}$$

In[ ]:=
```
(*This is βx in general*)
betaxGen = Simplify[
   Inverse[Px].Sum[p[k] (xvec[k] - xbvec) (wG[k] - 1), {k, {ab, aB, Ab, AB}}] /.
     {xb[ab] → 1 - xb[aB] - xb[Ab] - xb[AB]} /.
    {xb[aB] → xaBb, xb[Ab] → xAbb, xb[AB] → xABb}];

MatrixForm[betaxGen]
```

Out[ ]//MatrixForm=
```
( -wG[ab] + wG[aB] )
( -wG[ab] + wG[Ab] )
( -wG[ab] + wG[AB] )
```

In[ ]:=
```
(*This is the selection pointer βxp in general*)
betaxpGen = Inverse[Hx].betaxGen;

MatrixForm[FullSimplify[betaxpGen]]
```

Out[ ]//MatrixForm=
```
( (2+4 r (-1+xaBb+xAbb+2 xABb)) wG[ab]-2 (wG[aB]+r (-1+xAbb+xABb) wG[aB]+r (xaBb+xABb) wG[Ab]+r (-1+xaBb+xAbb+2 xABb) w[illegible]
  ---------------------------------------------------------------------------------------------------------------
                                                    -1+r
  (2+4 r (-1+xaBb+xAbb+2 xABb)) wG[ab]-2 r (xAbb+xABb) wG[aB]-2 (1+r (-1+xaBb+xABb)) wG[Ab]-2 r (-1+xaBb+xAbb+2 xABb) w[illegible]
  ---------------------------------------------------------------------------------------------------------------
                                                    -1+r
  2 (wG[ab]-2 r (xaBb+xAbb+2 xABb) wG[ab]+r (xAbb+xABb) wG[aB]+r (xaBb+xABb) wG[Ab]+(-1+r (xaBb+xAbb+2 xABb)) wG[AB[illegible]
  ---------------------------------------------------------------------------------------------------------------
                                                    -1+r                                                          )
```

In[ ]:=
```
(*This is mean absolute fitness*)
WbGen = FullSimplify[
  Sum[p[k] × p[l] × WG[k, l], {k, {ab, aB, Ab, AB}}, {l, {ab, aB, Ab, AB}}] /.
    {xb[ab] → 1 - xb[aB] - xb[Ab] - xb[AB]} /.
   {xb[aB] → xaBb, xb[Ab] → xAbb, xb[AB] → xABb}]
```

Out[ ]=
```
(-1 + xaBb + xAbb + xABb)^2 WG[ab, ab] - xaBb (-1 + xaBb + xAbb + xABb) WG[ab, aB] -
 xAbb (-1 + xaBb + xAbb + xABb) WG[ab, Ab] - xABb (-1 + xaBb + xAbb + xABb) WG[ab, AB] -
 xaBb (-1 + xaBb + xAbb + xABb) WG[aB, ab] + xaBb^2 WG[aB, aB] + xaBb xAbb WG[aB, Ab] +
 xaBb xABb WG[aB, AB] - xAbb (-1 + xaBb + xAbb + xABb) WG[Ab, ab] + xaBb xAbb WG[Ab, aB] +
 xAbb^2 WG[Ab, Ab] + xAbb xABb WG[Ab, AB] - xABb (-1 + xaBb + xAbb + xABb) WG[AB, ab] +
 xaBb xABb WG[AB, aB] + xAbb xABb WG[AB, Ab] + xABb^2 WG[AB, AB]
```

```
In[*]:= (*This proves that betax==1/2 1/Wb dWb/dxb assuming that Wij is independent of
          allele frequency and of linkage disequilibrium and if Wij==Wji*)

        Simplify[
         {betaxGen[[1, 1]] == 1/2 1/WbGen D[WbGen, xaBb], betaxGen[[2, 1]] == 1/2 1/WbGen D[WbGen,
                xAbb], betaxGen[[3, 1]] == 1/2 1/WbGen D[WbGen, xABb]} /.
              {wG[ab] → WG[ab] / WbGen, wG[aB] → WG[aB] / WbGen,
               wG[Ab] → WG[Ab] / WbGen, wG[AB] → WG[AB] / WbGen} /.
             {WG[ab] → Sum[p[k] × WG[ab, k], {k, {ab, aB, Ab, AB}}],
              WG[aB] → Sum[p[k] × WG[aB, k], {k, {ab, aB, Ab, AB}}],
              WG[Ab] → Sum[p[k] × WG[Ab, k], {k, {ab, aB, Ab, AB}}],
              WG[AB] → Sum[p[k] × WG[AB, k], {k, {ab, aB, Ab, AB}}]} /.
            {WG[aB, ab] → WG[ab, aB], WG[Ab, ab] → WG[ab, Ab], WG[AB, ab] → WG[ab, AB],
             WG[Ab, aB] → WG[aB, Ab], WG[AB, aB] → WG[aB, AB], WG[AB, Ab] → WG[Ab, AB]} /.
           {xb[ab] → 1 - xb[aB] - xb[Ab] - xb[AB]} /.
          {xb[aB] → xaBb, xb[Ab] → xAbb, xb[AB] → xABb}]

Out[*]=
        {True, True, True}

In[*]:= (*Gene content for locus i in haplytope k*)
        xG[i_, k_] :=
         If[i == 1 && (ToString[k] == ToString[ab] || ToString[k] == ToString[aB]), 0,
          If[i == 1 && (ToString[k] == ToString[Ab] || ToString[k] == ToString[AB]), 1,
           If[i == 2 && (ToString[k] == ToString[ab] || ToString[k] == ToString[Ab]), 0,
            If[i == 2 && (ToString[k] == ToString[aB] || ToString[k] == ToString[AB]), 1]]]]

In[*]:= (*Phenotype i of genotype kl*)
        z[m_, k_, l_] :=
         cb (1 + e[m]) (y[1, k, l, m] + y[2, k, l, m]) - 2 cb e[m] × y[1, k, l, m] × y[2, k, l, m]

        y[i_, k_, l_, m_] :=
         c[i] (1 + d[i, m]) (xG[i, k] + xG[i, l]) - 2 c[i] × d[i, m] × xG[i, k] × xG[i, l]

        zvec[k_, l_] := {{z[1, k, l]}, {z[2, k, l]}};
```

In[ ]:=
```
(*These are the mean phenotypes*)
zbvec[xaBbvar_, xAbbvar_, xABbvar_] = Simplify[
   Sum[p[k] × p[l] × zvec[k, l], {k, {ab, aB, Ab, AB}}, {l, {ab, aB, Ab, AB}}] /.
     {xb[ab] → 1 - xb[aB] - xb[Ab] - xb[AB]} /. {xb[aB] → xaBb, xb[Ab] → xAbb,
     xb[AB] → xABb} /. {xaBb → xaBbvar, xAbb → xAbbvar, xABb → xABbvar}];

(*Mean phenotypes evaluated at the
 chosen dominance and epistasis coefficients*)
Simplify[
 MatrixForm[zbvec[xaBb, xAbb, xABb] /. {cb → 1, c[1] → 1, c[2] → 1, d[1, 1] → 0,
    d[1, 2] → 0, d[2, 1] → 0, d[2, 2] → 0, e[1] → 0, e[2] → -1}]]

Simplify[
 MatrixForm[zbvec[xaBb, xAbb, xABb] /. {cb → 1, c[1] → 1, c[2] → 1, d[1, 1] → 0,
     d[1, 2] → 0, d[2, 1] → 0, d[2, 2] → 0, e[1] → 0, e[2] → -1}] /.
   {xaBb → (1 - x1b) x2b - Xb, xAbb → x1b (1 - x2b) - Xb, xABb → x1b x2b + Xb}]
```

Out[ ]//MatrixForm=

$$\begin{pmatrix} 2\,(xaBb + xAbb + 2\,xABb) \\ 4\,(xaBb\,(xAbb + xABb) + xABb\,(1 + xAbb + xABb)) \end{pmatrix}$$

Out[ ]//MatrixForm=

$$\begin{pmatrix} 2\,(x1b + x2b) \\ 4\,(2\,x1b\,x2b + Xb) \end{pmatrix}$$

In[ ]:=
```
(*This is fitness of genotype kl*)
W[k_, l_, θ1_, σ1_, θ2_, σ2_] := Exp[-(z[1, k, l] - θ1)^2/σ1^2 - (z[2, k, l] - θ2)^2/σ2^2]

(*This is mean fitness*)
Wb[θ1_, σ1_, θ2_, σ2_] := Simplify[Sum[p[k] × p[l] × W[k, l, θ1, σ1, θ2, σ2],
   {k, {ab, aB, Ab, AB}}, {l, {ab, aB, Ab, AB}}]]

WbFun[xaBbvar_, xAbbvar_, xABbvar_] :=
  Wb[θ1, σ1, θ2, σ2] /. {xb[ab] → 1 - xb[aB] - xb[Ab] - xb[AB]} /.
    {xb[aB] → xaBb, xb[Ab] → xAbb, xb[AB] → xABb} /.
   {xaBb → xaBbvar, xAbb → xAbbvar, xABb → xABbvar};

(*This is relative fitness of genotype kl*)
w[k_, l_, θ1_, σ1_, θ2_, σ2_] := W[k, l, θ1, σ1, θ2, σ2] / Wb[θ1, σ1, θ2, σ2]

(*This is the relative fitness of haplotype k*)
wG[k_, θ1_, σ1_, θ2_, σ2_] :=
  Sum[p[l] × w[k, l, θ1, σ1, θ2, σ2], {l, {ab, aB, Ab, AB}}];

(*This runs the numerical solutions*)

r = 0.5;

cb = 1;
```

```
c[1] = 1;
c[2] = 1;
d[1, 1] = 0;
d[1, 2] = 0;
d[2, 1] = 0;
d[2, 2] = 0;
e[1] = 0;
e[2] = -1;

θ1 = 2;
θ2 = 4;
σ1 = Sqrt[10];
σ2 = Sqrt[40];

tend = 100;

(*Initial conditions must be such that initial
 haplotype frequencies are between zero and one*)
xaBinit = {0.01, 1, .2};
xAbinit = {0.01, 1, .2};
xABinit = {0.01, 1, .2};

(*This is Δx bar to be used for numerical solutions*)

Deltax[xaBbvar_, xAbbvar_, xABbvar_] =
  Tx[xaBb, xAbb, xABb].betaxGen + zetax - xbFun[xaBb, xAbb, xABb] /.
      {wG[ab] → wG[ab, θ1, σ1, θ2, σ2], wG[aB] → wG[aB, θ1, σ1, θ2, σ2],
       wG[Ab] → wG[Ab, θ1, σ1, θ2, σ2], wG[AB] → wG[AB, θ1, σ1, θ2, σ2]} /.
     {xb[ab] → 1 - xb[aB] - xb[Ab] - xb[AB]} /. {xb[aB] → xaBb, xb[Ab] → xAbb,
     xb[AB] → xABb} /. {xaBb → xaBbvar, xAbb → xAbbvar, xABb → xABbvar};

pSol[k_, xaBbvar_, xAbbvar_, xABbvar_] :=
  p[k] /. {xb[ab] → 1 - xb[aB] - xb[Ab] - xb[AB]} /. {xb[aB] → xaBb, xb[Ab] → xAbb,
     xb[AB] → xABb} /. {xaBb → xaBbvar, xAbb → xAbbvar, xABb → xABbvar};

Table[If[0 ≤ pSol[ab, i, j, k] ≤ 1, {xaBbSol[0, i, j, k] = i,
    xAbbSol[0, i, j, k] = j, xABbSol[0, i, j, k] = k, xDbSol[0, i, j, k] =
     {{xaBbSol[0, i, j, k]}, {xAbbSol[0, i, j, k]}, {xABbSol[0, i, j, k]}},
    Table[{xDbSol[t + 1, i, j, k] = xDbSol[t, i, j, k] +
        Deltax[xaBbSol[t, i, j, k], xAbbSol[t, i, j, k], xABbSol[t, i, j, k]],
      xaBbSol[t + 1, i, j, k] = xDbSol[t + 1, i, j, k][[1, 1]],
      xAbbSol[t + 1, i, j, k] = xDbSol[t + 1, i, j, k][[2, 1]],
      xABbSol[t + 1, i, j, k] = xDbSol[t + 1, i, j, k][[3, 1]]}, {t, 0, tend}]}],
  {i, xaBinit[[1]], xaBinit[[2]], xaBinit[[3]]}, {j, xAbinit[[1]],
   xAbinit[[2]], xAbinit[[3]]},
  {k, xABinit[[1]], xABinit[[2]], xABinit[[3]]}];
```

In[ ]:=
```
(*This counts the number of initial conditions that meet the
 criterion that haplotype frequencies are between zero and one*)

Counter = 0;
Table[If[0 ≤ pSol[ab, i, j, k] ≤ 1, {Counter = Counter + 1;
    ActualInitialConditions[Counter] = {i, j, k}}],
  {i, xaBinit⟦1⟧, xaBinit⟦2⟧, xaBinit⟦3⟧}, {j, xAbinit⟦1⟧,
   xAbinit⟦2⟧, xAbinit⟦3⟧}, {k, xABinit⟦1⟧, xABinit⟦2⟧, xABinit⟦3⟧}];
CounterEnd = Counter
```

Out[ ]=
```
35
```

In[ ]:=
```
(*This makes a series of plots*)

(*This computes the final points to be plotted*)

pointSize = 0.05;

endPointsxP = Table[
   {xAbbSol[tend, ActualInitialConditions[Counter]⟦1⟧, ActualInitialConditions[
        Counter]⟦2⟧, ActualInitialConditions[Counter]⟦3⟧] + xABbSol[tend,
      ActualInitialConditions[Counter]⟦1⟧, ActualInitialConditions[Counter]⟦2⟧,
      ActualInitialConditions[Counter]⟦3⟧], xaBbSol[tend,
      ActualInitialConditions[Counter]⟦1⟧, ActualInitialConditions[Counter]⟦2⟧,
      ActualInitialConditions[Counter]⟦3⟧] + xABbSol[tend,
      ActualInitialConditions[Counter]⟦1⟧, ActualInitialConditions[Counter]⟦2⟧,
      ActualInitialConditions[Counter]⟦3⟧]}, {Counter, 1, CounterEnd}];

yAxisLimitz := 1.1
yAxisLimitP := .6

lThickness = 0.02; (*Line thickness*)

(*This plots the change of allele frequencies
 for both loci on top of the mean fitness contour*)
plot =
 Show[Table[ListLinePlot[Table[{xAbbSol[t, ActualInitialConditions[Counter]⟦1⟧,
        ActualInitialConditions[Counter]⟦2⟧,
        ActualInitialConditions[Counter]⟦3⟧] + xABbSol[t,
        ActualInitialConditions[Counter]⟦1⟧, ActualInitialConditions[Counter]⟦
         2⟧, ActualInitialConditions[Counter]⟦3⟧],
      xaBbSol[t, ActualInitialConditions[Counter]⟦1⟧, ActualInitialConditions[
          Counter]⟦2⟧, ActualInitialConditions[Counter]⟦3⟧] +
       xABbSol[t, ActualInitialConditions[Counter]⟦1⟧,
        ActualInitialConditions[Counter]⟦2⟧,
        ActualInitialConditions[Counter]⟦3⟧]}, {t, 0, tend}],
    PlotStyle → Thickness[lThickness], Frame → True, FrameStyle → Large,
```

```
     TicksStyle → Large,
     PlotRange → {{-0.1, 1.1}, {-.1, 1.1}},
     AspectRatio → 1,
     Epilog → {PointSize[pointSize], Point[endPointsxP]}],
    {Counter, 1, CounterEnd}]]
(*Rasterize the plot to eliminate vectorization artifacts*)
plot = Rasterize[plot, "Image", ImageResolution → 300];
Export[StringJoin[ToString[NotebookDirectory[]], "Fig.S3a", ".pdf"], plot];

(*This shows the change in allele frequencies
 in both loci and in linkage disquilibrium over time*)
Show[Table[ListLinePlot[{Table[
      xAbbSol[t, ActualInitialConditions[Counter]⟦1⟧, ActualInitialConditions[
          Counter]⟦2⟧, ActualInitialConditions[Counter]⟦3⟧] + xABbSol[t,
        ActualInitialConditions[Counter]⟦1⟧, ActualInitialConditions[Counter]⟦
         2⟧, ActualInitialConditions[Counter]⟦3⟧], {t, 0, tend}],
     Table[xaBbSol[t, ActualInitialConditions[Counter]⟦1⟧,
        ActualInitialConditions[Counter]⟦2⟧,
        ActualInitialConditions[Counter]⟦3⟧] + xABbSol[t,
        ActualInitialConditions[Counter]⟦1⟧, ActualInitialConditions[Counter]⟦
         2⟧, ActualInitialConditions[Counter]⟦3⟧], {t, 0, tend}],
     Table[(1 - xaBbSol[t, ActualInitialConditions[Counter]⟦1⟧,
           ActualInitialConditions[Counter]⟦2⟧,
           ActualInitialConditions[Counter]⟦3⟧] - xAbbSol[t,
           ActualInitialConditions[Counter]⟦1⟧, ActualInitialConditions[
             Counter]⟦2⟧, ActualInitialConditions[Counter]⟦3⟧] - xABbSol[t,
           ActualInitialConditions[Counter]⟦1⟧, ActualInitialConditions[
             Counter]⟦2⟧, ActualInitialConditions[Counter]⟦3⟧])
        xABbSol[t, ActualInitialConditions[Counter]⟦1⟧,
         ActualInitialConditions[Counter]⟦2⟧,
         ActualInitialConditions[Counter]⟦3⟧] -
       xaBbSol[t, ActualInitialConditions[Counter]⟦1⟧,
         ActualInitialConditions[Counter]⟦2⟧,
         ActualInitialConditions[Counter]⟦3⟧] ×
        xAbbSol[t, ActualInitialConditions[Counter]⟦1⟧,
         ActualInitialConditions[Counter]⟦2⟧,
         ActualInitialConditions[Counter]⟦3⟧], {t, 0, tend}]},
    PlotRange → {-0.2, yAxisLimitz}, TicksStyle → Large,
    PlotStyle →
     Thickness[lThickness]],
   {Counter, 1, CounterEnd}]]
Export[StringJoin[ToString[NotebookDirectory[]], "Fig.S3c", ".pdf"], %];

(*This plots the change in mean absolute fitness over time*)
Show[Table[
   ListLinePlot[{Table[WbFun[xaBbSol[t, ActualInitialConditions[Counter]⟦1⟧,
        ActualInitialConditions[Counter]⟦2⟧,
```

```
        ActualInitialConditions[Counter][[3]], xAbbSol[t,
        ActualInitialConditions[Counter][[1]], ActualInitialConditions[Counter][[
         2]], ActualInitialConditions[Counter][[3]], xABbSol[t,
        ActualInitialConditions[Counter][[1]], ActualInitialConditions[Counter][[
         2]], ActualInitialConditions[Counter][[3]]], {t, 0, tend}]},
    PlotRange → {0, yAxisLimitz}, TicksStyle → Large,
    PlotStyle → Thickness[lThickness]],
   {Counter, 1, CounterEnd}]]
(*Export[StringJoin[ToString[NotebookDirectory[]],"Fig.S3e",".pdf"],%];*)
```

Out[ ]=

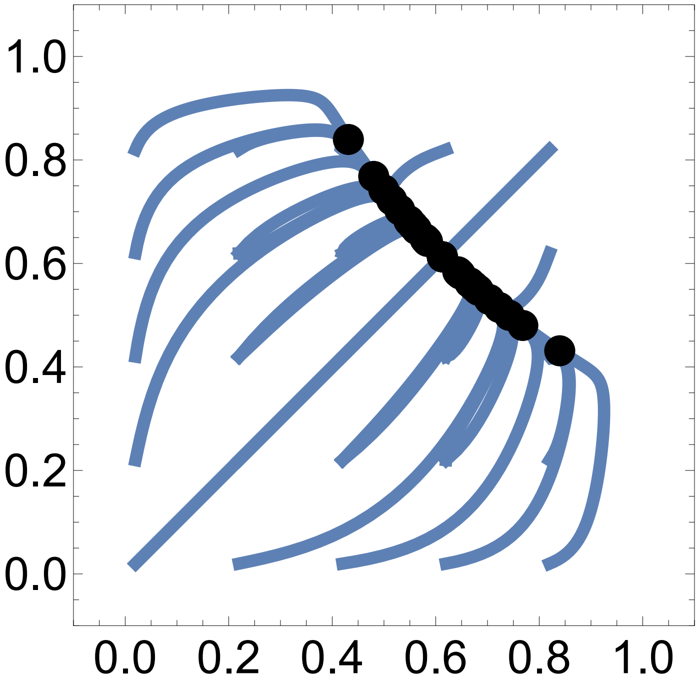

Out[ ]=

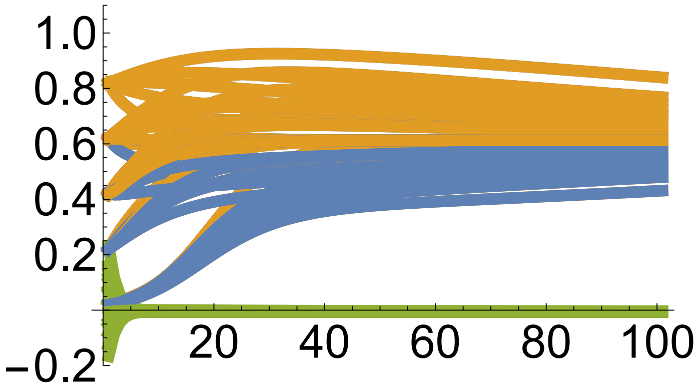

Out[ ]=

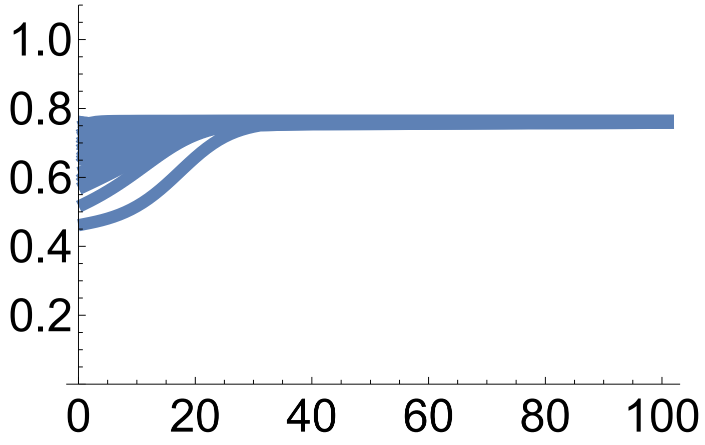

In[ ]:=
```
(*This plots mean z1 (blue) and mean z2 (orange) over time*)
Show[Table[
  ListLinePlot[{Table[zbvec[xaBbSol[t, ActualInitialConditions[Counter]⟦1⟧,
        ActualInitialConditions[Counter]⟦2⟧,
        ActualInitialConditions[Counter]⟦3⟧], xAbbSol[t,
        ActualInitialConditions[Counter]⟦1⟧, ActualInitialConditions[Counter]⟦
         2⟧, ActualInitialConditions[Counter]⟦3⟧], xABbSol[t,
        ActualInitialConditions[Counter]⟦1⟧, ActualInitialConditions[Counter]⟦
         2⟧, ActualInitialConditions[Counter]⟦3⟧]]⟦1, 1⟧, {t, 0, tend}],
    Table[zbvec[xaBbSol[t, ActualInitialConditions[Counter]⟦1⟧,
        ActualInitialConditions[Counter]⟦2⟧,
        ActualInitialConditions[Counter]⟦3⟧], xAbbSol[t,
        ActualInitialConditions[Counter]⟦1⟧, ActualInitialConditions[Counter]⟦
         2⟧, ActualInitialConditions[Counter]⟦3⟧], xABbSol[t,
        ActualInitialConditions[Counter]⟦1⟧, ActualInitialConditions[Counter]⟦
         2⟧, ActualInitialConditions[Counter]⟦3⟧]]⟦2, 1⟧, {t, 0, tend}],
    Table[θ1, {t, 0, tend}], Table[θ2, {t, 0, tend}]}, PlotRange →
    {0, 6}, TicksStyle → Large,
   PlotStyle → {Thickness[lThickness], Thickness[lThickness],
     {Magenta, Thickness[lThickness]},
     {Magenta, Thickness[lThickness]}}], {Counter, 1, CounterEnd}]]
Export[StringJoin[ToString[NotebookDirectory[]], "Fig.S3d", ".pdf"], %];
```

Out[ ]=

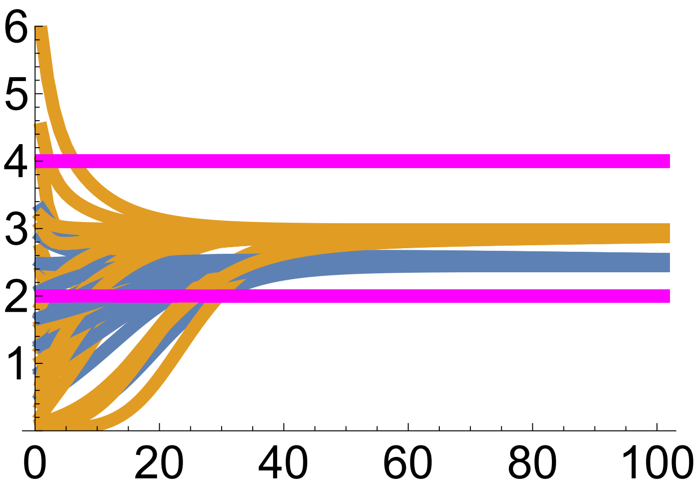

In[ ]:=
```
(*This plots the phase diagram for mean phenotypes*)

(*This computes the final points for the plots of the variance evolution*)

endPointszP = Table[{zbvec[xaBbSol[tend, ActualInitialConditions[Counter]⟦1⟧,
       ActualInitialConditions[Counter]⟦2⟧,
       ActualInitialConditions[Counter]⟦3⟧], xAbbSol[tend,
       ActualInitialConditions[Counter]⟦1⟧, ActualInitialConditions[Counter]⟦
        2⟧, ActualInitialConditions[Counter]⟦3⟧],
      xABbSol[tend, ActualInitialConditions[Counter]⟦1⟧,
```

```
      ActualInitialConditions[Counter]⟦2⟧,
      ActualInitialConditions[Counter]⟦3⟧]]⟦1, 1⟧,
   zbvec[xaBbSol[tend, ActualInitialConditions[Counter]⟦1⟧,
      ActualInitialConditions[Counter]⟦2⟧,
      ActualInitialConditions[Counter]⟦3⟧],
     xAbbSol[tend, ActualInitialConditions[Counter]⟦1⟧,
      ActualInitialConditions[Counter]⟦2⟧,
      ActualInitialConditions[Counter]⟦3⟧],
     xABbSol[tend, ActualInitialConditions[Counter]⟦1⟧,
      ActualInitialConditions[Counter]⟦2⟧,
      ActualInitialConditions[Counter]⟦3⟧]]⟦2, 1⟧}, {Counter, 1, CounterEnd}];

(*This plots mean z1 vs mean z2 with
 admissible developmental manifold in red*)

plot = Show[Table[
   ListLinePlot[{Table[{zbvec[xaBbSol[t, ActualInitialConditions[Counter]⟦1⟧,
          ActualInitialConditions[Counter]⟦2⟧,
          ActualInitialConditions[Counter]⟦3⟧], xAbbSol[t,
          ActualInitialConditions[Counter]⟦1⟧, ActualInitialConditions[
            Counter]⟦2⟧, ActualInitialConditions[Counter]⟦3⟧], xABbSol[t,
          ActualInitialConditions[Counter]⟦1⟧, ActualInitialConditions[
            Counter]⟦2⟧, ActualInitialConditions[Counter]⟦3⟧]]⟦1, 1⟧,
       zbvec[xaBbSol[t, ActualInitialConditions[Counter]⟦1⟧,
          ActualInitialConditions[Counter]⟦2⟧,
          ActualInitialConditions[Counter]⟦3⟧],
         xAbbSol[t, ActualInitialConditions[Counter]⟦1⟧,
          ActualInitialConditions[Counter]⟦2⟧,
          ActualInitialConditions[Counter]⟦3⟧],
         xABbSol[t, ActualInitialConditions[Counter]⟦1⟧,
          ActualInitialConditions[Counter]⟦2⟧,
          ActualInitialConditions[Counter]⟦3⟧]]⟦2, 1⟧}, {t, 0, tend}]},
    PlotRange → {{-0.1, 4.1}, {-0.1, 6.1}}, TicksStyle →
     Large, AspectRatio →
     1, PlotStyle →
     Thickness[0.5 lThickness],
    Epilog → {{PointSize[pointSize], Point[endPointszP]},
      {PointSize[pointSize], Magenta, Point[{θ1, θ2}]}}],
   {Counter, 1, CounterEnd}]]
(*Rasterize the plot to eliminate vectorization artifacts*)
plot = Rasterize[plot, "Image", ImageResolution → 300];
Export[StringJoin[ToString[NotebookDirectory[]], "Fig.S3b", ".pdf"], plot];
```

Out[ ]=

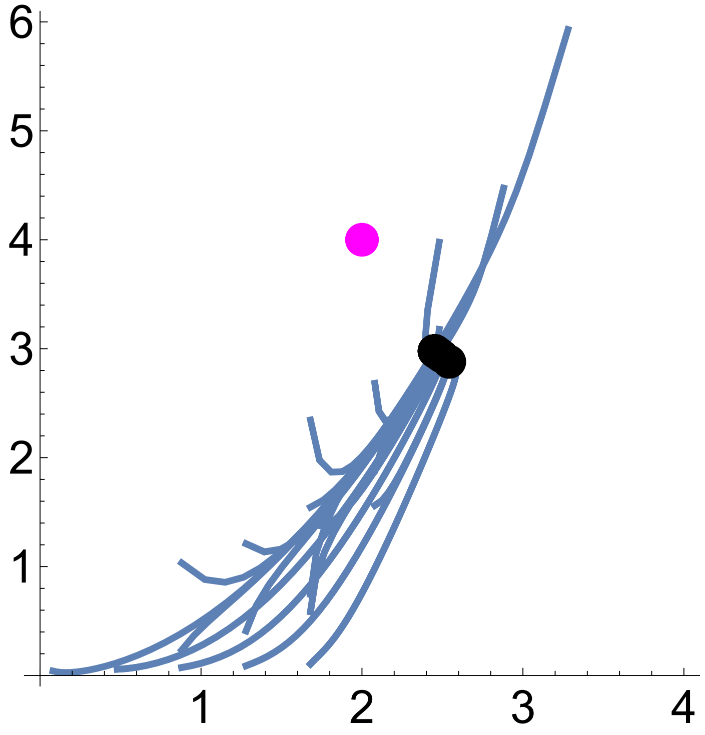

#### Example 5 exact evo-devo dynamics: one biallelic locus, explicit development of one phenotype, fitness depends on phenotype at final age only

In[ ]:=
```
Clear["Global`*"]
```

In[ ]:=
```
(*Gene content and offspring gene content*)
xa = 0; xA = 1;
xap = pa xa + pA (1/2 xA + 1/2 xa) ; xAp = pa (1/2 xA + 1/2 xa) + pA xA;

(*Haplotype frequencies*)
pa = (1 - xb);
pA = xb;
```

In[ ]:=
```
(*This is Tx*)
Tx[xvar_] = FullSimplify[
   Sum[p[i] (xp[i] - xb) (x[i] - xb), {i, 1, 2}] /. {p[1] → pa, p[2] → pA} /.
     {x[1] → xa, x[2] → xA} /. {xp[1] → xap, xp[2] → xAp}] /. {xb → xvar}

(*This is Px = var[x]*)
Px[xvar_] =
 FullSimplify[Sum[p[i] (x[i] - xb)^2, {i, 1, 2}] /. {p[1] → pa, p[2] → pA} /.
     {x[1] → xa, x[2] → xA} /. {xp[1] → xap, xp[2] → xAp}] /. {xb → xvar}
```

Out[ ]=

$$-\frac{1}{2}(-1+\text{xvar})\,\text{xvar}$$

Out[ ]=

$$-((-1+\text{xvar})\,\text{xvar})$$

In[ ]:=
```
(*This is zetax*)
zetax[xvar_] :=
 FullSimplify[Sum[p[i] × xp[i], {i, 1, 2}] /. {p[1] → pa, p[2] → pA} /.
    {xp[1] → xap, xp[2] → xAp}] /. {xb → xvar}
```

In[ ]:=
```
zetax[xb]
```

Out[ ]=

xb

In[ ]:=
```
(*Phenotype of genotype ij*)
z[i_, j_, a_] := (1 + y[i, j])^(a-1) z1

y[i_, j_] := c (1 + d) (x[i] + x[j]) - 2 c d x[i] × x[j]

(*Mean phenotype*)
zbfun[xvar_] := FullSimplify[
   Sum[p[i] × p[j] × z[i, j, Na], {i, 1, 2}, {j, 1, 2}] /. {p[1] → pa, p[2] → pA} /.
    {x[1] → xa, x[2] → xA}] /. {xb → xvar}

(*Phenotype variance*)
P[xvar_] :=
 FullSimplify[Sum[p[i] × p[j] (z[i, j, Na] - zbfun[xb])^2, {i, 1, 2}, {j, 1, 2}] /.
     {p[1] → pa, p[2] → pA} /. {x[1] → xa, x[2] → xA}] /. {xb → xvar}
```

In[ ]:=
```
zbfun[xb]

P[xb]
```

Out[ ]=

$$\left((-1+xb)^2-2\,(1+c+c\,d)^{-1+Na}\,(-1+xb)\,xb+(1+2\,c)^{-1+Na}\,xb^2\right)z1$$

Out[ ]=

$$xb^2\left(-(1+2\,c)^{-1+Na}+(-1+xb)^2-2\,(1+c+c\,d)^{-1+Na}\,(-1+xb)\,xb+(1+2\,c)^{-1+Na}\,xb^2\right)^2 z1^2+ 2\,(1-xb)\,xb \left(-(1+c+c\,d)^{-1+Na}+(-1+xb)^2-2\,(1+c+c\,d)^{-1+Na}\,(-1+xb)\,xb+(1+2\,c)^{-1+Na}\,xb^2\right)^2 z1^2+ (-1+xb)^2\left(z1-\left((-1+xb)^2-2\,(1+c+c\,d)^{-1+Na}\,(-1+xb)\,xb+(1+2\,c)^{-1+Na}\,xb^2\right)z1\right)^2$$

In[ ]:=
```
zbfun[xb] /. {d → 0, c → 1}
```

Out[ ]=

$$\left((-1+xb)^2-2^{Na}\,(-1+xb)\,xb+3^{-1+Na}\,xb^2\right)z1$$

```
In[•]:= (*This is fitness for genotype ij*)
        W[i_, j_, Na_, θ_, σ_] := Exp[-(z[i, j, Na] - θ)^2/σ^2] /. {x[1] → xa, x[2] → xA}

        (*Mean fitness*)
        Wb[xvar_, Na_, θ_, σ_] :=
         FullSimplify[Sum[p[i] × p[j] × W[i, j, Na, θ, σ], {i, 1, 2}, {j, 1, 2}] /.
            {p[1] → pa, p[2] → pA}] /. {xb → xvar}

        (*Relative fitness for genotype ij*)
        w[i_, j_, Na_, xb_, θ_, σ_] := W[i, j, Na, θ, σ] / Wb[xb, Na, θ, σ]

        (*This is absolute and relative fitness for haplotype i*)
        Wh[i_, Na_, θ_, σ_] :=
         Sum[p[j] × W[i, j, Na, θ, σ], {j, 1, 2}] /. {p[1] → pa, p[2] → pA}
        wh[i_, xb_, Na_, θ_, σ_] :=
         Sum[p[j] × w[i, j, Na, xb, θ, σ], {j, 1, 2}] /. {p[1] → pa, p[2] → pA}

In[•]:= (*This is cov[w,x]*)
        covwx[xvar_] := Simplify[Sum[p[i] (x[i] - xb) (wh[i, xb, Na, θ, σ] - 1), {i, 1, 2}] /.
             {x[1] → xa, x[2] → xA} /. {p[1] → pa, p[2] → pA}] /. {xb → xvar}

        (*This is betax*)
        betax[xbvar_, cvar_, dvar_, θvar_, σvar_, Navar_, z1var_] := covwx[xb]/Px[xb] /.
           {xb → xbvar, c → cvar, d → dvar, θ → θvar, σ → σvar, Na → Navar, z1 → z1var};
```

In[ ]:=
```
(*This runs the numerical solutions*)

Na = 3;
z1 = 0.1;

c = 1;
d = 2;

θ = 1;
σ = Sqrt[10];

tend = 500;

x1binit = {0.01, 1, .1};

InitialConditions =
  Table[{x1bSol[0, i] = i}, {i, x1binit[[1]], x1binit[[2]], x1binit[[3]]}];

Table[Table[{x1bSol[t + 1, i] =
     x1bSol[t, i] + (Tx[x1bSol[t, i]] × betax[x1bSol[t, i], c, d, θ, σ, Na, z1])},
   {t, 0, tend}], {i, x1binit[[1]], x1binit[[2]], x1binit[[3]]}];
```

```
In[ ]:= (*This plots allele frequencies, mean phenotype,
phenotype variance, and mean fitness over time*)

Plot1FileName =
  If[θ == 1.5 && d == 0, "a", If[θ == 1 && d == 0, "f", If[θ == 1 && d == 2, "k"]]];
Plot2FileName =
  If[θ == 1.5 && d == 0, "b", If[θ == 1 && d == 0, "g", If[θ == 1 && d == 2, "l"]]];
Plot3FileName =
  If[θ == 1.5 && d == 0, "c", If[θ == 1 && d == 0, "h", If[θ == 1 && d == 2, "m"]]];

yAxisLimitz[d_] := If[d ≥ 2, 1.3, 2]
yAxisLimitP[d_] := If[d ≥ 2, 0.6, .1]

lThickness = 0.02; (*Line thickness*)

Show[
 Table[ListLinePlot[{Table[x1bSol[t, i], {t, 0, tend}]}, PlotRange → {-0.1, 1.1},
   TicksStyle → Large, PlotStyle → Thickness[lThickness]],
  {i, x1binit[[1]], x1binit[[2]], x1binit[[3]]}]]
Export[StringJoin[ToString[NotebookDirectory[]],
   "Fig.S1", Plot1FileName, ".pdf"], %];

Show[Table[
  ListLinePlot[{Table[zbfun[x1bSol[t, i]], {t, 0, tend}], Table[θ, {t, 0, tend}]},
   PlotRange → {-0.1, yAxisLimitz[d]}, TicksStyle → Large,
   PlotStyle → {Thickness[lThickness], {Magenta, Thickness[lThickness]}}],
  {i, x1binit[[1]], x1binit[[2]], x1binit[[3]]}]]
Export[StringJoin[ToString[NotebookDirectory[]],
   "Fig.S1", Plot2FileName, ".pdf"], %];

Show[Table[ListLinePlot[{Table[P[x1bSol[t, i]], {t, 0, tend}]},
   PlotRange → {-0.1 yAxisLimitP[d], yAxisLimitP[d]}, TicksStyle → Large,
   PlotStyle → Thickness[lThickness]], {i, x1binit[[1]], x1binit[[2]], x1binit[[3]]}]]
Export[StringJoin[ToString[NotebookDirectory[]],
   "Fig.S1", Plot3FileName, ".pdf"], %];

Show[Table[ListLinePlot[{Table[Wb[x1bSol[t, i], Na, θ, σ], {t, 0, tend}]},
   PlotRange → {-0.1, 1.1}, TicksStyle → Large,
   PlotStyle → Thickness[lThickness]], {i, x1binit[[1]], x1binit[[2]], x1binit[[3]]}]]
```

Out[ ]=

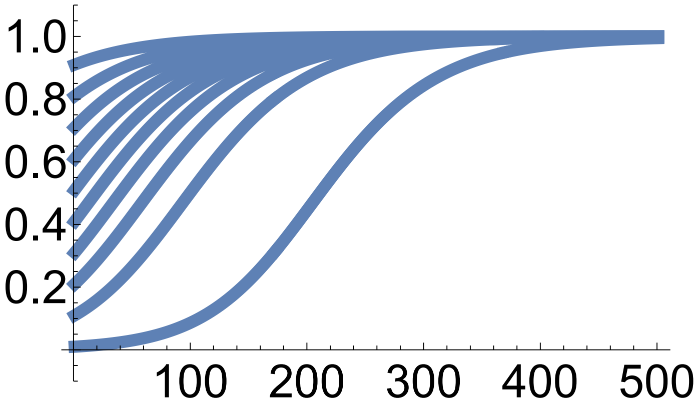

Out[ ]=

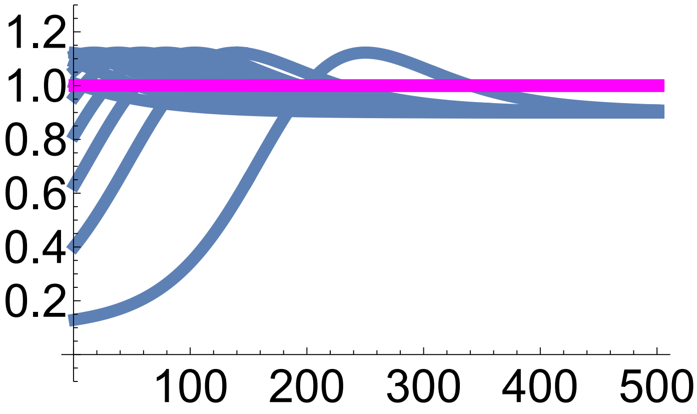

Out[ ]=

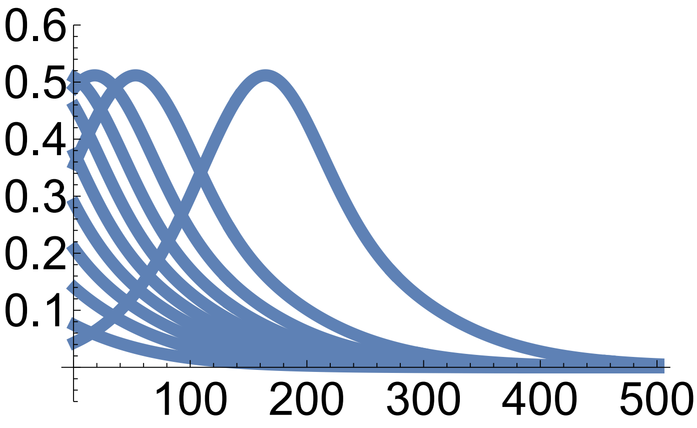

Out[ ]=

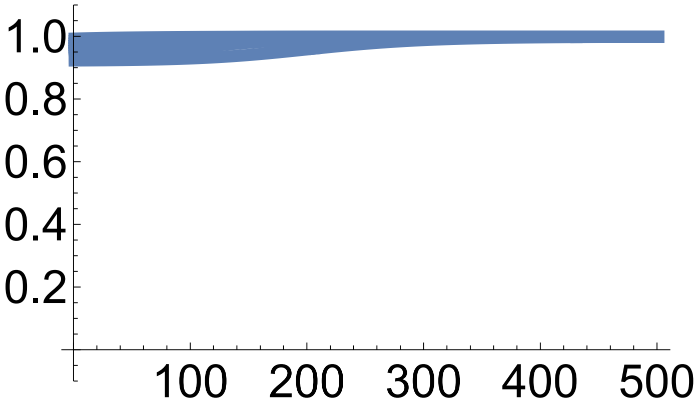

```
In[ ]:= (*This plots mean phenotype and mean fitness vs allele frequency*)

pointSize = 0.05;

Plot4FileName =
  If[θ == 1.5 && d == 0, "d", If[θ == 1 && d == 0, "i", If[θ == 1 && d == 2, "n"]]];
Plot5FileName =
  If[θ == 1.5 && d == 0, "e", If[θ == 1 && d == 0, "j", If[θ == 1 && d == 2, "o"]]];

endPoints = Table[{x1bSol[tend, i], zbfun[x1bSol[tend, i]]}, {i, x1binit⟦1⟧,
    x1binit⟦2⟧, x1binit⟦3⟧}, {j, x1binit⟦1⟧, x1binit⟦2⟧, x1binit⟦3⟧}];

p0 = Table[ListLinePlot[Table[{x1bSol[t, i], zbfun[x1bSol[t, i]]}, {t, 0, tend}],
    PlotRange → {{0, 1}, {0, 2}}, PlotStyle → {Thickness[lThickness]}],
   {i, x1binit⟦1⟧, x1binit⟦2⟧, x1binit⟦3⟧}];

plot =  Show[ContourPlot[Wb[xb, Na, θ, σ], {xb, 0, 1}, {zb, 0, yAxisLimitz[d]},
   FrameStyle → Large, PlotLegends → Automatic, LabelStyle → Large,
   Epilog → {PointSize[pointSize], Point[Flatten[endPoints, 1]]}],
  Plot[zbfun[x1b], {x1b, 0, 1}, PlotStyle → {Red, Thickness[3 lThickness]}],
  p0 /. Line[x_] :→ {Arrowheads[Table[{.1, .2}, {3}]], Arrow[x]}, ListLinePlot[
   Table[{xb, θ}, {xb, 0, 1}], PlotStyle → {Magenta, Thickness[lThickness]}]]

plot = Rasterize[plot, "Image", ImageResolution → 300];

Export[StringJoin[ToString[NotebookDirectory[]],
   "Fig.S1", Plot4FileName, ".pdf"], plot];

endPointsxP = Table[{x1bSol[tend, i], P[x1bSol[tend, i]]}, {i, x1binit⟦1⟧,
    x1binit⟦2⟧, x1binit⟦3⟧}, {j, x1binit⟦1⟧, x1binit⟦2⟧, x1binit⟦3⟧}];

p0 = Table[ListLinePlot[Table[{x1bSol[t, i], P[x1bSol[t, i]]}, {t, 0, tend}],
    PlotRange → {{0, 1}, {0, 1}}, PlotStyle → {Thickness[lThickness]}],
   {i, x1binit⟦1⟧, x1binit⟦2⟧, x1binit⟦3⟧}];

plot = Show[ContourPlot[Wb[xb, Na, θ, σ], {xb, 0, 1}, {P, 0, yAxisLimitP[d]},
   FrameStyle → Large, PlotLegends → Automatic, LabelStyle → Large,
   Epilog → {PointSize[pointSize], Point[Flatten[endPointsxP, 1]]}],
  Plot[P[x1b], {x1b, 0, 1}, PlotStyle → {Red, Thickness[3 lThickness]}],
  p0 /. Line[x_] :→ {Arrowheads[Table[{.1, .2}, {3}]], Arrow[x]}]

plot = Rasterize[plot, "Image", ImageResolution → 300];

Export[StringJoin[ToString[NotebookDirectory[]],
   "Fig.S1", Plot5FileName, ".pdf"], plot];
```

Out[ ]=

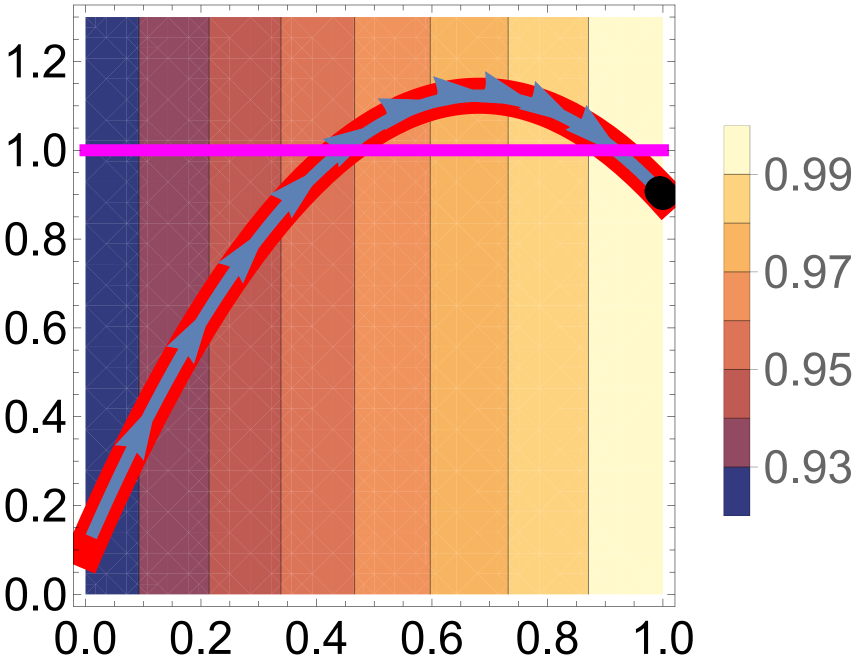
1.2
1.0
0.8
0.6
0.4
0.2
0.0
0.0
0.2
0.4
0.6
0.8
1.0
0.99
0.97
0.95
0.93

Out[ ]=

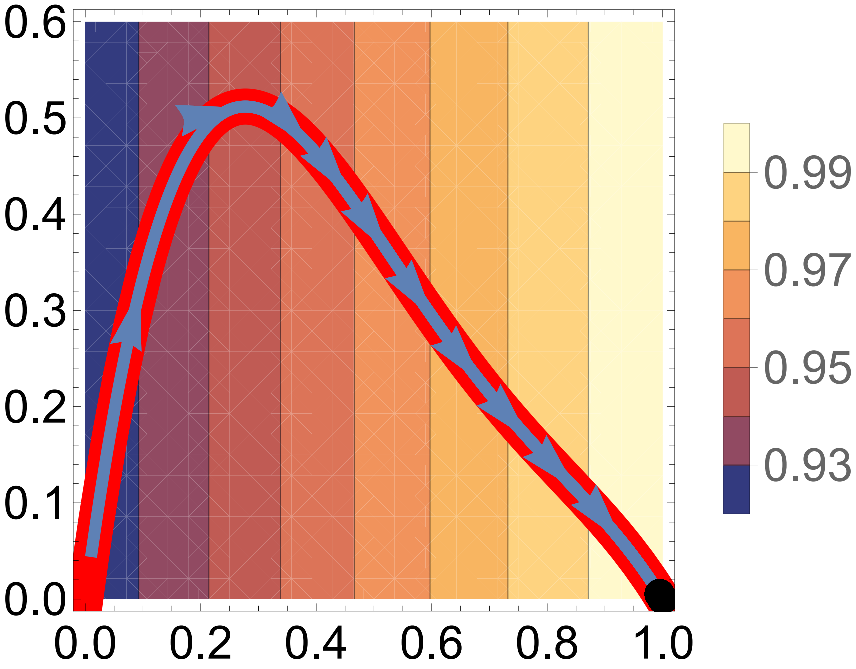
0.6
0.5
0.4
0.3
0.2
0.1
0.0
0.0
0.2
0.4
0.6
0.8
1.0
0.99
0.97
0.95
0.93

#### Example 5 approximated evo-devo dynamics: one biallelic locus, explicit development of one phenotype, fitness depends on phenotype at final age only

In[∘]:=
```
Clear["Global`*"]
```

In[∘]:=
```
(*This is Tx*)
Tx[xvar_] = -1/2 (-1 + xvar) xvar;
```

In[∘]:=
```
(*Developmental map*)
g[z_, xk_, xl_] := (1 + y[xk, xl]) z

y[xk_, xl_] := c (1 + d) (xk + xl) - 2 c d xk xl

z[xk_, xl_, a_] := (1 + y[xk, xl])^(a-1) z1;
```

In[∘]:=
```
(*This is D*)

De[xb_] = c (1 + d) - 2 c d xb;

(*This checks that dz/dx = D z*)
D[g[z, xk, xl], xk] == z De[xb] /. {xk → xb, xl → xb}
```

Out[∘]=
```
True
```

In[∘]:=
```
(*These are the indicated derivatives*)

dwdx[xb_] = dzdx[xb, Na] × dwdz[z[xb, xb, Na]];

dwdz[zb_] := 2/σ^2 (θ - zb)

dzdx[xb_, a_] :=
 If[a == 1, 0, If[a == 2, De[xb] z1, If[a > 2, (a - 1) De[xb] z1 (1 + y[xb, xb])^(a-2)]]]

(*This is the MAG variance*)

Lz[xb_, a_] := 2 dzdx[xb, a]^2 Tx[xb]

(*This is fitness*)
W[z_] := Exp[-(z - θ)^2/σ^2]
```

In[ ]:=
```
(*This runs the numerical solutions*)

Na = 3;
z1 = 0.1;

c = 1;
d = 2;

θ = 1;
σ = Sqrt[10];

tend = 500;

x1binit = {0.01, 1, .1};

InitialConditions =
  Table[{x1bSol[0, i] = i}, {i, x1binit⟦1⟧, x1binit⟦2⟧, x1binit⟦3⟧}];

Table[
  Table[{x1bSol[t + 1, i] = x1bSol[t, i] + (Tx[x1bSol[t, i]] × dwdx[x1bSol[t, i]])},
   {t, 0, tend}], {i, x1binit⟦1⟧, x1binit⟦2⟧, x1binit⟦3⟧}];
```

```
In[ ]:= (*This plots allele frequency, mean phenotype,
MAG variance, and mean fitness over time*)

Plot1FileName =
  If[θ == 1.5 && d == 0, "p", If[θ == 1 && d == 0, "u", If[θ == 1 && d == 2, "z"]]];
Plot2FileName =
  If[θ == 1.5 && d == 0, "q", If[θ == 1 && d == 0, "v", If[θ == 1 && d == 2, "aa"]]];
Plot3FileName =
  If[θ == 1.5 && d == 0, "r", If[θ == 1 && d == 0, "w", If[θ == 1 && d == 2, "ab"]]];

yAxisLimitz[d_] := If[d ≥ 2, 1.3, 2]
yAxisLimitP[d_] := If[d ≥ 2, 0.2, .05]

lThickness = 0.02; (*Line thickness*)

Show[
 Table[ListLinePlot[{Table[x1bSol[t, i], {t, 0, tend}]}, PlotRange → {-0.1, 1.1},
   TicksStyle → Large, PlotStyle → Thickness[lThickness]],
  {i, x1binit[[1]], x1binit[[2]], x1binit[[3]]}]]
Export[StringJoin[ToString[NotebookDirectory[]],
   "Fig.S1", Plot1FileName, ".pdf"], %];

Show[Table[ListLinePlot[
   {Table[z[x1bSol[t, i], x1bSol[t, i], Na], {t, 0, tend}], Table[θ, {t, 0, tend}]},
   PlotRange → {-0.1, yAxisLimitz[d]}, TicksStyle → Large,
   PlotStyle → {Thickness[lThickness], {Magenta, Thickness[lThickness]}}],
  {i, x1binit[[1]], x1binit[[2]], x1binit[[3]]}]]
Export[StringJoin[ToString[NotebookDirectory[]],
   "Fig.S1", Plot2FileName, ".pdf"], %];

Show[Table[ListLinePlot[{Table[Lz[x1bSol[t, i], Na], {t, 0, tend}]},
   PlotRange → {-0.1 yAxisLimitP[d], yAxisLimitP[d]}, TicksStyle → Large,
   PlotStyle → Thickness[lThickness]], {i, x1binit[[1]], x1binit[[2]], x1binit[[3]]}]]
Export[StringJoin[ToString[NotebookDirectory[]],
   "Fig.S1", Plot3FileName, ".pdf"], %];

Show[
 Table[ListLinePlot[{Table[W[z[x1bSol[t, i], x1bSol[t, i], Na]], {t, 0, tend}]},
   PlotRange → {-0.1, 1.1}, TicksStyle → Large,
   PlotStyle → Thickness[lThickness]], {i, x1binit[[1]], x1binit[[2]], x1binit[[3]]}]]
```

Out[ ]=

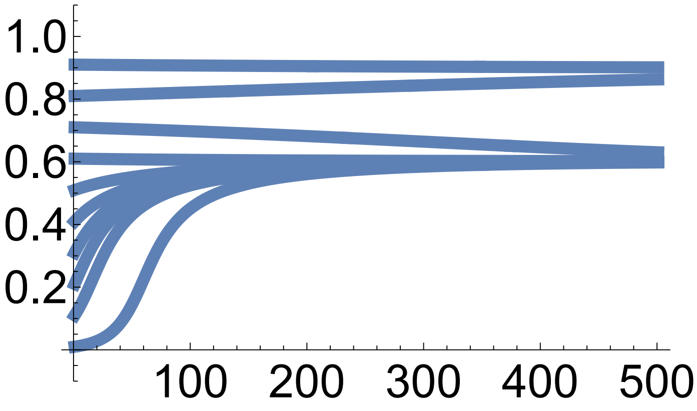

Out[ ]=

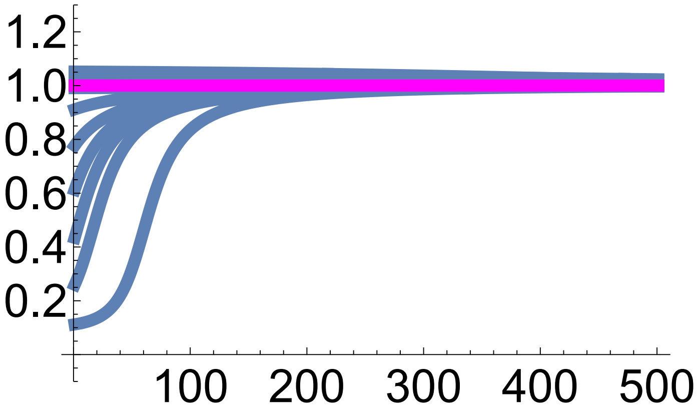

Out[ ]=

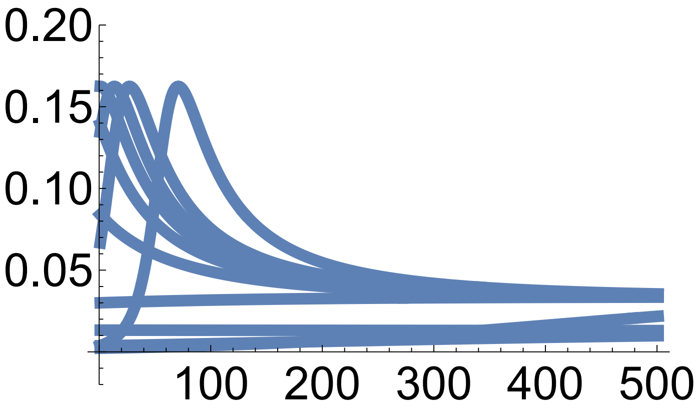

Out[ ]=

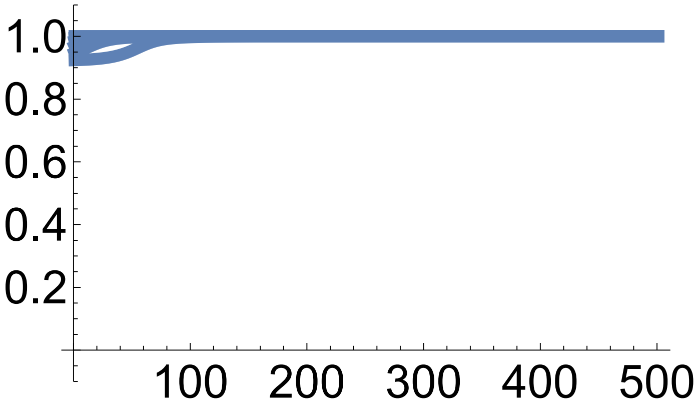

In[•]:=
```
(*This plots mean phenotype and MAG covariance vs allele frequency*)

pointSize = 0.05;

Plot4FileName =
  If[θ == 1.5 && d == 0, "s", If[θ == 1 && d == 0, "x", If[θ == 1 && d == 2, "ac"]]];
Plot5FileName =
  If[θ == 1.5 && d == 0, "t", If[θ == 1 && d == 0, "y", If[θ == 1 && d == 2, "ad"]]];

endPoints = Table[{x1bSol[tend, i], z[x1bSol[tend, i], x1bSol[tend, i], Na]},
   {i, x1binit⟦1⟧, x1binit⟦2⟧, x1binit⟦3⟧},
   {j, x1binit⟦1⟧, x1binit⟦2⟧, x1binit⟦3⟧}];

p0 = Table[ListLinePlot[
    Table[{x1bSol[t, i], z[x1bSol[t, i], x1bSol[t, i], Na]}, {t, 0, tend}],
    PlotRange → {{0, 1}, {0, 2}}, PlotStyle → {Thickness[lThickness]}],
   {i, x1binit⟦1⟧, x1binit⟦2⟧, x1binit⟦3⟧}];

plot =  Show[ContourPlot[W[zb], {xb, 0, 1}, {zb, 0, yAxisLimitz[d]},
   FrameStyle → Large, PlotLegends → Automatic, LabelStyle → Large,
   Epilog → {PointSize[pointSize], Point[Flatten[endPoints, 1]]}],
  Plot[z[x1b, x1b, Na], {x1b, 0, 1}, PlotStyle → {Red, Thickness[3 lThickness]}],
  p0 /. Line[x_] :→ {Arrowheads[Table[{.1, .2}, {3}]], Arrow[x]}, ListLinePlot[
   Table[{xb, θ}, {xb, 0, 1}], PlotStyle → {Magenta, Thickness[lThickness]}]]

plot = Rasterize[plot, "Image", ImageResolution → 300];

Export[StringJoin[ToString[NotebookDirectory[]],
   "Fig.S1", Plot4FileName, ".pdf"], plot];

endPointsxP = Table[{x1bSol[tend, i], Lz[x1bSol[tend, i], Na]}, {i, x1binit⟦1⟧,
    x1binit⟦2⟧, x1binit⟦3⟧}, {j, x1binit⟦1⟧, x1binit⟦2⟧, x1binit⟦3⟧}];

p0 = Table[ListLinePlot[Table[{x1bSol[t, i], Lz[x1bSol[t, i], Na]}, {t, 0, tend}],
    PlotRange → {{0, 1}, {0, 1}}, PlotStyle → {Thickness[lThickness]}],
   {i, x1binit⟦1⟧, x1binit⟦2⟧, x1binit⟦3⟧}];

plot  = Show[ContourPlot[W[z[xb, xb, Na]], {xb, 0, 1}, {P, 0, yAxisLimitP[d]},
   FrameStyle → Large, PlotLegends → Automatic, LabelStyle → Large,
   Epilog → {PointSize[pointSize], Point[Flatten[endPointsxP, 1]]}],
  Plot[Lz[x1b, Na], {x1b, 0, 1}, PlotStyle → {Red, Thickness[3 lThickness]}],
  p0 /. Line[x_] :→ {Arrowheads[Table[{.1, .2}, {3}]], Arrow[x]}]

plot = Rasterize[plot, "Image", ImageResolution → 300];

Export[StringJoin[ToString[NotebookDirectory[]],
   "Fig.S1", Plot5FileName, ".pdf"], plot];
```

Out[•]=

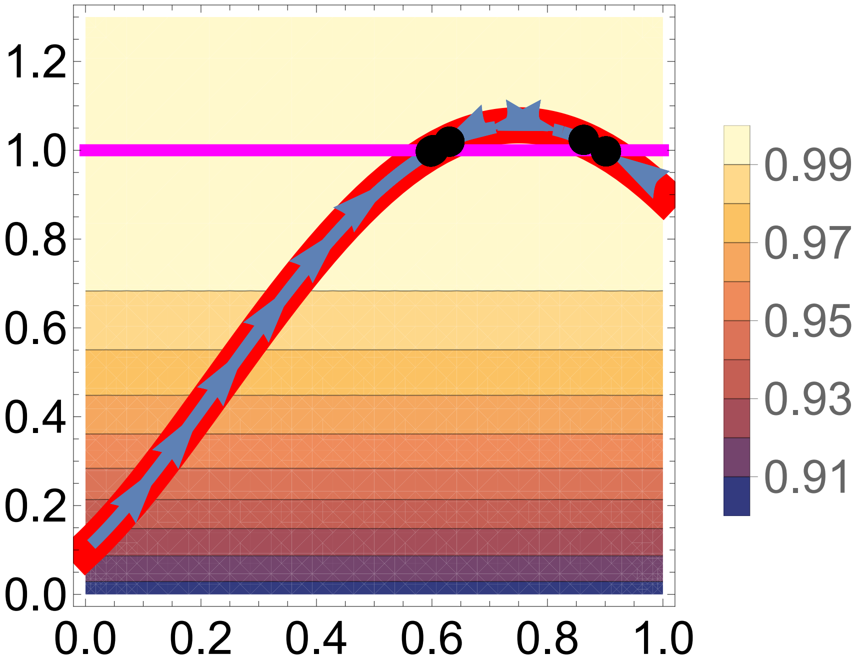
1.2
1.0
0.8
0.6
0.4
0.2
0.0
0.0 0.2 0.4 0.6 0.8 1.0
0.99
0.97
0.95
0.93
0.91

Out[•]=

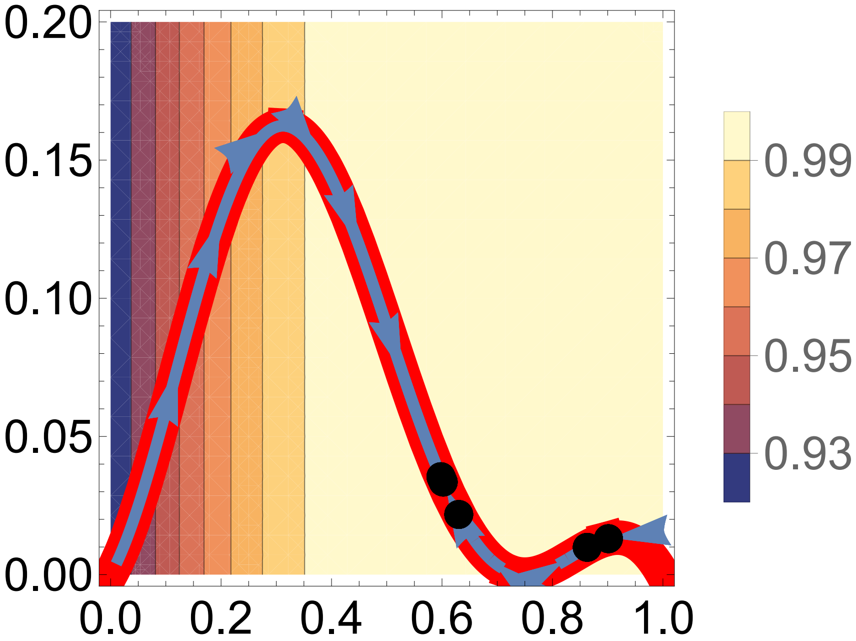
0.20
0.15
0.10
0.05
0.00
0.0 0.2 0.4 0.6 0.8 1.0
0.99
0.97
0.95
0.93

#### Example 6 exact evo-devo dynamics: one biallelic locus, explicit development of one phenotype, fitness depends on the phenotype at every age, weakening of selection with age

In[•]:=
```
Clear["Global`*"]
```

In[•]:=
```
(*Gene content and offspring gene content*)
xa = 0; xA = 1;
xap = pa xa + pA (1/2 xA + 1/2 xa) ; xAp = pa (1/2 xA + 1/2 xa) + pA xA;

(*Haplotype frequencies*)
pa = (1 - xb);
pA = xb;
```

In[•]:=
```
(*This is Tx*)
Tx[xvar_] = FullSimplify[
   Sum[p[i] (xp[i] - xb) (x[i] - xb), {i, 1, 2}] /. {p[1] → pa, p[2] → pA} /.
     {x[1] → xa, x[2] → xA} /. {xp[1] → xap, xp[2] → xAp}] /. {xb → xvar}

(*This is Px = var[x]*)
Px[xvar_] =
 FullSimplify[Sum[p[i] (x[i] - xb)^2, {i, 1, 2}] /. {p[1] → pa, p[2] → pA} /.
     {x[1] → xa, x[2] → xA} /. {xp[1] → xap, xp[2] → xAp}] /. {xb → xvar}
```

Out[•]=

$$-\frac{1}{2}(-1+\text{xvar})\,\text{xvar}$$

Out[•]=

$$-((-1+\text{xvar})\,\text{xvar})$$

In[•]:=
```
(*This is zetax*)
zetax[xvar_] :=
 FullSimplify[Sum[p[i] × xp[i], {i, 1, 2}] /. {p[1] → pa, p[2] → pA} /.
    {xp[1] → xap, xp[2] → xAp}] /. {xb → xvar}
```

```
In[ ]:= (*Phenotype of genotype ij*)
z[i_, j_, a_] := (1 + y[i, j])^(a-1) z1

y[i_, j_] := c (1 + d) (x[i] + x[j]) - 2 c d x[i] × x[j]

(*Mean phenotype at age a*)
zbfun[xvar_, a_] :=
 FullSimplify[Sum[p[i] × p[j] × z[i, j, a], {i, 1, 2}, {j, 1, 2}] /.
     {p[1] → pa, p[2] → pA} /. {x[1] → xa, x[2] → xA}] /. {xb → xvar}

(*Variance of phenotype at age a*)
P[xvar_, a_] :=
 FullSimplify[Sum[p[i] × p[j] (z[i, j, a] - zbfun[xb, a])^2, {i, 1, 2}, {j, 1, 2}] /.
     {p[1] → pa, p[2] → pA} /. {x[1] → xa, x[2] → xA}] /. {xb → xvar}

In[ ]:= zbfun[xb, a]

P[xb, a]
```

Out[ ]=

$$\left((-1+xb)^2-2\,(1+c+c\,d)^{-1+a}\,(-1+xb)\,xb+(1+2\,c)^{-1+a}\,xb^2\right)z1$$

Out[ ]=

$$xb^2\left(-(1+2\,c)^{-1+a}+(-1+xb)^2-2\,(1+c+c\,d)^{-1+a}\,(-1+xb)\,xb+(1+2\,c)^{-1+a}\,xb^2\right)^2 z1^2+$$
$$2\,(1-xb)\,xb$$
$$\left(-(1+c+c\,d)^{-1+a}+(-1+xb)^2-2\,(1+c+c\,d)^{-1+a}\,(-1+xb)\,xb+(1+2\,c)^{-1+a}\,xb^2\right)^2 z1^2+$$
$$(-1+xb)^2\left(z1-\left((-1+xb)^2-2\,(1+c+c\,d)^{-1+a}\,(-1+xb)\,xb+(1+2\,c)^{-1+a}\,xb^2\right)z1\right)^2$$

```
In[•]:= (*This is fitness for genotype ij*)
W[xvar_, i_, j_, Na_, θ_, σ_] :=
   1/τ[xb, Na] Sum[ϕ[xb, a] × f[i, j, a] + πf[xb, a, Na] × s[i, j, a], {a, 1, Na}] /.
     {x[1] → xa, x[2] → xA} /. {xb → xvar}
s[i_, j_, a_] := Exp[-(z[i, j, a] - θ)^2/σ^2] /. {x[1] → xa, x[2] → xA}
f[i_, j_, a_] := Exp[-(z[i, j, a] - θ)^2/σ^2] /. {x[1] → xa, x[2] → xA}
l[i_, j_, k_] := Product[s[i, j, a], {a, 1, k - 1}] /. {x[1] → xa, x[2] → xA}
πf[xvar_, a_, Na_] :=
  Sum[p[i] × p[j] 1/s[i, j, a] Sum[l[i, j, k] × f[i, j, k], {k, a + 1, Na}], {i, 1, 2},
      {j, 1, 2}] /. {x[1] → xa, x[2] → xA} /. {p[1] → pa, p[2] → pA} /. {xb → xvar}
ϕ[xvar_, a_] := Sum[p[i] × p[j] × l[i, j, a], {i, 1, 2}, {j, 1, 2}] /.
    {p[1] → pa, p[2] → pA} /. {xb → xvar}
τ[xvar_, Na_] :=
  Sum[p[i] × p[j] × Sum[a l[i, j, a] × f[i, j, a], {a, 1, Na}], {i, 1, 2}, {j, 1, 2}] /.
     {x[1] → xa, x[2] → xA} /. {p[1] → pa, p[2] → pA} /. {xb → xvar}

(*This is mean fitness*)

Wb[xvar_, Na_, θ_, σ_] :=
  Simplify[Sum[p[i] × p[j] × W[xb, i, j, Na, θ, σ], {i, 1, 2}, {j, 1, 2}] /.
     {p[1] → pa, p[2] → pA} /. {xb → xvar}]

(*This is relative fitness for genotype ij*)

w[xb_, i_, j_, Na_, θ_, σ_] := W[xb, i, j, Na, θ, σ] / Wb[xb, Na, θ, σ]

(*This is absolute and relative fitness for haplotype i*)
Wh[xvar_, i_, Na_, θ_, σ_] :=
  Sum[p[j] × W[xb, i, j, Na, θ, σ], {j, 1, 2}] /. {p[1] → pa, p[2] → pA} /. {xb → xvar}
wh[xvar_, i_, Na_, θ_, σ_] :=
  Sum[p[j] × w[xb, i, j, Na, θ, σ], {j, 1, 2}] /. {p[1] → pa, p[2] → pA} /. {xb → xvar}
```

```
In[•]:= Na = 3;

        (*This is cov[w,x]*)
        covwx[xvar_] := Simplify[Sum[p[i] (x[i] - xb) (wh[xb, i, Na, θ, σ] - 1), {i, 1, 2}] /.
              {x[1] → xa, x[2] → xA} /. {p[1] → pa, p[2] → pA}] /. {xb → xvar};

        (*This is betax*)
        betax[xbvar_, cvar_, dvar_, θvar_, σvar_, Navar_, z1var_] :=
          Simplify[covwx[xb] / Px[xb]] /.
           {xb → xbvar, c → cvar, d → dvar, θ → θvar, σ → σvar, Na → Navar, z1 → z1var};

        (*This runs the numerical solutions. It takes
         about 6 minutes. For faster runtime, reduce tend*)

        z1 = 0.1;

        c = 1;
        d = 2;

        (*Numerical solutions*)

        θ = 1;
        σ = Sqrt[10];

        tend = 1000;

        x1binit = {0.01, 1, .1};

        InitialConditions =
          Table[{x1bSol[0, i] = i}, {i, x1binit[[1]], x1binit[[2]], x1binit[[3]]}];

        Table[Table[{x1bSol[t + 1, i] =
              x1bSol[t, i] + (Tx[x1bSol[t, i]] × betax[x1bSol[t, i], c, d, θ, σ, Na, z1])},
           {t, 0, tend}], {i, x1binit[[1]], x1binit[[2]], x1binit[[3]]}];
```

```
In[ ]:= (*This plots allele frequency, mean phenotype,
phenotype variance, and mean fitness over time*)

Plot1FileName =
  If[θ == 1.5 && d == 0, "a", If[θ == 1 && d == 0, "f", If[θ == 1 && d == 2, "k"]]];
Plot2FileName =
  If[θ == 1.5 && d == 0, "b", If[θ == 1 && d == 0, "g", If[θ == 1 && d == 2, "l"]]];
Plot3FileName =
  If[θ == 1.5 && d == 0, "c", If[θ == 1 && d == 0, "h", If[θ == 1 && d == 2, "m"]]];

yAxisLimitz[d_] := If[d ≥ 2, 1.3, 2]
yAxisLimitP[d_] := If[d ≥ 2, 0.6, .1]

lThickness = 0.02; (*Line thickness*)

Show[
 Table[ListLinePlot[{Table[x1bSol[t, i], {t, 0, tend}]}, PlotRange → {-0.1, 1.1},
   TicksStyle → Large, PlotStyle → Thickness[lThickness]],
  {i, x1binit[[1]], x1binit[[2]], x1binit[[3]]}]]
Export[StringJoin[ToString[NotebookDirectory[]],
   "Fig.S2", Plot1FileName, ".pdf"], %];

Show[Table[ListLinePlot[
   {Table[zbfun[x1bSol[t, i], Na], {t, 0, tend}], Table[θ, {t, 0, tend}]},
   PlotRange → {-0.1, yAxisLimitz[d]}, TicksStyle → Large,
   PlotStyle → {Thickness[lThickness], {Magenta, Thickness[lThickness]}}],
  {i, x1binit[[1]], x1binit[[2]], x1binit[[3]]}]]
Export[StringJoin[ToString[NotebookDirectory[]],
   "Fig.S2", Plot2FileName, ".pdf"], %];

Show[Table[ListLinePlot[{Table[P[x1bSol[t, i], Na], {t, 0, tend}]},
   PlotRange → {-0.1 yAxisLimitP[d], yAxisLimitP[d]}, TicksStyle → Large,
   PlotStyle → Thickness[lThickness]], {i, x1binit[[1]], x1binit[[2]], x1binit[[3]]}]]
Export[StringJoin[ToString[NotebookDirectory[]],
   "Fig.S2", Plot3FileName, ".pdf"], %];

Show[Table[ListLinePlot[{Table[Wb[x1bSol[t, i], Na, θ, σ], {t, 0, tend}]},
   TicksStyle → Large, PlotRange → Full, PlotStyle → Thickness[lThickness]],
  {i, x1binit[[1]], x1binit[[2]], x1binit[[3]]}]]
```

Out[ ]=

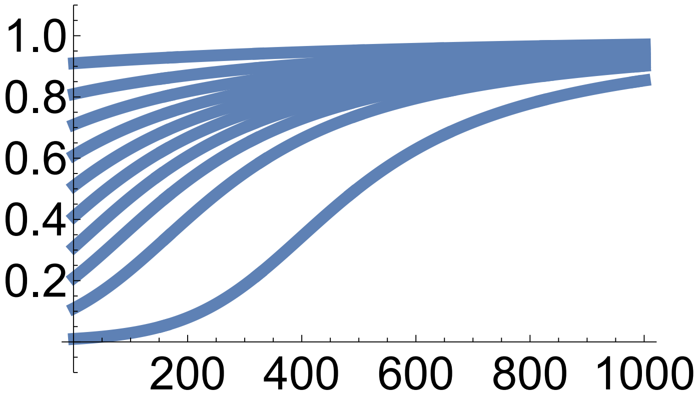

Out[ ]=

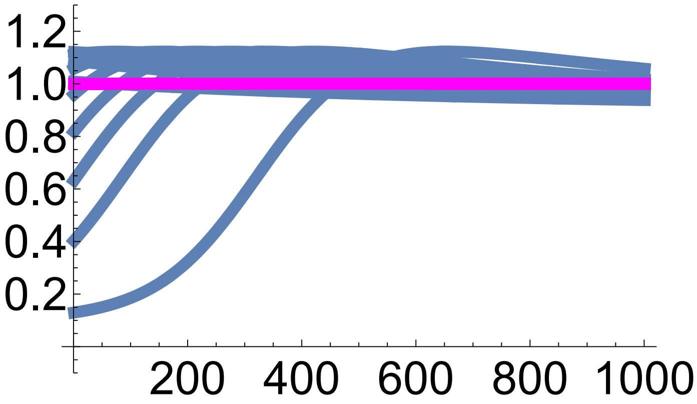

Out[ ]=

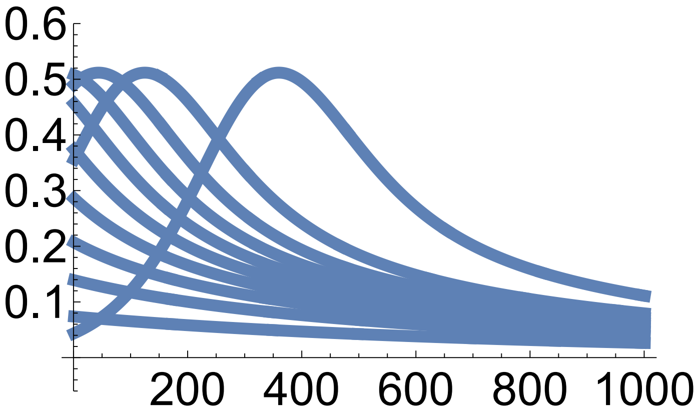

Out[ ]=

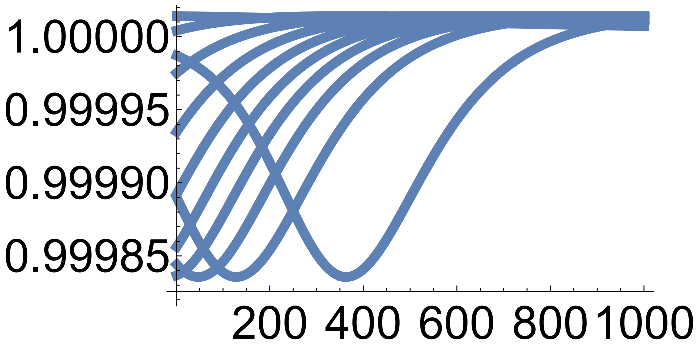

In[ ]:=
```
(*This plots mean phenotype and phenotype variance vs allele frequency*)

pointSize = 0.05;

Plot4FileName =
  If[θ == 1.5 && d == 0, "d", If[θ == 1 && d == 0, "i", If[θ == 1 && d == 2, "n"]]];
Plot5FileName =
  If[θ == 1.5 && d == 0, "e", If[θ == 1 && d == 0, "j", If[θ == 1 && d == 2, "o"]]];

endPoints = Table[{x1bSol[tend, i], zbfun[x1bSol[tend, i], Na]}, {i, x1binit[[1]],
    x1binit[[2]], x1binit[[3]]}, {j, x1binit[[1]], x1binit[[2]], x1binit[[3]]}];

p0 =
  Table[ListLinePlot[Table[{x1bSol[t, i], zbfun[x1bSol[t, i], Na]}, {t, 0, tend}],
    PlotRange → {{0, 1}, {0, 2}}, PlotStyle → {Thickness[lThickness]}],
   {i, x1binit[[1]], x1binit[[2]], x1binit[[3]]}];

plot =  Show[ContourPlot[Wb[xb, Na, θ, σ], {xb, 0, 1}, {zb, 0, yAxisLimitz[d]},
   FrameStyle → Large, PlotLegends → Automatic, LabelStyle → Large,
   Epilog → {PointSize[pointSize], Point[Flatten[endPoints, 1]]}],
  Plot[zbfun[x1b, Na], {x1b, 0, 1}, PlotStyle → {Red, Thickness[3 lThickness]}],
  p0 /. Line[x_] :→ {Arrowheads[Table[{.1, .2}, {3}]], Arrow[x]}, ListLinePlot[
   Table[{xb, θ}, {xb, 0, 1}], PlotStyle → {Magenta, Thickness[lThickness]}]]

plot = Rasterize[plot, "Image", ImageResolution → 300];

Export[StringJoin[ToString[NotebookDirectory[]],
   "Fig.S2", Plot4FileName, ".pdf"], plot];

endPointsxP = Table[{x1bSol[tend, i], P[x1bSol[tend, i], Na]}, {i, x1binit[[1]],
    x1binit[[2]], x1binit[[3]]}, {j, x1binit[[1]], x1binit[[2]], x1binit[[3]]}];

p0 = Table[ListLinePlot[Table[{x1bSol[t, i], P[x1bSol[t, i], Na]}, {t, 0, tend}],
    PlotRange → {{0, 1}, {0, 1}}, PlotStyle → {Thickness[lThickness]}],
   {i, x1binit[[1]], x1binit[[2]], x1binit[[3]]}];

plot  = Show[ContourPlot[Wb[xb, Na, θ, σ], {xb, 0, 1}, {P, 0, yAxisLimitP[d]},
   FrameStyle → Large, PlotLegends → Automatic, LabelStyle → Large,
   Epilog → {PointSize[pointSize], Point[Flatten[endPointsxP, 1]]}],
  Plot[P[x1b, Na], {x1b, 0, 1}, PlotStyle → {Red, Thickness[3 lThickness]}],
  p0 /. Line[x_] :→ {Arrowheads[Table[{.1, .2}, {3}]], Arrow[x]}]

plot = Rasterize[plot, "Image", ImageResolution → 300];

Export[StringJoin[ToString[NotebookDirectory[]],
   "Fig.S2", Plot5FileName, ".pdf"], plot];
```

Out[ ]=

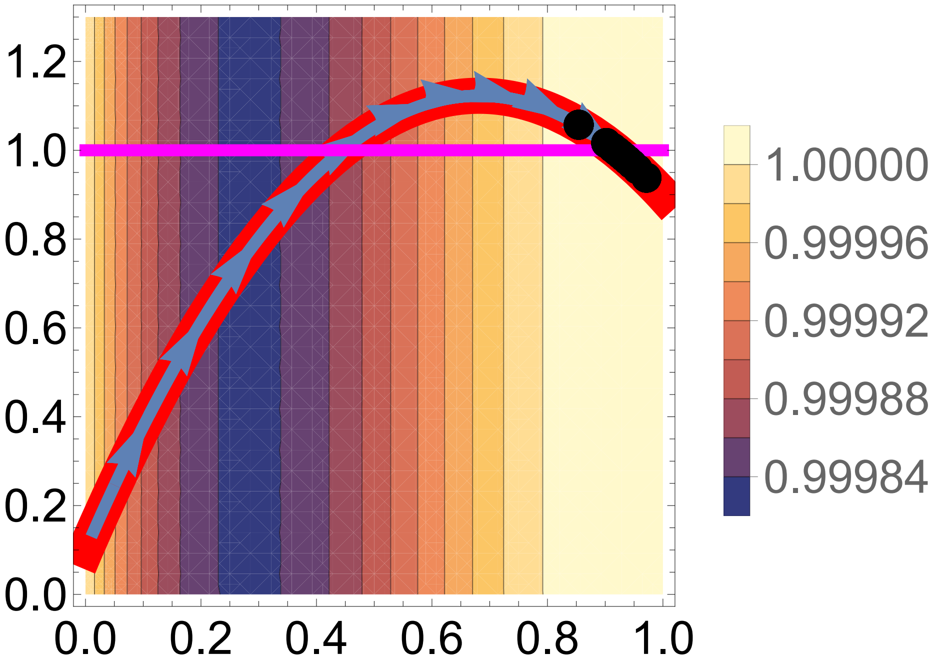
1.2
1.0
0.8
0.6
0.4
0.2
0.0
0.0
0.2
0.4
0.6
0.8
1.0
1.00000
0.99996
0.99992
0.99988
0.99984

Out[ ]=

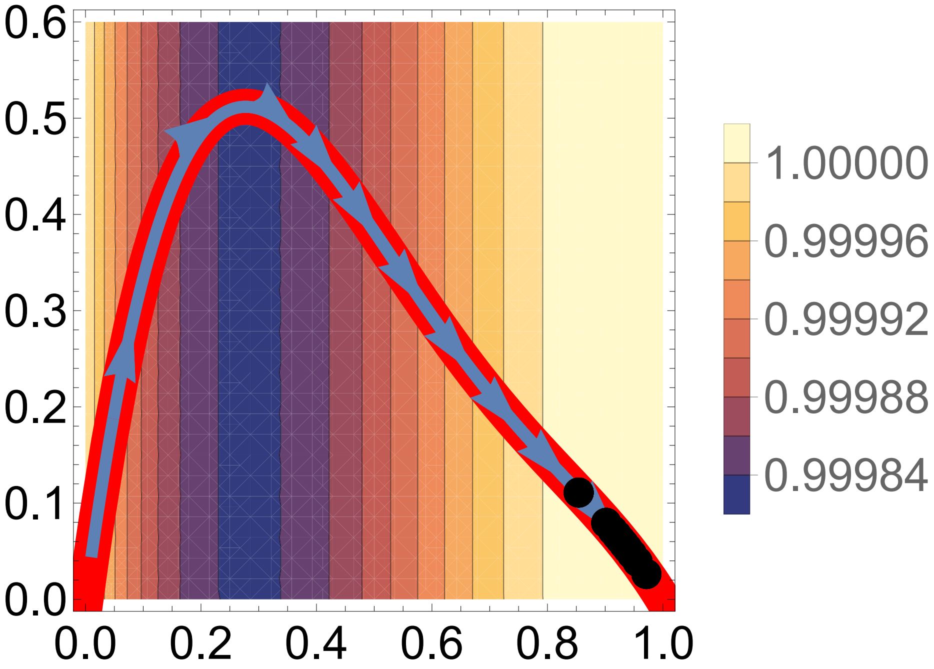
0.6
0.5
0.4
0.3
0.2
0.1
0.0
0.0
0.2
0.4
0.6
0.8
1.0
1.00000
0.99996
0.99992
0.99988
0.99984

#### Example 6 approximated evo-devo dynamics: one biallelic locus, explicit development of one phenotype, fitness depends on the phenotype at every age, weakening of selection with age

In[ ]:=
```
Clear["Global`*"]
```

In[ ]:=
```
(*This is Tx*)
Tx[xvar_] = -1/2 (-1 + xvar) xvar;

(*Developmental map*)
g[z_, xk_, xl_] := (1 + y[xk, xl]) z

y[xk_, xl_] := c (1 + d) (xk + xl) - 2 c d xk xl

z[xk_, xl_, a_] = (1 + y[xk, xl])^(a-1) z1;

(*These are the indicated derivatives*)

dwdx[xb_] := dzdx[xb].dwdz[z[xb, xb, 2], z[xb, xb, 3]];

dzdx[xb_] := {{dzdxa[xb, 1], dzdxa[xb, 2], dzdxa[xb, 3]}}

dzdxa[xb_, a_] :=
 If[a == 1, 0, If[a == 2, De[xb] z1, If[a > 2, (a - 1) De[xb] z1 (1 + y[xb, xb])^(a-2)]]]

(*This is D*)

De[xb_] = c (1 + d) - 2 c d xb

(*This is the MAG covariance*)

Lz[xb_] := 2 Tx[xb] × Transpose[dzdx[xb]].dzdx[xb]
```

Out[ ]=

$c\ (1 + d) - 2\ c\ d\ xb$

```
In[ ]:= (*This is fitness over Na life steps*)
        W[Na_, θ_, σ_] := 1/τ[Na] Sum[ϕ[a] × f[z[a]] + πf[a, Na] × s[z[a]], {a, 1, Na}]
        s[z_] := Exp[-(z - θ)^2/σ^2]
        f[z_] := Exp[-(z - θ)^2/σ^2]
        l[k_] := Product[s[zb[a]], {a, 1, k - 1}]
        πf[a_, Na_] := 1/s[zb[a]] Sum[l[k] × f[zb[k]], {k, a + 1, Na}]
        ϕ[a_] := l[a]
        τ[Na_] := Sum[a l[a] × f[zb[a]], {a, 1, Na}]

In[ ]:= Wfun[xvar_] = W[Na, θ, σ] /. {z[1] → z1, z[2] → z[xb, xb, 2], z[3] → z[xb, xb, 3]} /.
           {zb[1] → z1, zb[2] → z[xb, xb, 2], zb[3] → z[xb, xb, 3]} /. {xb → xvar};

In[ ]:= Simplify[Wfun[xb] /. z[a] → zb[a] /. Na → 3]
Out[ ]=
        1

In[ ]:= (*This is the selection gradient of the phenotype at age 1, 2, and 3*)

        dwdz1[z2b_, z3b_] =
          Simplify[D[W[3, θ, σ], z[1]] /. {z[1] → zb[1], z[2] → zb[2], z[3] → zb[3]}] /.
           {zb[1] → z1, zb[2] → z2b, zb[3] → z3b};

        dwdz2[z2b_, z3b_] =
          Simplify[D[W[3, θ, σ], z[2]] /. {z[1] → zb[1], z[2] → zb[2], z[3] → zb[3]}] /.
           {zb[1] → z1, zb[2] → z2b, zb[3] → z3b};

        dwdz3[z2b_, z3b_] =
          Simplify[D[W[3, θ, σ], z[3]] /. {z[1] → zb[1], z[2] → zb[2], z[3] → zb[3]}] /.
           {zb[1] → z1, zb[2] → z2b, zb[3] → z3b};

        (*This is the lifetime selection gradient*)

        dwdz[z2b_, z3b_] := {{dwdz1[z2b, z3b]}, {dwdz2[z2b, z3b]}, {dwdz3[z2b, z3b]}}
```

```
(*This runs the numerical solutions. In contrast to the exact
 method that takes 6 minutes, this takes about one second to run*)

Na = 3;
z1 = 0.1;

c = 1;
d = 2;

(*Numerical solutions*)

θ = 1;
σ = Sqrt[10];

(*This is fitness as a function of the phenotype at the final age*)
Wzfun[z3b_] = W[Na, θ, σ] /. {z[1] → z1, z[2] → z2b, z[3] → z3b} /.
   {zb[1] → z1, zb[2] → z[xb, xb, 2], zb[3] → z3b};

(*This is fitness as a function of allele frequency*)
Wfun[xvar_] = W[Na, θ, σ] /. {z[1] → z1, z[2] → z[xb, xb, 2], z[3] → z[xb, xb, 3]} /.
    {zb[1] → z1, zb[2] → z[xb, xb, 2], zb[3] → z[xb, xb, 3]} /. {xb → xvar};

tend = 1000;

x1binit = {0.01, 1, .1};

InitialConditions =
  Table[{x1bSol[0, i] = i}, {i, x1binit⟦1⟧, x1binit⟦2⟧, x1binit⟦3⟧}];

Table[Table[
   {x1bSol[t + 1, i] = x1bSol[t, i] + (Tx[x1bSol[t, i]] × dwdx[x1bSol[t, i]]⟦1, 1⟧)},
   {t, 0, tend}], {i, x1binit⟦1⟧, x1binit⟦2⟧, x1binit⟦3⟧}];
```

In[ ]:=
```
(*This plots allele frequency, mean phenotype,
MAG variance, and mean fitness over time*)

Plot1FileName =
  If[θ == 1.5 && d == 0, "p", If[θ == 1 && d == 0, "u", If[θ == 1 && d == 2, "z"]]];
Plot2FileName =
  If[θ == 1.5 && d == 0, "q", If[θ == 1 && d == 0, "v", If[θ == 1 && d == 2, "aa"]]];
Plot3FileName =
  If[θ == 1.5 && d == 0, "r", If[θ == 1 && d == 0, "w", If[θ == 1 && d == 2, "ab"]]];

yAxisLimitz[d_] := If[d ≥ 2, 1.3, 2]
yAxisLimitP[d_] := If[d ≥ 2, 0.2, .05]

lThickness = 0.02; (*Line thickness*)
Show[
 Table[ListLinePlot[{Table[x1bSol[t, i], {t, 0, tend}]}, PlotRange → {-0.1, 1.1},
   TicksStyle → Large, PlotStyle → Thickness[lThickness]],
  {i, x1binit[[1]], x1binit[[2]], x1binit[[3]]}]]
Export[StringJoin[ToString[NotebookDirectory[]],
   "Fig.S2", Plot1FileName, ".pdf"], %];
Show[Table[ListLinePlot[
   {Table[z[x1bSol[t, i], x1bSol[t, i], Na], {t, 0, tend}], Table[θ, {t, 0, tend}]},
   PlotRange → {-0.1, yAxisLimitz[d]}, TicksStyle → Large,
   PlotStyle → {Thickness[lThickness], {Magenta, Thickness[lThickness]}}],
  {i, x1binit[[1]], x1binit[[2]], x1binit[[3]]}]]
Export[StringJoin[ToString[NotebookDirectory[]],
   "Fig.S2", Plot2FileName, ".pdf"], %];
Show[Table[ListLinePlot[{Table[Lz[x1bSol[t, i]][[3, 3]], {t, 0, tend}]},
   PlotRange → {-0.1 yAxisLimitP[d], yAxisLimitP[d]}, TicksStyle → Large,
   PlotStyle → Thickness[lThickness]], {i, x1binit[[1]], x1binit[[2]], x1binit[[3]]}]]
Export[StringJoin[ToString[NotebookDirectory[]],
   "Fig.S2", Plot3FileName, ".pdf"], %];
Show[Table[ListLinePlot[{Table[Wfun[x1bSol[t, i]], {t, 0, tend}]},
   PlotRange → {-0.1, 1.1}, TicksStyle → Large,
   PlotStyle → Thickness[lThickness]], {i, x1binit[[1]], x1binit[[2]], x1binit[[3]]}]]
```

Out[ ]=

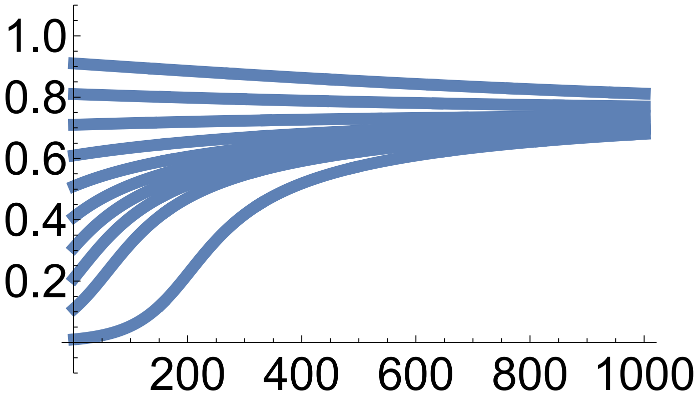

Out[ ]=

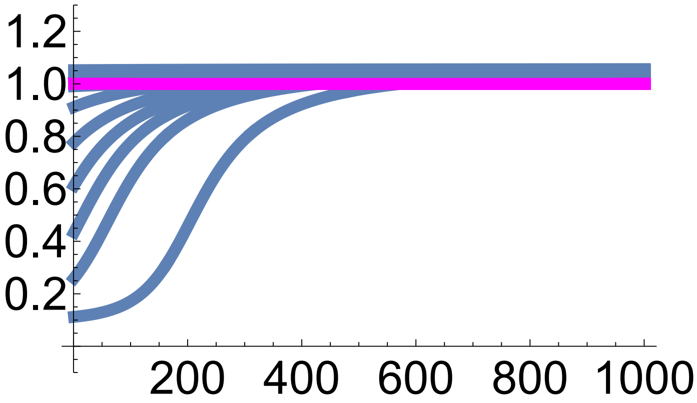

Out[ ]=

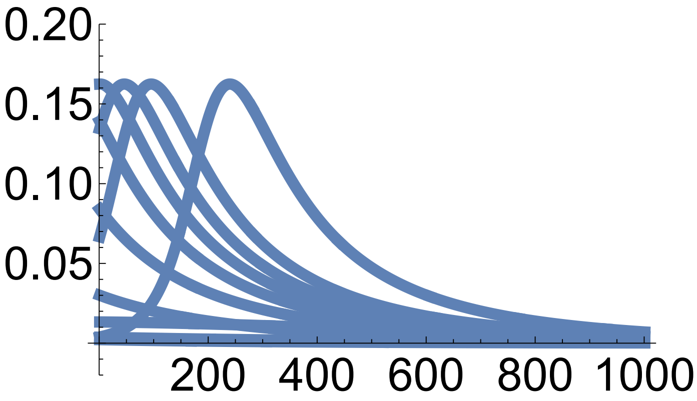

Out[ ]=

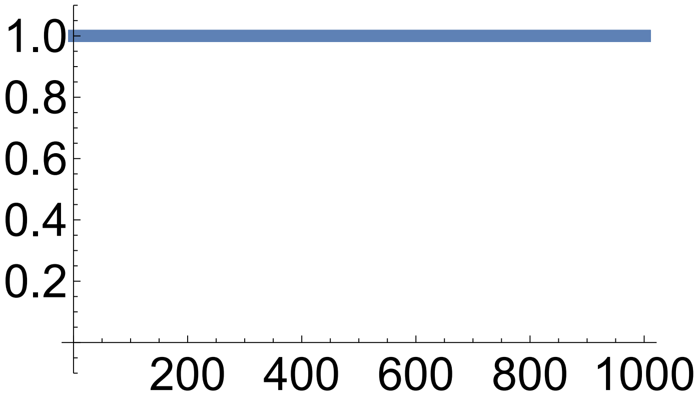

In[ ]:=

```
(*This plots mean phenotype and phenotype variance vs allele frequency*)

pointSize = 0.05;

Plot4FileName =
  If[θ == 1.5 && d == 0, "s", If[θ == 1 && d == 0, "x", If[θ == 1 && d == 2, "ac"]]];
Plot5FileName =
  If[θ == 1.5 && d == 0, "t", If[θ == 1 && d == 0, "y", If[θ == 1 && d == 2, "ad"]]];

endPoints = Table[{x1bSol[tend, i], z[x1bSol[tend, i], x1bSol[tend, i], Na]},
   {i, x1binit[[1]], x1binit[[2]], x1binit[[3]]},
   {j, x1binit[[1]], x1binit[[2]], x1binit[[3]]}];
```

```
p0 = Table[ListLinePlot[
     Table[{x1bSol[t, i], z[x1bSol[t, i], x1bSol[t, i], Na]}, {t, 0, tend}],
     PlotRange → {{0, 1}, {0, 2}}, PlotStyle → {Thickness[lThickness]}],
    {i, x1binit⟦1⟧, x1binit⟦2⟧, x1binit⟦3⟧}];

plot =  Show[ContourPlot[Wzfun[z3b], {xb, 0, 1}, {z3b, 0, yAxisLimitz[d]},
    FrameStyle → Large, PlotLegends → Automatic, LabelStyle → Large,
    Epilog → {PointSize[pointSize], Point[Flatten[endPoints, 1]]}],
   Plot[z[x1b, x1b, Na], {x1b, 0, 1}, PlotStyle → {Red, Thickness[3 lThickness]}],
   p0 /. Line[x_] :→ {Arrowheads[Table[{.1, .2}, {3}]], Arrow[x]}, ListLinePlot[
    Table[{xb, θ}, {xb, 0, 1}], PlotStyle → {Magenta, Thickness[lThickness]}]]

plot = Rasterize[plot, "Image", ImageResolution → 300];

Export[StringJoin[ToString[NotebookDirectory[]],
    "Fig.S2", Plot4FileName, ".pdf"], plot];

endPointsxP = Table[{x1bSol[tend, i], Lz[x1bSol[tend, i]]⟦3, 3⟧}, {i, x1binit⟦1⟧,
     x1binit⟦2⟧, x1binit⟦3⟧}, {j, x1binit⟦1⟧, x1binit⟦2⟧, x1binit⟦3⟧}];

p0 =
  Table[ListLinePlot[Table[{x1bSol[t, i], Lz[x1bSol[t, i]]⟦3, 3⟧}, {t, 0, tend}],
     PlotRange → {{0, 1}, {0, 1}}, PlotStyle → {Thickness[lThickness]}],
    {i, x1binit⟦1⟧, x1binit⟦2⟧, x1binit⟦3⟧}];

plot = Show[ContourPlot[Wfun[xb], {xb, 0, 1}, {P, 0, yAxisLimitP[d]},
    FrameStyle → Large, PlotLegends → Automatic, LabelStyle → Large,
    Epilog → {PointSize[pointSize], Point[Flatten[endPointsxP, 1]]}],
   Plot[Lz[x1b]⟦3, 3⟧, {x1b, 0, 1}, PlotStyle → {Red, Thickness[3 lThickness]}],
   p0 /. Line[x_] :→ {Arrowheads[Table[{.1, .2}, {3}]], Arrow[x]}]

plot = Rasterize[plot, "Image", ImageResolution → 300];

Export[StringJoin[ToString[NotebookDirectory[]],
    "Fig.S2", Plot5FileName, ".pdf"], plot];
```

Out[ ]=

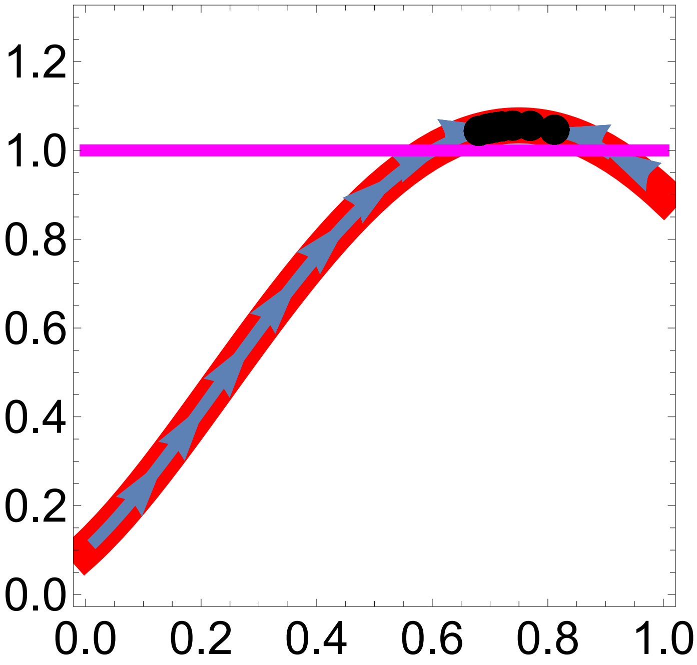
1.2
1.0
0.8
0.6
0.4
0.2
0.0
0.0
0.2
0.4
0.6
0.8
1.0

Out[ ]=

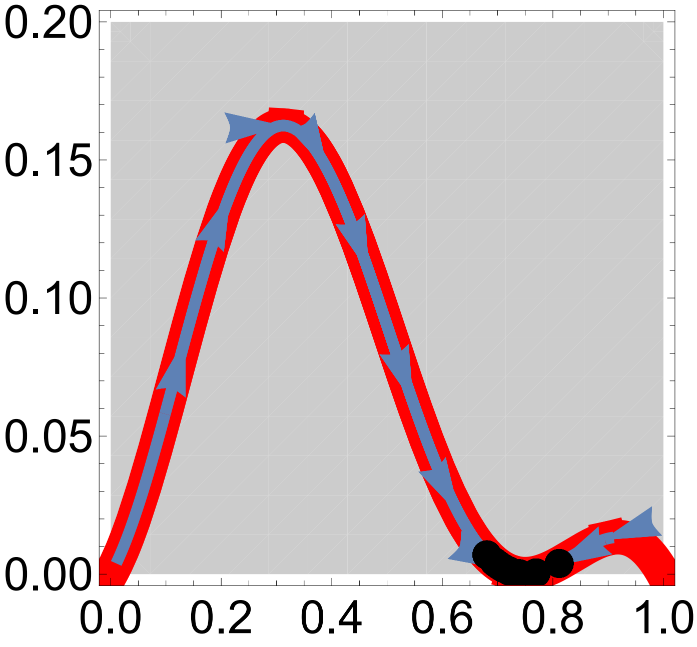
0.20
0.15
0.10
0.05
0.00
0.0
0.2
0.4
0.6
0.8
1.0
